# Quantitative Engineering and Investigation of Synthetic Sponge RNAs in *E. coli*

**DOI:** 10.64898/2026.05.19.726096

**Authors:** Scott B. Stacey, Kirill Sechkar, Marco Corrao, Harrison Steel, Antonis Papachristodoulou

## Abstract

Sponge RNAs (spRNAs) play an important regulatory role in bacterial small RNA (sRNA) networks, but their engineering and quantitative systems-level properties are unexplored. Here, we design, build, and quantitatively characterise synthetic spRNA-based gene circuits in *E. coli*. We establish multiple design strategies for synthetic spRNAs, engineering the first synthetic spRNAs. We show that these synthetic spRNAs can reversibly de-repress sRNA-regulated gene expression, demonstrate tuneable control of gene expression, and extend these designs to multi-target regulation. Through the use of time-resolved continuous-culture characterisation in Chi.Bio together with absolute fluorescent protein quantification, we generated a quantitative dynamical dataset for model fitting and mechanistic analysis. Sequential model development showed that recapitulating the observed circuit dynamics required incorporation of Hfq-mediated resource competition, often overlooked in models of sRNA-based synthetic gene circuits. The extended model captured promoter, sRNA, and sponge circuit behaviour and was used to investigate quantitative properties of spRNA-mediated regulation, the first such quantitative investigation of spRNA-based regulation. Model-based quantitative investigations further suggest that spRNAs can tune response functions, modulate thresholds and leakiness, alter response times, improve disturbance rejection in some regimes, increase effective specificity, and buffer regulatory output against sRNA mutation. Together, these results establish synthetic spRNAs as a new post-transcriptional tool for bacterial synthetic biology and provide a quantitative framework for understanding natural and engineered spRNA-mediated regulation.

## Introduction

Engineered biological systems hold great promise both for advancing understanding of natural biological systems and for enabling practical solutions in areas including healthcare, biomanufacturing, agriculture, and environmental remediation ^1,2^. A key challenge in synthetic biology, underpinning many of these outcomes, is the engineering of synthetic gene circuits with predictable, tuneable, and robust computational behaviour across changing conditions and contexts ^3–5^. Significant progress has been made in this area, including oscillators ^6,7^, switches ^8^, logic gates ^9^, and feedback circuits ^10^, typically based on transcription-factor regulation ^11,12^. In parallel, RNA-based regulation has also been used to build synthetic gene circuits ^13–26^, including those based on bacterial small RNAs (sRNAs) ^27–37^.

Small RNAs are a ubiquitous and biologically important form of gene regulation in bacteria ^38–40^. Bacterial sRNAs are typically short ncRNAs (50–200 nt) that generally, though not exclusively ^41^, regulate gene expression post-transcriptionally by base-pairing mRNAs, and are commonly grouped into *trans*- and *cis*-acting classes ^38^. In many bacteria, including *E. coli*, many known sRNAs rely on the chaperone Hfq for regulatory function ^42^, and Hfq-binding also contributes to their unusual stability among bacterial RNAs ^43,44^. Bacterial sRNAs typically operate by base pairing with target mRNAs (Fig. 1A), through which they can produce diverse regulatory outcomes (Fig. 1B) ^45^. These include binding at or near the mRNA RBS to occlude ribosome binding and reduce translation ^46^; altering mRNA structure to improve ribosome access and increase translation ^47^; promoting mRNA, and often sRNA, degradation ^48^; or blocking mRNA degradation and increasing transcript stability ^49^. Many endogenous sRNAs use relatively short binding regions and therefore have promiscuous target binding ^38^, yet they often regulate key cellular processes including central metabolism ^50^, stress responses ^51^, and biofilm formation ^52^.

**Figure 1:**
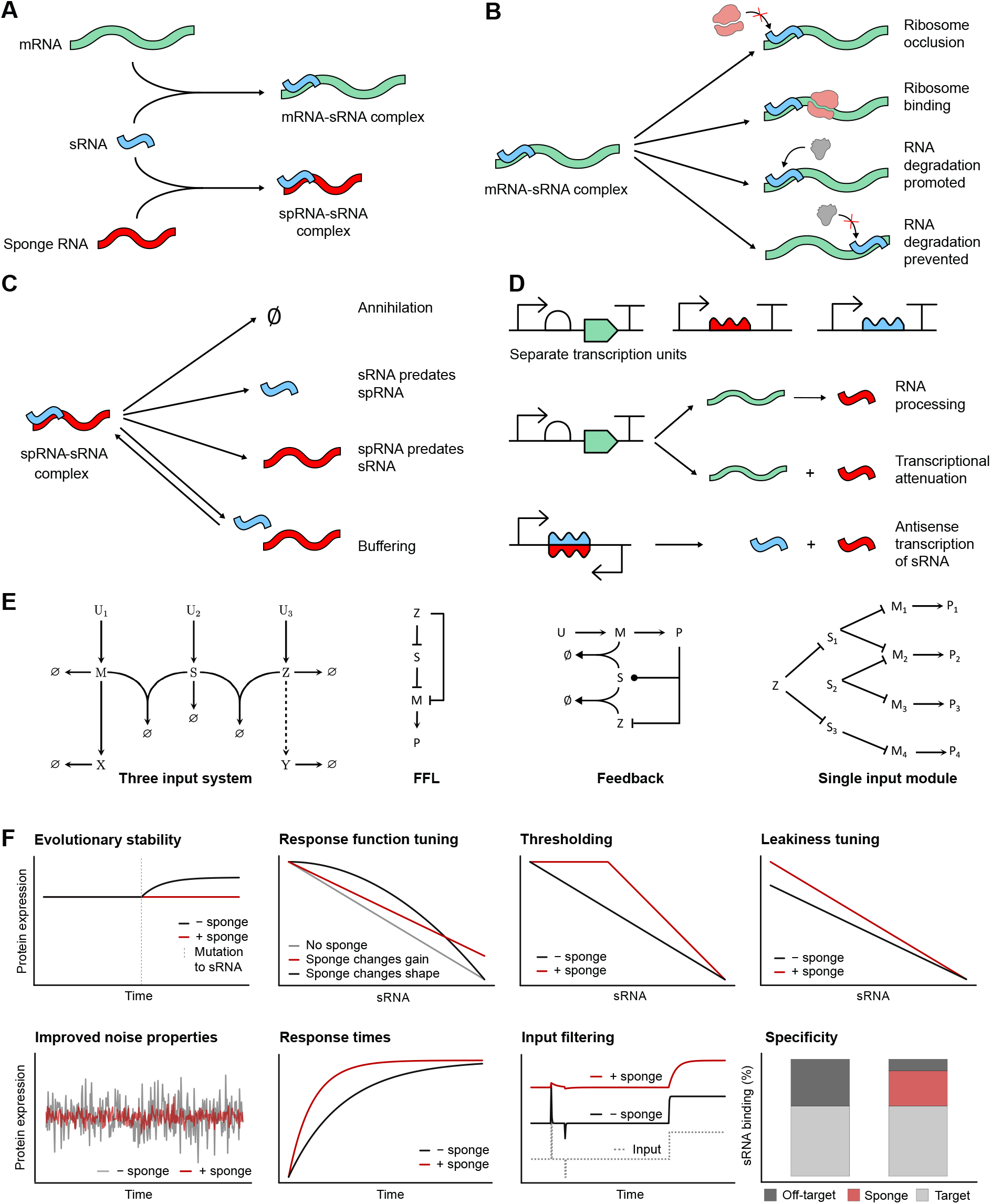
Concepts and hypotheses for sponge RNAs: **(A)** A sponge RNA is an alternative binding partner for an sRNA in addition to its mRNA target. **(B)** Upon sRNA binding to an mRNA target diverse regulatory activities can occur including translational repression or activation and/or mRNA protection or degradation. **(C)** Similarly it is thought that sponge RNAs could have diverse activities upon binding an sRNA including mutual annihilation, the sRNA predating the sponge RNA, the sponge RNA predating the sRNA, and true sponging or buffering. **(D)** In addition to diverse regulatory mechanisms sponge RNAs have diverse mechanisms of production. Sponge RNAs and sRNAs can be produced as individual transcription units or by processing mechanisms including premature transcriptional termination or post-transcriptional processing. It has also been hypothesised that a sponge RNA could be produced through antisense RNA transcription of an sRNA in the same way that sRNAs are sometimes produced through antisense transcription of an mRNA. **(E)** Sponge RNAs (*Z*) regulating sRNAs (*S*) may be involved in a variety of natural gene regulatory networks or synthetic gene circuits including multi-input systems, feedforward loops, feedback loops, or single-input larger regulons ultimately regulating target genes (*X, Y*, or *P*_*n*_). **(F)** Little is known about the quantitative impact of sponge RNAs on gene regulatory networks or synthetic gene circuits but plausible hypotheses include coupled regulation, evolutionary stability, response function tuning, faster response times, thresholding, leakiness tuning, improved (or enhanced) noise properties, input filtering, and improved specificity.

Quantitative studies of sRNA regulation have shown that sRNAs can produce characteristic threshold-linear responses ^53,54^. Other work has investigated noise properties ^55,56^ and response times ^57^. Small RNAs offer potential advantages in Engineering Biology, including small genetic footprint, easy programmability, and widely available computational design tools ^58–60^. These properties have motivated increasing use of sRNAs in synthetic systems, including metabolic engineering ^28,29^, control systems ^21,32^, and multiplexed regulation ^61^. Much of this work has focussed on the scaffold MicC ^28,32^, although there is growing work on engineering diverse scaffolds ^62–66^. Despite this progress, practical use of sRNA-based regulation in gene circuits faces challenges, including limited scaffold diversity, difficulty fine-tuning, and limited tools for switching off an sRNA regulator ^62,65,67^.

Sponge RNAs (spRNAs) are a recently uncovered regulator in endogenous sRNA-based gene regulatory networks, and have also been termed RNA predators or trap RNAs ^67–69^. Sponge RNAs are transcripts that provide alternative binding partners for sRNAs (Fig. 1A), effectively acting as “sRNAs for sRNAs” and introducing additional regulatory layers in sRNA networks. They sequester regulatory sRNAs through sequence-specific binding, reducing the effective amount of free sRNA available to regulate target mRNAs. Regulatory outcomes following sRNA–spRNA binding appear diverse (Fig. 1C), and in some cases remain unclear ^69,70^, but likely include mutual co-degradation (annihilation) ^71^, predatory/catalytic interactions (only one RNA in the pair is degraded) ^72^, and stable pair formation or buffering ^73^. Diverse modes of spRNA production have also been observed or proposed (Fig. 1D) ^69^, and spRNAs participate in a wide range of network architectures (Fig. 1E), including simple multi-input/open-loop systems ^71,74,75^, feedforward loops ^76^, feedback loops ^77^, and larger regulons ^72,78^.

While work has increasingly focused on discovering endogenous spRNAs and understanding their molecular interactions^69^, there has been little quantitative systems-level investigation of spRNA-mediated regulation and no prior engineering of synthetic spRNAs. Although systems-level roles for spRNAs have been proposed ^69,70^, these have not been extensively tested quantitatively or theoretically. Plausible functions of spRNA-mediated regulation (Fig. 1F) include enhanced evolutionary stability, response-function tuning, thresholding or mitigation of promoter leakiness, noise attenuation, altered response times, input filtering, increased specificity in the presence of off-targets, temporal control, and coupled or multi-target regulation. These behaviours may contribute both to natural spRNA-based regulation and to useful design of synthetic gene circuits.

A further challenge has been the quantitative investigation of RNA-based circuit dynamics. Many previous characterisations of sRNA-based circuits have relied on batch-culture microplate readers or low-time-resolution fluorescence, proteomic, or transcriptomic measurements ^32,71,79,80^. While such datasets are valuable for discovering regulatory RNAs, identifying targets, and determining overall regulatory outcomes, they are not effective for analysing gene-network dynamics. Infrequent sampling or steady-state-only measuremsaents can obscure important dynamical features ^81^. Batch culture introduces changing growth conditions that confound inference of circuit parameters and behaviour ^82,83^, particularly because growth rate affects both protein dilution and gene expression ^84^. Measurements in arbitrary fluorescence units limit comparability across experiments and reduce interpretability of fitted model parameters. Finally, simple one-sRNA synthetic circuits may obscure the impact of Hfq-mediated resource competition on regulatory behaviour, despite its well-established importance in natural sRNA regulation ^80,85–87^. Recent advances in continuous-culture characterisation platforms and absolute quantification methods make rigorous system identification of synthetic gene circuits in physically meaningful units increasingly feasible ^82,88–90^.

In this paper we develop and quantitatively investigate synthetic spRNAs in *E. coli*. We first design and characterise synthetic sRNAs and spRNAs, including synthetic spRNAs and sRNAs based on a natural mRNA-sRNA–spRNA system (ChiP–ChiX–ChbBC ^43,71,75,77,80,91^) and alternative spRNA design strategies for MicC-derived synthetic sRNAs. Using time-resolved characterisation in the Chi.Bio continuous-culture platform ^82^ with absolute fluorescent protein quantification ^88^, we generate a quantitative dynamical dataset for model fitting and mechanistic analysis, the first such dataset for sRNA or spRNA regulation. We develop a series of mathematical models and show that recapitulating the observed circuit dynamics requires modelling resource competition. Using our fitted model, we investigate quantitative properties of spRNA-mediated regulation, including tuneability of response functions, threshold and leakiness modulation, response-time effects, disturbance rejection, specificity in the presence of off-target binding, and robustness to sRNA mutations.

Together, our results expand the synthetic biology toolbox by engineering the first synthetic spRNAs and synthetic ChiX-based sRNAs, while providing an experimental framework for spRNA engineering. Our work also advances quantitative characterisation of synthetic gene circuits by combining dynamic measurements of circuit outputs and cell growth rates, continuous culturing in a bioreactor, and absolute protein quantification. Finally, our quantitative investigation of spRNAs provides a first step towards understanding spRNA systems biology, while suggesting roles for spRNAs in synthetic gene circuits.

## Materials and methods

### Bacterial strains and growth conditions

Marionette-Clo *E. coli* strain derived from *E. coli* DH10B ^92^ was used for all cloning, plasmid propagation, and characterisation experiments. Cells were grown in LB media for cloning, plasmid propagation, and protein expression. For circuit characterisation, cells were grown in M9 media supplemented with 0.4% glucose (w/v), 2 mm MgSO_4_, 100 µm CaCl_2_, 0.2% Casamino acids (w/v), and 1× trace elements (working concentrations: 134.32 µm EDTA Na_2_ · s2 H_2_O; 30.8 µm FeCl_3_; 6.16 µm ZnCl_2_; 0.76 µm CuCl_2_ · 2 H_2_O; 0.42 µm CoCl_2_; 1.62 µm H_3_BO_3_; and 81 nm MnCl_2_ · 4 H_2_O). All cultures were grown at 37 °C except protein expression experiments, which were at 30 °C. Standard techniques were used for plasmid propagation, preparation, transformation, and storage. Chloramphenicol, carbenicillin, and kanamycin were used at working concentrations of 5 µg mL^*−*1^, 100 µg mL^*−*1^, and 50 µg mL^*−*1^, respectively.

### Regulatory RNA design

Regulatory RNAs (sRNAs and spRNAs) were designed *in silico* by reprogramming binding regions to target desired RNAs and screening candidate sequences prior to experimental validation. IntaRNA^60,93^ was used to predict on-target and off-target interactions. Candidates were prioritised if they bound the intended site without mismatches, had predicted binding energies of at least Δ*G* ≤ − 20, and showed more favourable predicted binding to the intended target than to the most likely off-target. Secondary structures were predicted using Vienna RNAFold ^59^ or mfold ^58^ and visualised with RNACanvas ^94^ for comparison with appropriate natural RNA structures. As these tools provide only approximate guidance, multiple candidate regulatory RNAs were generally taken forward for experimental testing. Specific design strategies, candidate sets, and sequences are described in the Results section and Supplementary Materials.

### Molecular cloning

Plasmids were assembled using standard Golden Gate cloning with the EcoFlex MoClo kit ^95^, with transcription units assembled into level 1 plasmids and multiple level 1 plasmids cloned together to form synthetic gene circuits. Plasmid backbones were taken from the EcoFlex kit ^95^, with some modified (replacing chloramphenicol with kanamycin) to ensure compatibility with the Marionette strain ^92^. The putative GFPmut3 sequence was taken from the EcoFlex kit ^95^. All other DNA parts (including designed regulatory RNAs) were synthesised as EcoFlex-compatible gene fragments by Integrated DNA Technologies Inc. or Twist Biosciences. Some plasmids were engineered using NEB’s Q5 Site-Directed Mutagenesis Kit following the standard protocol and using the NEBaseChanger tool (New England Biolabs) to design primers. A list of DNA parts, primers, and plasmids used or assembled in this work is provided in the Supplementary Materials. Assembled plasmids were sequence-verified by full-plasmid Oxford Nanopore sequencing performed by Plasmidsaurus or PlasmidsNG with custom analysis and annotation.

### Microplate reader characterisation

Microplate reader assays employed a Tecan Spark instrument in 96-well plate format. Cells were transformed with a plasmid of interest; a single colony was selected to form a biological replicate and grown overnight in LB media with appropriate antibiotic at 37 °C shaking at 250 rpm. On the day of the assay, overnight cultures were pelleted, washed with PBS, and resuspended in fresh M9 media. Wells were filled with 199 µL M9 at appropriate inducer and antibiotic concentrations, to which 1 µL of culture was added. Blank wells (M9 only) and non-fluorescent negative controls were included. Plates were run for 24 hours at 37 °C with orbital shaking at 90 rpm (3 mm amplitude). GFPmut3 fluorescence was measured with 470 nm excitation and 515 nm emission; mScarlet-I with 570 nm excitation and 615 nm emission; and optical density at 600 nm. Red fluorescence, green fluorescence, and OD were generally recorded every 15 minutes.

### Chi.Bio characterisation

Most characterisation experiments were carried out in continuous culture using the Chi.Bio platform (Supplementary Fig. S1A) ^82^. Cells were transformed with plasmid(s); a single colony was grown overnight in LB with appropriate antibiotic at 37 °C shaking at 250 rpm. On the day of an experiment, 50 µL of overnight culture was diluted into 5 mL fresh M9 with antibiotics and grown for 2–3 hours at 37 °C shaking at 250 rpm. During pre-growth, Chi.Bio reactors were set up: autoclaved vials were cleaned with ethanol and dried; and media and waste tubing flushed with 70% ethanol; each vial was pumped to ∼20 mL M9; loaded into the reactor; and temperature control enabled at 37 °C with stirring at 0.5. After 15 minutes, OD was blanked in the Chi.Bio GUI and OD turbidostat control started (setpoint 0.5) with OD dithering enabled for growth rate calculation; temperature control remained at 37 °C and stirring at 0.5. Fluorescence settings were: GFPmut3, 457 nm excitation and 510 nm emission; mScarlet-I, either 523 nm excitation and 583 nm emission or 595 nm excitation and 670 nm emission. Experiments ran for a further 15 minutes to confirm OD blanking stability, after which any further blanking was performed. Vials were then inoculated with 200 µL culture by pipetting or syringe. Experiments were then run until required user intervention (e.g. media change, inducer step-change, or sampling).

Inducer step-changes were achieved by temporarily disabling OD control, emptying the input media tube into waste, swapping the media bottle at concentration *x* for one at concentration *y* where *y > x*, washing pumps through with new media into waste, adding 1 mL inducer solution (in M9) to each vial undergoing step-change, reconnecting pumps, and switching OD control back on. For a vial with *∼*20 mL culture volume, the inducer concentration required for the 1 mL addition is *z* = 21*y −* 20*x*. Induction timepoints were recorded or stored as metadata in the Chi.Bio GUI. Further description of Chi.Bio set-up is provided in ^82,90^.

### Data processing

Chi.Bio fluorescence data were processed by calculating mean fluorescence intensity in the 30 minutes prior to fluorescent protein induction to estimate background offset, and subtracting this from all fluorescence values. For experiments without absolute quantification, the mean background-corrected fluorescence in the final 30 minutes after GFPmut3 induction but prior to regulatory RNA induction was calculated as a corrected steady-state value; fluorescence was normalised to this value such that initial steady-state induced signal was 1. Finally, a 21-point symmetrical moving average of the normalised background-corrected fluorescence was applied to smooth shot-to-shot variability. When absolute quantification was used, processing was applied without normalisation to relative fluorescence.

When mScarlet-I was measured with 523 nm excitation and 583 nm emission in the presence of GFPmut3 expression, an additional correction was applied because GFPmut3 is detected in this channel. The ratio of GFPmut3 fluorescence in the green channel to that in the red channel was calculated, and the mean of the final 30 minutes of this ratio (when GFPmut3 is expressed but mScarlet-I is not) was used to correct mScarlet-I: mScarlet-I Fluorescence = Red Fluorescence −Red:Green Ratio × Green Fluorescence. This correction was not required when mScarlet-I was measured with 595 nm excitation and 670 nm emission, though this resulted in a noisier signal.

Microplate reader fluorescence intensity was processed by normalisation to blanks, negative controls, and OD:

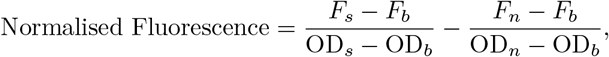

where *F*_*s*_ is sample fluorescence, *F*_*b*_ is mean fluorescence of media-only blanks, OD_*s*_ is sample OD, OD_*b*_ is mean OD of media-only blanks, *F*_*n*_ is mean fluorescence of non-fluorescent negative controls, and OD_*n*_ is mean OD of negative controls.

To generate promoter inducer response curves, data with OD*<* 0.2 or OD*>* 0.35 were excluded. Mean normalised fluorescence intensity was then taken as an approximation of steady-state fluorescence for each promoter and inducer concentration.

### Growth rate estimation

Growth rate estimation was performed from Chi.Bio OD traces ^82^ using the GOFFREDOpy package ^96^. OD data were pre-processed by discarding the first 5 OD measurements (≈5 minutes) following each inducer addition, and removing portions corresponding to known temporary hardware failures. For estimation, the standard deviation of the measurement noise was set to 0.005 and filter adaptivity to 0.15 (see ^96^ for more details). The default confidence level of 1% was used for outlier detection and rejection. Consistency of growth rate estimates was verified by testing for the absence of significant autocorrelation between consecutive innovations up to 5 lags. Growth rate estimates were finally smoothed with a 21-point symmetrical moving average prior to model fitting.

### Absolute quantification

Absolute quantification of fluorescent protein concentrations in Chi.Bio experiments assisted quantitative model fitting and enabled meaningful comparison between experiments and proteins. We adapted the FPCountR protocol^88,89^ to the Chi.Bio platform ^82^ by temporal sampling and plate-reader comparison.

Briefly (Supplementary Fig. S31), fluorescent proteins were overexpressed overnight; cells lysed and processed to yield a concentrated complex calibrant. Serial dilutions of calibrant were prepared and the ECMax assay performed, followed by the fluorescence assay ^88,89^. The FPCountR R package ^97^ was used to relate fluorescence intensity to moles of fluorescent protein for the microplate reader and its settings. Chi.Bio experiments were sampled around induction events and measured in the same microplate reader with the same settings. FPCountR^97^ was then used to convert these measurements to molar intracellular protein concentrations. For each Chi.Bio reactor, data across experiments were pooled and linear models of the form *y* = *mx* + *c* were fitted in R ^98^ describing conversion from processed Chi.Bio fluorescence to molar concentration. These were then used to convert Chi.Bio fluorescence timecourses into intracellular protein concentration timecourses. Details of our application of FPCountR ^88^, including a full protocol and troubleshooting tips, are provided in ^90^.

### Mathematical modelling

Parameter estimation was carried out in MATLAB. To reduce degrees of freedom and improve identifiability, parameters were shared across constructs where this was biologically justified: promoter parameters were shared across all circuits containing a given promoter, translation rates were shared across transcription units with the same RBS, and RNA-specific parameters were shared for a given regulatory RNA. This assumes that circuits were characterised under closely matched conditions and that the corresponding parts behave approximately modularly, which was supported here by Chi.Bio characterisation ^82,90^ and standardised transcription unit design (Supplementary Fig. S2). Nevertheless, conditions were never perfectly identical between experiments, and sequence-context effects may violate strict modularity. Despite these limitations, shared-parameter fitting allowed promoter and translation parameters to be estimated in relative isolation, reducing the extent to which they might otherwise be biased by downstream architecture choices in the RNA models.

Initial parameter estimates were bounded using *a priori* knowledge and updated in subsequent optimisation runs. Linear inequality constraints restricted promoter dynamic range between 50 and 2000. The optimisation used a three-level cost function. For each dataset, we calculated a normalised mean squared error,

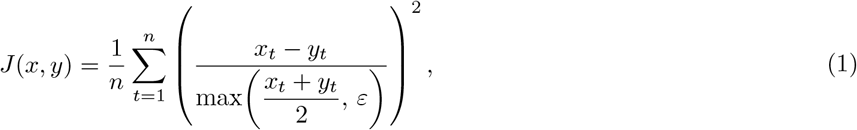

where *x*_*t*_ and *y*_*t*_ are the experimental and model values at timepoint *t, n* is the number of paired observations, and *ε* is a positive floor term chosen adaptively from the experimental data to avoid division by very small values. Experimental trajectories were resampled onto the model time grid prior to cost evaluation. Second, where multiple replicates existed for an objective, their costs were averaged as

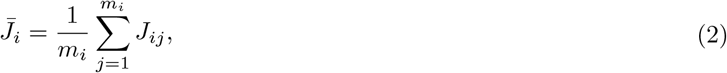

where *J*_*ij*_ is the cost for replicate *j* within objective *i*, and *m*_*i*_ is the number of replicates for that objective. The objectives included GFPmut3 fluorescence, mScarlet-I fluorescence where present, and, when fitting the growth feedback model, cell growth rate. Finally, the overall objective was defined as a weighted average over objective-specific mean costs,

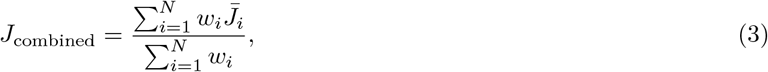

where *w*_*i*_ is the weighting for objective *i*, and *N* is the total number of objectives.

Because the objective function was non-convex, parameter estimation used a global–local optimisation strategy in which `GlobalSearch` was coupled to `fmincon`. Six optimisation runs were performed in parallel (`parfor`) using different initial guesses, and the lowest-cost solution was retained. Promoter-characterisation data were initially fit separately to reduce bias from downstream regulatory-RNA assumptions; in fitting the final model, promoter and RNA-circuit parameters were estimated jointly. Model fitting used experimentally estimated time-varying growth rates and inducer concentrations as exogenous inputs to the ODE system. Dataset-specific multiplicative scaling factors were also included for the RNA-circuit datasets to link model-predicted fluorescent protein concentrations to measured outputs and to absorb differences in scale not captured by the mechanistic parameters (see Supplementary Note S7.2). To investigate possible burden-mediated growth feedback, we also fit a variant of the final RNA-circuit model in which growth rate was predicted from modelled fluorescent protein expression and used as the dilution rate in the ODE system, rather than supplied directly from the experimental growth-rate measurements. Full details are provided in Supplementary Note S8.

Fit quality was not judged using a single universal metric, because datasets differed in the relative importance of particular features. For example, it was more important that a model captured observed rises and decreases in expression following sRNA or spRNA induction than that it marginally improved a global numerical goodness-of-fit metric. Accordingly, the weighted cost function was used to optimise parameters within a given model, while practical goodness-of-fit was assessed by the extent to which simulations reproduced the key qualitative and quantitative features of the data. These challenges further motivate the use of Chi.Bio-based characterisation, in which circuit dynamics are readily observable under approximately constant culture conditions. Throughout this work, references to model fit quality therefore generally refer to a model’s ability to capture both the qualitative and quantitative behaviour of the data.

All simulations were carried out by numerical integration using MATLAB’s ode15s ODE solver, both during parameter estimation and in subsequent simulations of spRNA behaviour. Steady states were determined numerically by simulating until convergence and taking the final simulated values as the steady state.

## Results

### Circuit conceptualisation and part selection

To test engineered synthetic spRNAs, we conceptualised a simple three-input circuit of the form shown in Fig. 1E. This architecture was chosen as a minimal and experimentally tractable system that still captures the core regulatory logic of sRNA-mediated knockdown and spRNA-mediated reversal, while allowing sequential induction of each regulatory RNA and direct observation of their effects on circuit dynamics. It is also broadly inspired by the natural ChiP–ChiX–ChbBC system, while remaining simpler than the endogenous architecture. The circuit comprises an mRNA, *M*, encoding a protein, *X*; *M* can be bound by a synthetic sRNA, *S*, designed to knock down *X* expression; *S* can in turn be bound by a synthetic spRNA, *Z*, sequestering *S* and reversing its regulatory effect. A second fluorescent protein, *Y*, may be co-transcribed with *Z* to enable online measurement of spRNA expression. To actualise this circuit, up to two orthogonal fluorescent proteins (*X* and *Y*) and three chemically inducible promoters (inducers *U*_1_, *U*_2_, and *U*_3_) were required.

After conceptualising the circuit, three promoters were chosen, validated, and characterised. Orthogonal promoters and fluorescent proteins were selected to minimise crosstalk between transcriptional inputs and reporter outputs, and strong terminators were used to reduce transcriptional readthrough between transcription units. To isolate promoter function from the context of the transcription units in which they would operate, each promoter was placed in a standardised transcription unit comprising the same plasmid backbone, ribozyme insulator, RBS, CDS, and terminator, differing only in the promoter. Three such transcription units (Supplementary Fig. S2) expressing GFPmut3 under promoters from the Marionette collection ^92^ (P_VanCC_, P_LuxB_, and P_Tac_) were assembled into the EcoFlex pTU1-A-LacZ backbone ^95^ to form pSS-01-001, pSS-01-002, and pSS-01-003, respectively. These plasmids were transformed into the Marionette-Clo strain ^92^ and characterised in microplate-reader batch-culture experiments (Supplementary Fig. S3). Each transcription unit showed desirable input-tuneable GFPmut3 expression. In addition, GFPmut3 and the second fluorescent protein, mScarlet-I, were tested in the Chi.Bio platform, validating observable fluorescence in that setup (Supplementary Fig. S4).

### Design and characterisation of ChiX-derived synthetic sRNAs

#### Design and initial functional validation

We next designed ChiX-derived synthetic sRNAs. The design process is summarised in Fig. 2A. Briefly, the binding region of the natural sRNA was extended, reprogrammed, and validated *in silico*. Using this approach, we designed ten synthetic sRNAs (Supplementary Note S3.1) targeting the GFPmut3 mRNA produced from pSS-01-001 (Supplementary Fig. S2A). Target sites of varying lengths (18–25 nucleotides) were selected within a region spanning 35 nucleotides upstream and downstream of the GFPmut3 RBS, generating variants targeting sequences downstream, upstream, or overlapping the RBS. Each synthetic sRNA was then assembled into a simple two-input circuit comprising inducible GFPmut3 expression and inducible sRNA expression (Fig. 2C; Supplementary Note S3.1).

**Figure 2:**
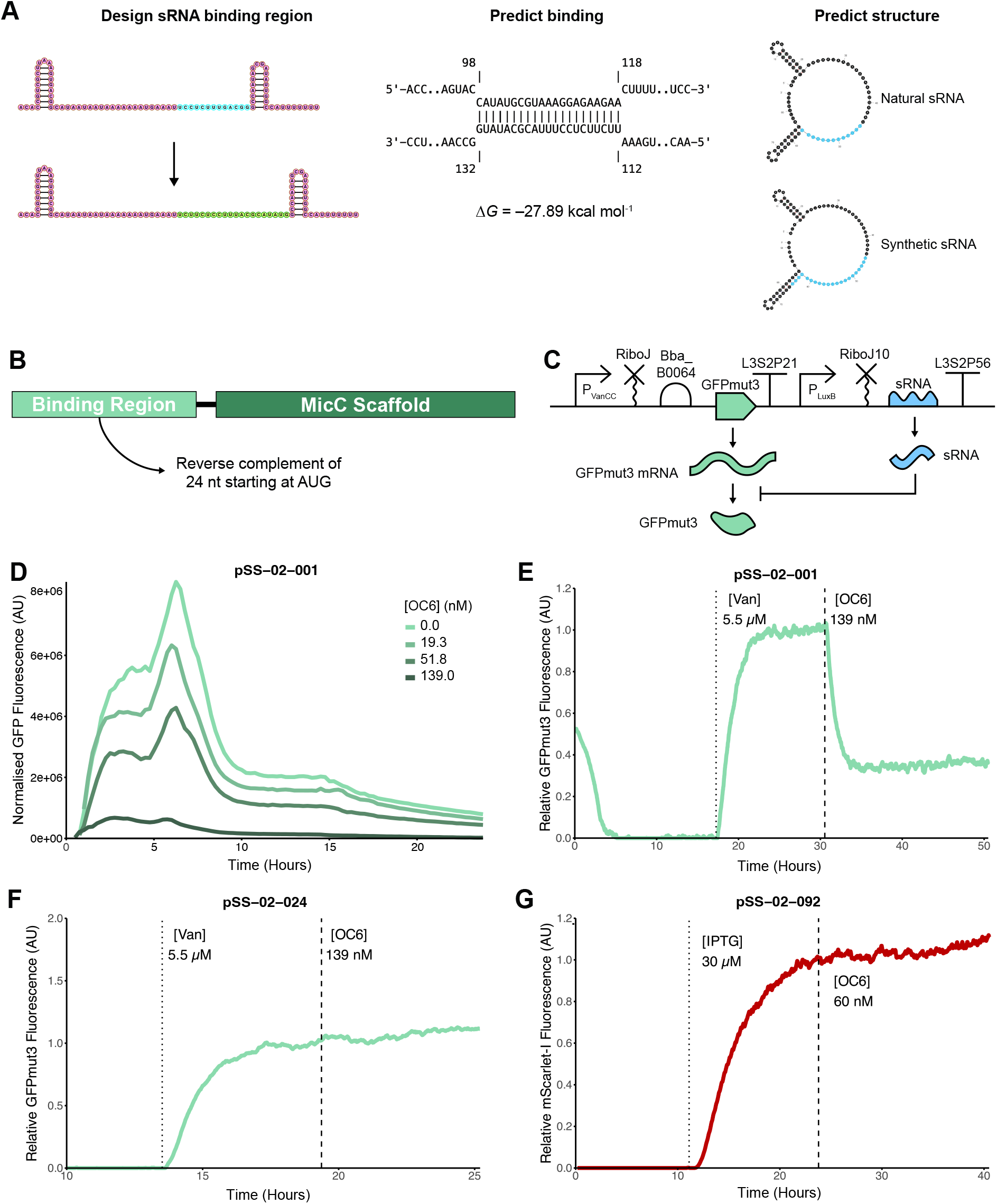
Design and characterisation of synthetic small RNAs: **(A)** To engineer synthetic ChiX-derived sRNAs, the natural ChiX sequence was used as a template and its binding region was lengthened and reprogrammed to target an mRNA of interest near its RBS. Designed sRNAs were rule-of-thumb validated *in silico* using binding predictions from IntaRNA ^60^ and structure prediction with mfold ^58^. **(B)** MicC-derived synthetic sRNAs were designed using the established approach ^28,29^, retaining the standard MicC scaffold and placing upstream of it the reverse complement of the first 24 nucleotides of the target mRNA beginning at the start codon. **(C)** SBOL Visual ^99^ schematic of the two-transcription-unit synthetic gene circuit used for synthetic sRNA validation. The first transcription unit expresses GFPmut3 in response to vanillic acid. The second transcription unit expresses a variant sRNA in response to OC6. **(D)** Representative example of microplate reader validation of synthetic sRNA for Syn-ChiX-20D in pSS-02-001. Darker green is indicative of higher sRNA induction and leads to lower normalised GFPmut3 fluorescence. **(E)** Representative characterisation of a functional synthetic sRNA (Syn-ChiX-20D in pSS-02-001) in the Chi.Bio platform. GFPmut3 rises to a steady-state fluorescence after induction, followed by induction of the synthetic sRNA, which leads to a *<* 0.4 fold-change. **(F)** The same experiment carried out with a negative control synthetic sRNA with a randomised binding region (Syn-ChiX-20NC) in pSS-02-024. As before GFPmut3 rises to a steady state but upon sRNA induction there is little change in expression. **(G)** To validate that Syn-ChiX-20D was orthogonal to mScarlet-I, it was tested in pSS-02-092; induction of Syn-ChiX-20D caused little change in mScarlet-I expression, as desired.

These synthetic gene circuits were first characterised in a microplate reader to validate sRNA functionality (Supplementary Fig. S17). Across all ten circuits, GFPmut3 was induced with 18 µm vanillic acid, with varying concentrations of OC6 used to induce the sRNAs. A representative example of desired sRNA behaviour is shown for Syn-ChiX-20D in pSS-02-001 (Fig. 2D). At the five-hour timepoint, fold-change in GFPmut3 expression for maximally induced sRNA ranged from 0.0438 for Syn-ChiX-18D (Supplementary Fig. S17A) to 0.936 for Syn-ChiX-22U (Supplementary Fig. S17F). Notably, the mean fold-change at five hours for sRNAs targeting upstream of the RBS was 0.822, compared to 0.105 for those targeting downstream; none of the upstream-targeting sRNAs were stronger repressors than the downstream-targeting sRNAs. This pattern may be indicative of design rules for ChiX-based synthetic sRNAs.

#### Continuous-culture characterisation

Two of the most-repressing sRNAs, Syn-ChiX-18D and Syn-ChiX-20D, were chosen for characterisation in continuous culture using the Chi.Bio platform ^82^. Cultures were maintained in exponential phase (Supplementary Fig. S1B) and exposed to sequential step-changes inducing GFPmut3 expression (5.5 µm vanillic acid) followed by sRNA expression (139 nm OC6). In pSS-02-001 (Syn-ChiX-20D), fold-changes of approximately 0.2–0.4 were achieved across several replicates (Fig. 2E and Supplementary Fig. S18A). In pSS-02-002 (Syn-ChiX-18D), across two replicates fold-change of ≤ 0.2 was achieved, though with more noise (Supplementary Fig. S18B–C). The three other synthetic sRNAs targeting downstream of the RBS (Syn-ChiX-22RD, Syn-ChiX-24RD, and Syn-ChiX-25D) were also tested in the Chi.Bio, showing functional repression of GFPmut3 (Supplementary Fig. S18D–F). Given its similar repression but cleaner, less noisy behaviour than Syn-ChiX-18D, Syn-ChiX-20D was taken forward for subsequent spRNA design.

After validation of Syn-ChiX-20D, several control experiments were carried out to confirm the intended mechanism (Supplementary Note S5.1). These included controls for the sRNA itself (Supplementary Fig. S23), the engineered binding region (Fig. 2F), and unintended target binding (Fig. 2G and Supplementary Fig. S24).

### Design and characterisation of synthetic sponge RNAs

#### Design of synthetic sponge RNAs

After validating Syn-ChiX-20D, we next designed a synthetic spRNA for this sRNA based on the natural ChiX spRNA ChbBC ^71,74,75,80^ (Fig. 3A). Briefly, the ChbBC binding region was identified, extended, and reprogrammed, followed by the same *in silico* validation workflow used for sRNA design (Fig. 2A). Full details of the design are provided in Supplementary Note S3.2.

**Figure 3:**
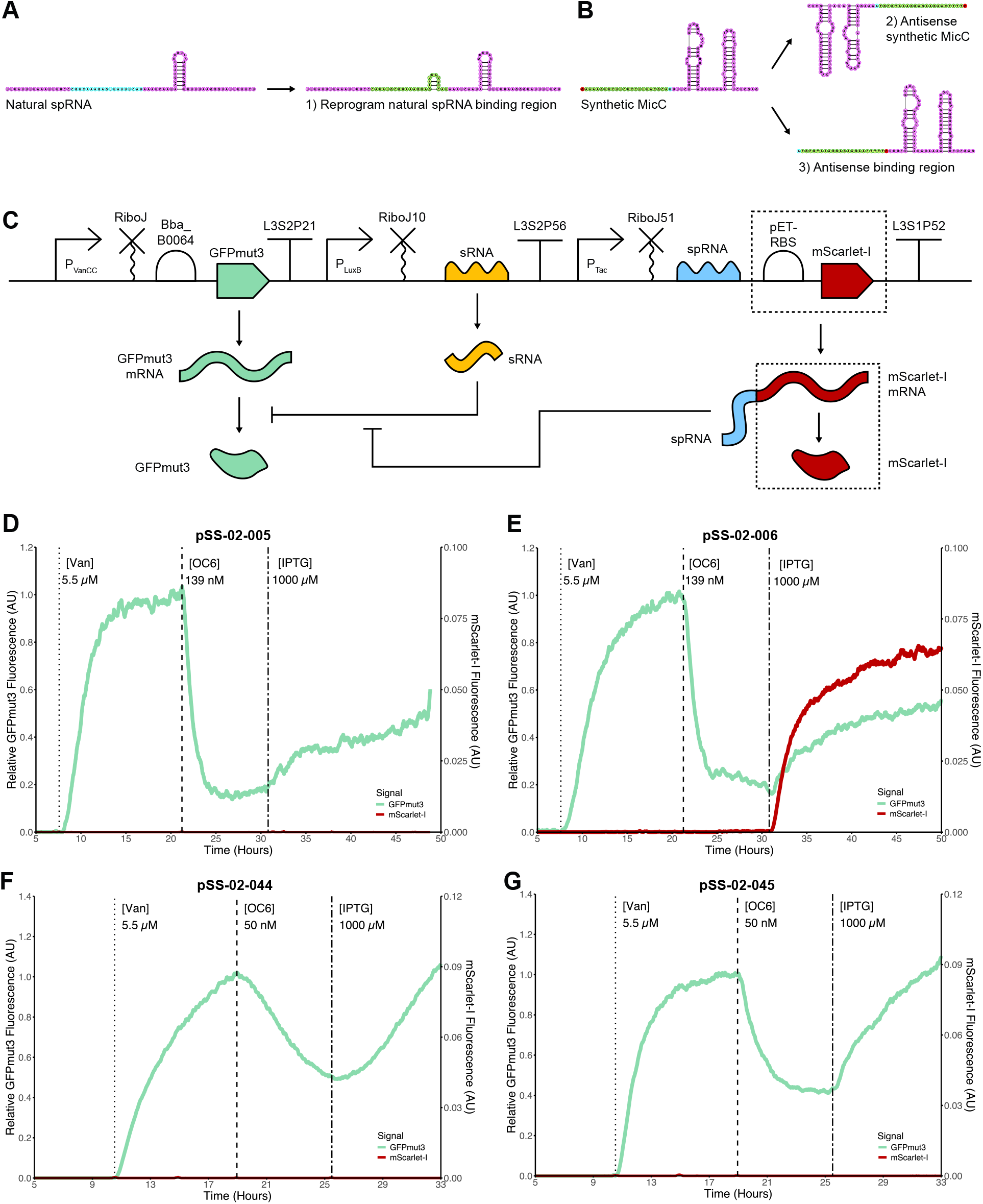
Design and characterisation of synthetic sponge RNAs: **(A)** Synthetic spRNAs can be designed by extending and reprogramming the binding region of a natural spRNA. **(B)** Alternative strategies include expressing the antisense of a synthetic sRNA or reprogramming its binding region to be antisense. **(C)** SBOL Visual ^99^ representation of the synthetic gene circuit used to test spRNAs. The regulatory RNAs are shown as placeholders to be replaced with a specific synthetic sRNA–spRNA pair according to the chosen design strategy. A second fluorescent protein, mScarlet-I, can optionally be placed downstream of the spRNA to enable dynamic measurement of spRNA expression. **(D)** Chi.Bio characterisation of pSS-02-005, containing GFPmut3 as the target mRNA, Syn-ChiX-20D as the sRNA, and Syn-ChbBC-20D as the sponge RNA. Upon sponge RNA expression, GFPmut3 repression is relieved as desired. **(E)** Chi.Bio characterisation of pSS-02-006, equivalent to pSS-02-005 but with mScarlet-I downstream of the spRNA. The circuit functions as intended, with GFPmut3 recovered and mScarlet-I confirming spRNA expression. **(F)** Chi.Bio characterisation of pSS-02-044, in which the sRNA is Syn-MicC-GFP and the spRNA is the antisense of the full sRNA. GFPmut3 is recovered as desired. **(G)** Chi.Bio characterisation of pSS-02-045, in which the sRNA is Syn-MicC-GFP and the spRNA uses the same scaffold with an antisense binding region. Again, the circuit functions as desired.

To broaden the applicability of synthetic spRNAs beyond natural sponge scaffolds, we also explored three alternative design strategies for the widely used MicC scaffold, which has no known natural spRNA (Fig. 3B): (1) expression of the full antisense sequence of a synthetic MicC-derived sRNA; (2) expression of a variant in which only the binding region is antisense; and (3) expression of ChiX as a spRNA for a MicC-derived sRNA. This first required the design of a synthetic MicC-derived sRNA targeting GFPmut3, following previously described methods ^28,29^ (Fig. 2B), yielding Syn-MicC-GFP (Supplementary Note S3.3). These strategies were then used to generate three corresponding synthetic spRNAs (Supplementary Note S3.4). The resulting ChbBC-derived and alternative synthetic spRNAs were assembled (Supplementary Note S3.5) into circuits of the form shown in Fig. 3C for experimental characterisation.

#### Characterisation of ChbBC-derived sponge RNA circuits

We characterised pSS-02-005 (Syn-ChiX-20D Syn-ChbBC-20D) and pSS-02-006 (Syn-ChiX-20D Syn-ChbBC-20D with mScarlet-I) in the Chi.Bio with sequential induction of GFPmut3, Syn-ChiX-20D, and the corresponding spRNA (Fig. 3D–E). In both circuits, induction of Syn-ChbBC-20D with 1000 µm IPTG led to recovery of GFPmut3 expression with the expected monotonic rise towards steady state (Fig. 3D–E). In pSS-02-006, this was accompanied by the expected mScarlet-I expression, validating spRNA expression (Fig. 3E). Replicate Chi.Bio experiments showed similar qualitative and quantitative behaviour (Supplementary Fig. S19).

Next, pSS-02-005 and pSS-02-006 were characterised with lower Syn-ChiX-20D induction (50 nm OC6 versus 139 nm) to determine whether higher sponge-mediated recovery of expression could be achieved when sRNA concentrations were lower. As predicted, higher recovery of GFPmut3 expression was achieved under these conditions (Supplementary Fig. S20A–B). However, unexpectedly for pSS-02-006, GFPmut3 expression rose above its initial steady state (Supplementary Fig. S20B). Normalising fluorescence by growth rate corrected this (Supplementary Fig. S20D), suggesting that growth-rate changes due to the burden imposed by gene expression could be responsible for this behaviour (see later investigation of growth feedback). As pSS-02-006 appeared to be the best-functioning spRNA circuit, it was further tested over four biological replicates with multiple step-changes in both sRNA and spRNA expression (Supplementary Fig. S21), further validating the consistent qualitative behaviour of the circuit.

Negative controls for the ChbBC-derived spRNA circuits are described in Supplementary Note S5.2. Briefly, these included a circuit lacking the spRNA (pSS-02-001), confirming that IPTG alone did not cause the response (Supplementary Fig. S25 and Fig. 4I); a version of Syn-ChbBC-20D with a randomised binding region (pSS-02-027), confirming that the engineered binding region was required (Supplementary Fig. S26 and Fig. 4J); and a circuit containing the spRNA but lacking the sRNA, confirming that the spRNA acts through sRNA sequestration (Supplementary Fig. S27). In a further experiment, GFPmut3 in pSS-02-005 was replaced with mScarlet-I and the sRNA and spRNA were redesigned to regulate mScarlet-I, generating pSS-02-040 (Supplementary Note S5.2.1), which functioned as desired (Supplementary Fig. S29), demonstrating the generalisability of spRNA design.

**Figure 4:**
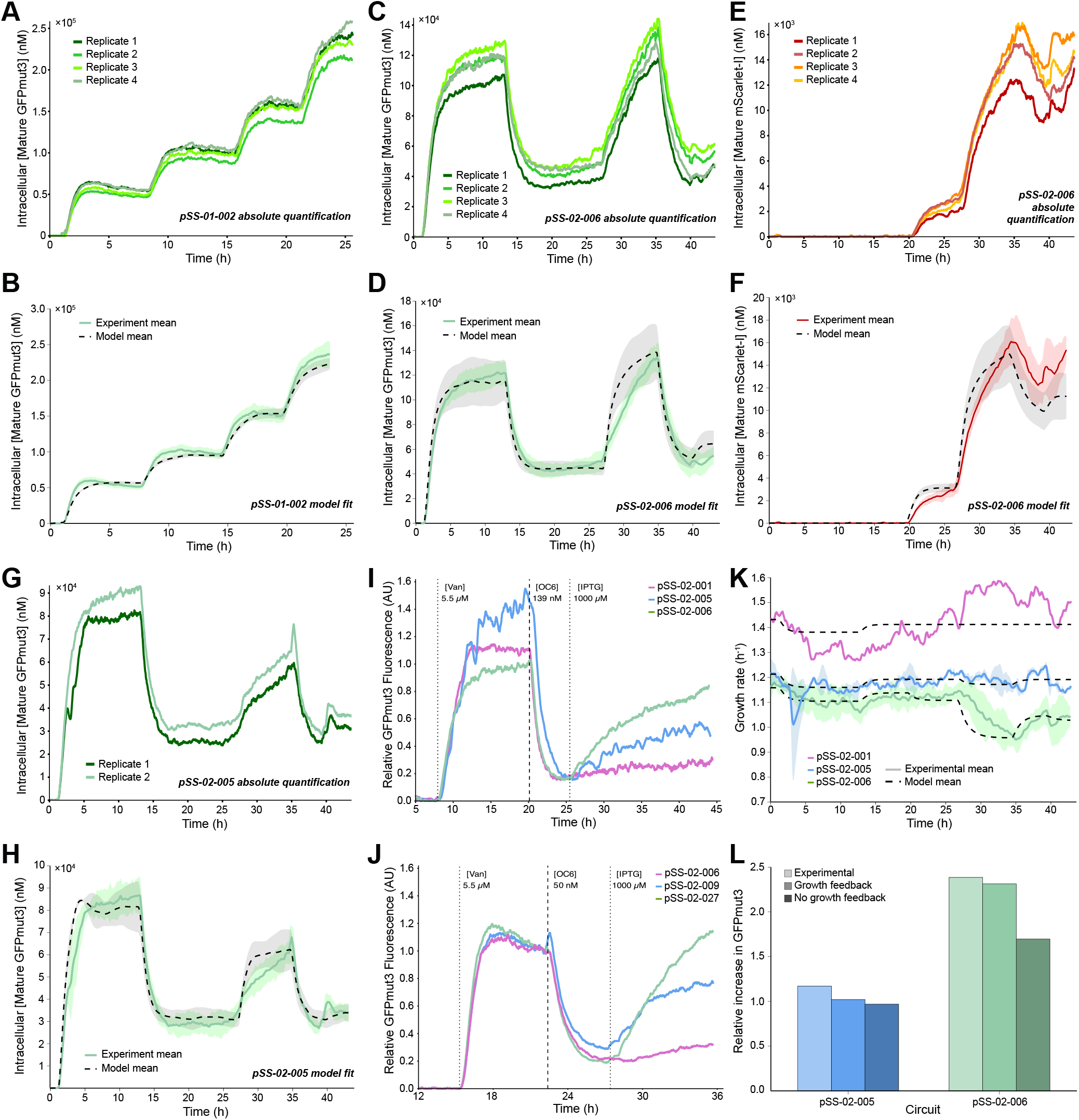
Quantitative characterisation with absolute quantification and model fitting of ChbBC-based synthetic sponge RNA circuits: **(A-B)** Representative Chi.Bio characterisation of promoter (pSS-01-002; P_LuxB_) and the corresponding fitted model mean against the experimental mean. **(C-D)** GFPmut3 signal for Chi.Bio characterisation of pSS-02-006 (sponge circuit with mScarlet-I) and the corresponding fitted model mean against the experimental mean. **(E-F)** mScarlet-I signal for Chi.Bio characterisation of pSS-02-006 and the corresponding fitted model mean against the experimental mean. **(G-H)** GFPmut3 signal for Chi.Bio characterisation of pSS-02-005 (sponge circuit without mScarlet-I) and the corresponding fitted model mean against the experimental mean. **(I)** Chi.Bio characterisation of spRNA circuits (no sponge, sponge without mScarlet-I, and sponge with mScarlet-I), showing that GFPmut3 recovery is greatest when mScarlet-I is present and absent without spRNA. **(J)** Introducing stop codons at the start of the mScarlet-I CDS to generate pSS-02-009 reduced gene expression burden with minimal change to the spRNA transcript sequence. Chi.Bio characterisation shows reduced GFPmut3 recovery relative to the sponge circuit with mScarlet-I (pSS-02-006), similar to the sponge circuit without mScarlet-I (pSS-02-005; Panel I), suggesting that burden partly explains the apparent improved performance of the circuit with mScarlet-I. Both circuits are compared with a negative control spRNA circuit containing a non-targeting binding region (pSS-02-027), which shows no spRNA-mediated GFPmut3 recovery. **(K)** Growth feedback models were fit using a cost function incorporating both estimated growth rates and protein concentrations. For both spRNA circuits, these fits recapitulated the experimental growth-rate changes. **(L)** The relative increase in GFPmut3 after spRNA induction in experiments, fixed-growth modelling, and growth-feedback modelling shows that including growth feedback improves prediction accuracy in both spRNA circuits, particularly in the case with mScarlet-I.

#### Characterisation of alternative spRNA design strategies

We next characterised the alternative spRNA design strategies in Chi.Bio experiments. We first explored strategy (1), in which a spRNA (Syn-MicC-spRNA-EA) was generated by expressing the reverse complement of the sRNA (Syn-MicC-GFP). This was tested in a circuit without mScarlet-I, pSS-02-044 (Fig. 3F), and in a circuit with mScarlet-I (Supplementary Fig. S22A), in both cases functioning as desired. Notably, different dynamics were observed compared to the previously characterised sRNA and spRNA circuits (Fig. 3D–E). We next investigated strategy (2), in which the spRNA (Syn-MicC-spRNA-AB) maintained the scaffold region of the sRNA (Syn-MicC-GFP) but the binding region was the antisense of the sRNA binding region. This circuit was again tested in the Chi.Bio, with circuits pSS-02-045 (Fig. 3G) and pSS-02-048 (Supplementary Fig. S22B) functioning as desired, although upon growth-rate normalisation of the fluorescence data the spRNA functioned only in the absence of mScarlet-I (Supplementary Fig. S22C–D). Finally, we investigated strategy (3), in which ChiX was co-opted as a spRNA for Syn-MicC-GFP (Syn-ChiX-spRNA). Two circuits were tested, one without mScarlet-I and one with mScarlet-I (Supplementary Fig. S22E–F). In this case, the spRNA functioned only when mScarlet-I was absent (Supplementary Fig. S22E).

#### Quantitative characterisation of ChbBC spRNA circuits

While the preceding characterisation showed that the ChbBC-based spRNA circuits functioned as expected and qualitatively similarly across replicates, we next characterised them quantitatively. We applied FPCountR-based ^88^ absolute quantification to Chi.Bio characterisation ^90^ across multiple replicates and reactors, confirming that circuit behaviour was quantitatively consistent across experiments and generating a rich dataset for subsequent model fitting, including under inducer conditions that revealed behaviours not evident from the initial characterisation (see subsequent investigation of resource competition).

These experiments included the standardised promoter transcription units (pSS-01-001, pSS-01-002, and pSS-01-003) (Supplementary Fig. S2), the sRNA circuit (pSS-02-001) (Fig. 2C with Syn-ChiX-20D as sRNA), and the two ChbBC-based spRNA circuits (pSS-02-005 (no mScarlet-I) and pSS-02-006 (with mScarlet-I)) (Fig. 3C with Syn-ChiX-20D and Syn-ChbBC-20D as sRNA and spRNA), each across multiple replicates and inducer conditions. Calibration of the microplate reader for the relevant fluorescent proteins is shown in Supplementary Fig. S32A–F, and quantification of Chi.Bio samples yielded highly explanatory reactor-specific conversion curves (Supplementary Figs. S34 and S35). These were used to quantify Chi.Bio fluorescence time courses (Fig. 4A, C, E, G, and Supplementary Fig. S36), yielding intracellular protein concentration estimates that were highly consistent across experiments and compatible with estimates of *E. coli* protein carrying capacity ^100^.

To distinguish changes in intracellular protein concentration arising from circuit behaviour from those due to altered growth-mediated dilution, we also estimated time-varying growth rates for each experiment using an *ad hoc* Bayesian estimator tailored to turbidostat data ^96^ (see Methods and example in Supplementary Fig. S37A). These estimates showed that induction of different circuit components could substantially alter growth rate during the experiment (Supplementary Fig. S37B), motivating explicit incorporation of growth dynamics in later modelling.

### Progressive model fitting supports Hfq resource competition

Using our quantitative characterisation data, we next fit a series of RNA-circuit models. Based on previous modelling efforts ^32,53^, we first tested a simple mass-action annihilation model (Supplementary Fig. S39), then extended it to include ChiX predation of mRNA (Supplementary Fig. S40) and other asymmetric RNA interactions (Supplementary Fig. S41) motivated by the reported predatory nature of ChiX regulation ^71^. While these models often captured the sRNA dynamics well, none could recapitulate the observed GFPmut3 response to spRNA induction, instead predicting increases in GFPmut3 after initial spRNA induction that were not observed experimentally (Supplementary Figs. S39, S40, and S41).

Because unexpected behaviours in synthetic gene circuits are often shaped by competition for limited shared cellular resources ^101^, and because ChiX is known to compete strongly for Hfq ^85,87^, we next incorporated resource competition using a recently published resource-aware modelling framework ^102^. This substantially improved fits to GFPmut3 in the spRNA circuits (Supplementary Fig. S42), but when mScarlet-I translation from the spRNA in pSS-02-006 was included, the model poorly captured both mScarlet-I and GFPmut3 dynamics (Supplementary Fig. S43). Finally, we extended the model to include explicit sRNA–mRNA and sRNA–spRNA complex formation, while allowing translation of mScarlet-I from the spRNA–sRNA complex, yielding our final model (Supplementary Equations (S13)). This model produced quantitatively good fits to GFPmut3 for the promoter constructs (Fig. 4B; Supplementary Fig. S44A–C), and for the spRNA circuits both with (Fig. 4D; Supplementary Fig. S44G–J) and without mScarlet-I (Fig. 4H; Supplementary Fig. S44E–F), while also capturing the mScarlet-I dynamics more accurately (Fig. 4F; Supplementary Fig. S44K–N). Together, these results support a model in which both Hfq resource competition and translation from sRNA–spRNA complexes contribute to the observed circuit behaviour. This final model and its parameters (Supplementary Tables S8 and S9) were taken forward for subsequent modelling; the full fitting process is described in Supplementary Note S7.

### Growth-feedback enhances apparent sponge RNA regulation

In multiple experiments, the spRNA with downstream mScarlet-I (pSS-02-006) appears to function as a stronger regulator than the same spRNA without mScarlet-I (pSS-02-005), leading to greater recovery of GFPmut3 expression (Fig. 3D–E; Fig. 4I; Supplementary Figs. S19 and S20). We hypothesised that this could arise from ribosome-mediated protection of the spRNA transcript from degradation during mScarlet-I translation, changes in transcript structure or function caused by inclusion of mScarlet-I, or growth-feedback effects caused by the burden of mScarlet-I expression.

#### Experimental investigation of mScarlet-I effect

To experimentally distinguish between these hypotheses, we introduced several stop codons at the beginning of the mScarlet-I CDS in pSS-02-006 (Supplementary Fig. S47) by site-directed mutagenesis, generating a spRNA circuit with almost the same sequence as pSS-02-006 but without mScarlet-I expression (pSS-02-009). This design should reduce gene-expression-related burden and ribosome protection while making minimal changes to the primary nucleotide sequence, and therefore minimal changes to spRNA structure and function. We characterised this stop-codon circuit alongside the spRNA circuit with mScarlet-I (pSS-02-006) and a spRNA circuit with a randomised binding region (pSS-02-027). In the stop-codon circuit, spRNA-mediated recovery was reduced compared to the spRNA circuit with mScarlet-I, suggesting that the stronger regulation is largely due to protein expression (burden or ribosome-mediated protection) rather than nucleotide changes associated with including mScarlet-I downstream (Fig. 4J).

#### Growth-feedback modelling supports a burden-mediated explanation

We next used modelling to distinguish between the burden and ribosome-protection hypotheses. Under the burden explanation, increased intracellular heterologous protein concentration arises through growth feedback: expression of synthetic protein slows growth, thereby reducing dilution of expressed proteins ^103^. We therefore fit a growth-feedback variant of our final RNA-circuit model, in which growth rate was predicted from circuit expression rather than supplied directly from experimental measurements (see Methods and Supplementary Note S8).

This model quantitatively captured the experimentally observed changes in growth rate for pSS-02-005 and pSS-02-006 (Fig. 4K; Supplementary Fig. S48A), while also fitting the fluorescent protein trajectories well (Supplementary Fig. S48B–D). By contrast, when growth feedback was omitted and growth was instead fixed (Supplementary Note S8), the model did not fully capture the relative GFPmut3 changes following spRNA induction (Fig. 4L). This mismatch was more pronounced for pSS-02-006, where growth feedback would be expected to be stronger because both GFPmut3 and mScarlet-I are expressed. This indicates that improved spRNA stability alone does not explain the stronger regulation in pSS-02-006.

Together, these results suggest that the improved regulatory strength of Syn-ChbBC-20D with mScarlet-I arises from both burden-mediated growth feedback and improved transcript stability. More broadly, these data and modelling further support the importance of accounting for circuit effects on growth during both design and characterisation ^104–106^, and provide an additional rationale for using continuous-culture platforms in which growth rate can be measured and, in principle, controlled.

### Tuneability of sponge RNA response

#### Expression-mediated tuning of spRNA regulation

Using our model (Supplementary Equations (S13)), we tested whether spRNA-mediated recovery of gene expression is tuneable by varying spRNA induction in pSS-02-006 (Fig. 5A) and pSS-02-005 (Fig. 5B). In both circuits, increasing spRNA induction produced dose-dependent recovery of GFPmut3 expression, with this effect much stronger in pSS-02-006 (Fig. 5A) than in pSS-02-005 (Fig. 5B), consistent with the lower inferred stability of the spRNA in pSS-02-005. These simulations included resource competition for Hfq but not growth feedback, suggesting that experimentally these differences may be larger in both circuits.

**Figure 5:**
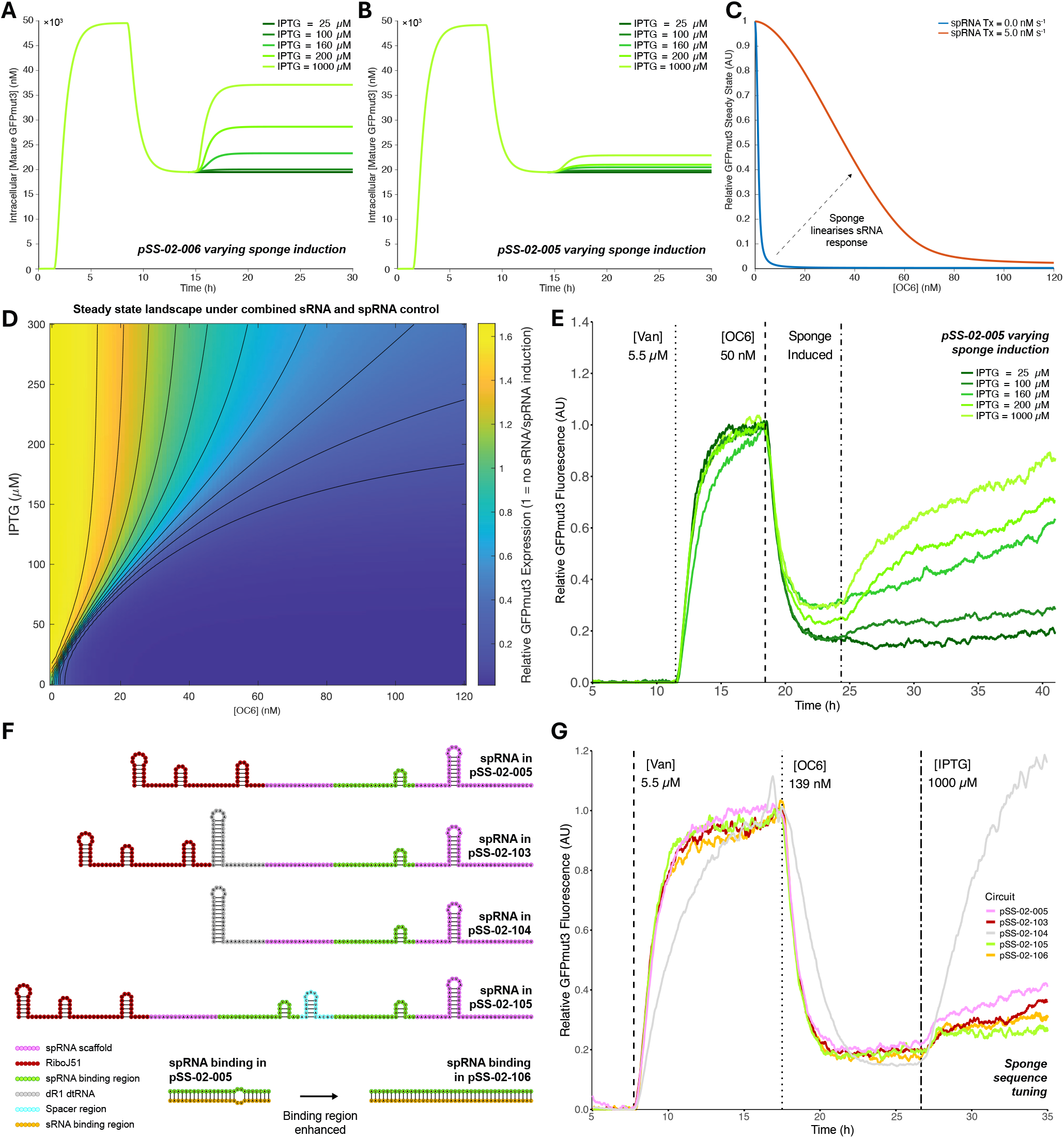
Tuneability of sponge RNA-controlled gene expression: **(A)** Deterministic simulations of pSS-02-006 (spRNA circuit with mScarlet-I). Sequential induction of GFPmut3 (5.5 µm vanillic acid), sRNA (50 nm OC6), and spRNA shows that gene expression can be tuned by varying spRNA inducer (IPTG) concentration. **(B)** Equivalent deterministic simulations of pSS-02-005 (spRNA circuit without mScarlet-I) shows reduced tuneability. **(C)** Deterministic steady-state GFPmut3 dose–response to OC6 induction of sRNA in the presence or absence of strong spRNA expression. Without spRNA, the response is switch-like, whereas spRNA expression partially linearises the system. **(D)** Deterministic steady-state GFPmut3 heat map from a 2D scan of sRNA (OC6) and spRNA (IPTG) inducer concentrations. Steady-state GFPmut3 at zero OC6 and IPTG was normalised to one. Increasing IPTG effectively linearises the response, requiring larger changes in OC6 to produce comparable knockdown, while also reducing sRNA leakiness and increasing dynamic range (relative steady state *>* 1). **(E)** Chi.Bio experiments testing the model in Panel B show that GFPmut3 expression was tuneable by varying spRNA expression. The observed tuneability was more similar to Panel A. **(F)** Sponge RNA regulation may also be tuned at the sequence level by incorporating degradation tuning RNAs (dtRNAs) ^20^, removing RiboJ51 and including dtRNA, duplicating binding regions, or removing binding mismatches. RNA structure components were predicted individually using Vienna RNAFold ^59^ and visualised with RNACanvas ^94^. **(G)** Experimental test of sequence tuning strategies from Panel F in Chi.Bio using sequential induction of GFPmut3, sRNA, and spRNA variants. The spRNA containing the dR1 dtRNA and lacking RiboJ51 showed significantly improved recovery of GFPmut3 expression after spRNA induction, whereas all other variants performed similarly to the original spRNA. A spike in GFPmut3 fluorescence prior to sRNA induction is observed in pSS-02-104 due to turbidostat hardware issues but does not affect the conclusion.

We next used simulations to test whether a two-input circuit (sRNA and spRNA) provides advantages for tuneability of gene expression. Simulating steady-state GFPmut3 expression in response to increasing concentrations of sRNA inducer (OC6) alone revealed steep switch-like behaviour, in which small changes in OC6 concentration led to large changes in GFPmut3 expression (Fig. 5C), presenting a potential challenge when fine-tuning of gene expression is required. We then included spRNA transcription and performed a 2D scan of spRNA (IPTG) and sRNA (OC6) induction for pSS-02-006 (Fig. 5D). In contrast to the sRNA-only circuit, inclusion of the spRNA produced regions of inducer combinations that achieved similar expression levels, and in some regimes larger changes in OC6 were required to achieve further knockdown (Fig. 5C–D), effectively linearising the system. Notably, spRNA expression in the absence of sRNA induction led to *>* 1 relative GFPmut3 expression (Fig. 5D), while at high spRNA expression a higher concentration of sRNA inducer was required to observe an initial effect, providing evidence for leakiness tuning and thresholding, respectively.

We tested tuneability experimentally in pSS-02-005, where growth feedback would be expected to have a smaller effect, by carrying out a Chi.Bio experiment using the same inducer step changes as the simulations (Fig. 5B). This demonstrated tuneability of GFPmut3 expression (Fig. 5E): no discernible recovery was achieved with 25 µm IPTG, while 100, 160, and 200 µm produced varying intermediate levels of recovery, and 1000 µm gave the highest recovery of GFPmut3 expression. Interestingly, the experimental results for pSS-02-005 (Fig. 5E) appear more similar to the simulations for pSS-02-006 (Fig. 5A) than to the simulations for pSS-02-005 (Fig. 5B), which may reflect growth feedback, bioreactor or experimental differences not captured in the simulations, or differing levels of initial sRNA knockdown.

#### Sequence-mediated tuning of spRNA regulation

We next investigated whether the regulatory strength of a spRNA could be engineered at the sequence level. We tested several sequence-engineering strategies (Fig. 5F) to improve spRNA binding or stability: (i) based on previous work improving stability of mRNAs and ncRNAs using degradation-tuning RNA (dtRNA) ^20^, we incorporated the best-performing dtRNA, dR1, upstream of the spRNA; (ii) replaced RiboJ51 with dR1; (iii) included two binding regions for the cognate sRNA; and (iv) reprogrammed the two-nucleotide mismatch in the spRNA to give full complementarity with the cognate sRNA. Circuits were assembled for each strategy (pSS-02-103 through pSS-02-106) and tested in a Chi.Bio experiment alongside pSS-02-005 (Fig. 5G). Strategy (ii) was the only one that improved performance, achieving full recovery of gene expression (Fig. 5G), while the other strategies performed similarly to, or slightly worse than, the original circuit. Although these other strategies may still work for different spRNAs, at a minimum this demonstrates the potential to enhance stability of natural spRNAs through inclusion of dtRNAs ^20^. Although it cannot be discounted that removing RiboJ51 contributed to the improved performance, previous results suggest that RiboJ elements can in fact improve transcript stability ^107^, while it would be unsurprising that the dtRNA, a 5^*′*^ stability hairpin, only worked when placed at the 5^*′*^ end of the transcript.

### Sponge RNAs can be extended to regulate multiple targets

We next tested whether spRNA circuits could be extended to regulate multiple genes simultaneously, inspired by previous work on fusion sRNAs ^61^ and natural multi-target sRNAs ^108^ and spRNAs ^73,78^. Following methods developed for fusion sRNAs ^61^, we first designed multi-target ChiX- and MicC-derived sRNAs containing two binding regions targeting GFPmut3 and mScarlet-I (Supplementary Note S3.6). Chi.Bio characterisation experiments involving sequential induction of mScarlet-I, GFPmut3, and the respective multi-target sRNAs showed strong simultaneous knockdown of both genes in both circuits (Supplementary Fig. S30). These results suggested two strategies for multi-target regulation using spRNAs: (i) a spRNA containing two binding regions, each targeting a different sRNA; and (ii) a spRNA containing a single binding region targeting one of the above multi-target sRNAs.

We first investigated strategy (i) using a circuit with constitutive expression of mScarlet-I and GFPmut3, two MicC-derived sRNAs targeting one of each protein, and a MicC-based spRNA with binding regions for both sRNAs (Fig. 6A; Supplementary Note S3.6). In Chi.Bio experiments, sequential induction of the sRNAs followed by the spRNA gave the expected behaviour: each sRNA orthogonally downregulated its target gene, and induction of the spRNA led to simultaneous recovery of both genes (Fig. 6C). Interestingly, the dynamics of both sRNA- and spRNA-mediated regulation differed meaningfully between the two target proteins, highlighting the potential for temporal control of two genes using multi-target RNAs.

**Figure 6:**
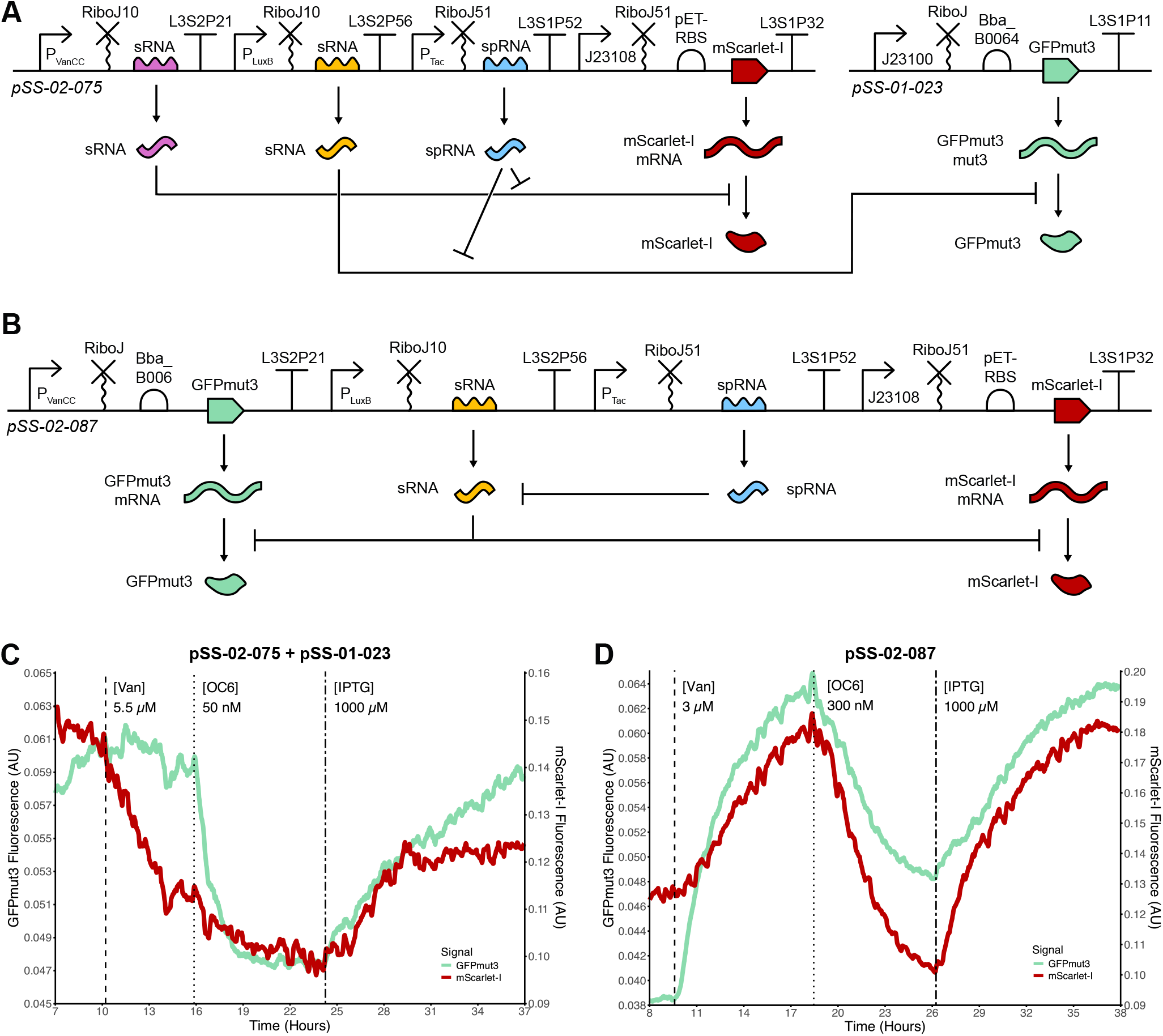
Design and characterisation of multi-target sRNA and sponge RNA circuits: **(A)** Two plasmids (pSS-02-075 and pSS-01-023) come together to form a multi-target sponge RNA circuit where two sRNAs downregulate two constitutively expressed targets orthogonally (GFPmut3 and mScarlet-I). A single sponge RNA binds both sRNAs de-repressing protein expression. **(B)** A similar circuit in which a single sRNA binds both mScarlet-I and GFPmut3, downregulating their expression; this sRNA can then be bound by a single sponge RNA, which de-represses expression of both proteins. **(C)** Chi.Bio characterisation of pSS-02-075 co-transformed with pSS-01-023. Both proteins are expressed constitutively, followed by induction of the sRNA for mScarlet-I leading to repression of mScarlet-I, followed by the sRNA for GFPmut3 leading to repression of GFPmut3. Finally the sponge RNA which binds both sRNAs is induced leading to de-repression of both fluorescent proteins. **(D)** Chi.Bio characterisation of pSS-02-087. mScarlet-I is expressed constitutively after which GFPmut3 is induced. This is followed by induction of the sRNA that binds both GFPmut3 and mScarlet-I mRNA leading to downregulation of both proteins. Finally a sponge RNA for that sRNA is induced leading to de-repression of both proteins.

We next investigated strategy (ii) using a circuit with constitutive expression of mScarlet-I, inducible expression of GFPmut3, a multi-target MicC-derived sRNA targeting both genes, and a spRNA with the same scaffold but an antisense binding region (Fig. 6B; Supplementary Note S3.6). This circuit was characterised in the Chi.Bio by sequential induction of GFPmut3, the sRNA, and the spRNA (Fig. 6D). As before, induction of the sRNA caused simultaneous knockdown of both genes, followed by simultaneous recovery upon spRNA induction (Fig. 6D). In contrast to strategy (i), the two genes appeared to be regulated on similar timescales (Fig. 6D). Taken together, these results provide a proof-of-concept demonstration that our synthetic spRNA design strategies can be used to engineer low-burden gene circuits capable of regulating multiple genes simultaneously.

### Quantitative Investigation of Synthetic spRNA Systems Biology

Using our fitted model (Supplementary Equations (S13) and Supplementary Tables S9 and S8), we next investigated quantitative characteristics of spRNA-based regulation, including the hypotheses outlined in Fig. 1F. Several of our results already support such behaviours, including temporal control (Fig. 6C), tuneability (Fig. 5), and thresholding and leakiness tuning (Fig. 5D). While quantitative characteristics of sRNA-based regulation have been investigated previously ^53,54,109^, they have not been investigated for spRNAs.

#### Sponge RNAs reshape response functions

One well described quantitative aspect of sRNA-mediated regulation is the threshold-linear response ^53,54^, in which expression of a regulated gene is strongly repressed below a critical transcription level but increases approximately linearly above this threshold. Using a mass-action annihilation model for our sRNA circuit (Supplementary Equations (S16)), we reproduced these behaviours: our fitted *γ*_*sm*_ yielded a non-ideal threshold-linear response, while increasing *γ*_*sm*_ produced behaviour approaching the idealised limit (Fig. 7A). By contrast, a perfectly predatory mass-action sRNA model (Supplementary Equations (S17)) gave an approximately linear response with slope tuned by binding strength (Fig. 7B). In our more realistic model with partial predation and resource competition (Supplementary Equations (S13)), however, we again observed close-to-idealised threshold-linear behaviour, but with the threshold now dependent not only on sRNA transcription rate but also on sRNA binding strength (Fig. 7C). When resource availability was held fixed (*R* = *R*_total_), the same model behaved more like the mass-action predation model, exhibiting an approximately linear response in which binding strength primarily tuned the slope (Fig. 7D).

**Figure 7:**
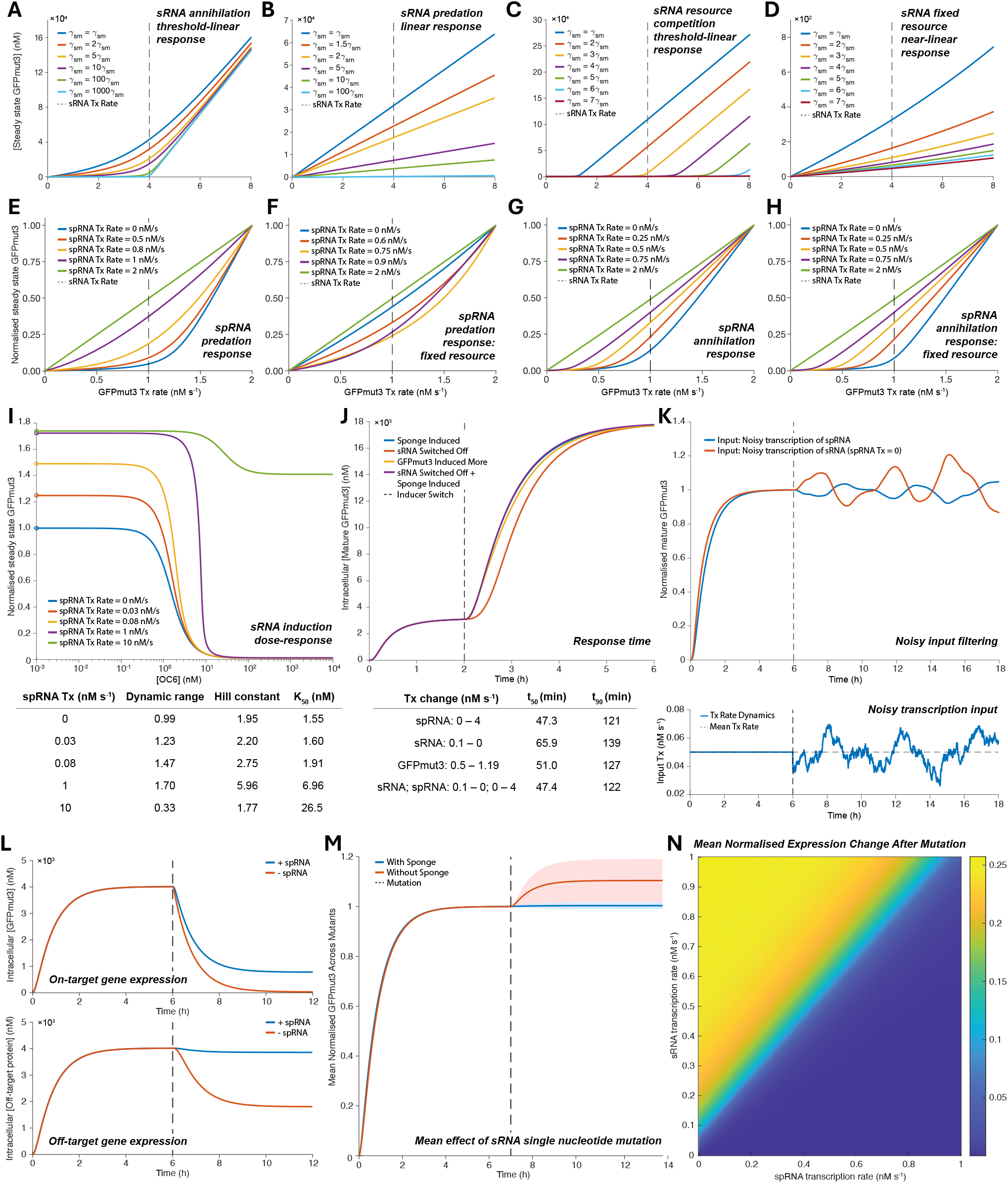
Quantitative sponge RNA regulatory behaviour: **(A–D)** GFPmut3 transcription-rate (Tx) response curves for fixed sRNA Tx and variable sRNA–mRNA binding strength for: **(A)** an annihilation model with threshold-linear response; **(B)** a predation model with linear response; **(C)** our resource competition model with threshold-linear response; and **(D)** the same model with fixed resources, with near-linear response. **(E–H)** Similar simulations for variable spRNA Tx, with normalised GFP expression plotted. In our resource competition model with predation, **(E)** increasing spRNA Tx progressively linearises the response. With fixed resources, **(F)** spRNA Tx instead introduces substantial non-linearity. In annihilation models, **(G–H)** increasing spRNA Tx shifts the threshold. **(I)** spRNA Tx alters Hill coefficients, dynamic ranges, and *K*_50_ of sRNA induction dose-response curves. **(J)** spRNAs can improve response times; for example, inducing spRNA gives a faster rise to steady state than switching off sRNA transcription. **(K)** spRNAs can reduce input disturbances in some regimes, here by damping noisy Tx input when applied to spRNA rather than sRNA. **(L)** spRNA expression can reduce effective off-target regulation: without spRNA, both on- and off-targets are downregulated, whereas with spRNA only the on-target is substantially downregulated. **(M)** Mean trajectories for all 164 single-point mutations across a 20-nucleotide sRNA binding site. After mutation, steady-state GFP expression changes substantially without spRNA, but minimally with spRNA. **(N)** Across plausible sRNA and spRNA Tx rates, spRNA reduces the effect of such mutations on output.

We next investigated how spRNA expression modulates this behaviour. Under fixed sRNA transcription, increasing spRNA transcription progressively linearised the response of GFPmut3 to increasing GFPmut3 transcription (Fig. 7E). In the absence of resource competition (*R* = *R*_total_), the model predicted linear behaviour both when spRNA was absent and when it was in excess, whereas intermediate spRNA transcription introduced non-linearity (Fig. 7F). When stoichiometric annihilation was enforced by setting *n* = 0, spRNA transcription tuned the threshold level in both resource-limiting and fixed-resource cases until spRNA was in excess, at which point the response again became linear (Fig. 7G–H).

Sponge RNAs can also tune Hill-type response functions. Simulating normalised steady-state GFPmut3 expression across sRNA inducer (OC6) concentrations for varying spRNA transcription rates showed that sponge transcription increased the maximum on-state with relatively little change in the minimum off-state, giving up to a 1.7-fold increase in dynamic range (Fig. 7I). This increase in dynamic range results from spRNA transcription tuning out promoter leakiness in sRNA expression. This spRNA-mediated leakiness tuning may be particularly important in sRNA biology, where promoter leakiness can have an outsized impact: best-in-class promoter leakiness has little effect when driving the protein itself, but applying the same leakiness to sRNA expression can strongly alter protein output (Supplementary Fig. S49), which may explain why spRNAs are common in important tightly-regulated cellular processes^69^. At excessively high spRNA transcription, however, the minimal off-state rose and dynamic range decreased (Fig. 7I). Sponge RNA transcription also shifted threshold position (*K*_50_) rightwards, and fitted Hill functions changed shape with spRNA, with effective cooperativity either increasing (*n* = 1.95 to *n* = 5.96) or decreasing (*n* = 1.77) depending on the regime (Fig. 7I). Together, these results show that spRNAs are not merely tools to reduce sRNA-mediated repression, but can meaningfully tune response gains, thresholds, and shapes. These characteristics could be useful for classifier-like modules, tight ON–OFF control without complex promoter re-engineering, and dynamic-range matching in multi-layered circuits.

#### Sponge RNAs may modulate dynamic behaviour

We next investigated how spRNAs affect circuit dynamics. Faster response times are a potential benefit of RNA-based gene regulation, though theoretical studies have reached conflicting conclusions, with some suggesting no improvement in response times ^109^ and others suggesting faster responses ^54,57^. In our modelling for trajectories rising from zero to a chosen steady state, inclusion of spRNA generally increased rise time relative to comparable sRNA-only solutions (Supplementary Note S9.2.1). However, when considering increasing expression of a gene already under sRNA regulation, switching on the spRNA gave the fastest 50% and 90% rise times for the specific step changes tested (Fig. 7J). Although the difference was modest (50% rise time: 47.3 mins vs. 65.9 mins) under idealised step changes, it became larger (50% rise time: 47.3 mins vs. 127.8 mins) when inducer dilution was included to reflect bioreactor (turbidostat) conditions in which inducer concentration cannot be rapidly decreased (Supplementary Fig. S52). These results suggest that, while spRNAs do not generally accelerate expression from zero, they may provide a useful means of rapidly switching off existing sRNA repression, particularly in biotechnological contexts where bioreactors are used. In natural systems, this role for spRNAs, as a specific means of switching off existing sRNA regulation ^67^, is also intuitively consistent with the sometimes relatively stable nature of sRNAs ^110^.

We also investigated how spRNA networks respond to input disturbances. For the parameter set tested, output noise was lower when a noisy transcription was applied to the spRNA than when comparable noise was applied directly to the sRNA (Fig. 7K). Across 50 repeated noisy inputs with matched standard deviation, the spRNA-input case also gave a lower mean Gain in Coefficient of Variation, indicating stronger attenuation of input noise (Supplementary Fig. S53). Although this does not prove a general disturbance-rejection advantage across all parameter regimes, it suggests that spRNAs can improve filtering in some regimes. Similar behaviour was also observed for other disturbance types, including spike and square-wave inputs (Supplementary Note S9.3).

#### spRNAs may improve specificity

Sponge RNAs may also improve specificity when an sRNA represses both an intended on-target and an unintended off-target. To test this, we simulated an extended version of our model with two sRNA-regulated targets, one which is bound more strongly than the other (Supplementary Note S9.4). Without spRNA, induction of the sRNA led to pronounced downregulation of both the on-target and off-target (Fig. 7L). By contrast, when an spRNA was co-expressed, the on-target remained repressed, while the off-target became largely insensitive to that level of sRNA (Fig. 7L). Consistent with this, arbitrary regulatory regimes (e.g. ≥ 40% on-target repression and ≤ 2% off-target repression) were achievable over a wider range of sRNA transcription rates as spRNA transcription increased (Supplementary Fig. S55). These results suggest that spRNAs may improve specificity in circuits where an sRNA has both on-target and off-target interactions.

#### spRNAs may increase robustness to sRNA mutation

Finally, we investigated whether spRNAs could buffer the effects of sRNA mutation. Using IntaRNA ^60^, we predicted binding energies for all 164 possible single-nucleotide mutations in the 20-nt Syn-ChiX-20D binding region and mapped these to mutant-specific effective binding rates for sRNA–mRNA and sRNA–spRNA interactions (Supplementary Note S9.5). Simulations of these mutants showed that, following mutation, GFPmut3 expression remained effectively unchanged in the presence of spRNA, compared with an average ∼10% increase in its absence (Fig. 7M). Variability across mutants was also substantially larger without spRNA (Fig. 7M), a pattern supported by individual mutant trajectories (Supplementary Fig. S56). In most cases, the presence of spRNA increased the stability of gene expression in response to sRNA mutation, and although some mutants showed the opposite, these typically involved small changes in expression (Supplementary Fig. S56E).

This buffering was not restricted to a single input condition. In a 2D sweep over biologically reasonable sRNA and spRNA transcription rates, increasing spRNA transcription strengthened mutational buffering, particularly at higher sRNA transcription rates, where the mean normalised change in expression decreased from ∼0.25 without spRNA to near zero at high spRNA expression (Fig. 7N). These results suggest that, under some conditions, spRNAs may increase robustness to sRNA mutation and thereby facilitate evolutionary exploration of new sRNA interactions. This conclusion should, however, be interpreted cautiously, because it depends on the assumed relationship between predicted binding energy and effective binding rate, and because the analysis is restricted to mutations in the binding region of the sRNA of this specific circuit architecture (see Supplementary Note S9.5.1). Nevertheless, these simulations highlight a plausible evolutionary advantage of spRNAs.

## Discussion

In this work, we designed, built, and quantitatively characterised synthetic sponge RNA (spRNA) gene circuits in *E. coli*. We established design approaches for reprogramming natural sRNA–spRNA pairs and for reprogramming synthetic sRNAs to function as spRNAs, expanding the synthetic biology toolbox with new ChiX-based sRNA scaffolds and the first synthetic implementations of spRNAs. Experimentally, we showed that synthetic spRNAs can fine-tune gene expression and can be extended to regulate multiple target genes. Through mechanistic modelling, we suggest that spRNAs may reshape response functions and tune effective cooperativity, leakiness, dynamic range, response timing, and potentially evolutionary stability, specificity, and disturbance rejection.

A central contribution of this work is the integration of experimental characterisation with quantitative model development. Our modelling approach progressed from a simple mass-action annihilation model widely used for sRNA circuits ^32,53^ to a final resource-aware model with explicit complex formation. This development was driven by clear mismatches between candidate model predictions and experimentally observed dynamics, while being guided by biochemically plausible mechanisms ^111,112^. In particular, we incorporated resource competition for Hfq, an sRNA chaperone, using a previously published quasi-steady-state modelling framework ^102^, and included explicit complex formation and translation from sRNA–spRNA complexes. The resulting model reproduced behaviour in promoter constructs, sponge circuits, and sRNA circuits, capturing key qualitative behaviours across replicates in data both seen and unseen during fitting. Without this combined modelling and experimental approach, the role of resource competition in explaining non-linear sponge-mediated regulation would likely have been missed, potentially leading to erroneous conclusions about spRNA function or the stoichiometric relationship between sRNA and spRNA. Despite strong evidence for the importance of Hfq competition in natural sRNA networks ^80,85,111,112^, previous modelling of synthetic sRNA circuits has often not incorporated Hfq-based resource competition ^32,53,57^, perhaps because it was not required to explain systems containing only a single synthetic sRNA.

In addition to Hfq-based competition, our results highlight the importance of other forms of host–circuit coupling, namely growth feedback, when interpreting the apparent regulatory performance of synthetic sRNAs and spRNAs. The stronger apparent sponge-mediated recovery observed in the mScarlet-I reporter circuit could not be attributed solely to the transcript-sequence change caused by inclusion of downstream mScarlet-I. Instead, experimental perturbations and modelling support a combined explanation involving both improved spRNA stability and burden-mediated growth feedback, whereby slowed growth reduces dilution and increases intracellular protein concentrations. As seen in the mScarlet-I reporter circuit, this growth feedback can easily be misinterpreted as improved regulator performance. This finding reinforces the importance of measuring and accounting for growth rate during circuit characterisation, especially when comparing architectures with different expression burdens ^113^.

These findings were enabled by a quantitative experimental framework as well as by the RNA parts themselves. Absolute protein quantification ^88^ allowed model parameters to be defined in absolute concentration units, making predictions of hypothetical circuit performance more interpretable and more useful for future engineering^90^. Frequent time-resolved measurements in a small-scale turbidostat enabled robust measurement of both transient dynamics and steady states in response to sequential inducer step changes under approximately constant environmental conditions ^82,83^. Our growth-rate estimator further linked growth-rate decreases following induction of circuit components to changes in intracellular protein levels. Given the importance of growth rate for circuit behaviour, the value of maintaining a constant culture environment becomes particularly clear: nutrient depletion in batch culture is known to generate complex emergent phenomena such as oscillations ^104–106^, which could easily obscure the dynamics caused by spRNA action. Together with standardised promoter constructs, these choices improved parameter identifiability, strengthened comparability across experiments, and made the resulting simulations more effective for model-guided circuit design. In this sense, the work contributes not only new RNA parts and circuit behaviours, but also a framework for quantitatively engineering RNA regulation.

Several findings from our modelling have immediate implications for both synthetic circuit design and natural spRNA function. First, spRNAs can act as more than simple “off-switches” for sRNAs: they can meaningfully reshape response functions, including modulating effective cooperativity. This suggests plausible uses of sRNA–spRNA modules as classifier-like elements in synthetic circuits or as post-transcriptional tuning layers for dynamic-range matching in multi-layered systems. Several of our simulations show that spRNAs can introduce switch-like thresholds while also mitigating leaky sRNA expression, where promoter leakiness can have disproportionately large effects on gene output. Both observations point to likely utility of spRNAs in natural gene regulatory networks, where transcripts may have been co-opted to provide these functions cheaply. Second, our investigation of response times indicates that spRNAs do not universally accelerate or slow responses. However, when de-repression of an sRNA-regulated system is required, spRNAs may provide a practical route to faster responses, which is intuitively consistent with the often relatively stable nature of sRNAs ^110^ when compared to mRNAs. In synthetic systems, these benefits may be especially relevant in bioreactor settings where removal of chemical inducers is challenging. Third, our simulations suggest that spRNAs can improve robustness through both mutation buffering and increased specificity. In natural systems, where sRNA binding is often weaker and more promiscuous than in engineered systems ^38,45,114^, the specificity benefits of spRNAs may be particularly important for maintaining intended regulation, while mutation buffering may promote both evolutionary stability and exploration of new targets through sRNA mutation. More broadly, our results highlight the need to understand and exploit both the molecular mechanism of sRNA or spRNA activity (predation versus annihilation) and the resource competition imposed by sRNA or spRNA expression when engineering predictable input–output behaviour in RNA-based gene circuits.

Our work also points to several directions for future research. Constructing and investigating larger libraries of spRNAs across scaffolds, targets, growth conditions, and bacterial species would help clarify design rules and portability, in line with similar studies for sRNAs ^29,30,35,66^. As additional natural spRNAs are identified ^69^, improved scaffolds and new design principles may emerge. Our synthetic circuits are composed of individual inducible transcription units, whereas in natural systems spRNAs are often produced by more complex mechanisms, including RNA processing ^72^, and potentially premature transcription termination and antisense transcription ^69^ (Fig. 1D). Engineering these production modes could enable ultra-compact synthetic gene circuits in which a target gene and two regulators (sRNA and spRNA) are generated from a single locus. Our demonstration of the importance of Hfq-based resource competition also suggests that further work should explore engineering sRNA and spRNA Hfq-binding scaffolds^115^, as well as tuning Hfq expression, in line with recent work modulating gene regulation through Hfq levels ^116^. Several of our hypotheses could also be tested further through stochastic modelling, long-term evolution experiments ^117^, and improved direct observation of RNA abundance and complex formation ^118,119^.

Small RNAs have recently been identified in lambda bacteriophage, where they enhance lytic infection by regulating *E. coli* and phage transcripts ^120^, while some *E. coli* prophage RNAs act as sponges for chromosomally encoded sRNAs ^121^. Bacteriophage may therefore use both sRNAs and spRNAs against bacterial hosts, raising the possibility of evolutionary arms races between bacterial and phage regulatory RNAs, and suggesting that further small RNA and sponge RNA biology may remain to be discovered in bacteriophage. In synthetic systems, this points to the possibility of using bacteriophage to deliver sRNAs or spRNAs to bacteria in applications such as phage therapy. A classic application of sRNAs has been high-throughput reprogramming of host and pathway gene expression using synthetic sRNA libraries for metabolic engineering ^28,122^; similarly, synthetic spRNA libraries could be used for high-throughput reprogramming of endogenous sRNA activity. A further application of sRNAs has been in feedback circuits^32^, and recent theoretical work on new antithetic feedback architectures ^123^ and multi-target feedback circuits ^124^ may lend themselves to implementation using sRNAs together with cognate spRNAs.

Overall, this study establishes synthetic sponge RNAs as a new tool for engineering bacterial gene circuits with post-transcriptional regulation. By combining rigorous continuous-culture characterisation, absolute quantification, and mechanistic, resource-aware quantitative modelling, we demonstrate the importance of resource competition in bacterial RNA-based circuits and provide a framework for understanding how spRNAs can shape gene-regulatory behaviour in both synthetic and natural systems. We anticipate that our spRNA design framework and quantitative investigation will support broader use of RNA-based regulation in bacterial synthetic biology, enabling more predictable, tuneable, and robust gene circuits.

## Author contributions statement

S.B.S., H.S., and A.P. conceived the project. H.S. and A.P. supervised the project. S.B.S. designed regulatory RNAs and gene circuits. S.B.S. performed all experiments. S.B.S. and K.S. performed mathematical modelling. S.B.S. analysed results. M.C. performed growth-rate estimation. S.B.S. wrote the manuscript. All authors reviewed, edited, and approved the final manuscript.

## Acknowledgments

S.B.S., M.C., H.S., and A.P. are in part supported by funding from the Engineering and Physical Sciences Research Council under the EEBio Programme Grant (UKRI-EPSRC; grant number EP/Y014073/1). S.B.S. and M.C. were supported by funding from the Biotechnology and Biological Sciences Research Council (UKRI-BBSRC; grant number BB/T008784/1). The EcoFlex kit was a kind gift from Paul Freemont (Addgene kit #1000000080). The Marionette Sensor Collection was a kind gift from Christopher Voigt (Addgene Kit #1000000137). The authors thank Eszter Csibra for advice on implementation of FPCountR and Claudia Driscoll for assistance with figure design.

## References

[1] Department for Science, Innovation & Technology. National Vision for Engineering Biology. Technical report, 2023.

[2] Government Office for Science. Engineering Biology Aspirations. Technical report, 2025.

[3] Carlos Barajas and Domitilla Del Vecchio. Synthetic biology by controller design. Current Opinion in Biotechnology, 2022.

[4] Shalni Kumar and Jeff Hasty. Stability, robustness, and containment: preparing synthetic biology for real-world deployment. Current Opinion in Biotechnology, 79:102880, 2023.

[5] Maurice Filo, Ching-Hsiang Chang, and Mustafa Khammash. Biomolecular feedback controllers: from theory to applications. Current Opinion in Biotechnology, 79:102882, 2023.

[6] Michael B. Elowitz and Stanislas Leibler. A synthetic oscillatory network of transcriptional regulators. Nature, 403(6767):335–338, January 2000.

[7] Laurent Potvin-Trottier, Nathan D. Lord, Glenn Vinnicombe, and Johan Paulsson. Synchronous long-term oscillations in a synthetic gene circuit. Nature, 538(7626):514–517, October 2016.

[8] Timothy S. Gardner, Charles R. Cantor, and James J. Collins. Construction of a genetic toggle switch in Escherichia coli. Nature, 403(6767):339–342, January 2000.

[9] A. A. K. Nielsen, B. S. Der, J. Shin, P. Vaidyanathan, V. Paralanov, E. A. Strychalski, D. Ross, D. Densmore, and C. A. Voigt. Genetic circuit design automation. Science, 352(6281):aac7341–aac7341, April 2016.

[10] Corentin Briat, Ankit Gupta, and Mustafa Khammash. Antithetic Integral Feedback Ensures Robust Perfect Adaptation in Noisy Biomolecular Networks. Cell Systems, 2(1):15–26, January 2016.

[11] Adrian L. Slusarczyk, Allen Lin, and Ron Weiss. Foundations for the design and implementation of synthetic genetic circuits. Nature Reviews Genetics, 13(6):406–420, June 2012.

[12] Jennifer A N Brophy and Christopher A Voigt. Principles of genetic circuit design. Nature Methods, 11(5):508–520, May 2014.

[13] Julius B. Lucks, Lei Qi, Vivek K. Mutalik, Denise Wang, and Adam P. Arkin. Versatile RNA-sensing transcriptional regulators for engineering genetic networks. Proceedings of the National Academy of Sciences, 108(21):8617–8622, May 2011.

[14] Zhen Xie, Liliana Wroblewska, Laura Prochazka, Ron Weiss, and Yaakov Benenson. Multi-Input RNAi-Based Logic Circuit for Identification of Specific Cancer Cells. Science, 333(6047):1307–1311, September 2011.

[15] Alexander A. Green, Pamela A. Silver, James J. Collins, and Peng Yin. Toehold Switches: De-Novo-Designed Regulators of Gene Expression. Cell, 159(4):925–939, November 2014.

[16] James Chappell, Kyle E Watters, Melissa K Takahashi, and Julius B Lucks. A renaissance in RNA synthetic biology: new mechanisms, applications and tools for the future. Current Opinion in Chemical Biology, 28:47–56, October 2015.

[17] James Chappell, Melissa K Takahashi, and Julius B Lucks. Creating small transcription activating RNAs. Nature Chemical Biology, 11(3):214–220, March 2015.

[18] James Chappell, Alexandra Westbrook, Matthew Verosloff, and Julius B. Lucks. Computational design of small transcription activating RNAs for versatile and dynamic gene regulation. Nature Communications, 8(1):1051, October 2017.

[19] Nicolaas M. Angenent-Mari, Alexander S. Garruss, Luis R. Soenksen, George Church, and James J. Collins. A deep learning approach to programmable RNA switches. Nature Communications, 11(1):5057, October 2020.

[20] Qi Zhang, Duo Ma, Fuqing Wu, Kylie Standage-Beier, Xingwen Chen, Kaiyue Wu, Alexander A. Green, and Xiao Wang. Predictable control of RNA lifetime using engineered degradation-tuning RNAs. Nature Chemical Biology, 17(7):828–836, 2021.

[21] F. Veronica Greco, Amir Pandi, Tobias J. Erb, Claire S. Grierson, and Thomas E. Gorochowski. Harnessing the central dogma for stringent multi-level control of gene expression. Nature Communications, 12(1):1738, 2021.

[22] Benjamin I. Tickman, Diego Alba Burbano, Venkata P. Chavali, Cholpisit Kiattisewee, Jason Fontana, Aset Khakimzhan, Vincent Noireaux, Jesse G. Zalatan, and James M. Carothers. Multi-layer CRISPRa/i circuits for dynamic genetic programs in cell-free and bacterial systems. Cell Systems, 13(3):215–229.e8, March 2022.

[23] Içvara Barbier, Hadiastri Kusumawardhani, Lakshya Chauhan, Pradyumna Vinod Harlapur, Mohit Kumar Jolly, and Yolanda Schaerli. Synthetic Gene Circuits Combining CRISPR Interference and CRISPR Activation in E. coli : Importance of Equal Guide RNA Binding Affinities to Avoid Context-Dependent Effects. ACS Synthetic Biology, 12(10):3064–3071, October 2023.

[24] Kasey S. Love, Christopher P. Johnstone, Emma L. Peterman, Stephanie Gaglione, Michael E. Birnbaum, and Kate E. Galloway. Model-guided design of microRNA-based gene circuits supports precise dosage of transgenic cargoes into diverse primary cells. Cell Systems, 16(6):101269, June 2025.

[25] Alice Boo, Harman Mehta, Rodrigo Ledesma-Amaro, and Guy-Bart Stan. Burden-aware feedback control of microbial consortia. Nature Communications, May 2026.

[26] Rongrong Du, Michael J. Flynn, Karan Mahe, Monique Honsa, Bo Gu, Dongyang Li, Sean E. McGeary, Viviana Gradinaru, Ralf Jungmann, and Michael B. Elowitz. miRNA modules for precise, tunable control of gene expression. Molecular Cell, 86(1):194–212.e7, January 2026.

[27] Guillermo Rodrigo, Thomas E. Landrain, and Alfonso Jaramillo. De novo automated design of small RNA circuits for engineering synthetic riboregulation in living cells. Proceedings of the National Academy of Sciences, 109(38):15271–15276, September 2012.

[28] Dokyun Na, Seung Min Yoo, Hannah Chung, Hyegwon Park, Jin Hwan Park, and Sang Yup Lee. Metabolic engineering of Escherichia coli using synthetic small regulatory RNAs. Nature Biotechnology, 31(2):170–174, 2013.

[29] Seung Min Yoo, Dokyun Na, and Sang Yup Lee. Design and use of synthetic regulatory small RNAs to control gene expression in Escherichia coli. Nature Protocols, 8(9):1694–1707, 2013.

[30] Allison Hoynes-O’Connor and Tae Seok Moon. Development of Design Rules for Reliable Antisense RNA Behavior in E. coli. ACS Synthetic Biology, 5(12):1441–1454, December 2016.

[31] Deepak K. Agrawal, Xun Tang, Alexandra Westbrook, Ryan Marshall, Colin S. Maxwell, Julius Lucks, Vincent Noireaux, Chase L. Beisel, Mary J. Dunlop, and Elisa Franco. Mathematical Modeling of RNA-Based Architectures for Closed Loop Control of Gene Expression. ACS Synthetic Biology, 7(5):1219–1228, May 2018.

[32] Ciarán L Kelly, Andreas W K Harris, Harrison Steel, Edward J Hancock, John T Heap, and Antonis Papachristodoulou. Synthetic negative feedback circuits using engineered small RNAs. Nucleic Acids Research, 46(18):9875–9889, 2018.

[33] Hsin-Ho Huang, Yili Qian, and Domitilla Del Vecchio. A quasi-integral controller for adaptation of genetic modules to variable ribosome demand. Nature Communications, 9(1):5415, December 2018.

[34] Nicolai Engelmann, Maik Molderings, and Heinz Koeppl. Tuning Ultrasensitivity in Genetic Logic Gates using Antisense RNA Feedback, July 2024.

[35] Jun Ren, Nuong Thi Nong, Phuong N. Lam Vo, Hyang-Mi Lee, and Dokyun Na. Rational Design of High-Efficiency Synthetic Small Regulatory RNAs and Their Application in Robust Genetic Circuit Performance Through Tight Control of Leaky Gene Expression. ACS Synthetic Biology, 13(10):3256–3267, October 2024.

[36] Jens Georg, Bork A. Berghoff, and Daniel Schindler. Harnessing Small RNAs as Synthetic Post-transcriptional Regulators in Bacteria. ACS Synthetic Biology, 14(7):2405–2417, July 2025.

[37] Daniel Kelvin, Erik Kubaczka, Marianna Karava, Heinz Koeppl, and Beatrix Suess. Iterative design of a NAND hybrid riboswitch by deep batch Bayesian optimization. Nucleic Acids Research, 54(5):gkag145, February 2026.

[38] Lauren S. Waters and Gisela Storz. Regulatory RNAs in Bacteria. Cell, 136(4):615–628, February 2009.

[39] S. Gottesman and G. Storz. Bacterial Small RNA Regulators: Versatile Roles and Rapidly Evolving Variations. Cold Spring Harbor Perspectives in Biology, 3(12):a003798–a003798, December 2011.

[40] Susan Gottesman. Bacterial Regulatory Circuits are Linked and Extended by Small RNAs. J. Mol. Biol., 437(11):169059, 2025.

[41] Kathrin S. Fröhlich and Kai Papenfort. Regulation outside the box: New mechanisms for small RNAs. Molecular Microbiology, 114(3):363–366, September 2020.

[42] Nicholas De Lay, Daniel J. Schu, and Susan Gottesman. Bacterial Small RNA-based Negative Regulation: Hfq and Its Accomplices. Journal of Biological Chemistry, 288(12):7996–8003, March 2013.

[43] Pierre Mandin and Susan Gottesman. A genetic approach for finding small RNAs regulators of genes of interest identifies RybC as regulating the DpiA/DpiB two-component system. Molecular Microbiology, 72(3):551–565, May 2009.

[44] Jörg Vogel and Ben F. Luisi. Hfq and its constellation of RNA. Nature Reviews Microbiology, 9(8):578–589, August 2011.

[45] Gisela Storz, Jörg Vogel, and Karen M. Wassarman. Regulation by Small RNAs in Bacteria: Expanding Frontiers. Molecular Cell, 43(6):880–891, September 2011.

[46] Takeshi Mizuno, Mei-Yin Chou, and Masayori Inouye. A unique mechanism regulating gene expression: translational inhibition by a complementary RNA transcript (micRNA). Proceedings of the National Academy of Sciences, 81:1966–1970, April 1984.

[47] Nadim Majdalani, Christofer Cunning, Darren Sledjeski, Tom Elliott, and Susan Gottesman. DsrA RNA regulates translation of RpoS message by an anti-antisense mechanism, independent of its action as an antisilencer of transcription. Proceedings of the National Academy of Sciences, 95(21):12462–12467, October 1998.

[48] Eric Massé, Freddy E. Escorcia, and Susan Gottesman. Coupled degradation of a small regulatory RNA and its mRNA targets in Escherichia coli. Genes & Development, 17(19):2374–2383, October 2003.

[49] Kathrin Sophie Fröhlich, Kai Papenfort, Agnes Fekete, and Jörg Vogel. A small RNA activates CFA synthase by isoform-specific mRNA stabilization. The EMBO Journal, 32(22):2963–2979, November 2013.

[50] Svetlana Durica-Mitic*, Yvonne Göpel*, and Boris Görke. Carbohydrate Utilization in Bacteria: Making the Most Out of Sugars with the Help of Small Regulatory RNAs. Microbiology Spectrum, 6(2):6.2.07, April 2018.

[51] Erik Holmqvist and E. Gerhart H. Wagner. Impact of bacterial sRNAs in stress responses. Biochemical Society Transactions, 45(6):1203–1212, December 2017.

[52] Jacob R. Chambers and Karin Sauer. Small RNAs and their role in biofilm formation. Trends in Microbiology, 21(1):39–49, January 2013.

[53] Erel Levine, Zhongge Zhang, Thomas Kuhlman, and Terence Hwa. Quantitative Characteristics of Gene Regulation by Small RNA. PLoS Biology, 5(9):e229, 2007.

[54] Pankaj Mehta, Sidhartha Goyal, and Ned S Wingreen. A quantitative comparison of sRNA-based and protein-based gene regulation. Molecular Systems Biology, 4(1):221, 2008.

[55] Erel Levine, Min Huang, Yingwu Huang, Thomas Kuhlman, Hualin Shi, Zhongge Zhang, and Terence Hwa. On noise and silence in small RNA regulation. Preprint, 2007.

[56] Ya Jia, Wangheng Liu, Anbang Li, Lijian Yang, and Xuan Zhan. Intrinsic noise in post-transcriptional gene regulation by small non-coding RNA. Biophysical Chemistry, 143(1–2):60–69, July 2009.

[57] Yishai Shimoni, Gilgi Friedlander, Guy Hetzroni, Gali Niv, Shoshy Altuvia, Ofer Biham, and Hanah Margalit. Regulation of gene expression by small non-coding RNAs: a quantitative view. Molecular Systems Biology, 3(1):138, January 2007.

[58] Michael Zuker. Mfold web server for nucleic acid folding and hybridization prediction. Nucleic Acids Research, 31(13):3406–3415, 2003.

[59] Ronny Lorenz, Stephan H. Bernhart, Christian Höner zu Siederdissen, Hakim Tafer, Christoph Flamm, Peter F. Stadler, and Ivo L. Hofacker. ViennaRNA Package 2.0. Algorithms for Molecular Biology, 6(1):26, 2011.

[60] Martin Mann, Patrick R. Wright, and Rolf Backofen. IntaRNA 2.0: enhanced and customizable prediction of RNA–RNA interactions. Nucleic Acids Research, 45(W1):W435–W439, 2017.

[61] Jinho Yeom, Jong Seong Park, Yong Min Jeon, Beom Seop Song, and Seung Min Yoo. Synthetic fused sRNA for the simultaneous repression of multiple genes. Applied Microbiology and Biotechnology, 106(7):2517–2527, 2022.

[62] Young Je Lee and Tae Seok Moon. Design rules of synthetic non-coding RNAs in bacteria. Methods, 143:58–69, July 2018.

[63] Minho Noh, Seung Min Yoo, Dongsoo Yang, and Sang Yup Lee. Broad-Spectrum Gene Repression Using Scaffold Engineering of Synthetic sRNAs. ACS Synthetic Biology, 8(6):1452–1461, June 2019.

[64] Jae Sung Cho, Dongsoo Yang, Cindy Pricilia Surya Prabowo, Mohammad Rifqi Ghiffary, Taehee Han, Kyeong Rok Choi, Cheon Woo Moon, Hengrui Zhou, Jae Yong Ryu, Hyun Uk Kim, and Sang Yup Lee. Targeted and high-throughput gene knockdown in diverse bacteria using synthetic sRNAs. Nature Communications, 14(1):2359, April 2023.

[65] Yang Wang, Guobin Yin, Huanjiao Weng, Luyao Zhang, Guocheng Du, Jian Chen, and Zhen Kang. Gene knockdown by structure defined single-stem loop small non-coding RNAs with programmable regulatory activities. Synthetic and Systems Biotechnology, 8(1):86–96, March 2023.

[66] Michel Brück, Tania S. Köbel, Sophie Dittmar, Adán A. Ramírez Rojas, Jens Georg, Bork A. Berghoff, and Daniel Schindler. A library-based approach allows systematic and rapid evaluation of seed region length and reveals design rules for synthetic bacterial small RNAs. iScience, 27(9):110774, September 2024.

[67] Katherine McJunkin and Susan Gottesman. What goes up must come down: off switches for regulatory RNAs. Genes & Development, 38(13–14):597–613, July 2024.

[68] Nara Figueroa-Bossi and Lionello Bossi. Sponges and Predators in the Small RNA World. Microbiology Spectrum, 6(4), July 2018.

[69] Emma L. Denham. The Sponge RNAs of bacteria – How to find them and their role in regulating the post-transcriptional network. Biochimica et Biophysica Acta (BBA) - Gene Regulatory Mechanisms, 1863(8):194565, August 2020.

[70] Laura N Vogt and Kathrin S Fröhlich. Impact of Small RNA Sponges on Regulatory RNA Networks in Bacteria. Annu. Rev. Genet., 59:69–89, 2025.

[71] Martin Overgaard, Jesper Johansen, Jakob Møller-Jensen, and Poul Valentin-Hansen. Switching off small RNA regulation with trap-mRNA. Molecular Microbiology, 73(5):790–800, 2009.

[72] Masatoshi Miyakoshi, Yanjie Chao, and Jörg Vogel. Cross talk between ABC transporter mRNAs via a target mRNA-derived sponge of the GcvB small RNA. The EMBO Journal, 34(11):1478–1492, June 2015.

[73] David Lalaouna, Marie-Claude Carrier, Szabolcs Semsey, Jean-Simon Brouard, Jing Wang, Joseph T. Wade, and Eric Massé. A 3′ External Transcribed Spacer in a tRNA Transcript Acts as a Sponge for Small RNAs to Prevent Transcriptional Noise. Molecular Cell, 58(3):393–405, May 2015.

[74] Nara Figueroa-Bossi, Martina Valentini, Laurette Malleret, and Lionello Bossi. Caught at its own game: regulatory small RNA inactivated by an inducible transcript mimicking its target. Genes & Development, 23:2004–2015, 2009.

[75] Jacqueline Plumbridge, Lionello Bossi, Jacques Oberto, Joseph T. Wade, and Nara Figueroa-Bossi. Interplay of transcriptional and small RNA-dependent control mechanisms regulates chitosugar uptake in E scherichia coli and S almonella: Coregulation of a sugar porin and PTS transporter. Molecular Microbiology, 92(4):648–658, 2014.

[76] Lillian G. Acuña, M. José Barros Diego Peñaloza, Paula I. Rodas, Daniel Paredes-Sabja, Juan A. Fuentes, Fernando Gil, and Iván L. Calderón. A feed-forward loop between SroC and MgrR small RNAs modulates the expression of eptB and the susceptibility to polymyxin B in Salmonella Typhimurium. Microbiology, 162(11):1996–2004, November 2016.

[77] Philip P Adams, Gabriele Baniulyte, Caroline Esnault, Kavya Chegireddy, Navjot Singh, Molly Monge, Ryan K Dale, Gisela Storz, and Joseph T Wade. Regulatory roles of Escherichia coli 5’ UTR and ORF-internal RNAs detected by 3’ end mapping. eLife, 10:e62438, 2021.

[78] Sylvain Durand, Adam Callan-Sidat, Josie McKeown, Stephen Li, Gergana Kostova, Juan R Hernandez-Fernaud, Mohammad Tauqeer Alam, Andrew Millard, Delphine Allouche, Chrystala Constantinidou, Ciarán Condon, and Emma L Denham. Identification of an RNA sponge that controls the RoxS riboregulator of central metabolism in Bacillus subtilis. Nucleic Acids Research, 49(11):6399–6419, June 2021.

[79] Jens Hör, Stanislaw A. Gorski, and Jörg Vogel. Bacterial RNA Biology on a Genome Scale. Molecular Cell, 70(5):785–799, June 2018.

[80] Jiandong Chen, Leann To, Francois de Mets, Xing Luo, Nadim Majdalani, Chin-Hsien Tai, and Susan Gottesman. A fluorescence-based genetic screen reveals diverse mechanisms silencing small RNA signaling in E. coli. PNAS, 118(27):e2106964118, 2021.

[81] Caleb Weinreb, Samuel Wolock, Betsabeh K. Tusi, Merav Socolovsky, and Allon M. Klein. Fundamental limits on dynamic inference from single-cell snapshots. Proc. Natl. Acad. Sci. U.S.A., 115(10):E2467–E2476, 2018.

[82] Harrison Steel, Robert Habgood, Ciarán L. Kelly, and Antonis Papachristodoulou. In situ characterisation and manipulation of biological systems with Chi.Bio. PLOS Biology, 18(7):e3000794, 2020.

[83] Shivang Hina-Nilesh Joshi, Chentao Yong, and Andras Gyorgy. Inducible plasmid copy number control for synthetic biology in commonly used e. coli strains. Nat. Commun., 13(6691):6691, 2022.

[84] Kirill Sechkar, Harrison Steel, Giansimone Perrino, and Guy-Bart Stan. A coarse-grained bacterial cell model for resource-aware analysis and design of synthetic gene circuits. Nature Communications, 15(1):1981, 2024.

[85] Kyung Moon and Susan Gottesman. Competition among Hfq-binding small RNAs in Escherichia coli. Molecular Microbiology, 82(6):1545–1562, 2011.

[86] Michael J. Ellis, Ryan S. Trussler, and David B. Haniford. Hfq binds directly to the ribosome-binding site of IS10 transposase mRNA to inhibit translation. Molecular Microbiology, 96(3):633–650, 2015.

[87] Jorjethe Roca, Andrew Santiago-Frangos, and Sarah A. Woodson. Diversity of bacterial small RNAs drives competitive strategies for a mutual chaperone. Nature Communications, 13(1):2449, May 2022.

[88] Eszter Csibra and Guy-Bart Stan. Absolute protein quantification using fluorescence measurements with FPCountR. Nature Communications, 13(1):6600, 2022.

[89] Eszter Csibra and Guy-Bart Stan. FPCount protocol - in-lysate (purification free) protocol. 2023.

[90] Scott B. Stacey, Ting An Lee, Olivia Gallupova, Eszter Csibra, Antonis Papachristodoulou, Harrison Steel, and Kirill Sechkar. Characterisation of synthetic gene circuits with absolute quantification in continuous culture. In Xiaojun Tian, editor, In Press. Springer US, 2025.

[91] Anders Aamann Rasmussen, Jesper Johansen, Jesper S. Nielsen, Martin Overgaard, Birgitte Kallipolitis, and Poul Valentin-Hansen. A conserved small RNA promotes silencing of the outer membrane protein YbfM. Molecular Microbiology, 72(3):566–577, May 2009.

[92] Adam J. Meyer, Thomas H. Segall-Shapiro, Emerson Glassey, Jing Zhang, and Christopher A. Voigt. Escherichia coli “Marionette” strains with 12 highly optimized small-molecule sensors. Nature Chemical Biology, 15(2):196–204, 2019.

[93] Martin Raden, Syed M Ali, Omer S Alkhnbashi, Anke Busch, Fabrizio Costa, Jason A Davis, Florian Eggenhofer, Rick Gelhausen, Jens Georg, Steffen Heyne, Michael Hiller, Kousik Kundu, Robert Kleinkauf, Steffen C Lott, Mostafa M Mohamed, Alexander Mattheis, Milad Miladi, Andreas S Richter, Sebastian Will, Joachim Wolff, Patrick R Wright, and Rolf Backofen. Freiburg RNA tools: a central online resource for RNA-focused research and teaching. Nucleic Acids Research, 46(W1):W25–W29, 2018.

[94] Philip Z Johnson and Anne E Simon. RNAcanvas: interactive drawing and exploration of nucleic acid structures. Nucleic Acids Research, 51(W1):W501–W508, 2023.

[95] Simon J. Moore, Hung-En Lai, Richard J. R. Kelwick, Soo Mei Chee, David J. Bell, Karen Marie Polizzi, and Paul S. Freemont. EcoFlex: A Multifunctional MoClo Kit for E. coli Synthetic Biology. ACS Synthetic Biology, 5(10):1059–1069, 2016.

[96] Marco Corrao. GOFFREDOpy: probabilistic tracking of growth rate and OD from turbidostat data. Available at https://github.com/marco-corrao/GOFFREDOpy, 2026.

[97] Eszter Csibra. Fpcountr: Fluorescent protein calibration for plate readers. r package, 2021.

[98] R Core Team. R: A Language and Environment for Statistical Computing. R Foundation for Statistical Computing, Vienna, Austria, 2020.

[99] Hasan Baig, Pedro Fontanarossa, James McLaughlin, James Scott-Brown, Prashant Vaidyanathan, Thomas Gorochowski, Goksel Misirli, Jacob Beal, and Chris Myers. Synthetic biology open language visual (SBOL visual) version 3.0. Journal of Integrative Bioinformatics, 18(3), 2021.

[100] Yasushi Ishihama, Thorsten Schmidt, Juri Rappsilber, Matthias Mann, F Ulrich Hartl, Michael J Kerner, and Dmitrij Frishman. Protein abundance profiling of the Escherichia coli cytosol. BMC Genomics, 9(1):102, 2008.

[101] Yili Qian, Hsin-Ho Huang, José I. Jiménez, and Domitilla Del Vecchio. Resource Competition Shapes the Response of Genetic Circuits. ACS Synthetic Biology, 6(7):1263–1272, July 2017.

[102] Armin M. Zand, Stanislav Anastassov, Timothy Frei, and Mustafa Khammash. Multi-Layer Autocatalytic Feedback Enables Integral Control Amidst Resource Competition and Across Scales. ACS Synth. Biol., 14(4):1041–1061, 2025.

[103] Stefan Klumpp and Terence Hwa. Bacterial growth: global effects on gene expression, growth feedback and proteome partition. Curr. Opin. Biotechnol., 28:96–102, 2014.

[104] Cheemeng Tan, Philippe Marguet, and Lingchong You. Emergent bistability by a growth-modulating positive feedback circuit. Nat. Chem. Biol., 5:842–848, 2009.

[105] Juan Melendez-Alvarez, Changhan He, Rong Zhang, Yang Kuang, and Xiao-Jun Tian. Emergent damped oscillation induced by nutrient-modulating growth feedback. ACS Synthetic Biology, 10(5):1227–1236, 2021.

[106] Dylan Hirsch and Domitilla Del Vecchio. Differential equation model for the population-level dynamics of a toggle switch with growth-feedback. In 2022 IEEE 61st Conference on Decision and Control (CDC), pages 06–09. IEEE.

[107] Kalen P. Clifton, Ethan M. Jones, Sudip Paudel, John P. Marken, Callan E. Monette, Andrew D. Halleran, Lidia Epp, and Margaret S. Saha. The genetic insulator RiboJ increases expression of insulated genes. Journal of Biological Engineering, 12(1):23, October 2018.

[108] Masatoshi Miyakoshi, Haruna Okayama, Maxence Lejars, Takeshi Kanda, Yuki Tanaka, Kaori Itaya, Miki Okuno, Takehiko Itoh, Noritaka Iwai, and Masaaki Wachi. Mining RNA-seq data reveals the massive regulon of GcvB small RNA and its physiological significance in maintaining amino acid homeostasis in Escherichia coli. Molecular Microbiology, 117(1):160–178, January 2022.

[109] Razika Hussein and Han N. Lim. Direct comparison of small RNA and transcription factor signaling. Nucleic Acids Research, 40(15):7269–7279, 2012.

[110] Jörg Vogel, Verena Bartels, Thean Hock Tang, Gennady Churakov, Jacoba G. Slagter-Jäger, Alexander Hüttenhofer, and E. Gerhart H. Wagner. RNomics in Escherichia coli detects new sRNA species and indicates parallel transcriptional output in bacteria. Nucleic Acids Research, 31(22):6435–6443, November 2003.

[111] R. Hussein and H. N. Lim. Disruption of small RNA signaling caused by competition for Hfq. Proceedings of the National Academy of Sciences, 108(3):1110–1115, January 2011.

[112] Raya Faigenbaum-Romm, Avichai Reich, Yair E. Gatt, Meshi Barsheshet, Liron Argaman, and Hanah Margalit. Hierarchy in Hfq Chaperon Occupancy of Small RNA Targets Plays a Major Role in Their Regulation. Cell Reports, 30(9):3127–3138.e6, March 2020.

[113] Francesca Ceroni, Rhys Algar, Guy-Bart Stan, and Tom Ellis. Quantifying cellular capacity identifies gene expression designs with reduced burden. Nature Methods, 12(5):415–418, May 2015.

[114] Chase L. Beisel and Gisela Storz. The Base-Pairing RNA Spot 42 Participates in a Multioutput Feedforward Loop to Help Enact Catabolite Repression in Escherichia coli. Molecular Cell, 41(3):286–297, February 2011.

[115] Yuta Sakai, Koichi Abe, Saki Nakashima, Wataru Yoshida, Stefano Ferri, Koji Sode, and Kazunori Ikebukuro. Improving the Gene-Regulation Ability of Small RNAs by Scaffold Engineering in Escherichia coli. ACS Synthetic Biology, 3(3):152–162, March 2014.

[116] Yu Jung Jung, Keun Ha Park, Tae Yeong Jang, and Seung Min Yoo. Gene expression regulation by modulating Hfq expression in coordination with tailor-made sRNA-based knockdown in Escherichia coli. Journal of Biotechnology, 388:1–10, June 2024.

[117] R E Lenski and M Travisano. Dynamics of adaptation and diversification: a 10,000-generation experiment with bacterial populations. Proceedings of the National Academy of Sciences, 91(15):6808–6814, July 1994.

[118] Jichuan Zhang, Jingyi Fei, Benjamin J. Leslie, Kyu Young Han, Thomas E. Kuhlman, and Taekjip Ha. Tandem Spinach Array for mRNA Imaging in Living Bacterial Cells. Scientific Reports, 5(1):17295, November 2015.

[119] Khalid K. Alam, Kwaku D. Tawiah, Matthew F. Lichte, David Porciani, and Donald H. Burke. A Fluorescent Split Aptamer for Visualizing RNA–RNA Assembly In Vivo. ACS Synthetic Biology, 6(9):1710–1721, September 2017.

[120] Aviezer Silverman, Raneem Nashef, Reut Bruner, Tamar Noy, and Sahar Melamed. Phage-encoded small RNA hijacks host replication machinery to support the phage lytic cycle, October 2024.

[121] Jai J. Tree, Sander Granneman, Sean P. McAteer, David Tollervey, and David L. Gally. Identification of Bacteriophage-Encoded Anti-sRNAs in Pathogenic Escherichia coli. Molecular Cell, 55(2):199–213, July 2014.

[122] Minho Noh, Seung Min Yoo, Won Jun Kim, and Sang Yup Lee. Gene Expression Knockdown by Modulating Synthetic Small RNA Expression in Escherichia coli. Cell Systems, 5(4):418–426.e4, October 2017.

[123] Maurice Filo, Stephanie K. Aoki, Mucun Hou, Stanislav Anastassov, and Mustafa Khammash. Engineering sensor-based antithetic integral controllers for enhanced dynamic performance and noise attenuation. Cell Systems, 17(3):101512, March 2026.

[124] Suchana Chakravarty, Rong Zhang, and Xiao-Jun Tian. Noise Reduction in Resource-Coupled Multi-Module Gene Circuits through Antithetic Feedback Control. In 2024 IEEE 63rd Conference on Decision and Control (CDC), pages 5566–5571, Milan, Italy, December 2024. IEEE.

[125] Jonathan A. Bernstein, Arkady B. Khodursky, Pei-Hsun Lin, Sue Lin-Chao, and Stanley N. Cohen. Global analysis of mRNA decay and abundance in Escherichia coli at single-gene resolution using two-color fluorescent DNA microarrays. Proceedings of the National Academy of Sciences, 99(15):9697–9702, July 2002.

[126] Enrique Balleza, J. Mark Kim, and Philippe Cluzel. Systematic characterization of maturation time of fluorescent proteins in living cells. Nature Methods, 15(1):47–51, January 2018.

[127] Talley J. Lambert. FPbase: a community-editable fluorescent protein database. Nature Methods, 16(4):277–278, April 2019.

[128] Chunbo Lou, Brynne Stanton, Ying-Ja Chen, Brian Munsky, and Christopher A Voigt. Ribozyme-based insulator parts buffer synthetic circuits from genetic context. Nature Biotechnology, 30(11):1137–1142, November 2012.

[129] Scott A Scholz, Chase D Lindeboom, and Lydia Freddolino. Genetic context effects can override canonical cis regulatory elements in Escherichia coli. Nucleic Acids Research, 50(18):10360–10375, October 2022.

[130] Bin Shao, Jayan Rammohan, Daniel A. Anderson, Nina Alperovich, David Ross, and Christopher A. Voigt. Single-cell measurement of plasmid copy number and promoter activity. Nature Communications, 12(1):1475, December 2021.

[131] Lisa Tietze and Rahmi Lale. Importance of the 5′ regulatory region to bacterial synthetic biology applications. Microbial Biotechnology, 14(6):2291–2315, November 2021.

[132] Esteban Martínez-García, Sofía Fraile, Elena Algar, Tomás Aparicio, Elena Velázquez, Belén Calles, Huseyin Tas, Blas Blázquez, Bruno Martín, Clara Prieto, Lucas Sánchez-Sampedro, Morten H H Nørholm, Daniel C Volke, Nicolas T Wirth, Pavel Dvořák, Lorea Alejaldre, Lewis Grozinger, Matthew Crowther, Angel Goñi-Moreno, Pablo I Nikel, Juan Nogales, and Víctor de Lorenzo. SEVA 4.0: an update of the Standard European Vector Architecture database for advanced analysis and programming of bacterial phenotypes. Nucleic Acids Research, 51(D1):D1558–D1567, January 2023.

[133] Daphne S. Bindels, Lindsay Haarbosch, Laura van Weeren, Marten Postma, Katrin E. Wiese, Marieke Mastop, Sylvain Aumonier, Guillaume Gotthard, Antoine Royant, Mark A. Hink, and Theodorus W. J. Gadella. mScarlet: a bright monomeric red fluorescent protein for cellular imaging. Nature Methods, 14(1):53–56, January 2017.

