## Supplementary Information for "Quantitative Engineering and Investigation of Synthetic Sponge RNAs in *E. coli*"

May 2026

### Contents

**S1 Chi.Bio Experimental Setup** 2

**S2 Part Validation** 3

**S3 Regulatory RNA Design** 5
S3.1 ChiX-Derived Synthetic sRNA Design 5
S3.1.1 Assembly of ChiX-Derived Synthetic sRNA Circuits 6
S3.2 ChbBC-Derived Sponge RNA Design 7
S3.3 MicC-Derived sRNA Design 8
S3.4 Alternative Sponge RNA Design Strategies 9
S3.5 Assembly of Synthetic Sponge RNA Circuits 9
S3.6 Design and Assembly of Multi-Target RNA Circuits 10

**S4 Regulatory RNA Characterisation** 12
S4.1 ChiX-Derived Synthetic sRNAs 12
S4.2 ChbBC-derived synthetic sponge RNAs 13
S4.3 Alternative sponge RNA design strategies characterisation 15

**S5 Regulatory RNA Control Experiments** 17
S5.1 ChiX sRNA Controls 17
S5.1.1 No sRNA Control 17
S5.1.2 Binding Region Control 17
S5.1.3 Off-Target Binding Control 17
S5.2 ChbBC Sponge RNA Controls 18
S5.2.1 ChbBC sponge RNA circuit regulating mScarlet-I 20
S5.3 Multi-target regulatory RNAs 22
S5.3.1 Multi-target small RNAs 22

**S6 Quantitative characterisation: absolute quantification and growth-rate estimation** 22
S6.1 Absolute quantification 22
S6.2 Growth-rate estimation 29

**S7 Model Fitting** 30
S7.1 Promoter response function fitting 30
S7.2 Dataset-specific scaling factors in RNA-circuit model fitting 32
S7.3 Mass-action annihilation-based model 33
S7.4 Incorporating RNA Predation in Models 36
S7.5 Incorporating Resource Competition 39
S7.6 Final model – explicit complexes and $Z$–$S$ complex translation 43
S7.7 Validation and Investigation of Model Issues 48
S7.7.1 Issues with sRNA Circuit Model 48
S7.7.2 Validating Model Against 'Unseen' Data 48

**S8 Growth-feedback investigation** 51

**S9 Additional Details for Quantitative Investigation of Synthetic spRNA Systems Biology** 53
S9.1 Promoter leakiness in sRNA gene expression 53
S9.2 Response-time investigation 53
S9.2.1 Response times for trajectories rising from zero expression 53
S9.2.2 Response times in turbidostat experiments 54
S9.3 Input filtering 56
S9.4 Specificity 58

S9.5 Evolutionary Stability . . . 59
S9.5.1 Limitations to the evolutionary analysis . . . 61

**S10 DNA Sequences and Plasmids** **61**

### S1 Chi.Bio Experimental Setup

Most characterisation experiments in this work were carried out using the Chi.Bio experimental platform for continuous culture [1].

Chi.Bio is a small-scale bioreactor platform that uses 20 mL glass vials with custom printed lids connected to media-in and media-out pumps. The media-in pump draws from bottles of fresh media, while the media-out pump removes culture to waste. The platform controls stirring and temperature and records OD and fluorescence measurements, with OD dithering enabling dynamic growth-rate estimation. The pumps and bioreactor are controlled through a connected computer and graphical user interface. A diagram of a general Chi.Bio setup is shown in Supplementary Fig. S1A.

Experiments were conducted in turbidostat mode with dither on, where the optical density (OD) of the culture was maintained at a setpoint of 0.5 by pumping in fresh media and removing spent media. A representative OD trace from a turbidostat experiment is shown in Supplementary Fig. S1B. The experimental setup used here is described in the Methods section of the main text, with fuller treatment provided in our published method on Chi.Bio characterisation with absolute quantification [2].

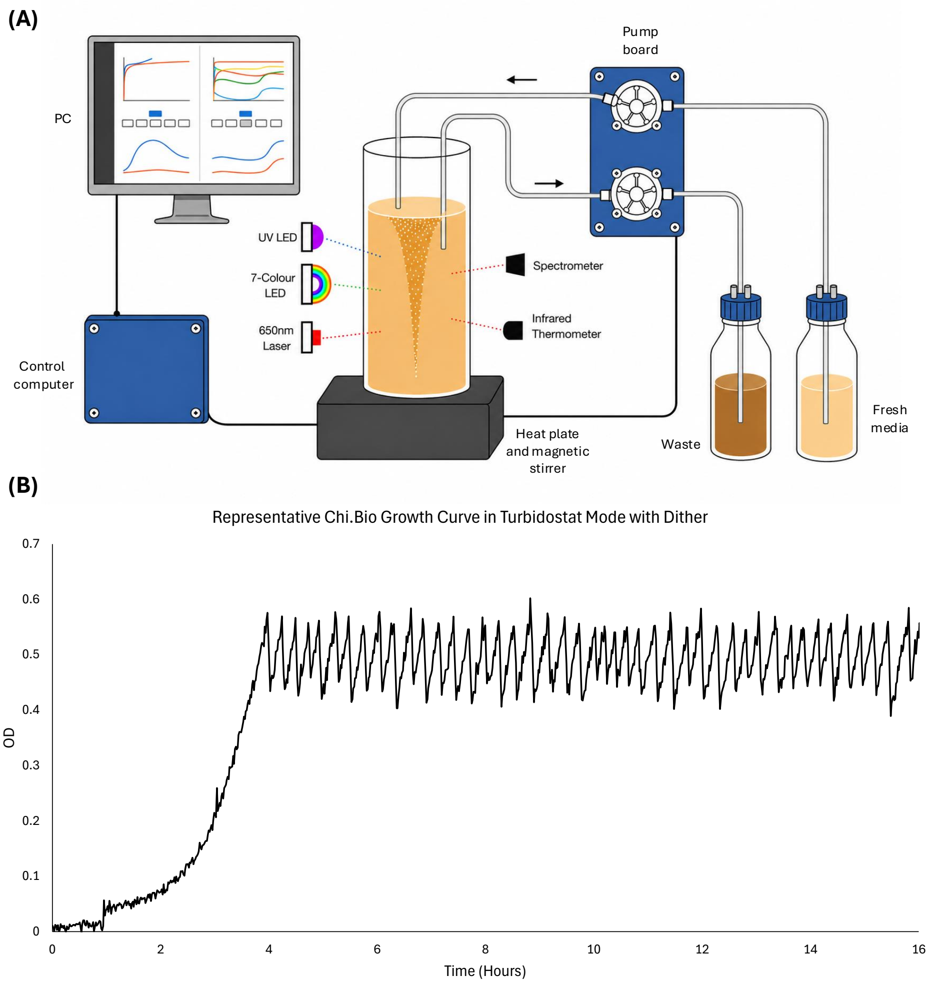


**Figure S1: Overview of Chi.Bio Experimental Setup: (A)** Diagram of Chi.Bio experimental setup. **(B)** Representative OD curve for a typical *E. coli* culture when grown in turbidostat mode with dither enabled with stirring = 0.5; temperature = 37°C; and OD setpoint = 0.5.

### S2 Part Validation

Before designing synthetic regulatory RNAs, we first conceptualised the target circuit and selected the fluorescent proteins and chemically inducible promoters required to build it. Three promoters from the Marionette collection [3] ($P_{VanCC}$, $P_{LuxB}$, and $P_{Tac}$) were chosen for initial testing in standardised GFPmut3 transcription units (Supplementary Fig. S2). Each transcription unit used the same ribozyme insulator (RiboJ), ribosome binding site (BBa_B0064), coding sequence (GFPmut3), terminator (L3S2P21), and plasmid backbone (pTU1-A-LacZ [4]), including the same origin of replication (pUC19-derived pMB1) and ampicillin-resistance gene. This design meant that differences in characterisation could be attributed primarily to the promoter, while also enabling model fitting with shared parameters across transcription units.

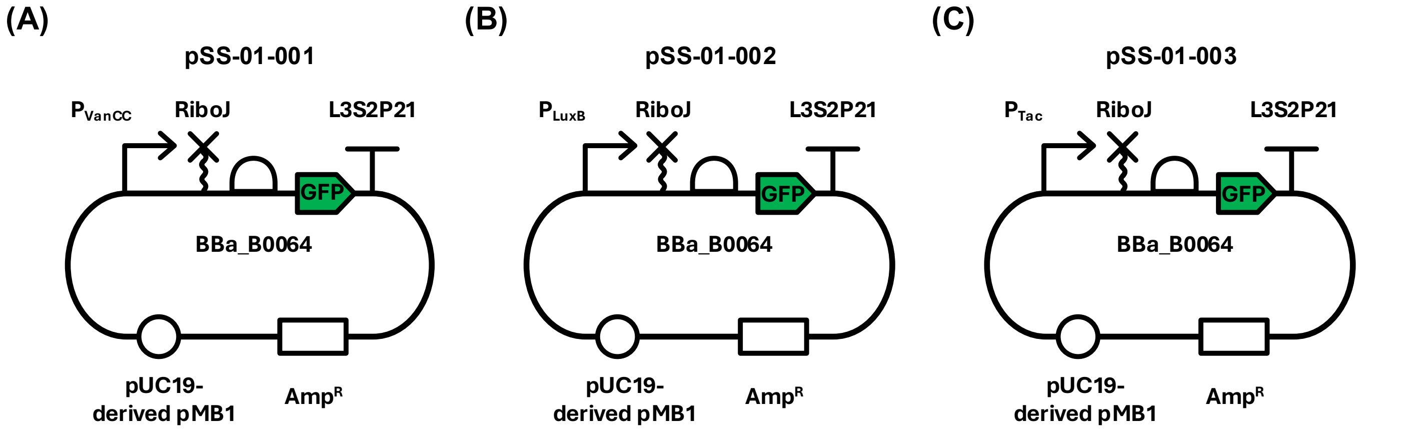


**Figure S2: Standardised transcription units used to test promoter function.** Each construct used the same RiboJ insulator, RBS (BBa_B0064), CDS (GFPmut3), terminator (L3S2P21), and plasmid backbone (pTU1-A-LacZ [4]), with expression driven by **(A)** $P_{VanCC}$ in pSS-01-001, **(B)** $P_{LuxB}$ in pSS-01-002, or **(C)** $P_{Tac}$ in pSS-01-003. Including the same terminator, RBS, and RiboJ insulator helped keep the resulting mRNAs as consistent as possible regardless of the promoter driving expression [5].

After assembling the standardised transcription units (Supplementary Fig. S2), we initially characterised them in microplate reader experiments (Supplementary Fig. S3). Dose-response curves were generated by varying the relevant inducer concentration for each promoter: vanillic acid for $P_{VanCC}$, OC6 for $P_{LuxB}$, and IPTG for $P_{Tac}$. Steady-state fluorescence was approximated by taking the mean normalised fluorescence from the early exponential phase window, excluding data with OD$< 0.2$ or OD$> 0.35$ as described in Methods. All three promoters showed a clear dose-dependent increase in fluorescence, high dynamic range, and typical sigmoidal response (Supplementary Fig. S3), and were therefore selected for further use.

Finally, after validating that the three standardised transcription units and their component parts were functioning as expected, we tested whether the two chosen fluorescent proteins (GFPmut3 and mScarlet-I) could be detected in the Chi.Bio under our chosen inducible promoters. In both cases the Chi.Bio measured clear changes in fluorescence upon induction (Supplementary Fig. S4), and therefore both fluorescent proteins were selected for further use.

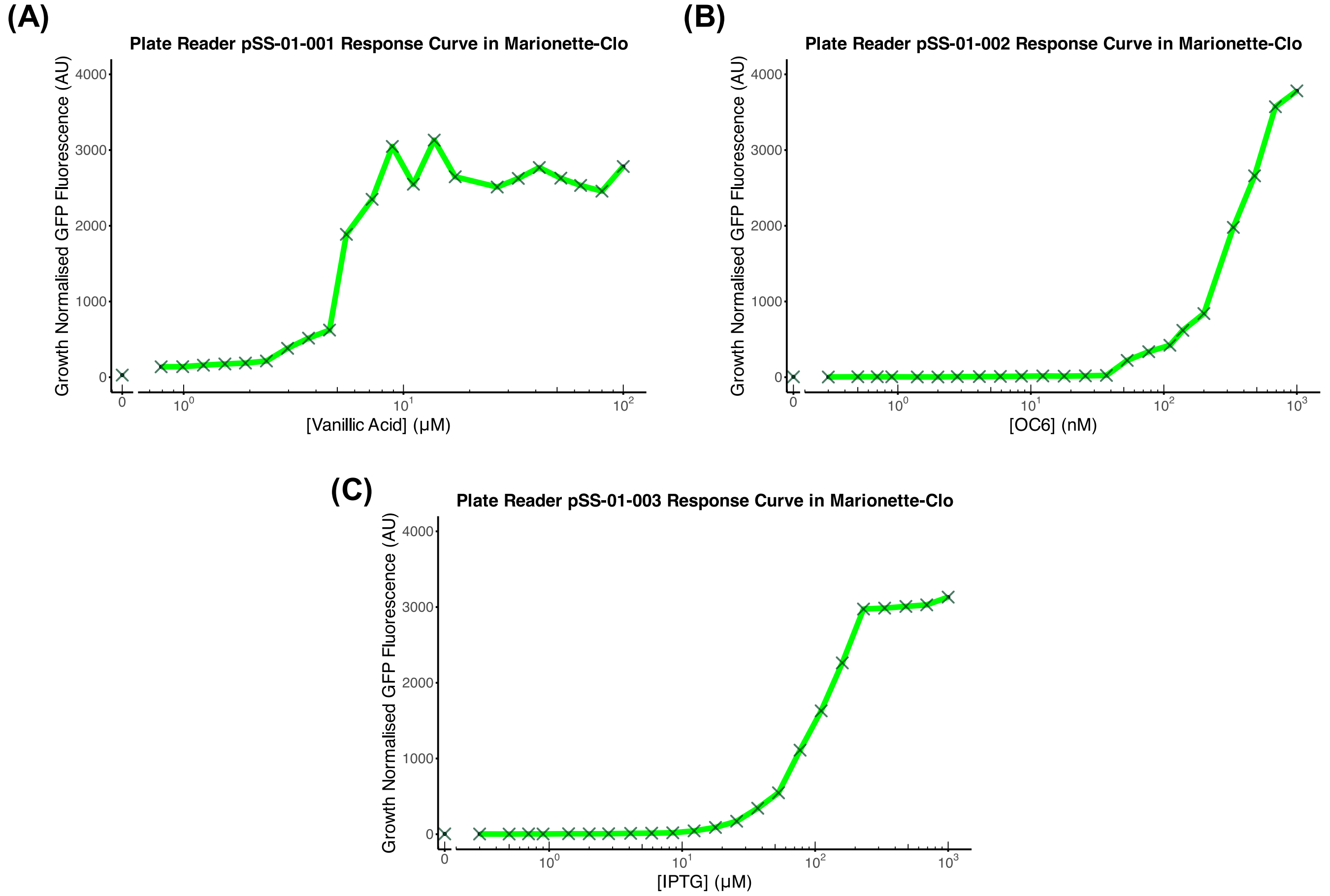


**Figure S3:** Standardised transcription units in plasmids pSS-01-001, pSS-01-002, and pSS-01-003 (Supplementary Fig. S2) were characterised in microplate reader experiments to generate response curves for **(A)** $P_{\mathrm{VanCC}}$, **(B)** $P_{\mathrm{LuxB}}$, and **(C)** $P_{\mathrm{Tac}}$. Fluorescence values were taken from early exponential phase and normalised to blanks, negative controls, and OD as described in Methods.

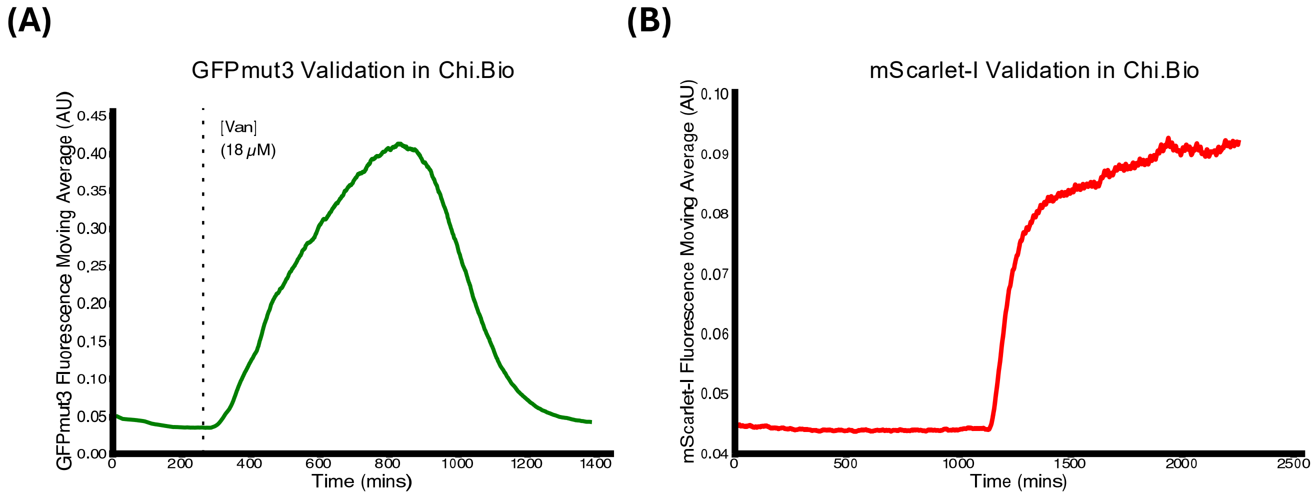


**Figure S4:** Chi.Bio [1] validation of the two fluorescent reporters, **(A)** GFPmut3 and **(B)** mScarlet-I. Each reporter produced a clear fluorescence increase after gene-expression induction.

### S3 Regulatory RNA Design

#### S3.1 ChiX-Derived Synthetic sRNA Design

To design synthetic versions of ChiX, a template sRNA sequence was adapted from the natural ChiX sRNA sequence (Supplementary Fig. S5):

acaccgtcgcttaaagtgacggcataataataaaaaaatgaaatTCCTCTTTGACGGgccaatagcgatattggccattttttt

**Figure S5: Natural sequence of ChiX:** Our determined consensus binding sequence is in blue and the invariable scaffold sequence is in black.

This template sequence contained a scaffold region and the consensus binding region of ChiX, which was defined using nucleotides known to bind one or both natural mRNA targets of ChiX, *dpiAB* [6] and *chiP* [7]. Natural target binding sites were predicted using IntaRNA [8] (Supplementary Fig. S6A–B). This consensus binding region (nucleotides 45 to 57 in ChiX) was considered to be the programmable region of the sRNA, responsible for the specificity of binding to mRNA targets. However, two nucleotides within this region are mismatched in ChiX binding to the sponge RNA ChbBC [9] (Supplementary Fig. S6C). These two nucleotides were therefore kept constant in the template in case they were important for spRNA function. All other nucleotides outside the programmable region were also kept constant, making up the invariable scaffold region of our synthetic ChiX-derived sRNAs. This process generated the ChiX-derived synthetic sRNA template that was used for subsequent design (Supplementary Fig. S7).

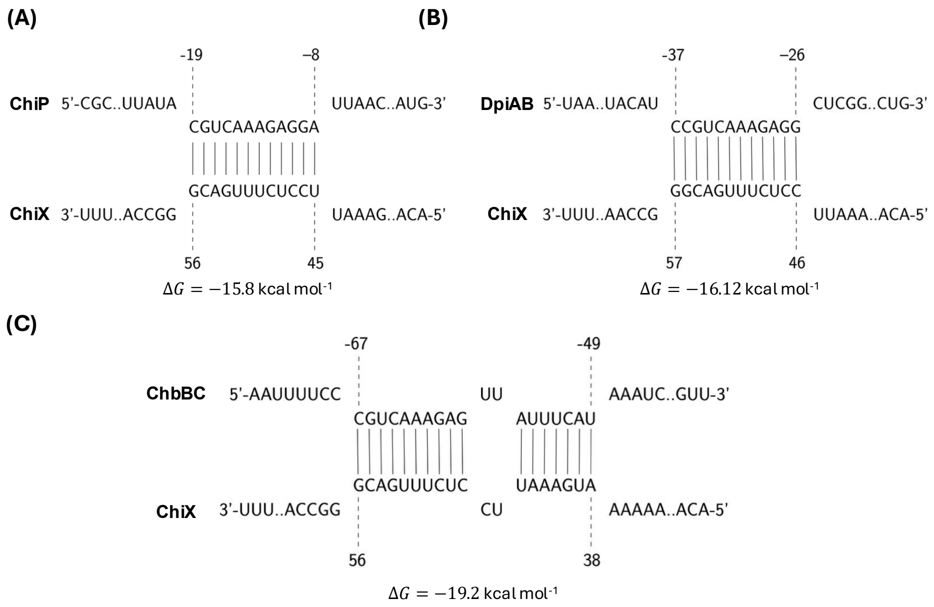


**Figure S6: IntaRNA [8] predicted binding of natural ChiX to its mRNA targets: (A)** ChiP and **(B)** DpiAB, as well as to its sponge RNA **(C)** ChbBC. Predicted binding interactions agree with reported binding for each interaction [9, 6, 10]. The IntaRNA-predicted free energy ($\Delta G$) is displayed.

acaccgtcgcttaaagtgacggcataataataaaaaaatgaaatTCN$_{(15\text{-}22)}$Ggccaatagcgatattggccattttttt

**Figure S7: Template sequence for the design of synthetic ChiX-derived sRNAs.** The invariable scaffold region is shown in black. Two nucleotides that are mismatched in ChiX–ChbBC binding (Supplementary Fig. S6C) are shown in red and maintained in the synthetic template. A single nucleotide that naturally pairs with DpiAB but not ChiP (Supplementary Fig. S6A–B) is shown in blue and was also kept constant. The programmable region is shown as a variable-length N region, with 15–22 variable nucleotides giving a total binding region of 18–25 nucleotides.

Next, this template sequence (Supplementary Fig. S7) was used to design synthetic ChiX-derived sRNAs. To identify potential target binding sites, the reverse complement of the complete binding region (i.e. the reverse complement of TCN$_{(15\text{-}22)}$G) was searched for in the region spanning 35 nucleotides upstream and downstream of the GFPmut3 RBS.

These potential target sites were investigated *in silico* using IntaRNA [8] for both on-target and off-target binding, and ten promising target sites were chosen. These included five sites upstream of the RBS and five downstream, with some overlapping and therefore directly binding the RBS. The resulting synthetic sRNAs were Syn-ChiX-18D, Syn-ChiX-18U, Syn-ChiX-20D, Syn-ChiX-20U, Syn-ChiX-22RD, Syn-ChiX-22U, Syn-ChiX-24RD, Syn-ChiX-24RU, Syn-ChiX-24U, and Syn-ChiX-25D. Predicted free energies for target and off-target binding are given in Supplementary Table S1. Structures for the ten synthetic sRNAs were predicted using Vienna RNA's RNAFold tool [11], visualised with RNACanvas [12], and compared with a structure prediction of natural ChiX (Supplementary Fig. S8), showing that these sRNAs generally maintained similar structural motifs to natural ChiX. Sequences of the ten designed synthetic ChiX-derived sRNAs are given in Supplementary File 1.

| **sRNA** | $\Delta G_{\mathrm{GFP}}$ (kcal mol$^{-1}$) | $\Delta G_{\mathrm{mScarlet}}$ (kcal mol$^{-1}$) | **Endogenous off-target** | $\Delta G$ (kcal mol$^{-1}$) |
|---|---|---|---|---|
| Syn-ChiX-18D | -25.76 | -7.85 | *aptD* | -16.13 |
| Syn-ChiX-18U | -19.97 | -10.48 | *yciV* | -17.22 |
| Syn-ChiX-20D | -27.89 | -8.41 | *moaE* | -17.08 |
| Syn-ChiX-20U | -22.49 | -19.90 | *yciV* | -17.37 |
| Syn-ChiX-22RD | -30.40 | -6.19 | *aptD* | -16.10 |
| Syn-ChiX-22U | -27.63 | -20.44 | *yciV* | -17.21 |
| Syn-ChiX-24RD | -35.52 | -7.77 | *msrQ* | -17.47 |
| Syn-ChiX-24RU | -18.86 | -6.07 | *yedD* | -16.16 |
| Syn-ChiX-24U | -28.09 | -22.07 | *rcsD* | -18.30 |
| Syn-ChiX-25D | -33.37 | -5.54 | *dgcP* | -20.50 |
| Syn-ChiX-20NC | -6.93 | -7.98 | *yeiL* | -16.97 |

**Table S1: Predicted binding free energies for ChiX-derived synthetic sRNAs against intended and off-target transcripts.** Predicted binding free energies ($\Delta G$) are shown for each ChiX-derived synthetic sRNA against the intended GFPmut3 target, the synthetic mScarlet-I off-target, and the most favourable endogenous off-target predicted by IntaRNA [8]. Small RNA sequences input into IntaRNA included RiboJ10, the synthetic sRNA sequence, and the L3S2P56 terminator. Endogenous off-target predictions were carried out against the *E. coli* DH10B reference genome (NC_010473). Predictions against GFPmut3 and mScarlet-I used putative mRNA sequences including the ribozyme insulator, RBS, coding sequence, and terminator. Sequences used for all IntaRNA predictions are provided in Supplementary File 1. All predictions were carried out using default settings with the IntaRNA online tool [8, 13].

##### S3.1.1 Assembly of ChiX-Derived Synthetic sRNA Circuits

These synthetic sRNAs were synthesised for assembly into a simple two-input synthetic gene circuit. The first transcription unit was the $\mathrm{P_{VanCC}}$–GFPmut3 unit in pSS-01-001. A second transcription unit was assembled into EcoFlex pTU1-B-LacZ [4], comprising $\mathrm{P_{LuxB}}$, a RiboJ10 insulator, one of the synthetic sRNAs, and the L3S2P56 terminator, generating plasmids pSS-01-sRNA-01 through pSS-01-sRNA-10. Ten two-input synthetic gene circuits of the form shown in Fig. 2C were then assembled by combining pSS-01-001 with one of the pSS-01-sRNA plasmids into EcoFlex pTU2-a-RFP [4].

The ten synthetic sRNAs generated by this design process were Syn-ChiX-18D, Syn-ChiX-18U, Syn-ChiX-20D, Syn-ChiX-20U, Syn-ChiX-22RD, Syn-ChiX-22U, Syn-ChiX-24RD, Syn-ChiX-24RU, Syn-ChiX-24U, and Syn-ChiX-25D, targeting sequences downstream (D), upstream (U), or overlapping the RBS (R).

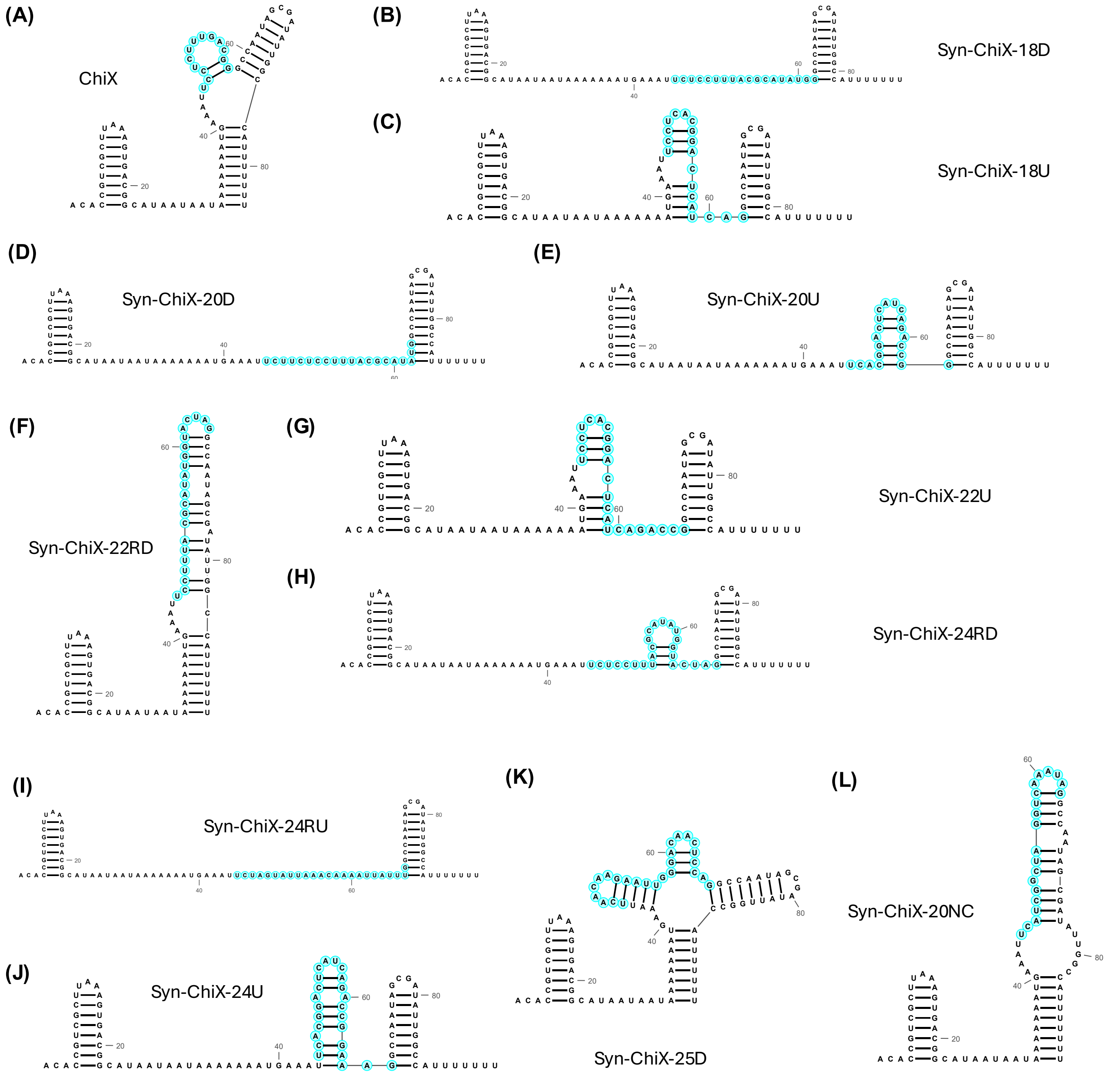


**Figure S8:** Vienna RNAFold [11] structure predictions visualised by RNACanvas [12] for natural ChiX, the designed synthetic ChiX variants, and the non-binding negative control ChiX (Syn-ChiX-20NC). RNA structure predictions were carried out using default RNAFold parameters using the online web server (http://rna.tbi.univie.ac.at/cgi-bin/RNAWebSuite/RNAfold.cgi). For each RNA, the RNA sequence and minimum free energy (MFE) structure in Vienna format was loaded into RNACanvas [12], where putative binding regions were outlined in blue. Sequences used for structure predictions were the sRNA sequences alone (i.e. not including RiboJs, terminators, etc.).

#### S3.2 ChbBC-Derived Sponge RNA Design

Two natural sponge RNAs have been described for ChiX: the 5' mRNA fragment ChiZ [14] and the intergenic region ChbBC [15, 9, 10, 16]. To design a synthetic ChbBC-derived spRNA for Syn-ChiX-20D, the full intergenic region between ChbB and ChbC was analysed for binding to ChiX using IntaRNA [8]. This identified the likely sRNA–spRNA binding region (Supplementary Fig. S10A), with the predicted interaction matching reported binding interactions [15, 16]. Part of the ChbBC–ChiX interaction occurs in the ChiX–mRNA binding region, which was reprogrammed to generate Syn-ChiX-20D, while part occurs in the invariant scaffold of Syn-ChiX-20D. Therefore, a template sequence for synthetic ChbBC was generated with an extended binding region to bind the extended 20 nucleotide binding region of Syn-ChiX-20D, while nucleotides involved in binding invariant parts of Syn-ChiX-20D were kept constant (Supplementary Fig. S9). This template was then used to generate Syn-ChbBC-20D with a binding region antisense to the Syn-ChiX-20D binding region. The sequence for Syn-ChbBC-20D is provided in Supplementary File 1.

TTTATTTTAAATTTTCC NNNNNNNNNNNNNNNNNNNN TT ATTTCAT AAATCAATACCGCAATATT TAAATTGCGGTTTTTAAGGGTATTTTTCT

**Figure S9: Synthetic ChbBC-based sponge RNA design template**

**(A)**

```
                         -67                      -49
                          |                        |
ChbBC      5'-AAUUUUCC              UU        AAAUC..GUU-3'
                       CGUCAAAGAG  AUUUCAU
                       ||||||||||  |||||||          ΔG = −19.2 kcal mol^-1
                       GCAGUUUCUC  UAAAGUA
ChiX     3'-UUU..ACCGG              CU        AAAAA..ACA-5'
                          |                        |
                         56                        38
```

**(B)**

```
                                    ΔG = −25.73 kcal mol^-1
                          86                                   112
                           |                                    |
Syn-ChbBC-20D-GFP  5'-ACC..UUUCC                      UU        AAAUC..AAA-3'
                                CAUAUGCGUAAAGGAGAA  AUUUCAU
                                ||||||||||||||||||  |||||||
                                GUAUACGCAUUUCCUCUU  UAAAGUA
Syn-ChiX-20D-GFP   3'-CCU..AACCG                      CU        AAAAA..CAA-5'
                           |                                    |
                          132                                  106
```

**Figure S10: IntaRNA [8] predicted binding between ChiX sRNA and ChbBC spRNA: (A)** natural ChiX and ChbBC; **(B)** synthetic sRNA–spRNA pair, Syn-ChiX-20D and Syn-ChbBC-20D.

IntaRNA [8] predicted the desired interaction between Syn-ChiX-20D and Syn-ChbBC-20D (Supplementary Fig. S10B), with $\Delta G = -25.73$ kcal mol$^{-1}$. The most favourable predicted endogenous off-target was *yrbG*, with $\Delta G = -20.26$ kcal mol$^{-1}$. Structures were predicted with Vienna RNAFold [11] and visualised with RNACanvas [12]. Although Syn-ChbBC-20D was predicted to differ structurally from natural ChbBC, several features were maintained, including a likely important stem loop and substantial single-stranded portions of the sRNA binding site (Supplementary Fig. S11).

In addition to the above-described design, the same process was applied to design an equivalent circuit regulating mScarlet-I rather than GFPmut3, to demonstrate the generalisability of our regulatory parts. In this design, the Syn-ChiX variant binds mScarlet-I mRNA while the Syn-ChbBC variant binds the Syn-ChiX variant. Supplementary Table S2 shows the IntaRNA predictions [8], again showing favourable binding to the desired targets and less favourable off-target binding.

**Table S2: Predicted binding free energies for mScarlet-I-regulating RNA circuit.**

| RNA | Type | Target | $\Delta G$ | Endogenous off-target | $\Delta G$ |
|---|---|---|---|---|---|
| Syn-ChiX-20D-mSc | sRNA | mScarlet-I mRNA | -30.15 | *moaA* | -17.45 |
| Syn-ChbBC-20D-mSc | spRNA | Syn-ChiX-20D-mSc | -33.44 | *yejG* | -19.36 |

#### S3.3 MicC-Derived sRNA Design

Syn-MicC-GFP was generated by following previously described design protocols [17, 18] for our GFPmut3 mRNA. Syn-MicC-GFP had the desired predicted binding, with a favourable $\Delta G = -32.29$ kcal mol$^{-1}$ (Supplementary Fig. S12).

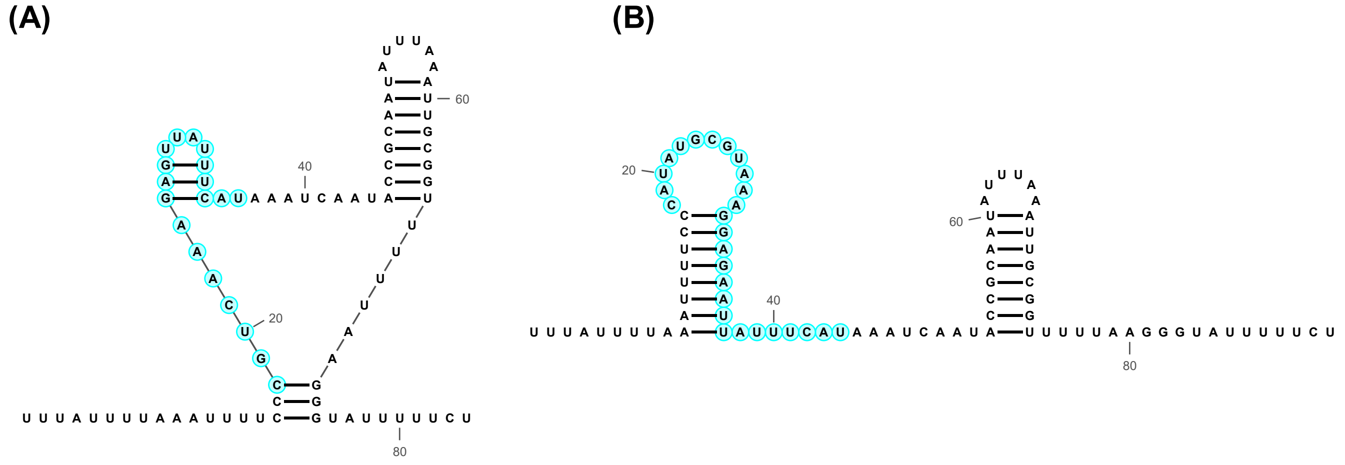


**Figure S11: Vienna RNAFold [11] predicted structures visualised with RNACanvas [12]: (A)** natural ChbBC and **(B)** Syn-ChbBC-20D. A stem loop is maintained and, in both structures, significant portions of the sRNA binding region (labelled in blue) are single stranded.

The sequence for Syn-MicC-GFP is provided in Supplementary File 1.

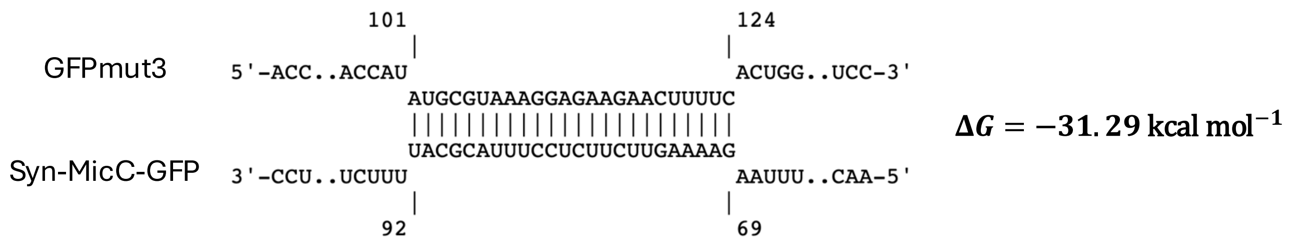


**Figure S12: IntaRNA [8] predicted binding between Syn-MicC-GFP and the GFPmut3 mRNA sequence.**

#### S3.4 Alternative Sponge RNA Design Strategies

After design of Syn-MicC-GFP, synthetic spRNAs were designed by taking the antisense of the whole sequence to generate Syn-MicC-spRNA-EA (entirely antisense), maintaining the scaffold and taking the antisense of the binding region to generate Syn-MicC-spRNA-AB (antisense binding site), or changing the binding region of ChiX to the 24 antisense nucleotides of the Syn-MicC-GFP binding region to generate Syn-ChiX-spRNA. All three designs had the desired IntaRNA-predicted binding and favourable $\Delta G$ values (Supplementary Fig. S13). Off-target binding was less favourable for all of these alternative sponge RNAs: $-20.09$ kcal mol$^{-1}$, $-20.98$ kcal mol$^{-1}$, and $-20.46$ kcal mol$^{-1}$ for Syn-MicC-spRNA-EA binding *ydbL*, Syn-MicC-spRNA-AB binding *aqpZ*, and Syn-ChiX-spRNA binding *yrbG*, respectively. The sequences for these spRNAs are provided in Supplementary File 1.

#### S3.5 Assembly of Synthetic Sponge RNA Circuits

After design and synthesis of the synthetic spRNA variants, several synthetic gene circuits of the form shown in Fig. 3C were assembled for characterisation. Each circuit contained three transcription units: (i) a transcription unit expressing GFPmut3 in response to vanillic acid ($P_{VanCC}$, RiboJ, BBa_B0064 RBS, GFPmut3 CDS, and L3S2P21); (ii) a transcription unit expressing one of the synthetic sRNAs (Syn-ChiX-20D or Syn-MicC-GFP) in response to OC6 ($P_{LuxB}$, RiboJ10, the sRNA, and L3S2P56); and (iii) a transcription unit expressing one of the synthetic spRNAs (Syn-ChbBC-20D, Syn-MicC-spRNA-EA (entirely antisense), Syn-MicC-spRNA-AB (antisense binding), or Syn-ChiX-spRNA) in response to IPTG ($P_{Tac}$, RiboJ51, the spRNA, and L3S1P52). Optionally, pET-RBS and mScarlet-I were placed downstream of the spRNA and upstream of the terminator (L3S1P52) to enable live measurement of spRNA expression. Complete details of each single-target sponge RNA circuit described in this section are provided in Supplementary File 2.

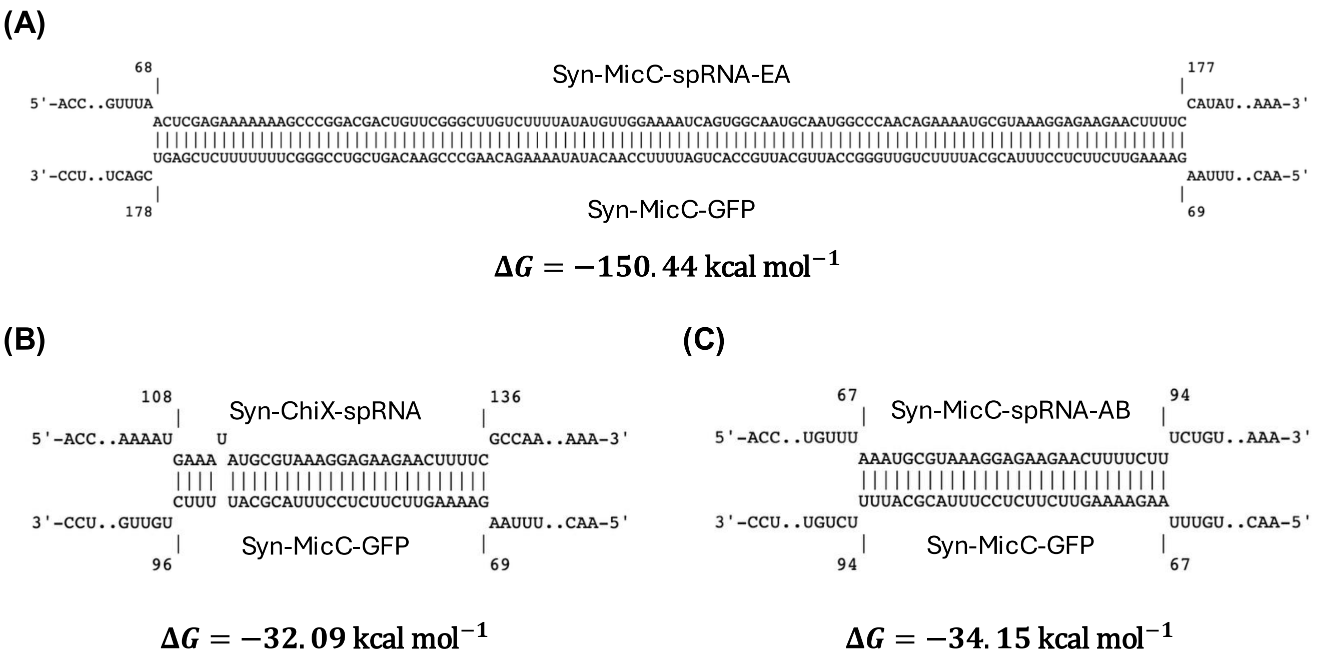


**Figure S13: IntaRNA [8] predicted binding for alternative design sponge RNAs: (A)** Syn-MicC-spRNA-EA (entirely antisense) binding to Syn-MicC-GFP; **(B)** Syn-ChiX-spRNA binding to Syn-MicC-GFP; and **(C)** Syn-MicC-spRNA-AB (antisense binding site) binding to Syn-MicC-GFP.

#### S3.6 Design and Assembly of Multi-Target RNA Circuits

Following methods developed for fusion sRNAs [19], we designed multi-target versions of ChiX- and MicC-derived sRNAs containing two binding regions targeting GFPmut3 and mScarlet-I. These designs were based on the templates in Supplementary Figures S14 and S15. IntaRNA predictions for these synthetic multi-target sRNAs showed strong predicted binding to both desired mRNA targets, with lower predicted binding to off-targets (Supplementary Table S3). We assembled these designs into multi-target sRNA circuits (Supplementary Fig. S16).

ACACCGTCGCTTAAAGTGACGGCATAATAATAAAAAAATGAAAT NNNNNNNNNNNNNNNNNNNNN AAA NNNNNNNNNNNNNNNNNNNNN GCCAATAGCGATATTGGCCATTTTTTT

**Figure S14: Two-binding-region synthetic ChiX-derived sRNA template**

NNNNNNNNNNNNNNNNNNNNNNNNNN AAA NNNNNNNNNNNNNNNNNNNNNNNNNN TTTCTGTTGGGC CATTGCATTGCCACTGATTTTCCAACATATAAAAAGACAAGCCCGAACAGTCGTCCGGGCTTTTT TTCTCGAG

**Figure S15: Two-binding-region synthetic MicC-derived sRNA template**

**Table S3: Predicted binding free energies for sRNAs design for multi-target gene circuits.** Predicted binding free energies ($\Delta G$) are shown for each synthetic sRNA against GFPmut3, mScarlet-I, and the most favourable endogenous off-target predicted by IntaRNA.

| **sRNA** | $\Delta G_{\text{GFP}}$ (kcal mol$^{-1}$) | $\Delta G_{\text{mScarlet}}$ (kcal mol$^{-1}$) | **Endogenous off-target** | $\Delta G$ (kcal mol$^{-1}$) |
|---|---|---|---|---|
| Syn-MicC-Multi | -31.75 | -49.66 | *zapC* | -23.40 |
| Syn-ChiX-Multi | -23.09 | -35.28 | *ypeC* | -15.82 |
| Syn-MicC-mSc | – | -42.36 | *zapC* | -20.72 |

These designs suggested two strategies for generating synthetic sponge RNAs capable of regulating multiple genes simultaneously. For strategy (i), we designed a circuit comprising constitutive expression of mScarlet-I and GFPmut3,

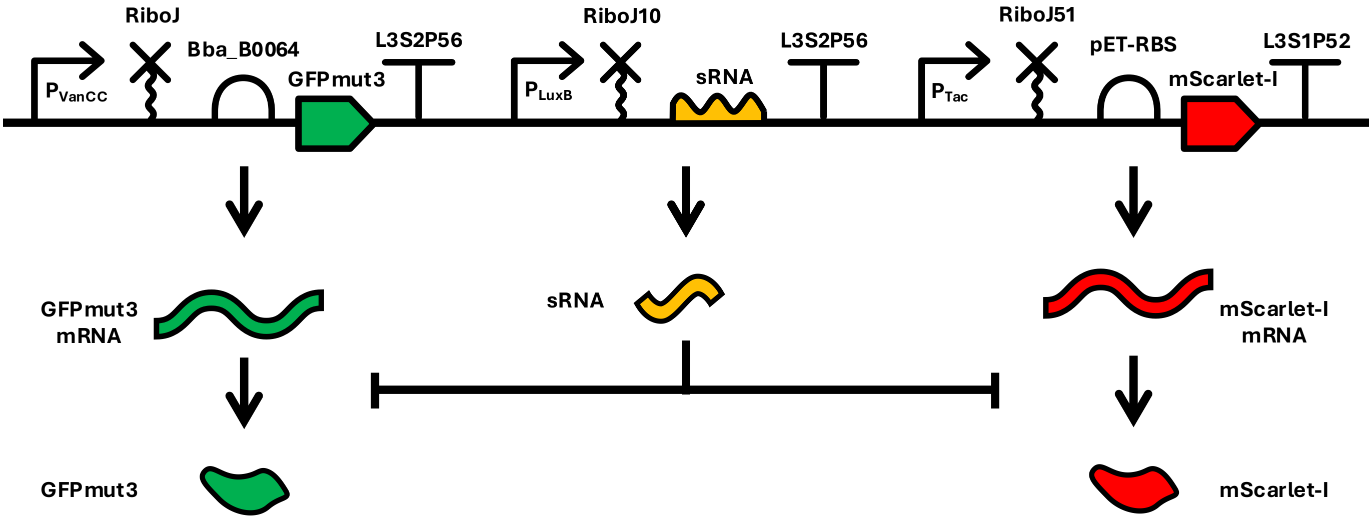


**Figure S16: Circuit architecture for multi-target sRNA circuits.**

two MicC-derived sRNAs targeting GFPmut3 or mScarlet-I respectively, and a MicC-based sponge RNA containing binding regions for both sRNAs (using the same design template as Supplementary Fig. S15). IntaRNA predictions again showed strong predicted binding between each synthetic sRNA and the multi-target sponge RNA (Supplementary Table S4). We then assembled these parts into a multi-target spRNA circuit (Fig. 6A; pSS-02-075 + pSS-01-023).

**Table S4: Predicted binding free energies for spRNAs used in multi-target spRNA gene circuits.** Predicted binding free energies ($\Delta G$) are shown for the synthetic sponge RNA against its cognate synthetic sRNAs and the most favourable endogenous off-target predicted by IntaRNA.

| spRNA | sRNA 1 | $\Delta G$ | sRNA 2 | $\Delta G$ | Endogenous off-target | $\Delta G$ |
|---|---|---|---|---|---|---|
| Syn-MicC-Multi-spRNA | Syn-MicC-GFP | -33.45 | Syn-MicC-mSc | -35.32 | *yebU* | -20.72 |
| Syn-MicC-spRNA-AB | Syn-MicC-Multi | -32.54 | – | – | *aqpZ* | -20.98 |

For strategy (ii), we designed a circuit comprising constitutive expression of mScarlet-I, inducible expression of GFPmut3, a multi-target MicC-derived sRNA targeting both genes (Syn-MicC-Multi; see above), and a sponge RNA that binds at least one of the two binding sites of Syn-MicC-Multi (Syn-MicC-spRNA-AB; see above). IntaRNA predictions again showed strong predicted binding between Syn-MicC-spRNA-AB and Syn-MicC-Multi (Supplementary Table S4). A circuit of the form shown in Fig. 6B was then assembled (pSS-02-087).

### S4 Regulatory RNA Characterisation

#### S4.1 ChiX-Derived Synthetic sRNAs

Initial validation of synthetic sRNA functionality was performed in microplate reader assays with varying OC6 concentrations (sRNA induction) and fixed GFPmut3 expression for each Syn-ChiX variant. In general, sRNAs targeting at or downstream of the RBS were functional (Supplementary Fig. S17). After this initial validation, the sRNAs that appeared to function well in microplate reader assays were further characterised in Chi.Bio experiments, again showing functional downregulation of GFPmut3 expression during exponential growth (Fig. 2E; Supplementary Fig. S18).

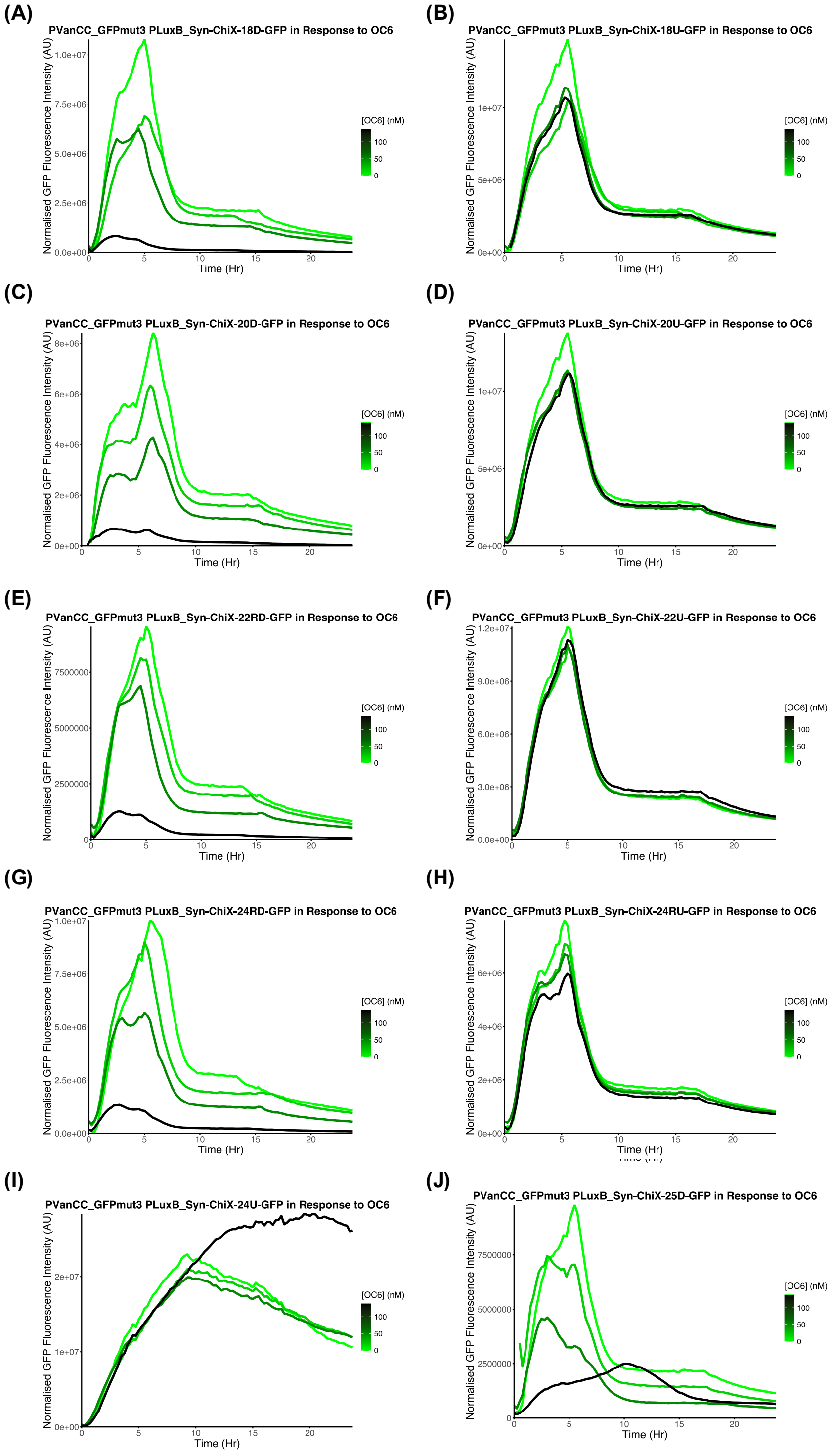


**Figure S17:** Characterisation of synthetic ChiX-derived sRNAs in microplate reader assays at varying OC6 concentrations (sRNA induction) and fixed GFPmut3 expression.

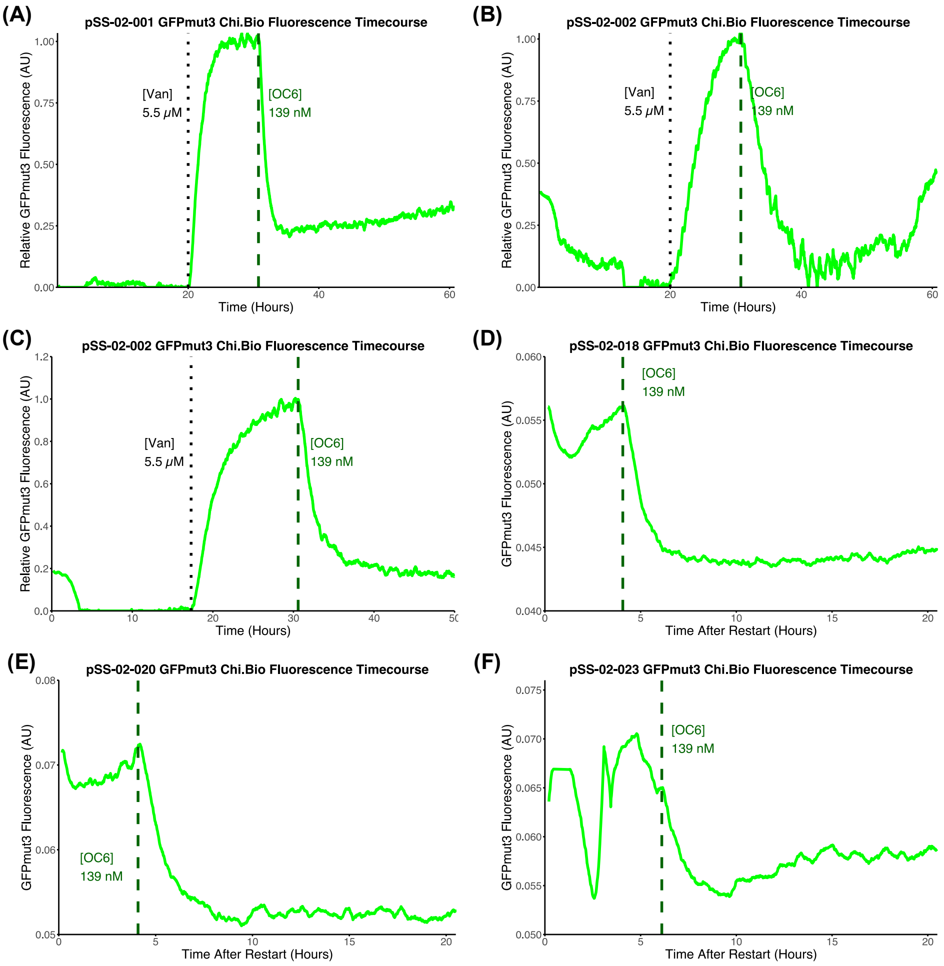


**Figure S18:** Chi.Bio characterisation of synthetic ChiX-derived sRNA circuits by sequential induction of GFPmut3 (5.5 µM vanillic acid) followed by sRNA (139 nM OC6) for **(A)** a second replicate for Syn-ChiX-20D (pSS-02-001), **(B–C)** two replicates for Syn-ChiX-18D (pSS-02-002), **(D)** Syn-ChiX-22RD (pSS-02-018), **(E)** Syn-ChiX-24RD (pSS-02-020), and **(F)** Syn-ChiX-25D (pSS-02-023). In **(D–F)**, data are shown after a restart following Chi.Bio malfunction, by which point GFPmut3 was already induced.

#### S4.2 ChbBC-derived synthetic sponge RNAs

Initial characterisation of ChbBC-derived synthetic sponge RNAs is described in the Results section of the main text. In these Chi.Bio experiments, GFPmut3 was first induced with 5.5 µM vanillic acid, followed by induction of the sRNA with 139 nM OC6 and induction of the sponge RNA with 1000 µM IPTG (Fig. 3D–E). A second Chi.Bio experiment under the same conditions showed very similar qualitative behaviour (Supplementary Fig. S19), with the spRNA in pSS-02-005 (ChbBC spRNA circuit without mScarlet-I) leading to lower recovery of GFPmut3 expression than in pSS-02-006 (ChbBC spRNA circuit with mScarlet-I). This validated synthetic sponge RNA function across multiple experiments.

As described in the main text, we next tested whether higher levels of GFPmut3 recovery could be achieved by sponge RNA expression when sRNA induction was lower, because the partial GFPmut3 recovery observed in previous characterisation experiments could be due to sRNA being in excess. We therefore carried out a similar characterisation experiment but induced the sRNA with 50 nM OC6 rather than 139 nM OC6 (Supplementary Fig. S20). GFPmut3 recovery was higher for both circuits in these experiments (Supplementary Fig. S20A–B) compared to previous characterisations with 139 nM OC6 (Fig. 3C–D; Supplementary Fig. S19). In pSS-02-006, GFPmut3 reached higher expression levels after spRNA induction than the steady-state GFPmut3 expression prior to sRNA induction (i.e. normalised fluorescence > 1), which may be due to leaky sRNA expression or growth-rate changes (e.g. due to gene expression burden from expressing both mScarlet-I and GFPmut3). To examine this further, we normalised GFPmut3 fluorescence trajectories to dynamically estimated Chi.Bio growth rates. This reduced the normalised fluorescence value below the initial steady-state value, suggesting that the higher GFPmut3 fluorescence is explained by reductions in growth rate.

The circuit with mScarlet-I (pSS-02-006) was further characterised in quadruplicate with additional changes in inducer

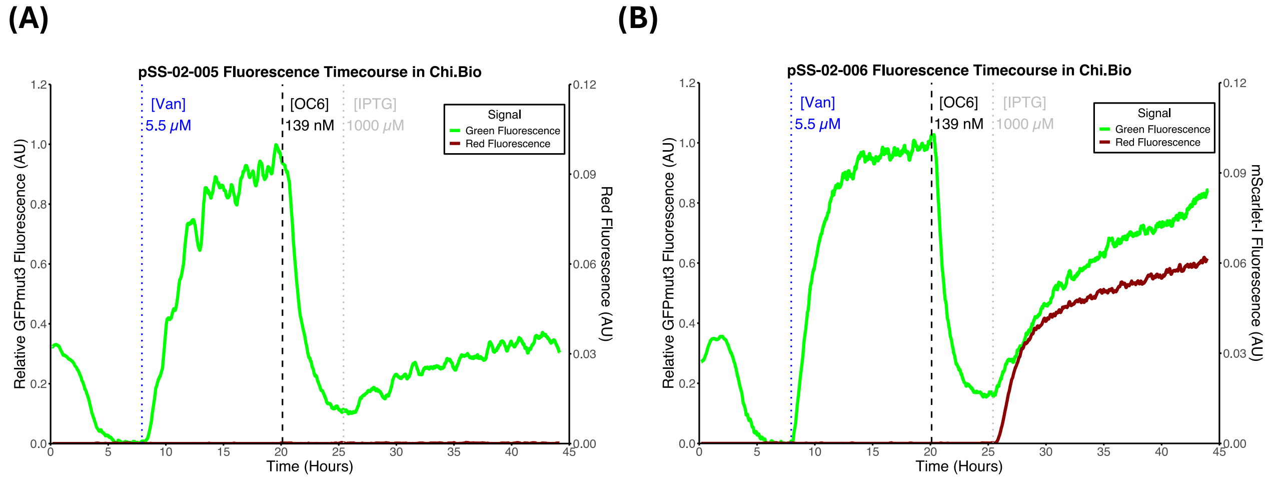


**Figure S19: Repeat Chi.Bio characterisation of synthetic ChbBC-derived sponge RNA circuits:** characterisation was carried out by sequential induction of GFPmut3, followed by sRNA, followed by spRNA, through step changes in inducer from 0 to 5.5 μM, 139 nM, and 1000 μM for vanillic acid, OC6, and IPTG, respectively, with induction timepoints shown by vertical lines. GFPmut3 fluorescence is normalised so that steady-state fluorescence after initial induction is set to 1. Characterisations are shown for **(A)** the ChbBC sponge RNA circuit without mScarlet-I (pSS-02-005) and **(B)** the ChbBC sponge RNA circuit with mScarlet-I (pSS-02-006).

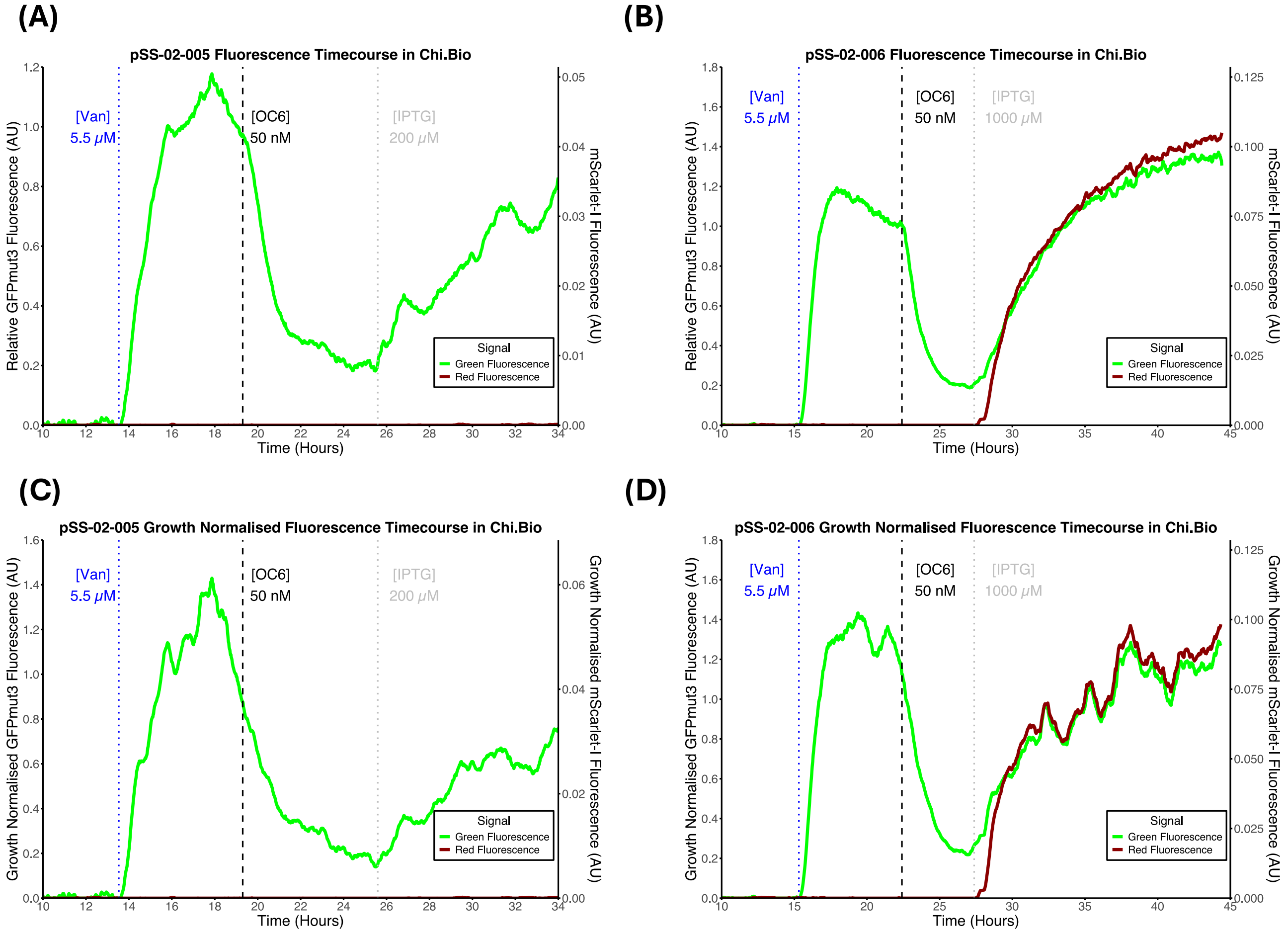


**Figure S20: ChbBC-derived synthetic sponge RNA circuit characterisation with lower sRNA induction:** pSS-02-005 and pSS-02-006 were tested with lower initial induction of Syn-ChiX-20D (50 nM versus 139 nM). For both circuits, similar levels of sRNA-mediated GFPmut3 knockdown were observed compared to previous characterisations, but both pSS-02-005 **(A)** and pSS-02-006 **(B)** showed higher spRNA-mediated recovery of GFPmut3. Growth normalisation of GFPmut3 trajectories **(C–D)** reduced the apparent spRNA-mediated recovery, but recovery remained higher than in previous experiments where OC6 = 139 nM.

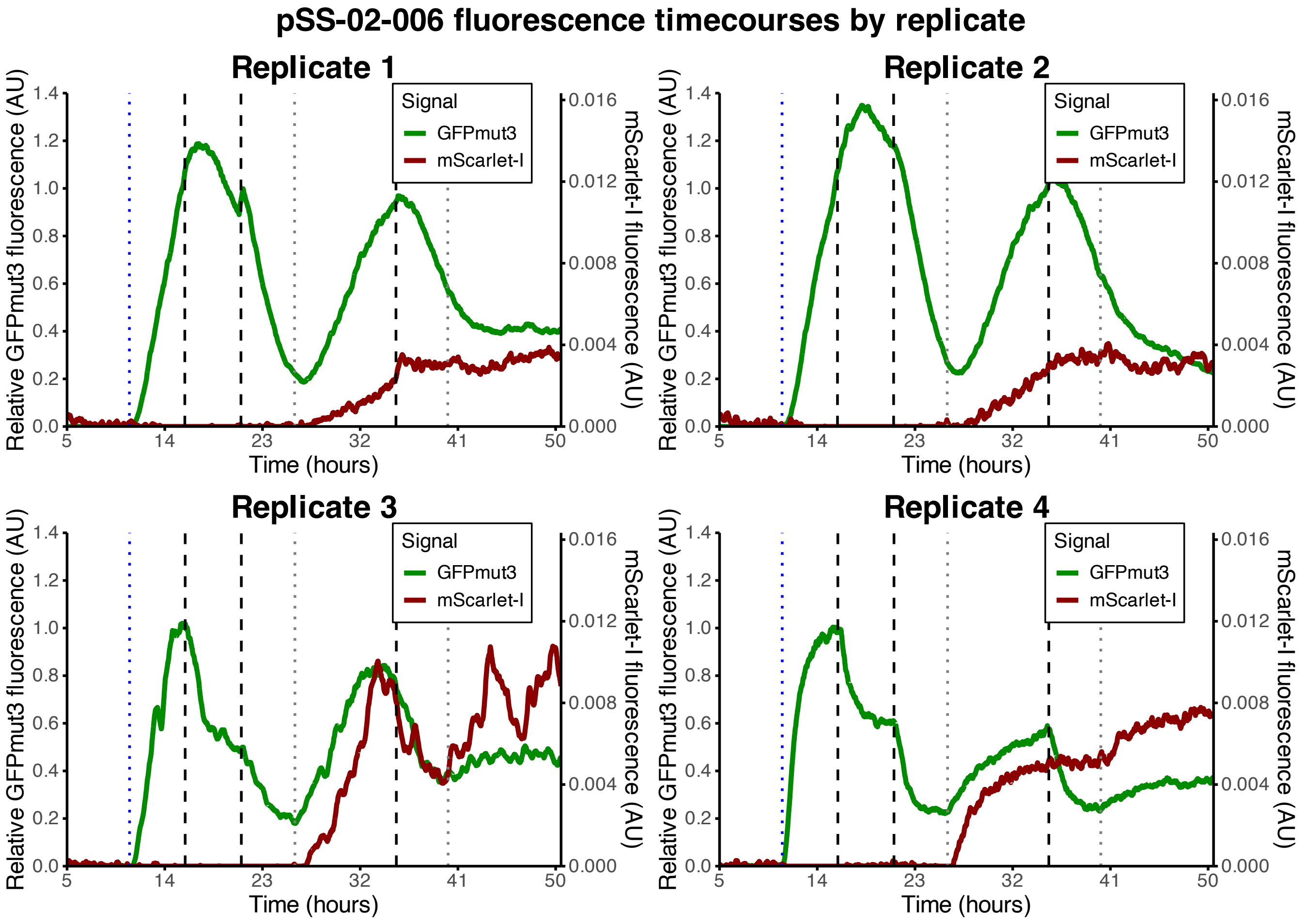


**Figure S21: Quadruplicate characterisation of pSS-02-006 in response to sequential induction step changes.** All inducer concentration step changes occur at the vertical dotted/dashed lines and are in order from 0 to 5.5 µM vanillic acid, 0 to 12 nM OC6, 12 to 50 nM OC6, 0 to 200 µM IPTG, 50 to 300 nM OC6, and 200 to 1000 µM IPTG.

conditions (Supplementary Fig. S21). Although there were quantitative differences across replicates, the circuit behaved similarly across the four replicates qualitatively. In particular, the timescales on which the circuit operated were consistent, given that these circuits were characterised over the course of 50 hours.

#### S4.3 Alternative sponge RNA design strategies characterisation

As described in the main text, here we present additional data for Chi.Bio characterisation of the alternative design strategy sponge RNA circuits (Supplementary Fig. S22). These data show growth-normalised fluorescence curves (fluorescence multiplied by growth rate) for the alternative sponge RNA circuits, showing that some circuits did not function when mScarlet-I was co-transcribed with the spRNA (Supplementary Fig. S22D,F).

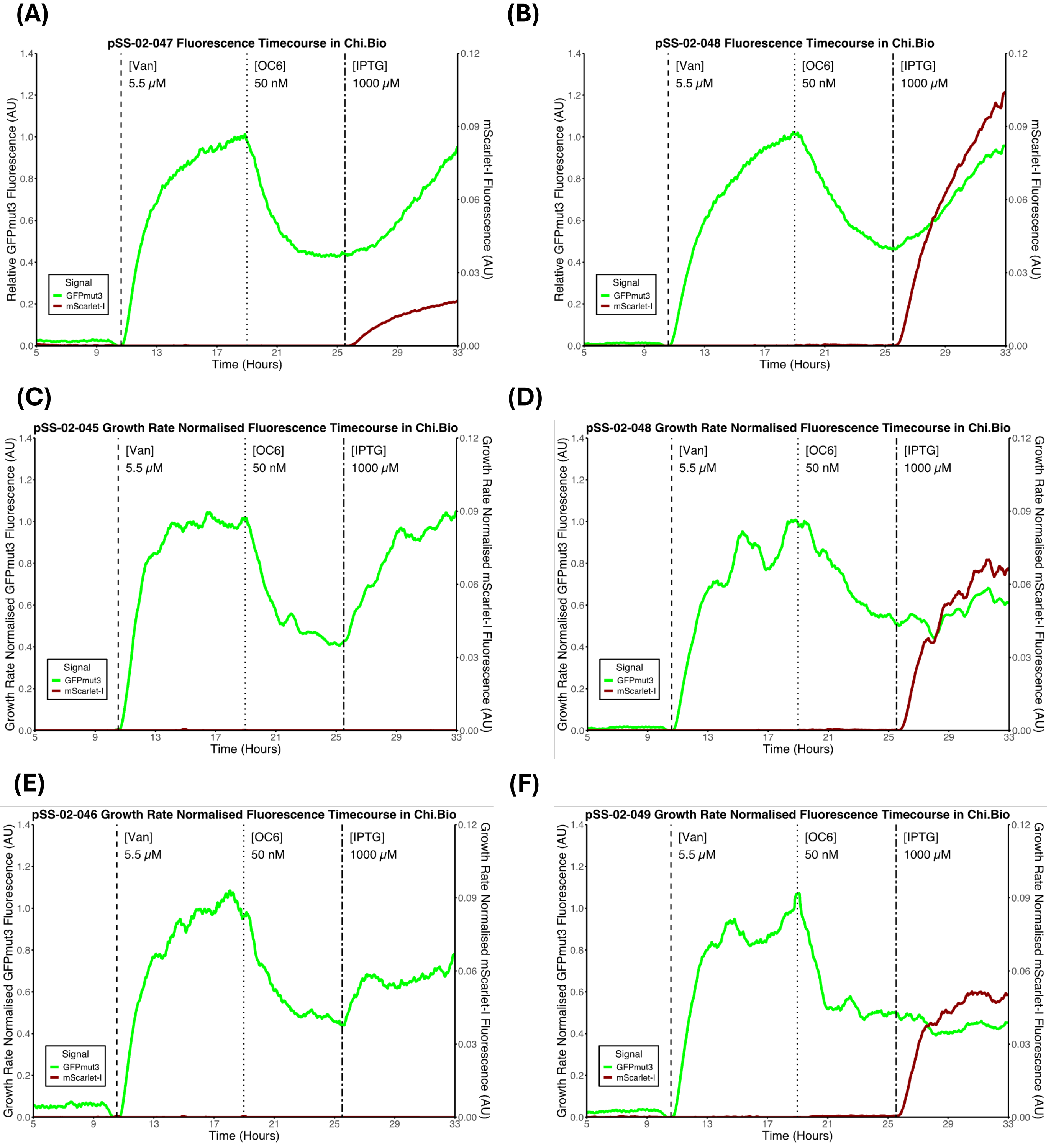


**Figure S22: Additional data for characterisation of alternative design sponge RNAs: (A)** characterisation of pSS-02-047, a sponge circuit with Syn-MicC-spRNA-EA as the spRNA and mScarlet-I co-transcribed with it, showing a functional spRNA circuit. **(B)** Characterisation of pSS-02-048, a spRNA circuit with Syn-MicC-spRNA-AB as the spRNA and mScarlet-I co-transcribed with it, showing an apparent functional spRNA circuit. **(C–D)** Growth-normalised fluorescence curves for pSS-02-045 (Syn-MicC-spRNA-AB) and pSS-02-048 (Syn-MicC-spRNA-AB with mScarlet-I) show the spRNA only functions in the absence of co-transcribed mScarlet-I. **(E–F)** Characterisation of pSS-02-046 (Syn-ChiX-spRNA) and pSS-02-049 (Syn-ChiX-spRNA with mScarlet-I) shows a functional spRNA circuit only in the absence of the co-transcribed mScarlet-I.

### S5 Regulatory RNA Control Experiments

#### S5.1 ChiX sRNA Controls

##### S5.1.1 No sRNA Control

After validating functional sRNA behaviour for Syn-ChiX-20D, we first tested whether the OC6 step-change alone could explain the observed downregulation. Therefore, pSS-01-001 ($P_{VanCC}$-GFPmut3) was re-characterised under the same inducer step-changes (5.5 µM vanillic acid and 139 nM OC6). As expected, GFPmut3 was not downregulated after the OC6 step-change (Supplementary Fig. S23).

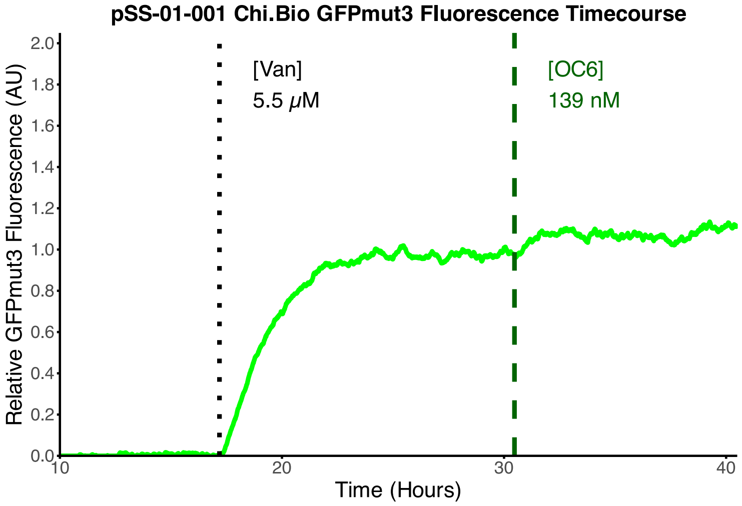


**Figure S23:** $P_{VanCC}$-GFPmut3 (pSS-01-001) tested in response to vanillic acid and OC6 step-changes in a Chi.Bio experiment. Addition of OC6 did not downregulate GFPmut3, supporting that the observed GFPmut3 knockdown in the sRNA circuit is sRNA-dependent.

##### S5.1.2 Binding Region Control

To test whether the engineered binding region of Syn-ChiX-20D was responsible for the observed regulatory activity on GFPmut3, a negative-control version of this sRNA was designed by randomising the 20 nucleotide binding region, generating a putative non-binding sRNA (Syn-ChiX-20NC). Binding by this negative-control sRNA was predicted *in silico* with IntaRNA [8], showing weak predicted binding to the putative GFPmut3 and mScarlet-I mRNAs and to endogenous off-targets (Supplementary Table S1).

In addition to the ten $P_{LuxB}$-sRNA transcription units described in the main Results section, a further $P_{LuxB}$-controlled sRNA transcription unit producing Syn-ChiX-20NC was assembled to form pSS-01-sRNA-NC. This was assembled together with pSS-01-001 to form pSS-02-024, giving a circuit expected to express both GFPmut3 and a non-regulating sRNA. This circuit was tested in the Chi.Bio by inducing GFPmut3 with a step-change from 0 to 5.5 µM vanillic acid, followed by induction of sRNA expression with a step-change from 0 to 139 nM OC6 (Fig. 2E). After induction of the negative-control sRNA, there was no noticeable change in GFPmut3 expression (Fig. 2E), confirming that the programmed binding region is required for GFPmut3 regulation.

##### S5.1.3 Off-Target Binding Control

In addition to testing whether the engineered binding region of Syn-ChiX-20D was required for GFPmut3 regulation, we also tested whether Syn-ChiX-20D unintentionally regulated mScarlet-I. This was important because the intended sponge RNA circuits use both GFPmut3 and mScarlet-I. To do this, an mScarlet-I-expressing transcription unit was assembled: pSS-01-035 ($P_{Tac}$–RiboJ51–pET-RBS–mScarlet-I–L3S2P21). This transcription unit was assembled with pSS-01-sRNA-03 ($P_{LuxB}$-Syn-ChiX-20D) to form pSS-02-092. This circuit was tested in the Chi.Bio platform by inducing mScarlet-I with a step-change in IPTG concentration from 0 to 30 µM, followed by induction of Syn-ChiX-20D with a step-change in OC6 from 0 to 60 nM (Fig. 2F). As expected, there was no meaningful change in mScarlet-I fluorescence after induction of Syn-ChiX-20D (Fig. 2F). A similar microplate-reader experiment was carried out with a version of the circuit lacking RiboJ51 and containing a different RBS (pSS-02-025), again showing no relationship between Syn-ChiX-20D expression and mScarlet-I fluorescence (Supplementary Fig. S24).

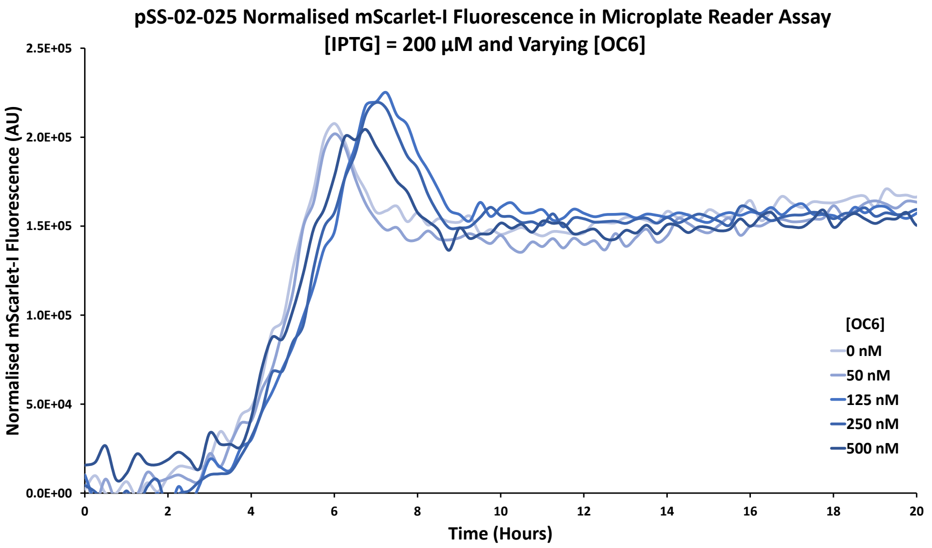


**Figure S24:** Control microplate reader experiment in which Syn-ChiX-20D was induced at varying levels in the presence of mScarlet-I expression. No relationship between Syn-ChiX-20D induction and mScarlet-I fluorescence was observed, supporting that Syn-ChiX-20D regulates GFPmut3 without measurably regulating mScarlet-I.

#### S5.2 ChbBC Sponge RNA Controls

As described in the main text, several control experiments were carried out to verify that the observed sponge RNA-mediated regulation was due to the sponge RNA, required the engineered binding region, and acted through sequestration of the synthetic sRNA. These controls are described in more detail here.

Initial negative controls exposed pSS-02-001 (an equivalent circuit to pSS-02-005 and pSS-02-006 but lacking the sponge RNA/mScarlet-I transcription unit) to the same inducer step-changes, to test whether IPTG addition itself was responsible for the apparent sponge RNA activity. As expected, across three Chi.Bio experiments (two with 139 nM OC6 and 1000 µM IPTG, and one with 50 nM OC6 and 200 µM IPTG), there was no noticeable increase in GFPmut3 after exposure to IPTG (Fig. 4I; Supplementary Fig. S25A–B). This validates that the sponge RNA transcription unit is required for the increase in GFPmut3, supporting the intended mechanism in which the sponge RNA sequesters the sRNA.

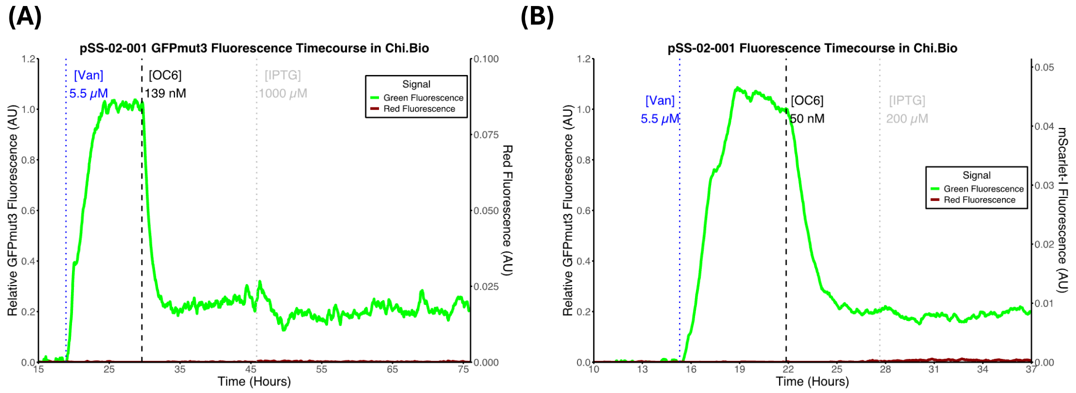


**Figure S25:** Two control experiments testing whether the sponge RNA transcription unit is required for the observed increase in GFPmut3. pSS-02-001, the sRNA circuit, is equivalent to pSS-02-005 or pSS-02-006 (the sponge circuits) but lacks the sponge RNA/mScarlet-I transcription unit. It was characterised in two Chi.Bio experiments under the same inducer step-changes used for sponge RNA circuit characterisation. In both cases, no equivalent increase in GFPmut3 was observed upon IPTG step-change, supporting that IPTG alone does not cause the observed increase in GFPmut3.

Next, a version of Syn-ChbBC-20D was generated in which the programmable binding region was replaced with randomised nucleotides. This produced a sponge RNA with limited IntaRNA [8] predicted binding to Syn-ChiX-20D, generating Syn-ChbBC-20NC ($\Delta G = -10.4$ kcal mol$^{-1}$, compared to $\Delta G = -25.73$ kcal mol$^{-1}$ for Syn-ChbBC-20D). This negative control sponge RNA was assembled into the same circuit architecture as pSS-02-006 (sponge RNA circuit with mScarlet-I), with Syn-ChbBC-20NC replacing Syn-ChbBC-20D, forming pSS-02-027. Across two experiments

(Supplementary Fig. S26 and Fig. 4J), no increase in GFPmut3 was observed upon induction of Syn-ChbBC-20NC, suggesting, as expected, that Syn-ChiX-20D was not sequestered. At the same time, mScarlet-I fluorescence was observed, validating that Syn-ChbBC-20NC was expressed (Supplementary Fig. S26). Together, these results validate that the engineered binding region of Syn-ChbBC-20D is required for the desired regulatory functionality observed in pSS-02-006 and pSS-02-005 (Fig. 3D–E; Fig. 4C–E; Supplementary Fig. S19; Supplementary Fig. S20; Supplementary Fig. S21).

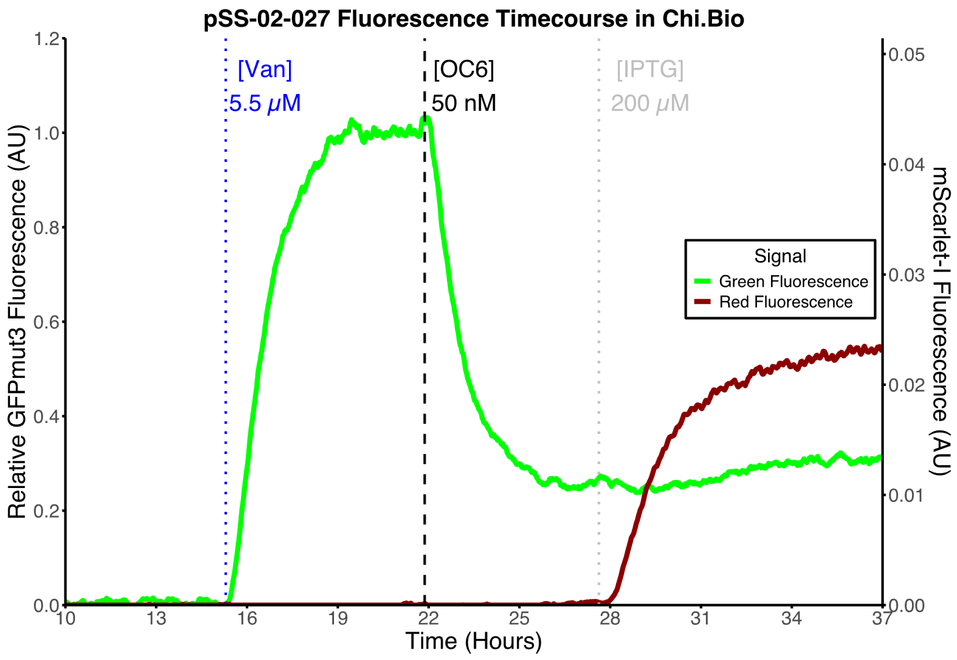


**Figure S26:** Chi.Bio characterisation with relative GFPmut3 fluorescence and normalised mScarlet-I fluorescence for pSS-02-027, a circuit equivalent to pSS-02-006 (sponge circuit with mScarlet-I) but with a randomised spRNA binding region (Syn-ChbBC-20NC) for the sRNA, Syn-ChiX-20D. Upon sponge RNA induction, no increase in GFPmut3 is observed, consistent with loss of binding between sRNA and spRNA, while mScarlet-I fluorescence is still observed, validating expression of Syn-ChbBC-20NC.

To further test, in combination with the previous control, whether Syn-ChbBC-20D operates through binding to Syn-ChiX-20D, we asked whether Syn-ChbBC-20D (with mScarlet-I) could increase GFPmut3 in the absence of Syn-ChiX-20D. A new circuit (pSS-02-NoS) was therefore made equivalent to pSS-02-006 but lacking the sRNA transcription unit. This circuit was tested in the Chi.Bio (Supplementary Fig. S27A). After induction with 50 nM OC6, no change in GFPmut3 was observed, as expected in the absence of sRNA. However, upon induction of Syn-ChbBC-20D (and mScarlet-I) with 200 μM IPTG, GFPmut3 increased approximately 2-fold. This increase in GFPmut3 was likely a result of burden-mediated growth feedback due to high simultaneous expression of both mScarlet-I and GFPmut3 in the absence of the sRNA. We therefore growth-rate normalised the fluorescence signals by multiplying fluorescence by growth rate (Supplementary Fig. S27C). After this normalisation, GFPmut3 remained approximately unchanged after induction of Syn-ChbBC-20D, supporting that Syn-ChbBC-20D only causes an increase in GFPmut3 when Syn-ChiX-20D is present. The same experiment and analysis were carried out with a version of the sponge circuit in which Syn-ChiX-20D was replaced with Syn-ChiX-20NC, and the same behaviour was observed (Supplementary Fig. S27B,D).

Finally, to test more directly whether Syn-ChiX-20D and Syn-ChbBC-20D bind each other, we used a circuit lacking the GFPmut3 transcription unit. This circuit (pSS-02-090) contained a Syn-ChbBC-20D mScarlet-I transcription unit under the $P_{Tac}$ promoter and a Syn-ChiX-20D transcription unit under the $P_{LuxB}$ promoter. The circuit was tested in the Chi.Bio by first inducing Syn-ChbBC-20D mScarlet-I with 1000 μM IPTG, followed by induction of Syn-ChiX-20D with 139 nM OC6 (Supplementary Fig. S28). In this experiment, mScarlet-I was downregulated, though less strongly than GFPmut3 in the corresponding reporter circuit. Together with the other control experiments, this supports the conclusion that Syn-ChiX-20D and Syn-ChbBC-20D are binding partners. This also suggests that Syn-ChiX-20D can downregulate the Syn-ChbBC-20D mScarlet-I transcription unit to some extent, while other experiments show that Syn-ChbBC-20D downregulates Syn-ChiX-20D (Fig. 3D–E; Fig. 4C–E; Supplementary Fig. S19; Supplementary Fig. S20; Supplementary Fig. S21). Together, these data are consistent with a mutual annihilation-like interaction between Syn-ChiX-20D and Syn-ChbBC-20D, which our final mathematical model represents as symmetric mutual annihilation (Supplementary Note S7). Other plausible mechanisms could include Syn-ChiX-20D sequestering Hfq, reducing Syn-ChbBC-20D mScarlet-I transcript stability and therefore reducing its expression level. However, in

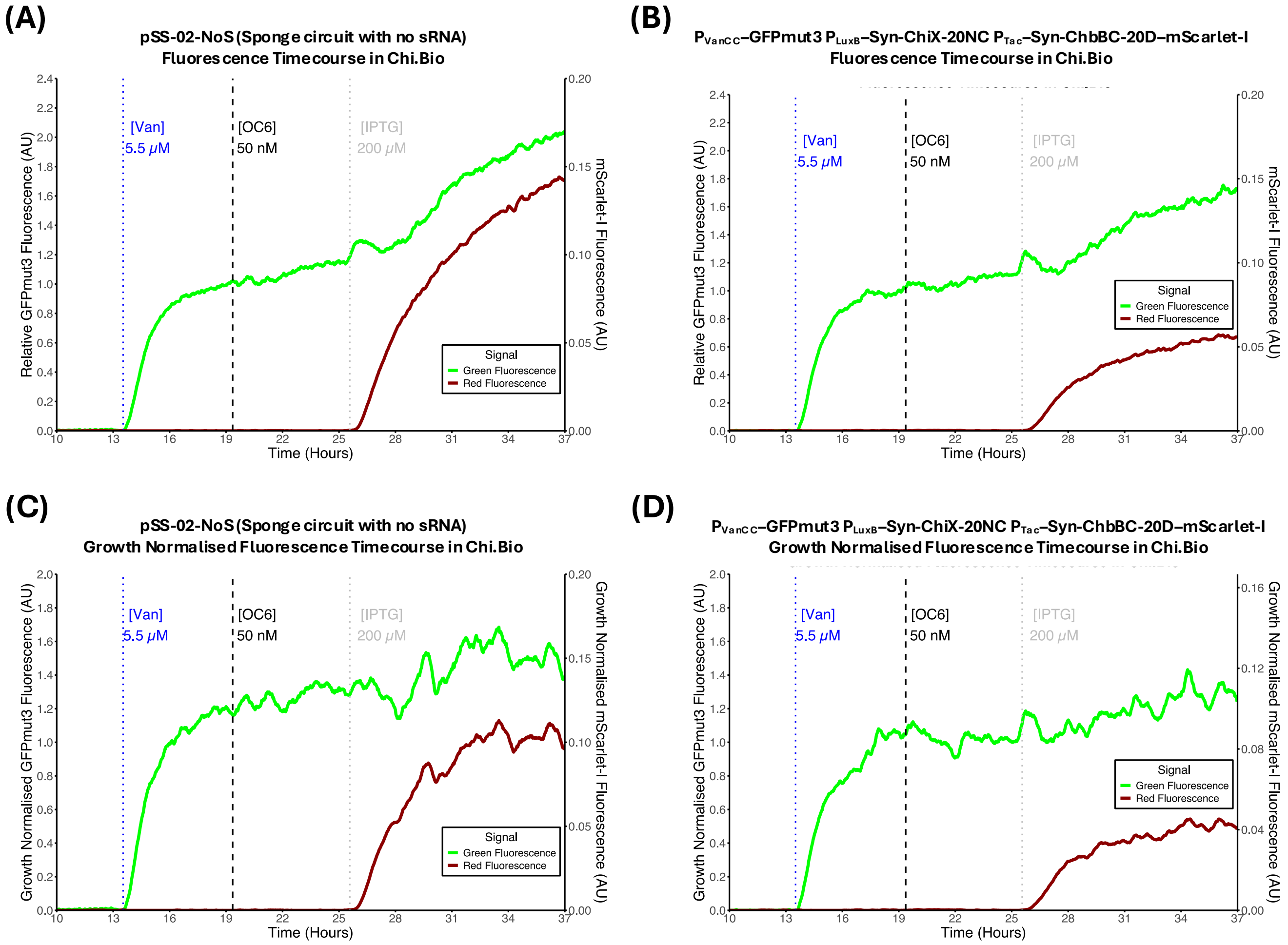


**Figure S27:** Syn-ChbBC-20D negative controls testing whether Syn-ChbBC-20D increases GFPmut3 through specific base-pairing to Syn-ChiX-20D. **(A)** In pSS-02-NoS, no sRNA is present and therefore Syn-ChbBC-20D should not have a binding partner. Addition of 50 nM OC6 did not downregulate GFPmut3, as expected in the absence of sRNA. Induction of Syn-ChbBC-20D with 200 μM IPTG produced an approximately 2-fold increase in GFPmut3 while mScarlet-I was also expressed. **(B)** Syn-ChbBC-20D is again present, but Syn-ChiX-20D is replaced by a negative-control version with a randomised binding region (Syn-ChiX-20NC), so binding between the spRNA and sRNA should be strongly reduced. The same behaviour is observed in response to the sequential induction steps. **(C–D)** Changes in GFPmut3 expression were no longer observed when fluorescence timecourses were normalised by growth rate, suggesting that the observed changes in Panels A and B resulted from gene-expression burden and further supporting that Syn-ChbBC-20D operates by specific base-pairing to Syn-ChiX-20D.

combination with our other data, a direct interaction appears more likely.

##### S5.2.1 ChbBC sponge RNA circuit regulating mScarlet-I

As described in Supplementary Note S3.2, we also engineered new versions of Syn-ChiX and Syn-ChbBC in which Syn-ChiX was designed to target an mScarlet-I mRNA, generating Syn-ChiX-20D-mSc and a cognate synthetic spRNA, Syn-ChbBC-20D-mSc (Supplementary Table S2). These new regulatory RNAs, together with an mScarlet-I transcription unit, were assembled to form the circuit pSS-02-040 (same form as pSS-02-005; Fig. 3C), which was tested in duplicate in the Chi.Bio platform (Supplementary Fig. S29). The circuit behaved as desired, with Syn-ChiX-20D-mSc expression leading to a reduction in mScarlet-I fluorescence, followed by partial recovery of that fluorescence upon Syn-ChbBC-20D-mSc expression (Supplementary Fig. S29). In this case, our one-shot design of both the sRNA and spRNA demonstrates that the design process can, at least in some cases, be adapted to an arbitrary protein of interest without re-optimisation.

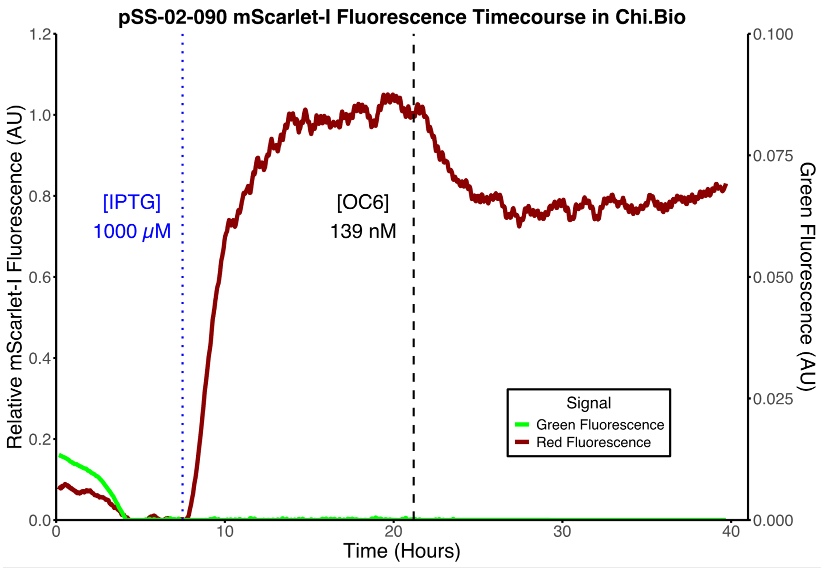


**Figure S28:** Chi.Bio characterisation of pSS-02-090, a circuit containing Syn-ChiX-20D and a Syn-ChbBC-20D–mScarlet-I transcription unit but lacking GFPmut3. Syn-ChbBC-20D and mScarlet-I were first induced with 1000 µM IPTG, producing the expected increase in mScarlet-I fluorescence. This was followed by induction of Syn-ChiX-20D with 139 nM OC6, which led to a reduction in mScarlet-I fluorescence, supporting a direct interaction between Syn-ChiX-20D and Syn-ChbBC-20D.

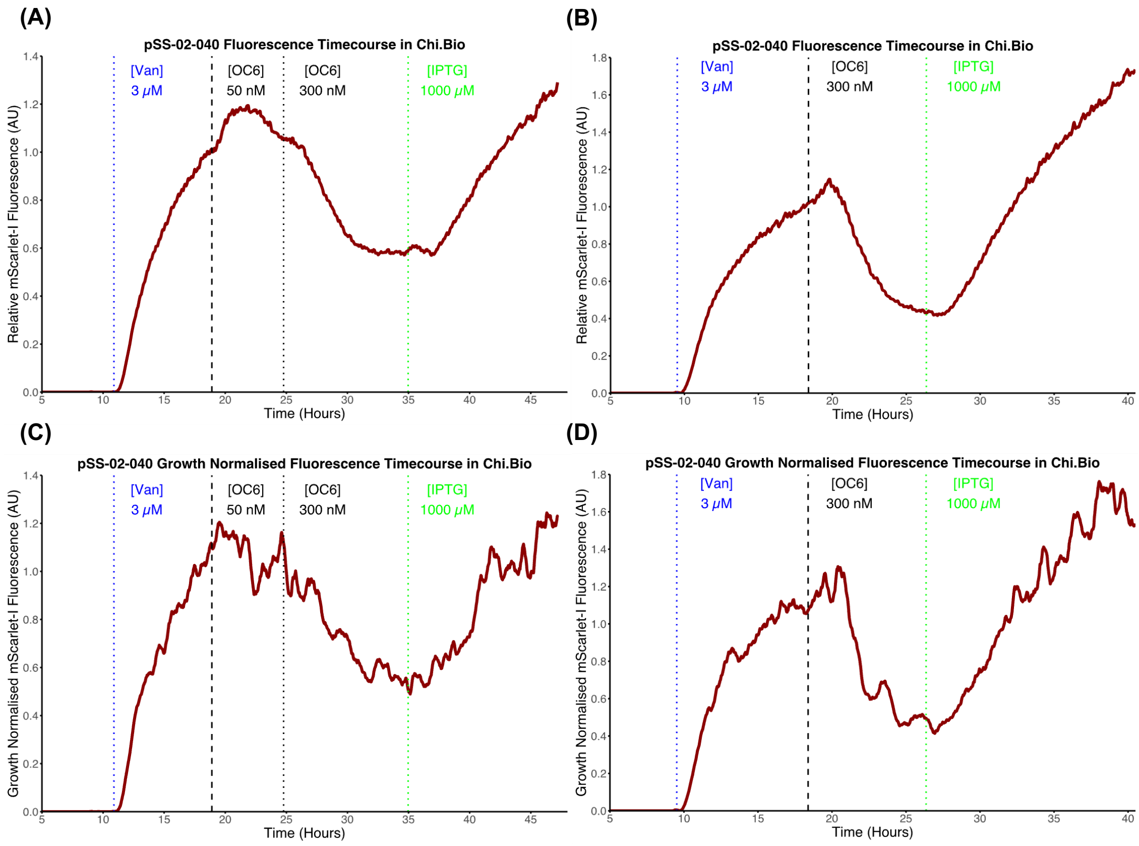


**Figure S29: Chi.Bio characterisation of pSS-02-040**, an equivalent circuit to pSS-02-005 (sponge circuit without mScarlet-I) but where the sRNA regulates mScarlet-I instead of GFPmut3, and the spRNA binds to this new sRNA. Following sequential induction of each transcription unit, the circuit behaves as expected, with initial mScarlet-I expression knocked down by sRNA induction and recovered by spRNA induction, validating that our sRNA–spRNA system can be used to regulate arbitrary protein targets. **(A–B)** show two replicates for pSS-02-040 characterisation. **(C–D)** show the same experiments but fluorescence is normalised to growth rate, validating that changes in mScarlet-I expression are not due to changes in dilution rate.

#### S5.3 Multi-target regulatory RNAs

##### S5.3.1 Multi-target small RNAs

As described in Supplementary Note S3.6, ChiX-derived and MicC-derived multi-target sRNAs were designed and assembled into circuits for the regulation of GFPmut3 and mScarlet-I. The ChiX-based multi-target sRNA was tested in pSS-02-070, and the MicC-based multi-target sRNA was tested in pSS-02-071, both of the form shown in Supplementary Fig. S16. In Chi.Bio characterisation, both circuits worked as intended, leading to simultaneous knockdown of both target genes (Supplementary Fig. S30).

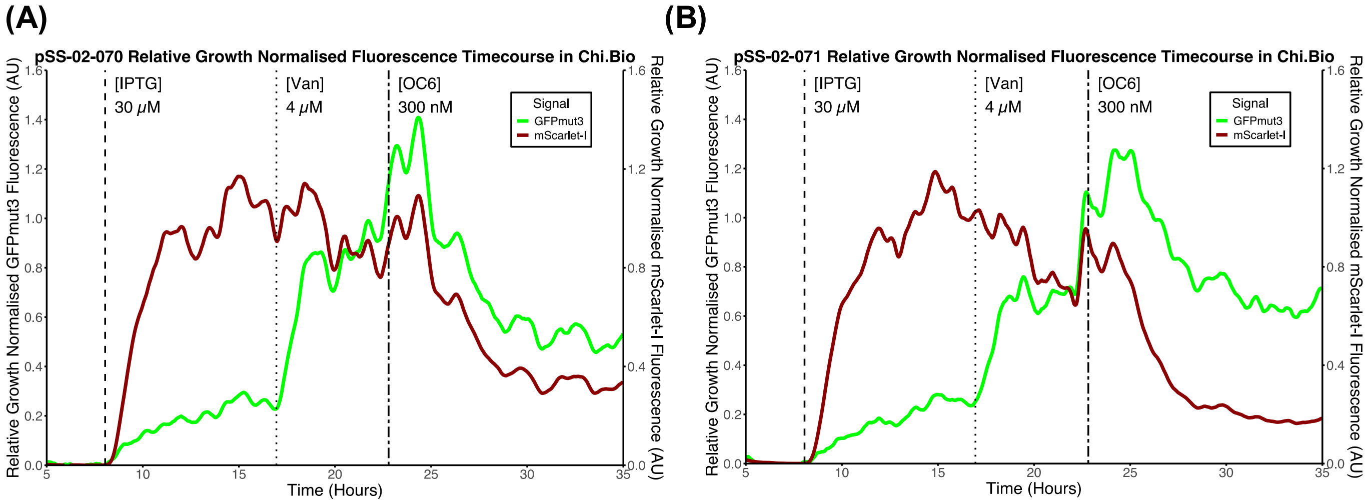


**Figure S30: Chi.Bio characterisation of multi-target sRNA circuits: (A)** pSS-02-070 and **(B)** pSS-02-071, based on ChiX and MicC respectively.

### S6 Quantitative characterisation: absolute quantification and growth-rate estimation

#### S6.1 Absolute quantification

Absolute quantification of intracellular protein concentrations was used in this study to generate quantitative time-courses of gene circuit output expression in the Chi.Bio [1]. This was based on the previously described FPCountR method [20], which we previously adapted for use with Chi.Bio [2]. A brief overview of the process is provided in Supplementary Fig. S31. Applying absolute quantification to these synthetic gene circuit measurements enabled quantitatively meaningful model fitting and comparison across experiments, fluorescent proteins, reactors, and synthetic gene circuits.

First, the microplate reader was calibrated according to the previously described FPCountR method [20, 21]. Overnight overexpression cultures for each fluorescent protein of interest (GFPmut3 and mScarlet-I) were lysed and prepared in serial dilution plates. These plates were then analysed in the plate reader using FPCountR's ECMax assay [20] and the FPBase maximum absorbance for each fluorescent protein [22]. This generated absorbance spectra for each dilution for both GFPmut3 (Supplementary Fig. S32A) and mScarlet-I (Supplementary Fig. S32B) in our specific microplate reader. Data from all dilutions were then used to fit linear models of dilution versus concentration for GFPmut3 (Supplementary Fig. S32C) and mScarlet-I (Supplementary Fig. S32D). By application of the FPCountR fluorescence assay [20] we then related the calculated concentration values of each protein to the measured fluorescence (Supplementary Fig. S32E–F).

Next, samples were taken from Chi.Bio reactors at different timepoints across experiments, and sample fluorescence and OD were measured in triplicate in the calibrated microplate reader. Representative examples are shown in Supplementary Fig. S33, including both the Chi.Bio fluorescence timecourses with sampling points indicated and the associated molar concentrations of fluorescent protein measured in the calibrated microplate reader.

These paired measurements of microplate-reader quantified concentration and Chi.Bio fluorescence at corresponding timepoints were used to fit linear models relating Chi.Bio fluorescence to molar concentration, enabling subsequent conversion from Chi.Bio fluorescence timecourses to molar concentration timecourses. Linear models were fit for each fluorescent protein in each Chi.Bio reactor. The resulting models for GFPmut3 (Supplementary Fig. S34) and mScarlet-I (Supplementary Fig. S35) were all statistically significant ($p \ll 0.05$) and highly explanatory ($R^2 > 0.93$).

These fitted conversion curves were then used to convert Chi.Bio fluorescence intensity per cell timecourses, measured in arbitrary units, into intracellular GFPmut3 and mScarlet-I concentration timecourses (Supplementary Fig. S36). These quantified data were subsequently used for model fitting.

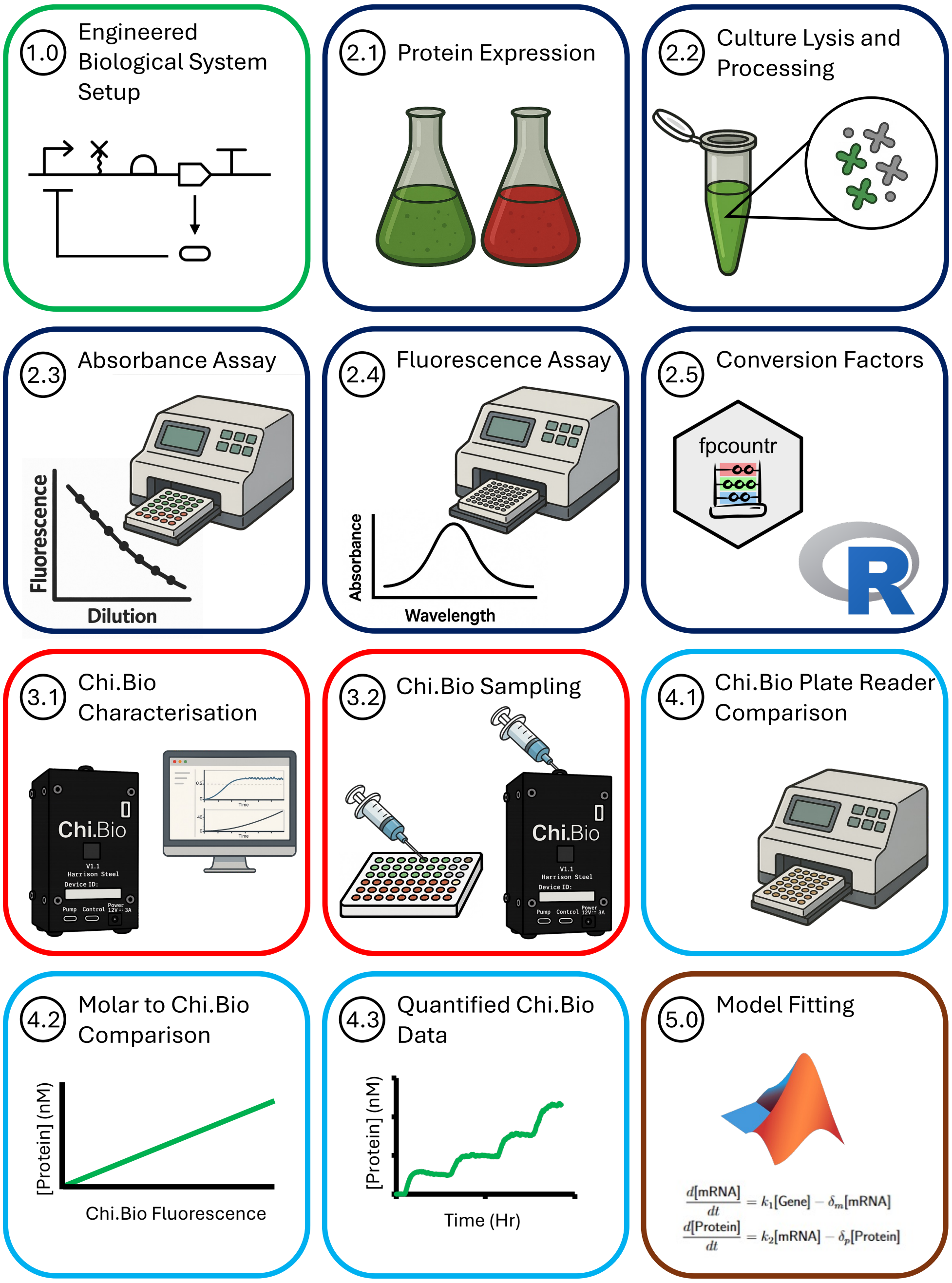


**Figure S31: Graphical summary of our absolute quantification protocol** [2] for use with the Chi.Bio [1], adapted from FPCountR [20]. Briefly, (1.0) a synthetic gene circuit is set up; fluorescent proteins are then (2.1) expressed and (2.2) lysed, concentrated, and prepared in a serial dilution plate. This plate is measured in a microplate reader using FPCountR's (2.3) absorbance and (2.4) fluorescence assays. (2.5) The FPCountR R package [23] is then used to calculate plate-reader-specific conversion factors from fluorescence intensity to moles of fluorescent protein. Next, (3.1) the circuit is characterised in the Chi.Bio, during which (3.2) samples are collected over time and measured in the same plate reader. These measurements are (4.1) converted to intracellular protein concentrations and compared with raw Chi.Bio fluorescence at the corresponding sampling times. (4.2) Linear models are then fitted across all timepoints to relate Chi.Bio fluorescence to molar concentration for each reactor. Finally, (4.3) these fitted models are used to convert Chi.Bio fluorescence timecourses into protein concentration timecourses, enabling downstream applications such as model fitting (5.0).

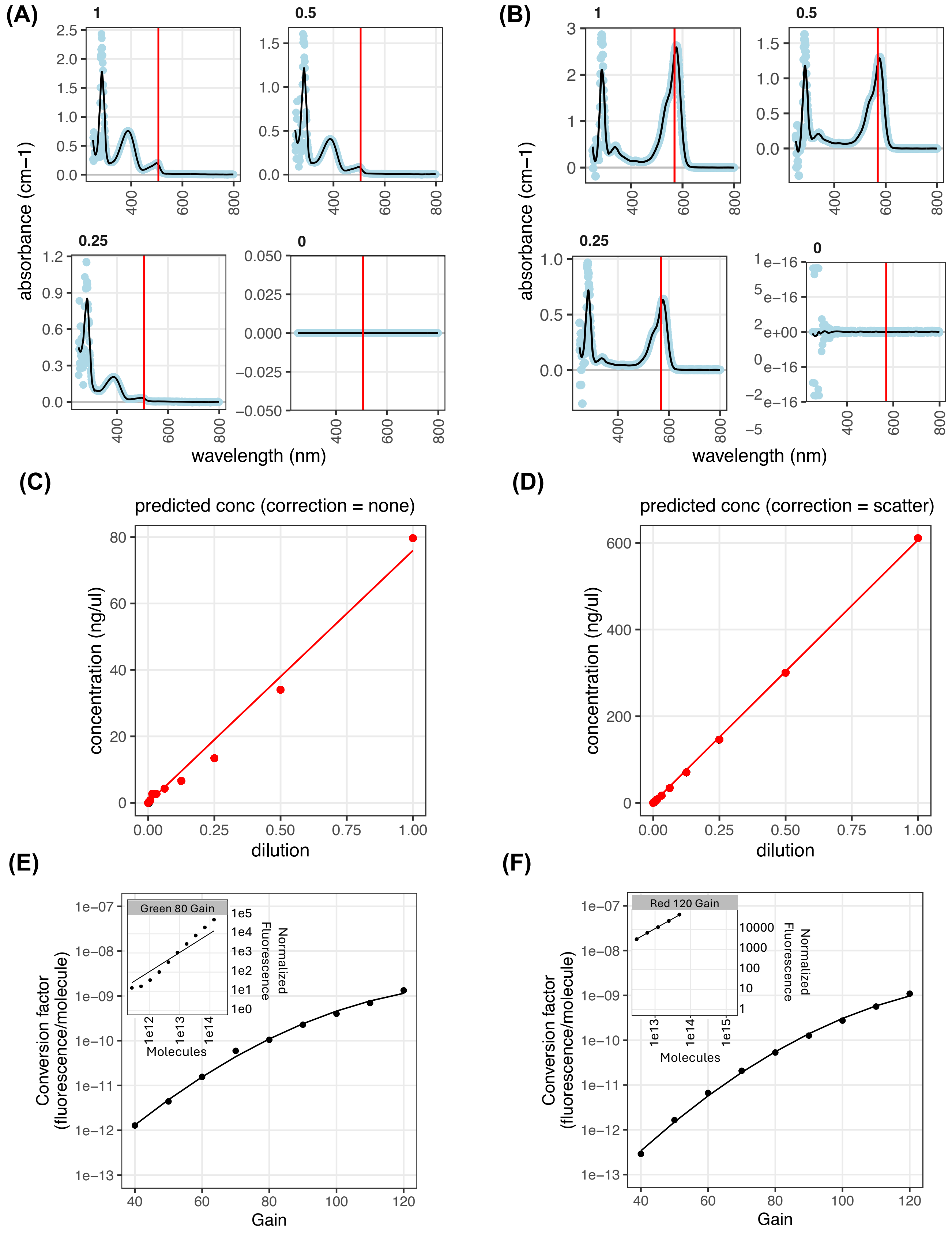


**Figure S32: Absolute quantification calibration of the microplate reader for GFPmut3 and mScarlet-I. (A–B)** Absorbance scans of serial dilution plates for lysed cultures expressing GFPmut3 and mScarlet-I, respectively. The black line shows a LOESS fitted model and the red line shows the FPBase [22] absorbance peak for each fluorescent protein, with each graph corresponding to a different dilution. **(C–D)** Linear fits of dilution versus concentration for each fluorescent protein calculated through the ECMax assay [20]. **(E–F)** Calculated conversion factors between fluorescence and number of molecules for each gain value for each fluorescent protein.

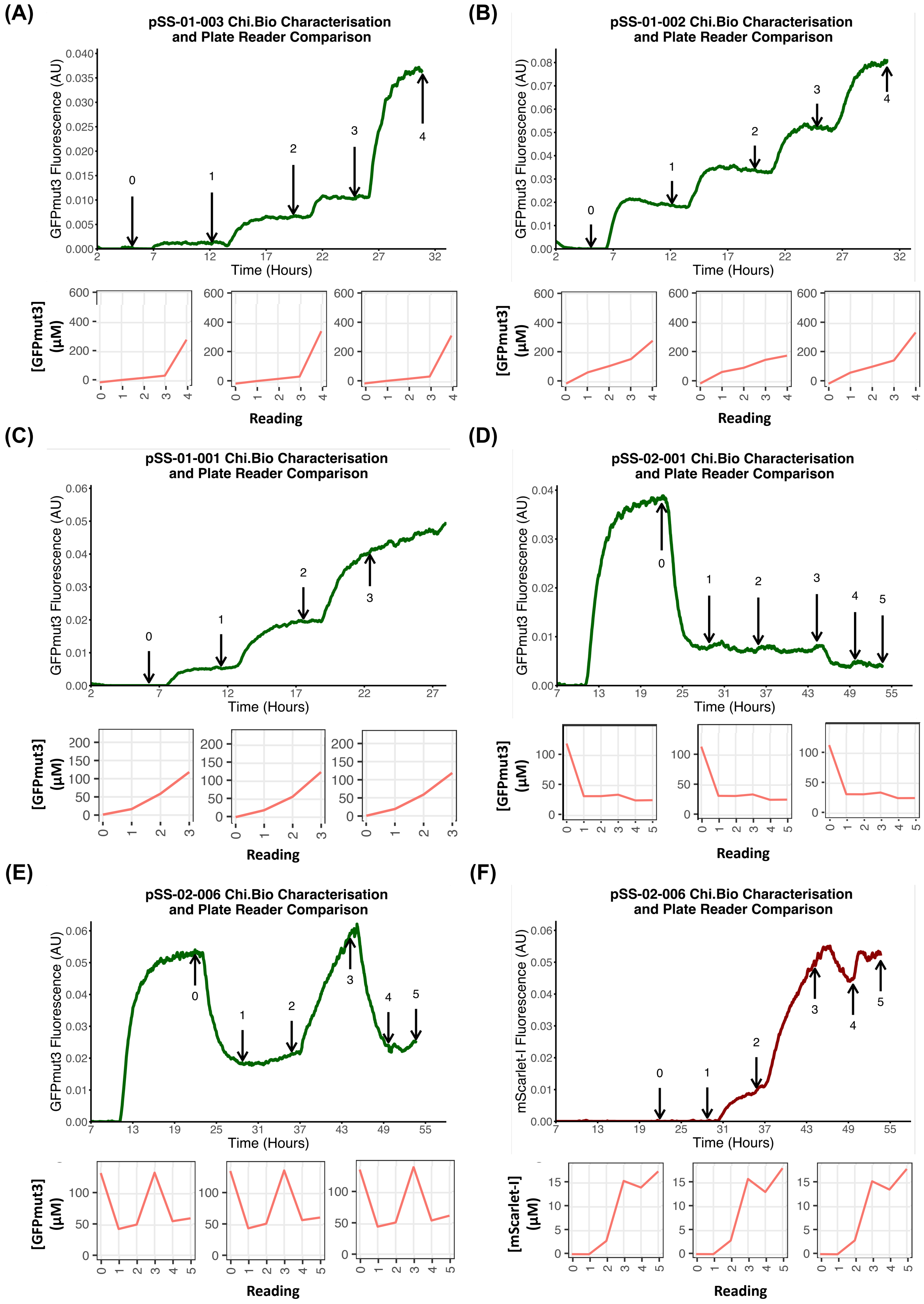


**Figure S33: Chi.Bio experiment sampling and plate reader comparison.** Chi.Bio experiments were sampled and measured in the calibrated microplate reader in triplicate to give measured fluorescent protein concentration at each sampling timepoint. Arrows in fluorescence timecourses are sampling timepoints which correspond to readings in the displayed concentration plots. Representative plots are shown for **(A)** pSS-01-003 ($P_{Tac}$-GFP); **(B)** pSS-01-002 ($P_{LuxB}$-GFP); **(C)** pSS-01-001 ($P_{VanCC}$-GFP); **(D)** pSS-02-001 (sRNA circuit); **(E)** pSS-02-006 GFP signal (sponge circuit with mScarlet-I); and **(F)** pSS-02-006 mScarlet-I signal.

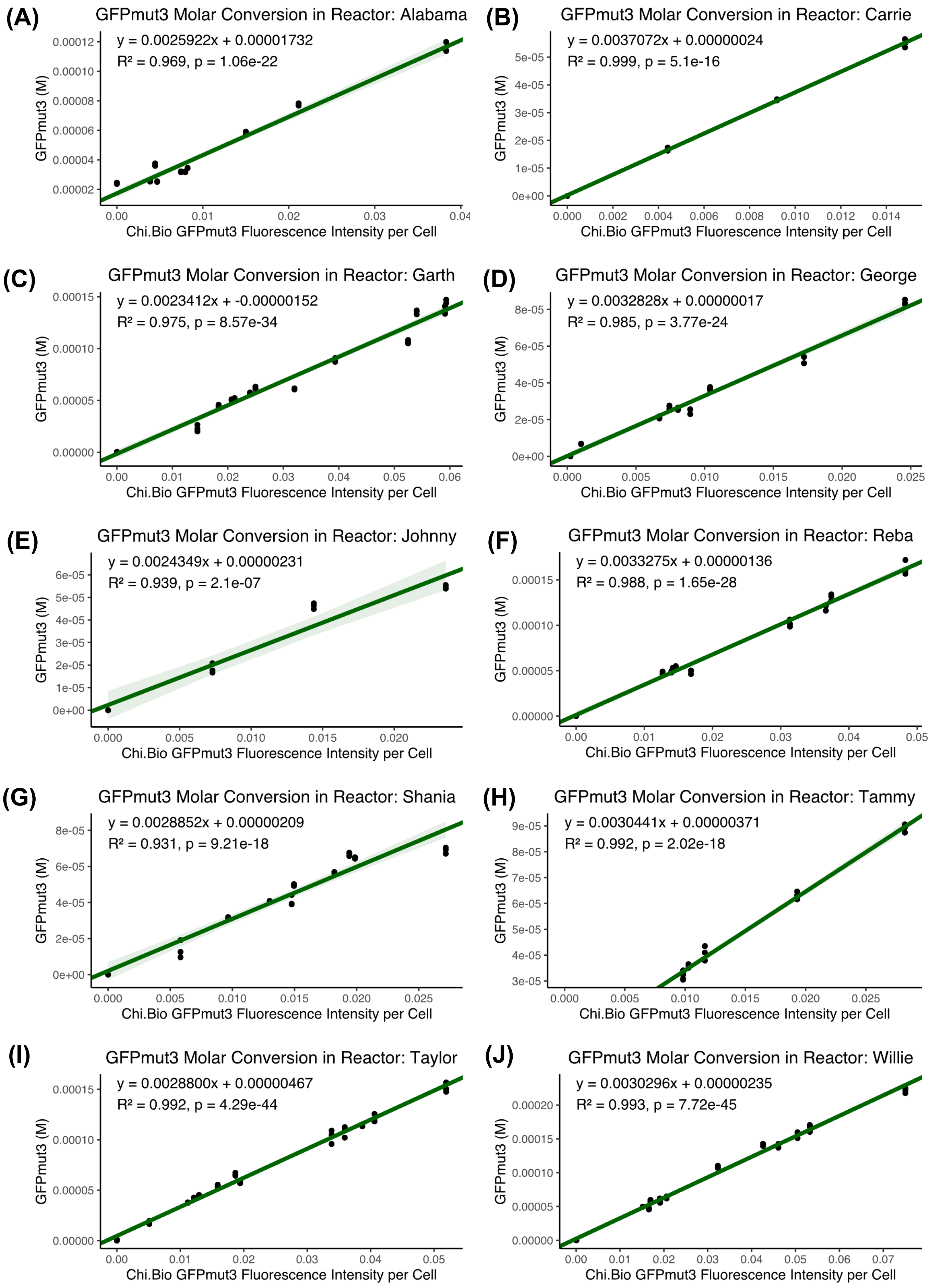


**Figure S34: Chi.Bio molar conversion curves for GFPmut3:** Linear models of the form $y = mx + c$ were fitted to Chi.Bio-to-plate-reader comparison data and used to convert GFPmut3 fluorescence intensity per cell in arbitrary units to intracellular GFPmut3 concentration. Graphs are shown for each unique Chi.Bio reactor to which absolute quantification was applied. All graphs additionally show the fitted parameters, $R^2$, and $p$.

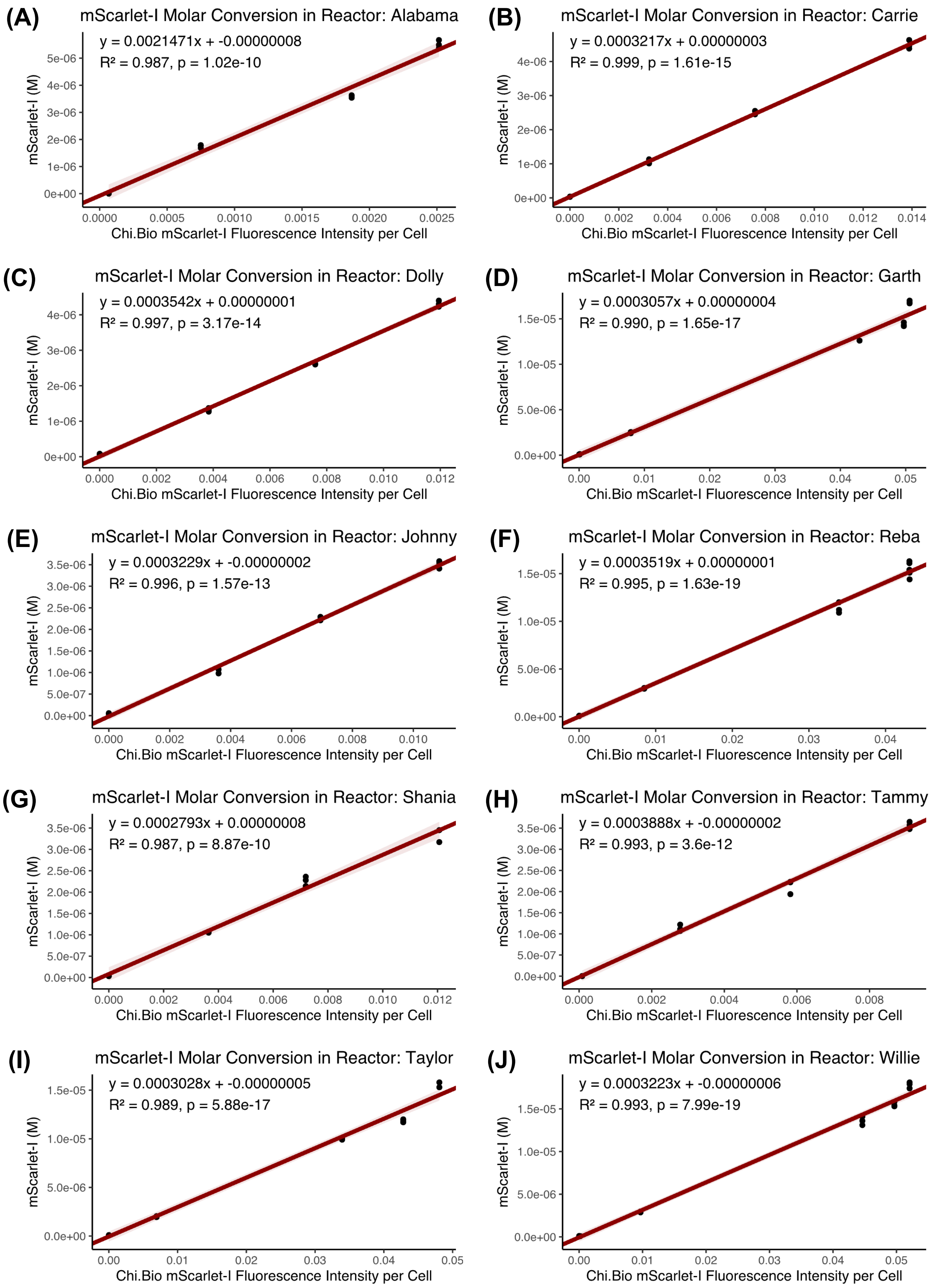


**Figure S35: Chi.Bio molar conversion curves for mScarlet-I:** Linear models of the form $y = mx + c$ were fitted to Chi.Bio-to-plate-reader comparison data and used to convert mScarlet-I fluorescence intensity per cell in arbitrary units to intracellular mScarlet-I concentration. Graphs are shown for each unique Chi.Bio reactor to which absolute quantification was applied. All graphs additionally show the fitted parameters, $R^2$, and $p$.

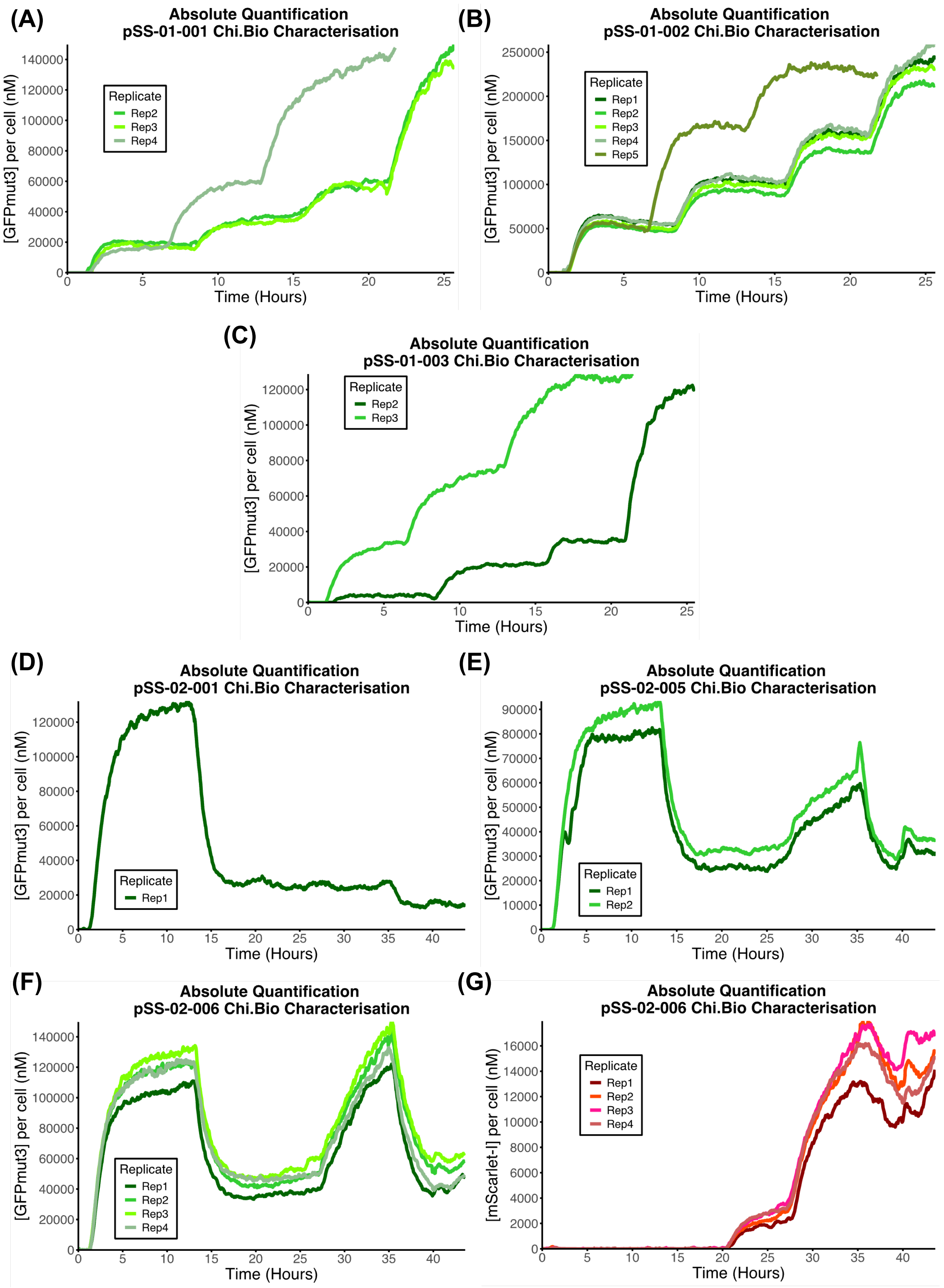


**Figure S36: Intracellular fluorescent protein timecourses from Chi.Bio characterisation:** **(A)** Three quantified timecourses for pSS-01-001 ($P_{VanCC}$-GFPmut3) in response to sequential induction step changes of 3 µM, 4 µM, 5 µM, and 7 µM vanillic acid. One repeat was carried out in a separate experiment and did not use 4 µM. **(B)** Five quantified timecourses for pSS-01-002 ($P_{LuxB}$-GFPmut3) in response to sequential induction step changes of 10 nM, 15 nM, 25 nM, and 50 nM OC6. One replicate was carried out in a separate experiment and instead used 10 nM, 30 nM, and 50 nM OC6. **(C)** Two quantified timecourses for pSS-01-003 ($P_{Tac}$-GFPmut3) in response to sequential induction step changes of 15 µM, 30 µM, 40 µM, and 80 µM or 40 µM, 60 µM, and 80 µM IPTG. **(D)** Quantified timecourse for pSS-02-001 (sRNA circuit) in response to sequential induction step changes of 5.5 µM vanillic acid, 20 nM OC6, and 100 nM OC6. **(E)** Duplicate quantified timecourses for pSS-02-005 (spRNA circuit without mScarlet-I) in response to sequential induction step changes of 5.5 µM vanillic acid, 20 nM OC6, 200 µM IPTG, 100 nM OC6, and 1000 µM IPTG. **(F–G)** Quadruplicate quantified timecourses for pSS-02-006 (spRNA circuit with mScarlet-I) in response to sequential induction step changes of 5.5 µM vanillic acid, 20 nM OC6, 200 µM IPTG, 100 nM OC6, and 1000 µM IPTG.

#### S6.2 Growth-rate estimation

As described in the Results and Methods sections of the main text, in addition to absolute quantification of fluorescent protein concentrations, we used GOFFREDOpy [24] for more accurate estimation of turbidostat culture growth rates for quantitative characterisation and model fitting of sponge RNA circuits. In this section, we briefly present example data demonstrating the resulting growth-rate predictions (Supplementary Fig. S37). In addition to the quantified fluorescent protein timecourses, we used these growth-rate estimates in model fitting.

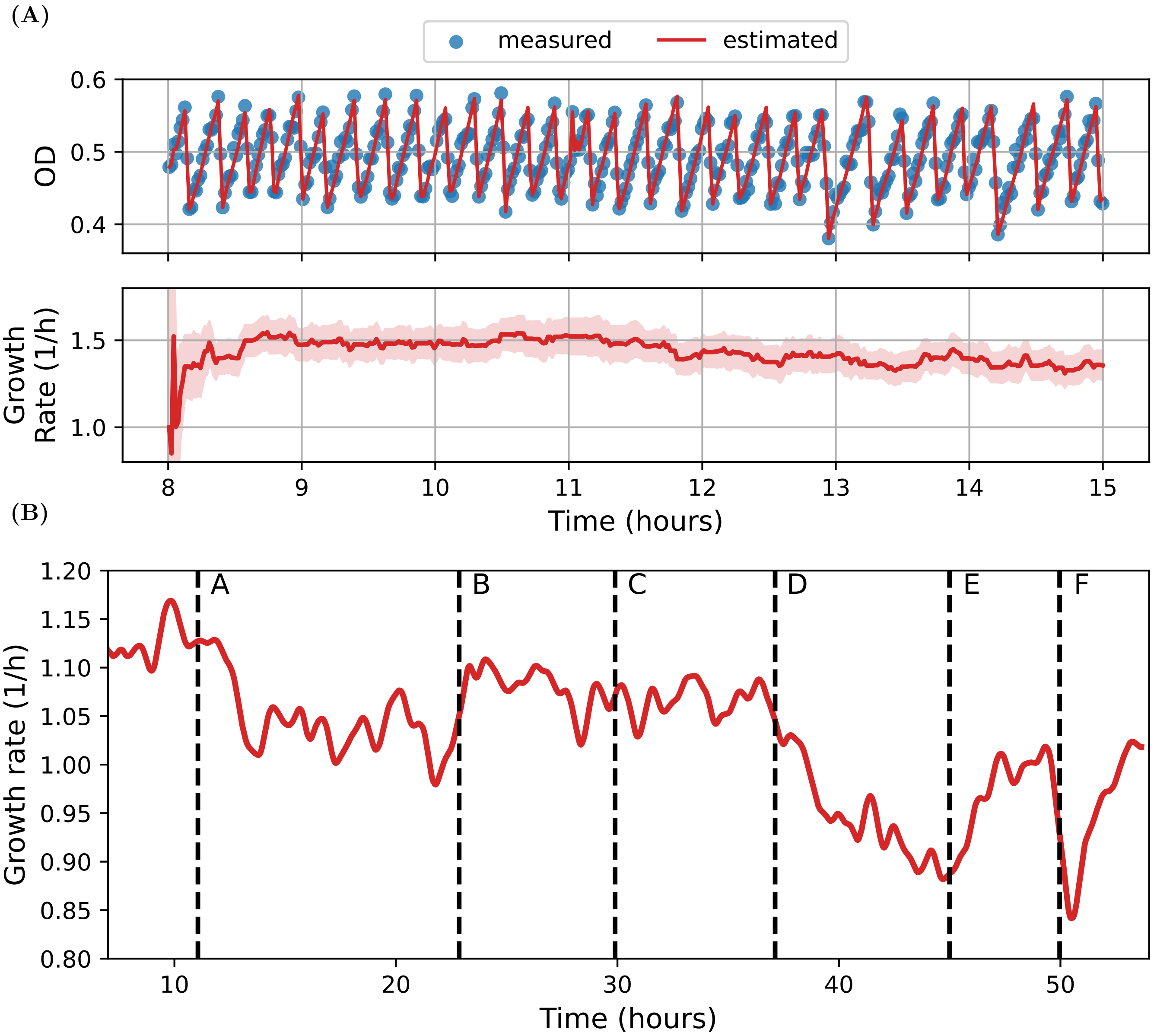


**Figure S37: Dynamic growth-rate estimation from measured OD traces in the turbidostat. (A)** Example output generated by applying the growth-rate estimation pipeline to measured OD traces in the Chi.Bio turbidostat. The tool generates a probabilistic estimate of the OD and growth rate of the culture (red lines) based on noisy OD measurements from the turbidostat (blue points), assuming an underlying exponential growth model. The ribbon around the growth-rate estimate represents the 95% credible interval around the mean estimate. For clarity, uncertainty bands are only reported for the growth-rate estimates. **(B)** Tracking growth changes during the experiments used for model fitting. Shown here are example growth-rate estimates generated from OD measurements in a multi-step induction experiment involving the pSS-02-006 construct (see main text for details). Labelled timepoints (dashed vertical lines) correspond to: A: induction of GFPmut3 with 5.5 µM vanillic acid; B: induction of GFPmut3-targeting sRNA with 20 nM OC6; C: induction of the sRNA-targeting spRNA with 60 µM IPTG; D: further induction of the spRNA (200 µM IPTG); E: further induction of the sRNA (100 nM OC6); F: further induction of the spRNA (1000 µM IPTG). The growth-rate changes accompanying each experimental phase reflect different levels of cellular burden associated with changing expression levels of the fluorescent proteins involved in the circuit. Resolving these changes and accounting for them during model fitting allows more accurate estimation of model parameters.

### S7 Model Fitting

Across all modelling stages, parameter estimation used the shared-parameter framework described in the Methods, with promoter, translation, and RNA-specific parameters tied across constructs where biologically justified. The optimisation approach and weighted-average cost function are also described in the Methods section of the main text.

#### S7.1 Promoter response function fitting

Promoter-characterisation data for pSS-01-001, pSS-01-002, and pSS-01-003 were fit independently of the RNA-circuit datasets using a Hill-type promoter response function coupled to an ODE model of transcription, translation, and fluorescent protein maturation. Parameters were estimated simultaneously across all promoter datasets ($P_{\mathrm{VanCC}}$, $P_{\mathrm{LuxB}}$, and $P_{\mathrm{Tac}}$) using the shared-parameter fitting framework described in the Methods section of the main text. Promoter parameters were fit independently of the RNA-circuit datasets in order to reduce the extent to which assumptions about downstream regulatory mechanisms could bias the promoter parameter estimates.

For each promoter, transcription was described by the response function,

$$f(U(t)) = k_0 + (k_1 - k_0)\,\frac{U(t)^n}{U(t)^n + K^n}, \tag{S1}$$

where $f(U(t))$ is the transcription rate as a function of inducer concentration $U(t)$; $k_0$ and $k_1$ are the basal and maximal transcription rates respectively; $K$ is the half-maximal inducer concentration; and $n$ is the Hill coefficient.

This response function was embedded in the ODE model,

$$\begin{aligned}
\frac{dM}{dt} &= f(U(t)) - (\mu(t) + \delta_m)\,M \\
\frac{dP}{dt} &= \theta M - \tau P - \mu(t)P \\
\frac{dF}{dt} &= \tau P - \mu(t)F,
\end{aligned} \tag{S2}$$

where $M$ is mRNA concentration, $P$ is immature fluorescent protein concentration, and $F$ is mature fluorescent protein concentration. Here, $\mu(t)$ is the experimentally estimated growth rate, $\delta_m$ is the mRNA degradation rate, $\theta$ is the GFPmut3 translation rate, and $\tau$ is the GFPmut3 maturation rate. During promoter fitting, $\theta$ was estimated from the data, while $\delta_m$ and $\tau$ were fixed based on reported values (Supplementary Table S5).

Promoter model fits are shown in Supplementary Fig. S38 and Fig. 4B. These fits provided good agreement with the experimental data across replicates and were used to obtain initial promoter and GFPmut3 translation parameter estimates that were later carried forward into RNA-circuit fitting. Estimated and fixed parameter values are reported in Supplementary Table S5. The GFPmut3 mRNA degradation rate was calculated from a previously reported YFP mRNA half-life of 6.8 minutes [25], giving $\delta_m = \ln 2/(6.8 \times 60) = 0.001699\ \mathrm{s}^{-1}$. This value is in line with typical mRNA half-lives in *E. coli* [26], and was considered reasonable in the absence of direct mRNA stability measurements. The GFPmut3 maturation rate was calculated from a reported maturation half-time of 4.1 minutes [22], giving $\tau = \ln 2/(4.1 \times 60) = 0.002818\ \mathrm{s}^{-1}$.

Although the promoter and translation parameters fit pSS-01-001, pSS-01-002, and pSS-01-003 well, there are limitations in the underlying assumptions. Without direct measurements of mRNA abundance and degradation, identifiability is constrained by redundancy between transcription and translation rates, although time-resolved Chi.Bio data and simultaneous fitting with shared parameters partially mitigate this relative to steady-state measurements of isolated transcription units alone.

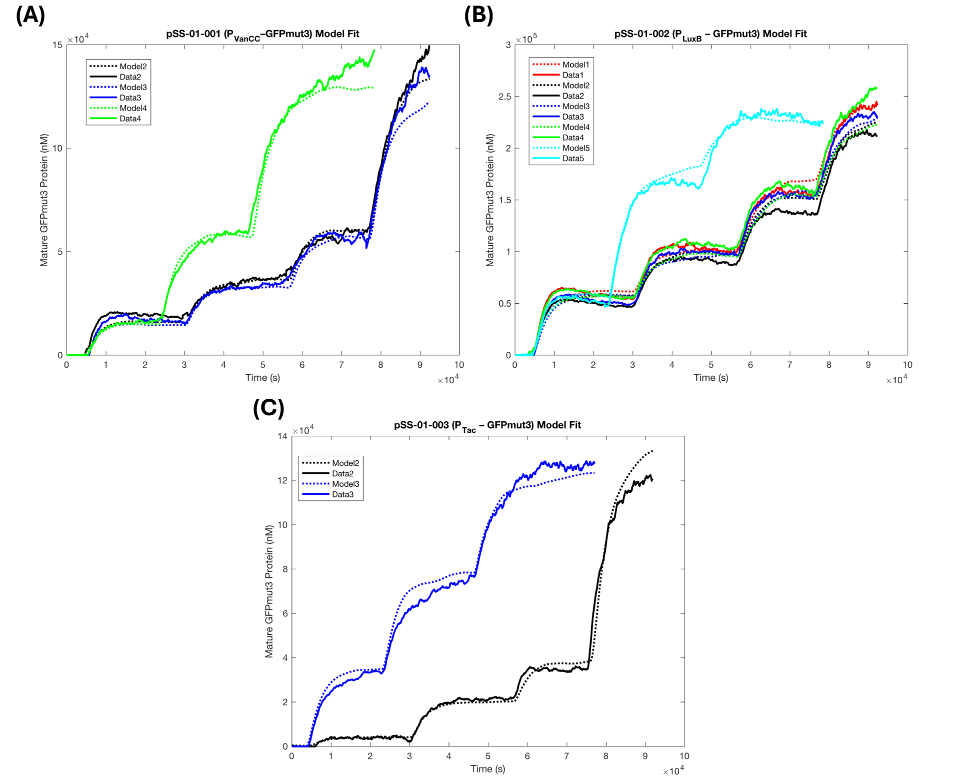


**Figure S38: Initial model fits to standardised promoter constructs: (A)** $P_{VanCC}$–GFPmut3 (pSS-01-001) over three replicates, **(B)** $P_{LuxB}$–GFPmut3 (pSS-01-002) over five replicates, and **(C)** $P_{Tac}$–GFPmut3 (pSS-01-003) over two replicates. Fitting was carried out simultaneously across all constructs and replicates, with shared values for parameters describing common elements of the three standardised constructs (translation rate, GFPmut3 maturation rate, and mRNA degradation rate). The model structure is given by Supplementary Equations S2, and the resulting parameters are given in Supplementary Table S5.

**Table S5: Fitted and chosen parameters for promoter construct model fitting (pSS-01-001, pSS-01-002, and pSS-01-003)**

| **Parameter** | **Description** | **Value** | **Units** | **Source** |
|---|---|---|---|---|
| $k_{0,\mathrm{Van}}$ | Basal transcription rate of the $P_{VanCC}$ promoter | 0.004689 | $\mathrm{nM\,s^{-1}}$ | Fitted in this work |
| $k_{1,\mathrm{Van}}$ | Maximal transcription rate of the $P_{VanCC}$ promoter | 8.193 | $\mathrm{nM\,s^{-1}}$ | Fitted in this work |
| $K_{\mathrm{Van}}$ | Half-maximal activation constant for the $P_{VanCC}$ promoter | 8591 | nM | Fitted in this work |
| $n_{\mathrm{Van}}$ | Hill coefficient for the $P_{VanCC}$ promoter | 2.823 | – | Fitted in this work |
| $k_{0,\mathrm{Lux}}$ | Basal transcription rate of the $P_{LuxB}$ promoter | 0.05624 | $\mathrm{nM\,s^{-1}}$ | Fitted in this work |
| $k_{1,\mathrm{Lux}}$ | Maximal transcription rate of the $P_{LuxB}$ promoter | 5.717 | $\mathrm{nM\,s^{-1}}$ | Fitted in this work |
| $K_{\mathrm{Lux}}$ | Half-maximal activation constant for the $P_{LuxB}$ promoter | 18.77 | nM | Fitted in this work |
| $n_{\mathrm{Lux}}$ | Hill coefficient for the $P_{LuxB}$ promoter | 1.772 | – | Fitted in this work |
| $k_{0,\mathrm{Tac}}$ | Basal transcription rate of the $P_{Tac}$ promoter | 0.004691 | $\mathrm{nM\,s^{-1}}$ | Fitted in this work |
| $k_{1,\mathrm{Tac}}$ | Maximal transcription rate of the $P_{Tac}$ promoter | 5.783 | $\mathrm{nM\,s^{-1}}$ | Fitted in this work |
| $K_{\mathrm{Tac}}$ | Half-maximal activation constant for the $P_{Tac}$ promoter | 79770 | nM | Fitted in this work |
| $n_{\mathrm{Tac}}$ | Hill coefficient for the $P_{Tac}$ promoter | 2.470 | – | Fitted in this work |
| $\theta$ | Translation rate of GFPmut3 | 0.02879 | $\mathrm{s^{-1}}$ | Fitted in this work |
| $\delta_m$ | Degradation rate of GFPmut3 mRNA | 0.001699 | $\mathrm{s^{-1}}$ | Calculated from YFP mRNA half-life in [25] |
| $\tau$ | Maturation rate of GFPmut3 | 0.002818 | $\mathrm{s^{-1}}$ | Calculated from GFPmut3 maturation half-time in [22] |

#### S7.2 Dataset-specific scaling factors in RNA-circuit model fitting

Promoter response and translation parameters were first estimated using isolated promoter–reporter constructs (Supplementary Note S7.1), in which GFPmut3 expression was driven directly by $P_{\mathrm{VanCC}}$, $P_{\mathrm{LuxB}}$, or $P_{\mathrm{Tac}}$. When these parameters were subsequently applied to the RNA-regulatory circuit datasets, the model captured the qualitative expression dynamics but showed modest systematic differences in absolute fluorescence-derived protein concentration between nominally comparable constructs and experimental replicates. We therefore introduced dataset-specific multiplicative scaling factors during RNA-circuit model fitting to account for differences in effective expression scale not explicitly represented by the mechanistic RNA-regulatory model. Here we provide more detail on the scaling factors.

For each RNA-circuit dataset $j$, the model describes the internal concentration of mature fluorescent protein, $F_j(t)$, using the same transcription, translation, maturation, dilution, and RNA-interaction equations described in this section. The corresponding measured fluorescent protein concentration was linked to the model prediction through a fitted multiplicative factor, $SF_j$, such that

$$F_{j,\mathrm{obs}}(t) \approx SF_j F_j(t).$$

Equivalently, when initialising the model from the first measured fluorescent protein value, the initial mature fluorescent protein state was set to $F_{j,\mathrm{obs}}(0)/SF_j$, and model predictions were multiplied by $SF_j$ before comparison with the experimental data. For example, for GFPmut3 in pSS-02-005 replicate 1,

$$F_{\mathrm{GFP,obs}}(t) \approx SF_{5,1} F_{\mathrm{GFP,model}}(t),$$

and analogous factors were used for pSS-02-005 replicate 2, pSS-02-006 replicates 1–4, and the sRNA-only circuit pSS-02-001. In pSS-02-006, the same replicate-specific scaling factor was applied to both GFPmut3 and mScarlet-I measurements from the same dataset, reflecting a dataset-level scale correction rather than a fluorescent protein-specific fitted adjustment.

The scaling factors were included only in the observation model used to compare simulated mature fluorescent protein concentrations with the experimental measurements. They did not alter the mechanistic RNA-regulatory terms, including sRNA–mRNA interaction, sRNA–spRNA interaction, resource-dependent complex formation, RNA degradation, protein maturation, or dilution by growth. Thus, these factors absorb modest differences in absolute expression scale while preserving the inferred dynamic behaviour and relative regulatory effects imposed by the mechanistic model.

Several sources could contribute to these differences. First, promoter and translation parameters estimated from isolated promoter–reporter constructs may not transfer perfectly to full RNA-regulatory circuits due to local genetic context, transcript architecture, and sequence-context effects. Second, nominally matched biological constructs can differ in effective expression due to variation in promoter context, untranslated regions, transcript stability, copy number, or burden. Third, differences between experimental runs, bioreactors, calibration, growth conditions, or background subtraction can introduce modest dataset-level shifts in fluorescence or expression. The fitted scaling factors therefore provide a pragmatic way to link independently characterised expression parameters to the RNA-circuit datasets without forcing scale discrepancies to be absorbed by mechanistic parameters.

The scaling factors were fitted jointly with the RNA-circuit parameters as part of the combined optimisation. To avoid arbitrary rescaling, the factors were constrained to remain close to their initial values during optimisation, and the fitted values corresponded to modest fold changes in expression scale. In our final model the fitted scaling factors corresponded to modest fold changes ($\sim$1.3–2.3$\times$), consistent with reported promoter context dependence [27, 28, 29]. These values indicate that only modest corrections to the absolute predicted protein scale were required. All scaling factors fitted in this work are reported in the appropriate parameter tables.

Importantly, the scaling factors were not used to change the shape or timing of the simulated responses. Because each $SF_j$ multiplies the entire predicted mature fluorescent protein trajectory for a given dataset, it cannot by itself reproduce changes in induction threshold, response timing, curvature, dynamic range, or relative responses to different input conditions within that dataset. These features remain determined by the fitted mechanistic parameters and the measured time-varying inducer and growth-rate inputs. The scaling factors therefore serve as dataset-level calibration parameters rather than as substitutes for the RNA-regulatory mechanism.

#### S7.3 Mass-action annihilation-based model

We next turned to modelling the RNA circuits. In line with previous modelling of sRNA-based gene circuits [30, 31], we first considered a simple mass-action annihilation model in which sRNA–mRNA and sRNA–spRNA interactions were represented as mutual annihilation reactions described by bimolecular mass-action terms. At this stage, proteins encoded by the sponge RNA (i.e. mScarlet-I in pSS-02-006) were not included in the modelling process. Therefore, pSS-02-005 and pSS-02-006 had the same model structure, while pSS-02-001 was modelled without the sponge RNA species.

For the sponge RNA circuits, this model is given by

$$\begin{aligned}
\frac{dM}{dt} &= f\left(V(t)\right) - \left(\mu(t) + \delta_m\right) M - \gamma_{sm} S M \\
\frac{dP}{dt} &= \theta M - \tau P - \mu(t) P \\
\frac{dF}{dt} &= \tau P - \mu(t) F \\
\frac{dS}{dt} &= f\left(O(t)\right) - \left(\mu(t) + \delta_s\right) S - \gamma_{sm} S M - \gamma_{sz} S Z \\
\frac{dZ}{dt} &= f\left(I(t)\right) - \left(\mu(t) + \delta_z\right) Z - \gamma_{sz} S Z \,,
\end{aligned} \tag{S3}$$

where $M$, $P$, $F$, $S$, and $Z$ are the time-varying concentrations of GFPmut3 mRNA, immature GFPmut3, mature GFPmut3, Syn-ChiX-20D, and Syn-ChbBC-20D, respectively. Parameters are defined as in Supplementary Equations S2 and S1, with the addition of $V(t)$, $I(t)$, and $O(t)$, which are the experimentally time-varying concentrations of vanillic acid, IPTG, and OC6, and $\gamma_{sm}$, $\gamma_{sz}$, $\delta_s$, and $\delta_z$, which represent the sRNA–mRNA binding rate, sRNA–spRNA binding rate, sRNA degradation rate, and sponge RNA degradation rate, respectively.

Fits for this model were carried out using the promoter and translation parameters obtained above (Supplementary Table S5), while most of the new RNA-specific parameters were fitted. The degradation rate of Syn-ChiX-20D was fixed rather than fitted and was assumed to match the degradation rate of natural ChiX. It was therefore calculated from the published ChiX half-life of 32 minutes [32], giving $\delta_s = \ln 2/(32 \times 60) = 0.0003610$ s$^{-1}$. To allow construct-specific differences, separate sponge RNA degradation rates and sRNA–spRNA binding rates were fitted for pSS-02-005 (sponge RNA circuit without mScarlet-I) and pSS-02-006 (sponge RNA circuit with mScarlet-I). The previously described scaling factors were also fitted per circuit and replicate (see Supplementary Note S7.2). Fits for this model are shown in Supplementary Fig. S39, and fitted parameter values are given in Supplementary Table S6.

This model was able to fit the pSS-02-001 experimental data well (Supplementary Fig. S39A). However, despite fitting over a large range of biologically feasible parameters, this initial model architecture was unable to recapitulate the experimentally observed behaviour for pSS-02-005 and pSS-02-006 (Supplementary Fig. S39B–G). In particular, upon initial induction of the sponge RNA with 60 µM IPTG at approximately the 20-hour timepoint, the fitted model predicted an increase in GFPmut3 expression (Supplementary Fig. S39B–G). The experimental data showed no such increase in GFPmut3 (Fig. 4C,G). Observed expression of mScarlet-I in pSS-02-006 in the same experiments confirmed that the lack of increase was not due to the sponge RNA being unexpressed (Fig. 4E), suggesting that the discrepancy between the model and experimental data may reflect missing or oversimplified RNA-interaction mechanisms.

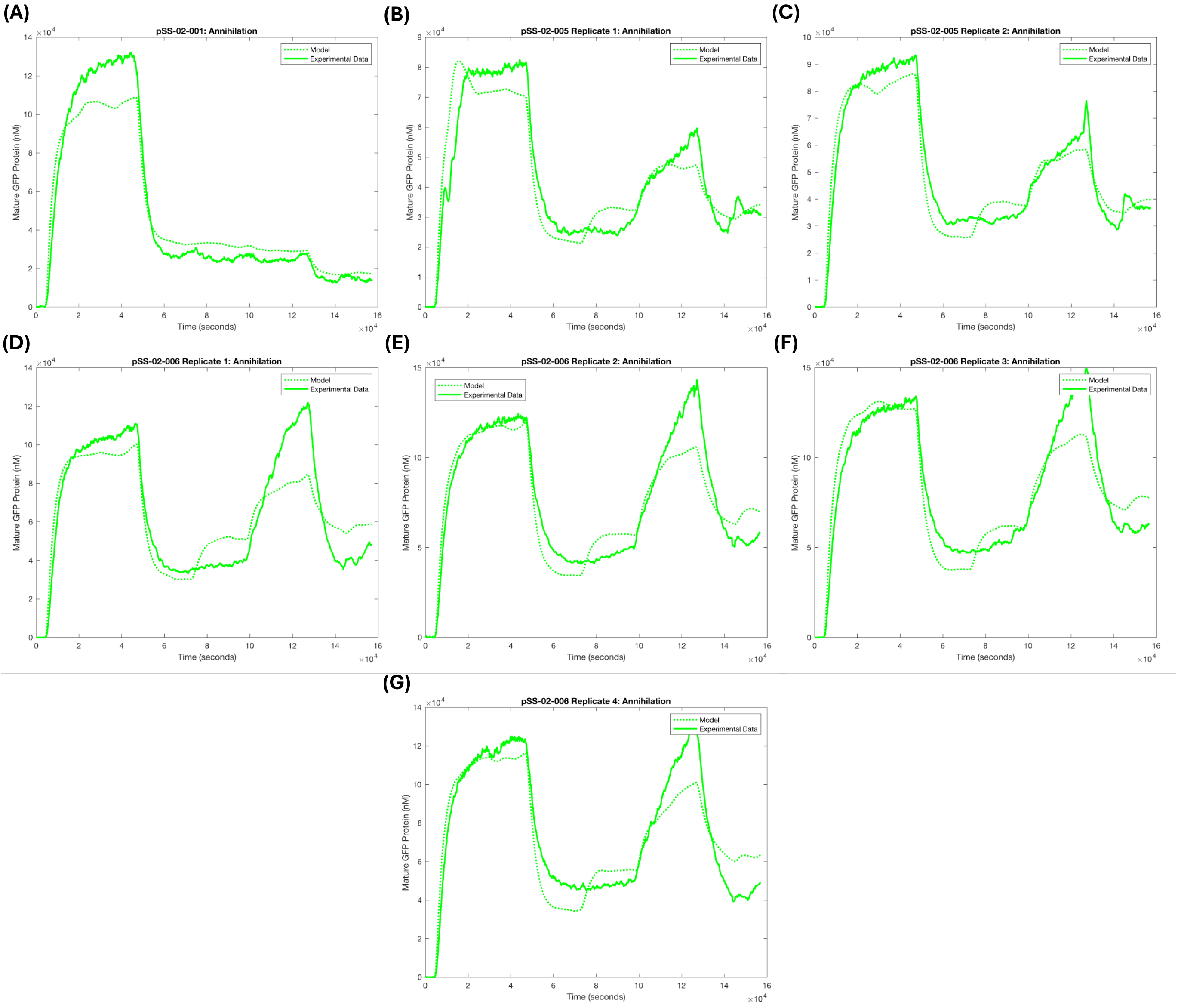


**Figure S39: Mass-action annihilation model fits for RNA circuits: (A)** the sRNA circuit (pSS-02-001); **(B–C)** the sponge RNA circuit without mScarlet-I (pSS-02-005); and **(D–G)** the sponge RNA circuit with mScarlet-I (pSS-02-006). All sponge-circuit models predict a GFPmut3 increase after low spRNA induction at around $7.5 \times 10^4$ seconds, which was not observed experimentally.

**Table S6: Fitted and chosen parameters for RNA regulatory circuit models for the simple mass-action annihilation-based model**

| Parameter | Description | Value | Units | Source |
|---|---|---|---|---|
| $\delta_{z,6}$ | Degradation rate of Syn-ChbBC-20D in pSS-02-006 | 0.002190 | $s^{-1}$ | Fitted in this work |
| $\delta_{z,5}$ | Degradation rate of Syn-ChbBC-20D in pSS-02-005 | 0.0002172 | $s^{-1}$ | Fitted in this work |
| $\gamma_{sz,6}$ | Annihilation rate of sRNA and spRNA in pSS-02-006 | $5.826 \times 10^{-6}$ | $nM^{-1}\,s^{-1}$ | Fitted in this work |
| $\gamma_{sz,5}$ | Annihilation rate of sRNA and spRNA in pSS-02-005 | $6.716 \times 10^{-7}$ | $nM^{-1}\,s^{-1}$ | Fitted in this work |
| $\gamma_{sm}$ | Annihilation rate of sRNA and GFPmut3 mRNA | $1.757 \times 10^{-6}$ | $nM^{-1}\,s^{-1}$ | Fitted in this work |
| $SF_{6,1}$ | Multiplicative scaling factor by which GFPmut3 in pSS-02-006 replicate 1 was scaled | 1.384 | – | Fitted in this work |
| $SF_{6,2}$ | Multiplicative scaling factor by which GFPmut3 in pSS-02-006 replicate 2 was scaled | 1.457 | – | Fitted in this work |
| $SF_{6,3}$ | Multiplicative scaling factor by which GFPmut3 in pSS-02-006 replicate 3 was scaled | 1.633 | – | Fitted in this work |
| $SF_{6,4}$ | Multiplicative scaling factor by which GFPmut3 in pSS-02-006 replicate 4 was scaled | 1.518 | – | Fitted in this work |
| $SF_{5,1}$ | Multiplicative scaling factor by which GFPmut3 in pSS-02-005 replicate 1 was scaled | 1.007 | – | Fitted in this work |
| $SF_{5,2}$ | Multiplicative scaling factor by which GFPmut3 in pSS-02-005 replicate 2 was scaled | 1.180 | – | Fitted in this work |
| $SF_s$ | Multiplicative scaling factor by which GFPmut3 in pSS-02-001 was scaled | 1.743 | – | Fitted in this work |
| $\delta_s$ | Degradation rate of Syn-ChiX-20D | 0.0003610 | $s^{-1}$ | Calculated based on reported half-life of ChiX in [32] |

#### S7.4 Incorporating RNA Predation in Models

That the simple annihilation model could not recapitulate the experimentally observed sponge-circuit dynamics, despite working well for pSS-02-001, suggested that additional mechanistic detail was required to capture the non-linear relationship between sponge RNA expression and GFPmut3 recovery. Given the previously reported catalytic or predatory behaviour of ChiX [15], the relatively uncertain mechanism of ChbBC-mediated regulation, and the poor fits of the annihilation model, we next tested models incorporating combinations of catalytic regulation.

To introduce sRNA predation of mRNA, we modified the annihilation model such that sRNA–mRNA binding still reduced mRNA according to

$$\frac{dM}{dt} = \cdots - \gamma_{sm} SM \tag{S4}$$

but reduced sRNA according to

$$\frac{dS}{dt} = \cdots - n\gamma_{sm} SM, \tag{S5}$$

where $n$ is a predation parameter, with $n = 0$ corresponding to perfect predation and $n = 1$ corresponding to perfect annihilation.

We first investigated the case in which Syn-ChiX-20D predates GFPmut3 mRNA, while the rest of the model remained unchanged compared with the annihilation-based model (Supplementary Equations S3). The promoter parameters were again held constant (Supplementary Table S5), and the parameters previously described for the annihilation-based model (Supplementary Table S6) were refitted together with an additional predation parameter, $n$. Although this model reduced the calculated cost function relative to the simple annihilation model, it was still unable to recapitulate the observed sponge-circuit dynamics. In particular, it retained the predicted GFPmut3 increase after low spRNA induction at approximately $t = 7.5 \times 10^4$ seconds (Supplementary Fig. S40), as in the simple annihilation model (Supplementary Fig. S39).

We next investigated other predation-based variants:

1. $S$ predates $M$ and $Z$ predates $S$ (Supplementary Fig. S41A).
2. $S$ predates both $M$ and $Z$ (Supplementary Fig. S41B).
3. $Z$ predates $S$ (Supplementary Fig. S41C).
4. $S$ predates $Z$ (Supplementary Fig. S41D).

with the asymmetrical RNA-reduction terms described in Supplementary Equations S4 and S5 adapted accordingly. Fitting was again carried out as above, with fixed promoter parameters (Supplementary Table S5), the previously described RNA-model parameters refitted, and up to two different predation parameters ($n_i$) fitted for the asymmetric RNA interactions. Once again, none of these model variants were able to qualitatively recapitulate the observed sponge-circuit behaviour (Supplementary Fig. S41), with the experimental data at approximately $t = 7.5 \times 10^4$ seconds again diverging from the model predictions.

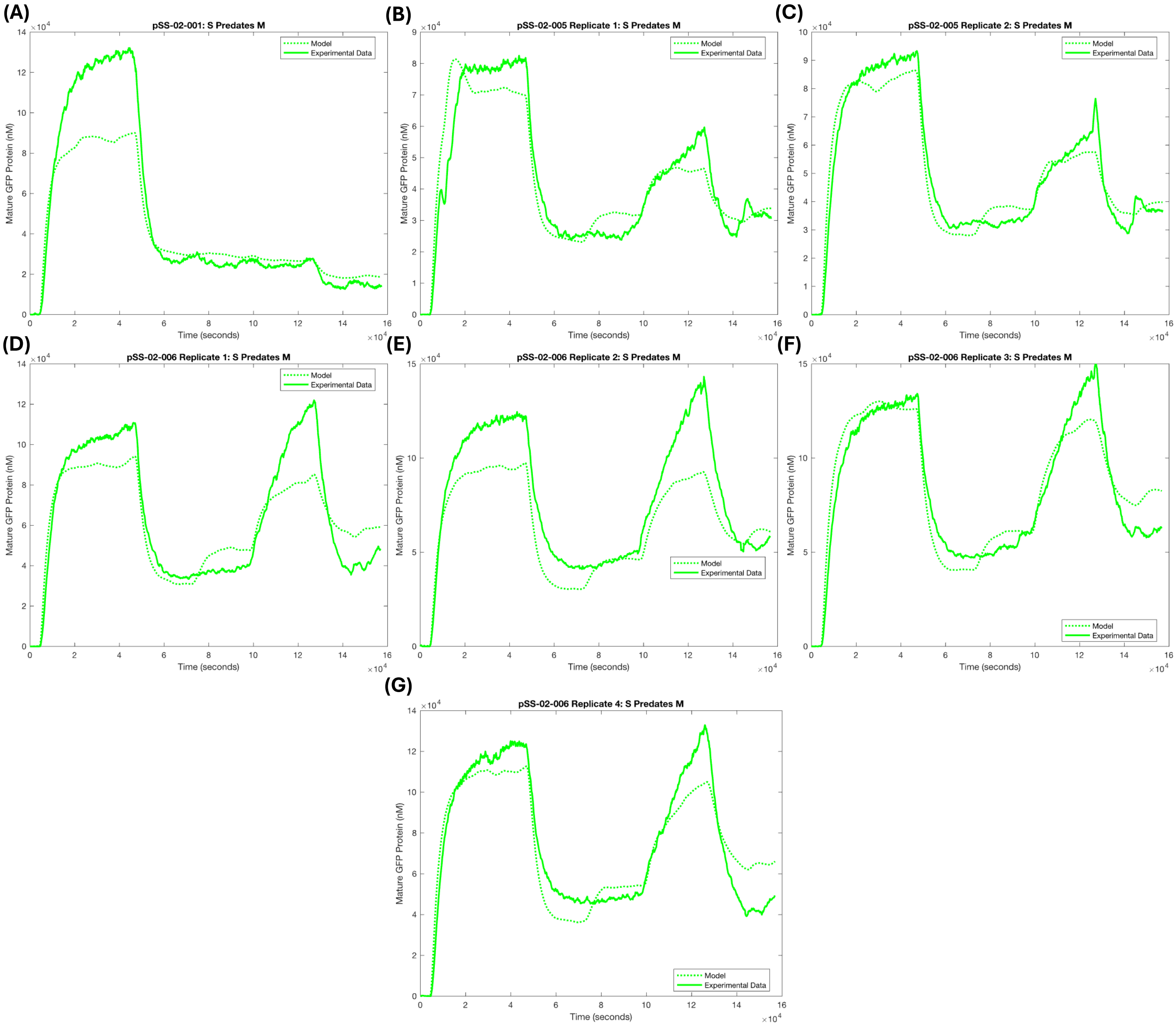


**Figure S40: Model fits for a mass-action model in which Syn-ChiX-20D predates GFPmut3 mRNA: (A)** the sRNA circuit (pSS-02-001); **(B–C)** the sponge RNA circuit without mScarlet-I (pSS-02-005); and **(D–G)** the sponge RNA circuit with mScarlet-I (pSS-02-006). All sponge-circuit models predict a GFPmut3 increase after low spRNA induction at around $7.5 \times 10^4$ seconds, which was not observed experimentally.

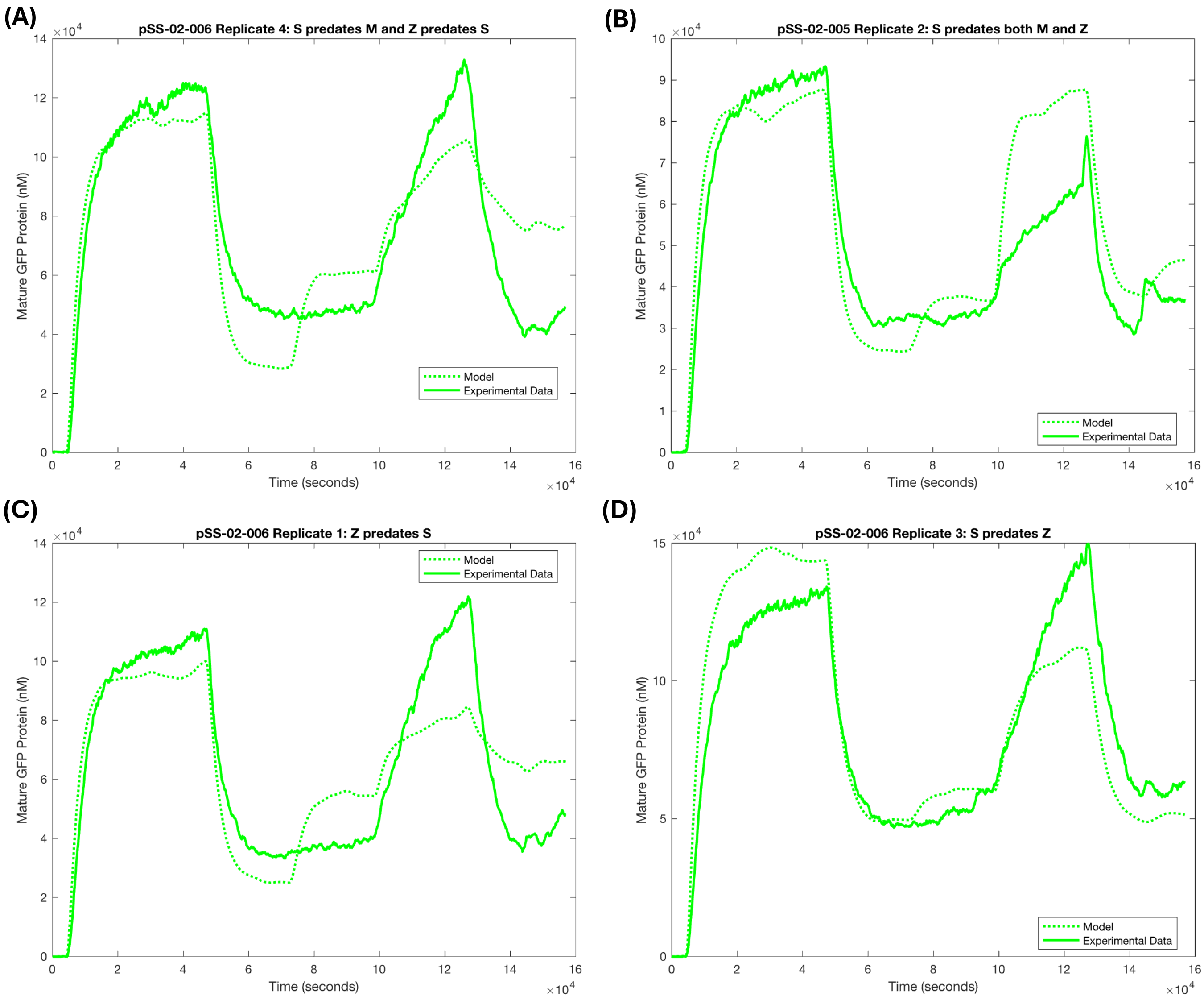


**Figure S41: Representative examples of mass-action fits with various asymmetrical RNA regulatory interactions: (A)** example fit for pSS-02-006 where Syn-ChiX-20D predates GFPmut3 mRNA and Syn-ChbBC-20D predates Syn-ChiX-20D; **(B)** example fit for pSS-02-005 where Syn-ChiX-20D predates both GFPmut3 mRNA and Syn-ChbBC-20D; **(C)** example fit for pSS-02-006 where Syn-ChbBC-20D predates Syn-ChiX-20D; **(D)** example fit for pSS-02-006 where Syn-ChiX-20D predates Syn-ChbBC-20D. All other RNA interactions are represented as annihilation reactions. None of these models qualitatively captured RNA-circuit regulatory dynamics.

#### S7.5 Incorporating Resource Competition

To account for possible coupling through a shared limiting resource, we extended the predation-based RNA model to include dependence of sRNA–mRNA and sRNA–spRNA interactions on a common resource pool. We interpret this resource, $R$, as the effective free concentration of Hfq available to the circuit components. Rather than modelling Hfq binding mechanistically, we used a coarse-grained rapid-equilibrium approximation in which the total resource $R_{\text{total}}$ is conserved and depleted by formation of interacting sRNA–mRNA and sRNA–spRNA pairs based on a previously published resource-aware modelling framework [33]. Under this approximation, the free resource is given by

$$R(t) = \frac{R_{\text{total}}}{1 + \dfrac{S(t)M(t)}{K_{sm}} + \dfrac{S(t)Z(t)}{K_{sz}}}, \tag{S6}$$

where $S(t)$, $M(t)$, and $Z(t)$ denote the concentrations of sRNA, mRNA, and spRNA, respectively, and $K_{sm}$ and $K_{sz}$ are effective dissociation constants describing sequestration of the shared resource by sRNA–mRNA and sRNA–spRNA interacting pairs.

For modelling pSS-02-001, which does not contain the sponge RNA species, the corresponding expression reduces to

$$R(t) = \frac{R_{\text{total}}}{1 + \dfrac{S(t)M(t)}{K_{sm}}}. \tag{S7}$$

Resource dependence was incorporated by scaling the sequestration fluxes by the free resource $R(t)$, giving

$$v_{sm}(t) = \frac{\gamma_{sm}}{K_{sm}} R(t)\, S(t)\, M(t), \tag{S8}$$

for sRNA–mRNA sequestration, and

$$v_{sz}(t) = \frac{\gamma_{sz}}{K_{sz}} R(t)\, S(t)\, Z(t), \tag{S9}$$

for sRNA–spRNA sequestration, where $\gamma_{sm}$ and $\gamma_{sz}$ are effective rate constants.

Because sRNA predation of mRNA gave the lowest cost among the predation variants, these fluxes were substituted into that model to generate the resource-aware ODE system (Supplementary Equations (S10)).

$$\begin{aligned}
\frac{dM}{dt} &= f(V(t)) - (\mu(t) + \delta_m)\, M - \frac{\gamma_{sm}}{K_{sm}} SMR \\
\frac{dP}{dt} &= \theta M - \mu(t)P - \tau P \\
\frac{dF}{dt} &= \tau P - \mu(t)F \\
\frac{dS}{dt} &= f(O(t)) - (\mu(t) + \delta_s)\, S - n\frac{\gamma_{sm}}{K_{sm}} SMR - \frac{\gamma_{sz}}{K_{sz}} SZR \\
\frac{dZ}{dt} &= f(I(t)) - (\mu(t) + \delta_z)\, Z - \frac{\gamma_{sz}}{K_{sz}} SZR
\end{aligned} \tag{S10}$$

Parameters were then estimated as before, with promoter and translation parameters held constant (Supplementary Table S5). RNA-circuit parameters were either fitted or fixed, as reported in detail in Supplementary Table S7. This resource-aware model substantially improved fits to GFPmut3 data for pSS-02-005 and pSS-02-006 (Supplementary Fig. S42), supporting resource competition as a plausible explanation for the observed non-linearity. This improvement came at the expense of lower-quality fits for the sRNA circuit (pSS-02-001; Supplementary Fig. S42A), which may be because resource competition is unnecessary in a circuit with a single regulatory RNA interaction pair, or because of issues with the pSS-02-001 dataset (Supplementary Note S7.7).

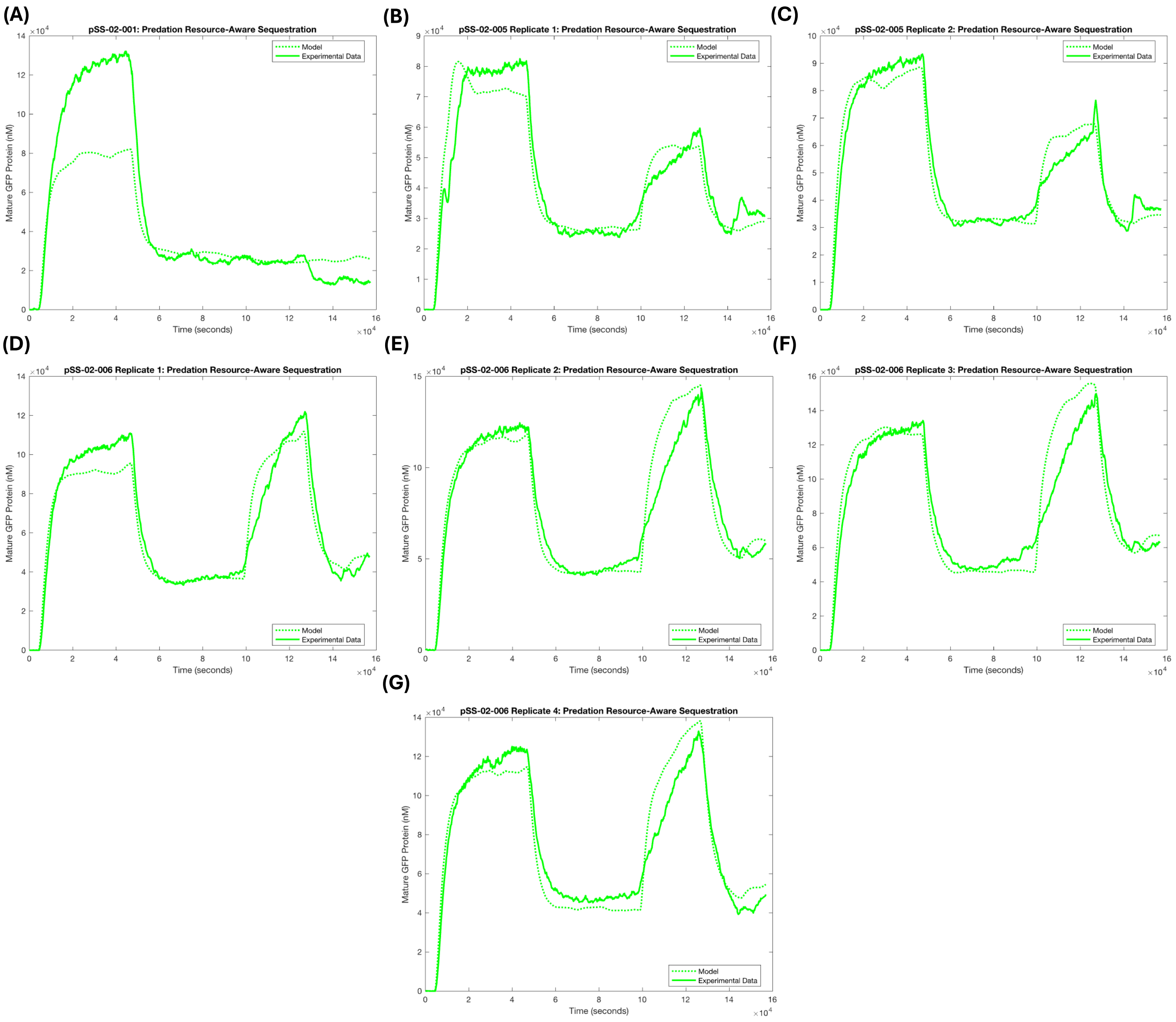


**Figure S42: GFPmut3 model fits for RNA circuits with resource-aware RNA sequestration: (A)** the sRNA circuit (pSS-02-001); **(B–C)** the spRNA circuit without mScarlet-I (pSS-02-005); and **(D–G)** the spRNA circuit with mScarlet-I (pSS-02-006). RNA interactions are modelled as resource-aware sequestration reactions, with Syn-ChiX-20D predating GFPmut3 mRNA. Unlike previous fits, this model qualitatively reproduced the experimental data, supporting the inclusion of resource competition.

We next extended the resource-aware model to include translation of mScarlet-I directly from $Z$ for pSS-02-006 (Supplementary Equations (S11)),

$$\begin{aligned}
\frac{dM}{dt} &= f(V(t)) - (\mu(t) + \delta_m)\, M - \frac{\gamma_{sm}}{K_{sm}} SMR \\
\frac{dP}{dt} &= \theta M - \mu(t)P - \tau P \\
\frac{dF}{dt} &= \tau P - \mu(t)F \\
\frac{dS}{dt} &= f(O(t)) - (\mu(t) + \delta_s)\, S - n\frac{\gamma_{sm}}{K_{sm}} SMR - \frac{\gamma_{sz}}{K_{sz}} SZR \\
\frac{dZ}{dt} &= f(I(t)) - (\mu(t) + \delta_z)\, Z - \frac{\gamma_{sz}}{K_{sz}} SZR \\
\frac{dP_2}{dt} &= \theta_m Z - \mu P_2 - \tau_m P_2 \\
\frac{dF_2}{dt} &= \tau_m P_2 - \mu F_2\,,
\end{aligned} \tag{S11}$$

where newly introduced species $P_2$ and $F_2$ represent immature and mature mScarlet-I, respectively, and newly introduced parameters $\theta_m$ and $\tau_m$ describe the translation rate of $Z$ into mScarlet-I and the maturation rate of mScarlet-I, respectively. All previous RNA-circuit parameters were re-estimated for this model in the same way as before, using fixed promoter and translation parameters and fixed values for $\delta_m$, $\tau$, $R_{\mathrm{total}}$, and $\delta_s$. The new mScarlet-I translation parameter, $\theta_m$, was fitted simultaneously. The mScarlet-I maturation rate, $\tau_m$, was fixed based on a reported maturation time of 25.7 minutes [34], giving $\tau_{\mathrm{mScarlet-I}} = \ln 2/(25.7 \times 60) = 0.0004495\ \mathrm{s}^{-1}$.

After parameter estimation, this model produced observably worse fits to the GFPmut3 traces (Supplementary Fig. S43A–G) and failed to capture the observed mScarlet-I traces (Supplementary Fig. S43H–K). Therefore, incorporating resource competition on its own was insufficient to explain the full observed circuit behaviour.

**Table S7: Fitted and chosen parameters for RNA regulatory circuit models in which Syn-ChiX-20D predates GFPmut3 mRNA with resource coupling between RNA interactions**

| Parameter | Description | Value | Units | Source |
|---|---|---|---|---|
| $\delta_{z,6}$ | Degradation rate of Syn-ChbBC-20D in pSS-02-006 | 0.03356 | $\mathrm{s}^{-1}$ | Fitted in this work |
| $\delta_{z,5}$ | Degradation rate of Syn-ChbBC-20D in pSS-02-005 | 0.1020 | $\mathrm{s}^{-1}$ | Fitted in this work |
| $\gamma_{sz,6}$ | Resource-coupled interaction rate of Syn-ChiX-20D and Syn-ChbBC-20D in pSS-02-006 | 0.627 | $\mathrm{s}^{-1}$ | Fitted in this work |
| $\gamma_{sz,5}$ | Resource-coupled interaction rate of Syn-ChiX-20D and Syn-ChbBC-20D in pSS-02-005 | 0.639 | $\mathrm{s}^{-1}$ | Fitted in this work |
| $\gamma_{sm}$ | Resource-coupled interaction rate of Syn-ChiX-20D and GFPmut3 mRNA | $3.300 \times 10^{-4}$ | $\mathrm{s}^{-1}$ | Fitted in this work |
| $SF_{6,1}$ | Multiplicative scaling factor by which GFPmut3 in pSS-02-006 replicate 1 was scaled | 1.766 | – | Fitted in this work |
| $SF_{6,2}$ | Multiplicative scaling factor by which GFPmut3 in pSS-02-006 replicate 2 was scaled | 1.929 | – | Fitted in this work |
| $SF_{6,3}$ | Multiplicative scaling factor by which GFPmut3 in pSS-02-006 replicate 3 was scaled | 2.169 | – | Fitted in this work |
| $SF_{6,4}$ | Multiplicative scaling factor by which GFPmut3 in pSS-02-006 replicate 4 was scaled | 2.004 | – | Fitted in this work |
| $SF_{5,1}$ | Multiplicative scaling factor by which GFPmut3 in pSS-02-005 replicate 1 was scaled | 1.355 | – | Fitted in this work |
| $SF_{5,2}$ | Multiplicative scaling factor by which GFPmut3 in pSS-02-005 replicate 2 was scaled | 1.628 | – | Fitted in this work |
| $SF_s$ | Multiplicative scaling factor by which GFPmut3 in pSS-02-001 was scaled | 1.788 | – | Fitted in this work |
| $R_{\mathrm{total}}$ | Total resource concentration representing the effective cellular Hfq pool | 4000 | nM | Fixed based on [35] |
| $n$ | Predation constant describing fractional loss of Syn-ChiX-20D during predation of GFPmut3 mRNA | 0.1005 | – | Fitted in this work |
| $K_{sm}$ | Effective dissociation constant governing Syn-ChiX-20D to GFPmut3 mRNA binding and resource usage | 7152 | $\mathrm{nM}^2$ | Fitted in this work |
| $K_{sz}$ | Effective dissociation constant governing Syn-ChiX-20D to Syn-ChbBC-20D binding and resource usage | $1.573 \times 10^5$ | $\mathrm{nM}^2$ | Fitted in this work |
| $\delta_s$ | Degradation rate of Syn-ChiX-20D | 0.0003610 | $\mathrm{s}^{-1}$ | Calculated based on published half-life of ChiX in [32] |

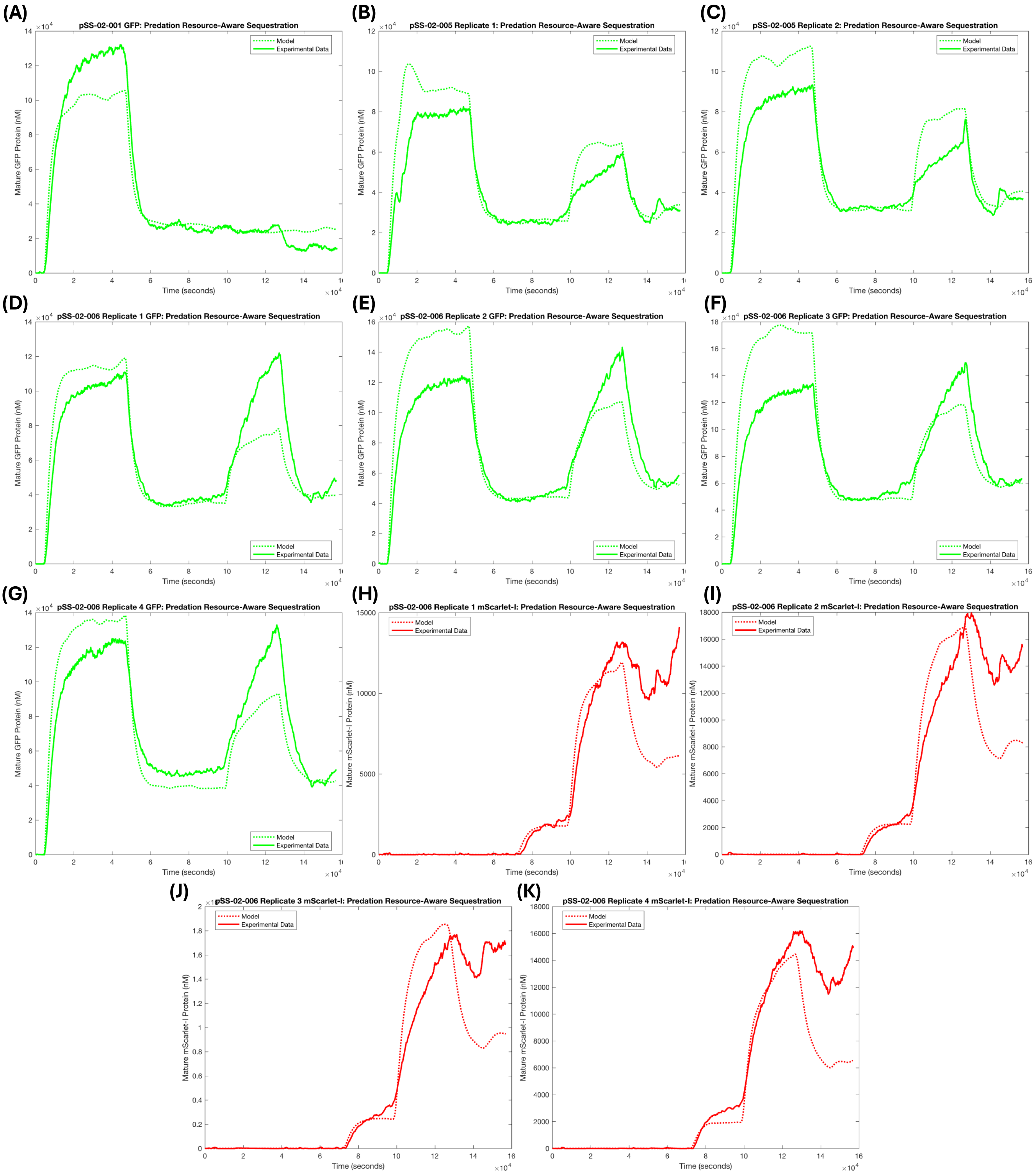


**Figure S43: Including mScarlet-I translation from the ChbBC–mScarlet-I transcript reduces the quality of the resource-aware model fit: (A)** GFPmut3 model fit for the sRNA circuit (pSS-02-001); **(B–C)** GFPmut3 model fit for the spRNA circuit without mScarlet-I (pSS-02-005); **(D–G)** GFPmut3 model fit for the spRNA circuit with mScarlet-I (pSS-02-006); and **(H–K)** mScarlet-I model fit for pSS-02-006.

#### S7.6 Final model – explicit complexes and $Z$–$S$ complex translation

In the previous models, RNA regulatory interactions were treated effectively, without explicitly representing intermediate complexes. However, the inability of these models to capture the observed mScarlet-I dynamics suggested that the spRNA–sRNA complex may be sufficiently long-lived to persist as a distinct species. Because the designed sRNA–spRNA binding region lies upstream of the mScarlet-I RBS in pSS-02-006, it is also plausible that the spRNA remains translatable while in complex. We therefore extended the model to include explicit sRNA–mRNA and sRNA–spRNA complexes, and allowed translation of mScarlet-I from both free spRNA and the spRNA–sRNA complex, yielding our final model.

This model is described by the following chemical reaction network:

$$
\begin{aligned}
&\varnothing \xrightarrow{\mathrm{f(V(t))}} \mathrm{M} && \mathrm{M} \xrightarrow{\delta_m+\mu(\mathrm{t})} \varnothing \\
&\varnothing \xrightarrow{\mathrm{f(O(t))}} \mathrm{S} && \mathrm{S} \xrightarrow{\delta_s+\mu(\mathrm{t})} \varnothing \\
&\varnothing \xrightarrow{\mathrm{f(I(t))}} \mathrm{Z} && \mathrm{Z} \xrightarrow{\delta_z+\mu(\mathrm{t})} \varnothing \\
&\mathrm{M} \xrightarrow{\theta} \mathrm{M}+\mathrm{P} && \mathrm{P} \xrightarrow{\mu(\mathrm{t})} \varnothing \\
&\mathrm{P} \xrightarrow{\tau} \mathrm{F} && \mathrm{F} \xrightarrow{\mu(\mathrm{t})} \varnothing \\
&\mathrm{S}+\mathrm{M} \xrightarrow{\frac{\gamma_{\mathrm{sm}}}{\mathrm{K_{sm}}}\mathrm{R(t)}} \mathrm{C_1} && \mathrm{C_1} \xrightarrow{\mu(\mathrm{t})} \varnothing \\
&\mathrm{C_1} \xrightarrow{\delta_c} n\,\mathrm{S} && \mathrm{S}+\mathrm{Z} \xrightarrow{\frac{\gamma_{\mathrm{sz}}}{\mathrm{K_{sz}}}\mathrm{R(t)}} \mathrm{C_2} \\
&\mathrm{C_2} \xrightarrow{\delta_c+\mu(\mathrm{t})} \varnothing && \mathrm{Z} \xrightarrow{\theta_m} \mathrm{Z}+\mathrm{P_2} \\
&\mathrm{C_2} \xrightarrow{\theta_m} \mathrm{C_2}+\mathrm{P_2} && \mathrm{P_2} \xrightarrow{\mu(\mathrm{t})} \varnothing \\
&\mathrm{P_2} \xrightarrow{\tau_m} \mathrm{F_2} && \mathrm{F_2} \xrightarrow{\mu(\mathrm{t})} \varnothing\,,
\end{aligned}
\tag{S12}
$$

where $M$ is GFPmut3 mRNA, $S$ is sRNA, $Z$ is spRNA, $C_1$ is the sRNA–mRNA complex, $C_2$ is the sRNA–spRNA complex, $P$ and $F$ are immature and mature GFPmut3, respectively, and $P_2$ and $F_2$ are immature and mature mScarlet-I, respectively.

For pSS-02-006, this reaction network gives rise to the ODE system,

$$
\begin{aligned}
\frac{dM}{dt} &= f(V(t)) - (\mu(t)+\delta_m)\,M - \frac{\gamma_{sm}}{K_{sm}}SMR \\
\frac{dP}{dt} &= \theta M - \mu(t)P - \tau P \\
\frac{dF}{dt} &= \tau P - \mu(t)F \\
\frac{dS}{dt} &= f(O(t)) - (\mu(t)+\delta_s)\,S - \frac{\gamma_{sm}}{K_{sm}}SMR - \frac{\gamma_{sz}}{K_{sz}}SZR + n\delta_c C_1 \\
\frac{dZ}{dt} &= f(I(t)) - (\mu(t)+\delta_z)\,Z - \frac{\gamma_{sz}}{K_{sz}}SZR \\
\frac{dC_1}{dt} &= \frac{\gamma_{sm}}{K_{sm}}SMR - (\mu(t)+\delta_c)\,C_1 \\
\frac{dC_2}{dt} &= \frac{\gamma_{sz}}{K_{sz}}SZR - (\mu(t)+\delta_c)\,C_2 \\
\frac{dP_2}{dt} &= \theta_m\,(Z+C_2) - \tau_m P_2 - \mu(t)P_2 \\
\frac{dF_2}{dt} &= \tau_m P_2 - \mu(t)F_2\,,
\end{aligned}
\tag{S13}
$$

where $f(V(t))$, $f(O(t))$, and $f(I(t))$ are the promoter response functions for the vanillic acid-, OC6-, and IPTG-inducible promoters, respectively; $R(t)$ is the free shared resource as defined in Supplementary Equation (S6); $\delta_s$ and

$\delta_z$ are sRNA and spRNA degradation rates; $\gamma_{sm}$ and $\gamma_{sz}$ are effective binding-rate constants for sRNA–mRNA and sRNA–spRNA interactions; $\delta_c$ is the complex degradation rate, assumed equal for $C_1$ and $C_2$; $\theta_m$ and $\tau_m$ are the translation and maturation rates of mScarlet-I; and $n$ is a predation/recycling parameter describing the fraction of $C_1$ degradation that recycles into $S$ ($n = 1$ corresponds to perfect recycling and $n = 0$ to perfect annihilation).

A reduced version of our final model for pSS-02-001 is provided in Supplementary Equations S14,

$$\begin{aligned}
\frac{dM}{dt} &= f(V(t)) - (\mu(t) + \delta_m)\, M - \frac{\gamma_{sm}}{K_{sm}} SMR \\
\frac{dP}{dt} &= \theta M - \mu(t)P - \tau P \\
\frac{dF}{dt} &= \tau P - \mu(t)F \\
\frac{dS}{dt} &= f(O(t)) - (\mu(t) + \delta_s)\, S - \frac{\gamma_{sm}}{K_{sm}} SMR + n\delta_c C_1 \\
\frac{dC_1}{dt} &= \frac{\gamma_{sm}}{K_{sm}} SMR - (\mu(t) + \delta_c)\, C_1 \,,
\end{aligned} \tag{S14}$$

where all parameters are as previously described, but $R$ is now defined by Supplementary Equation (S7). Similarly, a reduced version of our final model for pSS-02-005 is provided in Supplementary Equations (S15),

$$\begin{aligned}
\frac{dM}{dt} &= f(V(t)) - (\mu(t) + \delta_m)\, M - \frac{\gamma_{sm}}{K_{sm}} SMR \\
\frac{dP}{dt} &= \theta M - \mu(t)P - \tau P \\
\frac{dF}{dt} &= \tau P - \mu(t)F \\
\frac{dS}{dt} &= f(O(t)) - (\mu(t) + \delta_s)\, S - \frac{\gamma_{sm}}{K_{sm}} SMR - \frac{\gamma_{sz}}{K_{sz}} SZR + n\delta_c C_1 \\
\frac{dZ}{dt} &= f(I(t)) - (\mu(t) + \delta_z)\, Z - \frac{\gamma_{sz}}{K_{sz}} SZR \\
\frac{dC_1}{dt} &= \frac{\gamma_{sm}}{K_{sm}} SMR - (\mu(t) + \delta_c)\, C_1 \\
\frac{dC_2}{dt} &= \frac{\gamma_{sz}}{K_{sz}} SZR - (\mu(t) + \delta_c)\, C_2 \,,
\end{aligned} \tag{S15}$$

where all parameters are as previously described, and $R$ is defined as in Supplementary Equation (S6).

Parameters for all three models were fitted simultaneously, with shared parameters constrained as described in the Methods section of the main text. Because this was the final model, with the RNA regulatory interactions fixed to the structure described above, we also refitted the promoter and translation parameters using both the promoter-construct and RNA-circuit data. These promoter and translation parameters were bounded within ±5% of their previously fitted values (Supplementary Table S5), and the weighted cost function was updated to include both the promoter constructs and the RNA circuits. This allowed promoter parameters to be refined while accounting for both promoter-characterisation and RNA-circuit data. The final promoter and translation parameters are provided in Supplementary Table S8, the RNA-circuit parameters are provided in Supplementary Table S9, and the dataset-specific scaling factors are provided in Supplementary Table S10.

These fitted parameters produced good fits across the individual replicates (Supplementary Fig. S44). The combined fit retained quantitatively good agreement with GFPmut3 data for the promoter constructs (mean example in Fig. 4B; individual replicates in Supplementary Fig. S44A–C), comparable to the previous promoter-only fits (Supplementary Fig. S38). GFPmut3 fits for both sponge circuits, with mScarlet-I (pSS-02-006; Fig. 4D; Supplementary Fig. S44G–J) and without mScarlet-I (pSS-02-005; Fig. 4H; Supplementary Fig. S44E–F), remained qualitatively comparable to the best previous GFPmut3 fits for the sponge circuits (Supplementary Fig. S42). In addition, the final model captured the mScarlet-I dynamics more accurately than previous attempts to include mScarlet-I in the model (Fig. 4F; Supplementary Figs. S43H–K and S44K–N). Overall, these fit improvements support the hypothesis that both resource competition and translation from sRNA–spRNA complexes may contribute to the observed circuit behaviour. Despite these improvements, the model still did not quantitatively capture the dynamics of the sRNA circuit (pSS-02-001; Supplementary Fig. S44D).

**Table S8: Fitted and chosen promoter, translation, degradation, and maturation parameters used in the final RNA regulatory circuit model**

| Parameter | Description | Value | Units | Source |
|---|---|---|---|---|
| $k_{0,\mathrm{Van}}$ | Basal transcription rate of the $P_{\mathrm{VanCC}}$ promoter | 0.004803 | $\mathrm{nM\,s^{-1}}$ | Fitted in this work |
| $k_{1,\mathrm{Van}}$ | Maximal transcription rate of the $P_{\mathrm{VanCC}}$ promoter | 8.167 | $\mathrm{nM\,s^{-1}}$ | Fitted in this work |
| $K_{\mathrm{Van}}$ | Half-maximal activation constant for the $P_{\mathrm{VanCC}}$ promoter | 8565 | nM | Fitted in this work |
| $n_{\mathrm{Van}}$ | Hill coefficient for the $P_{\mathrm{VanCC}}$ promoter | 2.815 | – | Fitted in this work |
| $k_{0,\mathrm{Lux}}$ | Basal transcription rate of the $P_{\mathrm{LuxB}}$ promoter | 0.05546 | $\mathrm{nM\,s^{-1}}$ | Fitted in this work |
| $k_{1,\mathrm{Lux}}$ | Maximal transcription rate of the $P_{\mathrm{LuxB}}$ promoter | 5.622 | $\mathrm{nM\,s^{-1}}$ | Fitted in this work |
| $K_{\mathrm{Lux}}$ | Half-maximal activation constant for the $P_{\mathrm{LuxB}}$ promoter | 18.85 | nM | Fitted in this work |
| $n_{\mathrm{Lux}}$ | Hill coefficient for the $P_{\mathrm{LuxB}}$ promoter | 1.768 | – | Fitted in this work |
| $k_{0,\mathrm{Tac}}$ | Basal transcription rate of the $P_{\mathrm{Tac}}$ promoter | 0.004710 | $\mathrm{nM\,s^{-1}}$ | Fitted in this work |
| $k_{1,\mathrm{Tac}}$ | Maximal transcription rate of the $P_{\mathrm{Tac}}$ promoter | 5.820 | $\mathrm{nM\,s^{-1}}$ | Fitted in this work |
| $K_{\mathrm{Tac}}$ | Half-maximal activation constant for the $P_{\mathrm{Tac}}$ promoter | 80530 | nM | Fitted in this work |
| $n_{\mathrm{Tac}}$ | Hill coefficient for the $P_{\mathrm{Tac}}$ promoter | 2.507 | – | Fitted in this work |
| $\theta$ | Translation rate of GFPmut3 | 0.02855 | $\mathrm{s^{-1}}$ | Fitted in this work |
| $\delta_m$ | Degradation rate of GFPmut3 mRNA | 0.001699 | $\mathrm{s^{-1}}$ | Calculated from YFP mRNA half-life in [25] |
| $\tau$ | Maturation rate of GFPmut3 | 0.002818 | $\mathrm{s^{-1}}$ | Calculated from maturation time in [22] |
| $\tau_m$ | Maturation rate of mScarlet-I | 0.0004495 | $\mathrm{s^{-1}}$ | Calculated from maturation time in [34] |

**Table S9: Fitted and chosen RNA-interaction and resource-coupling parameters for the final RNA regulatory circuit model**

| Parameter | Description | Value | Units | Source |
|---|---|---|---|---|
| $\gamma_{sm}$ | Resource-coupled interaction rate of Syn-ChiX-20D and GFPmut3 mRNA | $3.222 \times 10^{-4}$ | $\mathrm{s^{-1}}$ | Fitted in this work |
| $R_{\mathrm{total}}$ | Total resource concentration representing the effective cellular Hfq pool | 4000 | nM | Fixed based on reported Hfq hexamer concentration in [35] |
| $K_{sm}$ | Effective dissociation constant governing Syn-ChiX-20D–GFPmut3 mRNA binding and resource usage | 7152 | $\mathrm{nM^2}$ | Fitted in this work |
| $K_{sz}$ | Effective dissociation constant governing Syn-ChiX-20D–Syn-ChbBC-20D binding and resource usage | $1.573 \times 10^{5}$ | $\mathrm{nM^2}$ | Fitted in this work |
| $\gamma_{sz}$ | Resource-coupled interaction rate of Syn-ChiX-20D and Syn-ChbBC-20D in pSS-02-006 and pSS-02-005 | 0.6237 | $\mathrm{s^{-1}}$ | Fitted in this work |
| $\delta_{z,6}$ | Degradation rate of Syn-ChbBC-20D in pSS-02-006 | 0.03297 | $\mathrm{s^{-1}}$ | Fitted in this work |
| $\delta_{z,5}$ | Degradation rate of Syn-ChbBC-20D in pSS-02-005 | 0.1014 | $\mathrm{s^{-1}}$ | Fitted in this work |
| $\theta_m$ | Translation rate of mScarlet-I from free Syn-ChbBC-20D and the Syn-ChbBC-20D–Syn-ChiX-20D complex | 0.03079 | $\mathrm{s^{-1}}$ | Fitted in this work |
| $\delta_c$ | Degradation or turnover rate of RNA complexes | 0.07477 | $\mathrm{s^{-1}}$ | Fitted in this work |
| $n$ | Predation constant describing fractional gain of Syn-ChiX-20D during degradation of the Syn-ChiX-20D–GFPmut3 mRNA complex | 0.8948 | – | Fitted in this work |
| $\delta_s$ | Degradation rate of Syn-ChiX-20D | 0.0003610 | $\mathrm{s^{-1}}$ | Calculated based on published half-life of ChiX in [32] |

**Table S10: Fitted dataset-specific scaling factors for the final RNA regulatory circuit model**

| Parameter | Description | Value | Units | Source |
|---|---|---|---|---|
| $SF_{6,1}$ | Multiplicative scaling factor by which mature fluorescent protein concentrations in pSS-02-006 replicate 1 were scaled | 1.702 | – | Fitted in this work |
| $SF_{6,2}$ | Multiplicative scaling factor by which mature fluorescent protein concentrations in pSS-02-006 replicate 2 were scaled | 1.836 | – | Fitted in this work |
| $SF_{6,3}$ | Multiplicative scaling factor by which mature fluorescent protein concentrations in pSS-02-006 replicate 3 were scaled | 2.254 | – | Fitted in this work |
| $SF_{6,4}$ | Multiplicative scaling factor by which mature fluorescent protein concentrations in pSS-02-006 replicate 4 were scaled | 2.024 | – | Fitted in this work |
| $SF_{5,1}$ | Multiplicative scaling factor by which GFPmut3 in pSS-02-005 replicate 1 was scaled | 1.371 | – | Fitted in this work |
| $SF_{5,2}$ | Multiplicative scaling factor by which GFPmut3 in pSS-02-005 replicate 2 was scaled | 1.606 | – | Fitted in this work |
| $SF_s$ | Multiplicative scaling factor by which GFPmut3 in pSS-02-001 was scaled | 1.846 | – | Fitted in this work |

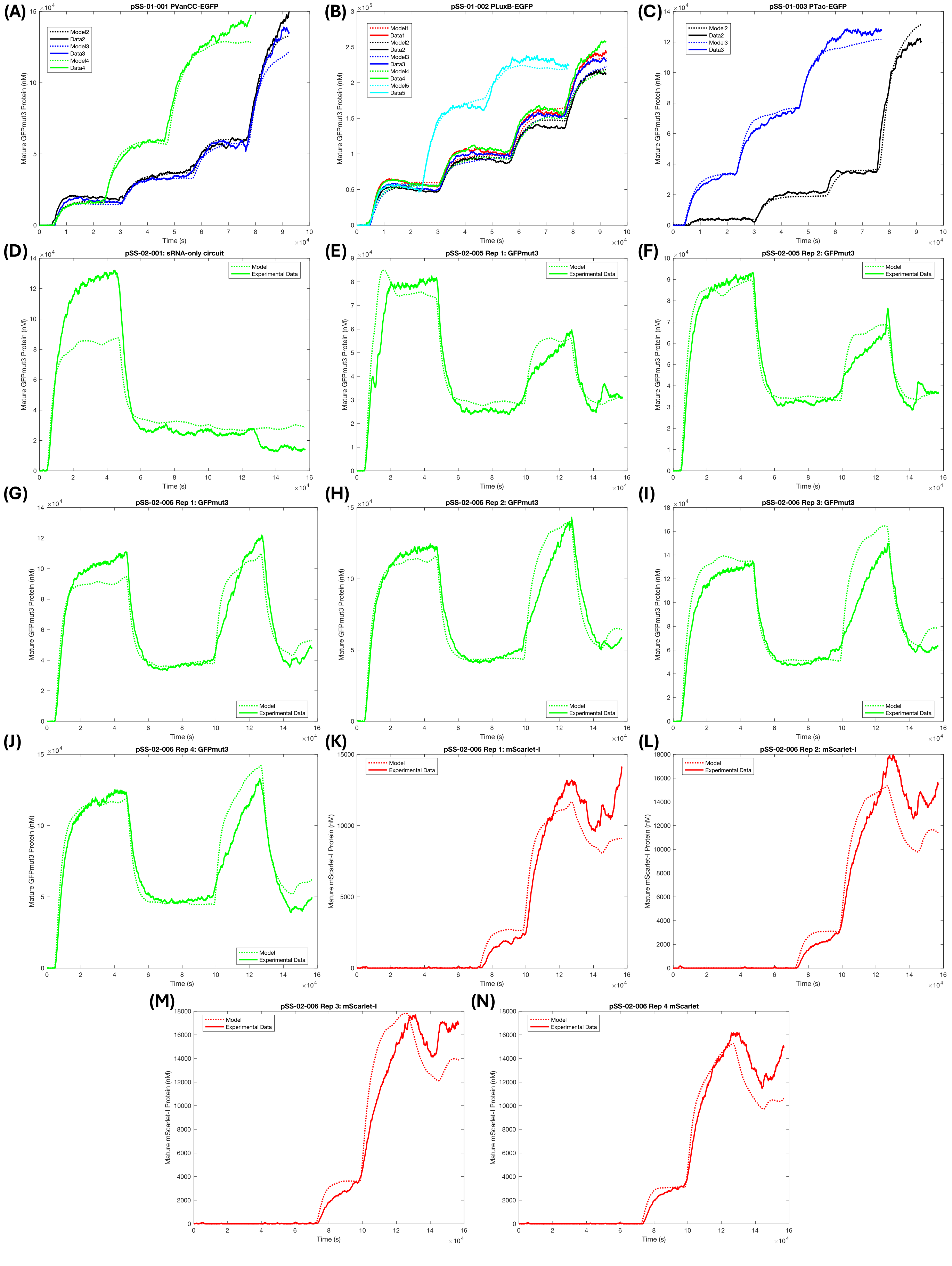


**Figure S44: Final RNA-circuit model fits for individual replicates:** **(A–C)** the standardised promoter constructs (pSS-01-001/$P_{VanCC}$, pSS-01-002/$P_{LuxB}$, and pSS-01-003/$P_{Tac}$); **(D)** the sRNA circuit (pSS-02-001); **(E–F)** the spRNA circuit without mScarlet-I (pSS-02-005); **(G–J)** GFPmut3 in the spRNA circuit with mScarlet-I (pSS-02-006); and **(K–N)** mScarlet-I in pSS-02-006.

#### S7.7 Validation and Investigation of Model Issues

##### S7.7.1 Issues with sRNA Circuit Model

Although the final model quantitatively captured promoter-construct and sponge-circuit behaviour, it did not qualitatively capture the behaviour of the fitted pSS-02-001 dataset well (Supplementary Fig. S44D). Our circuit design and Chi.Bio characterisation were optimised for repeatability, but occasional issues can arise in the data, including biofilm formation, hardware failures, and biochemical issues. In this instance, there was only a single experiment ($n = 1$) for pSS-02-001, and an unusually high growth rate was observed (Fig. 4K). This reduced its comparability with the other datasets and suggests that the fitted pSS-02-001 dataset may not be fully representative.

##### S7.7.2 Validating Model Against 'Unseen' Data

To investigate this further and guard against overfitting, we next tested the final model against data that were not used in the fitting process for both the sponge and sRNA circuits. We first validated the pSS-02-006 model and its fitted parameters against additional pSS-02-006 characterisation data that were not used in the initial model fitting (Supplementary Fig. S21). To achieve this, model structure and all fitted parameters were held constant. Because these additional datasets were not quantified in absolute units, only two additional parameters, $m$ and $c$, were fitted for each experiment to convert model output $y$ in nM to fluorescence in arbitrary units according to,

$$x = \frac{y - c}{m}.$$

After fitting these conversion parameters, the model qualitatively recapitulated experimentally observed behaviour in the four unseen replicates for pSS-02-006 (Supplementary Fig. S45A–D), despite substantially different growth rates compared with the fitted data (Supplementary Fig. S45E). This suggests that the model captures sponge RNA-mediated regulation across differing physiological conditions.

We next carried out a similar test on 'unseen' data for pSS-02-001. We compared our final model (Supplementary Equations (S14)) and parameters (Supplementary Tables S8 and S9) against simple mass-action models with annihilation (Supplementary Equations (S16)) or predation (Supplementary Equations (S17)) fitted to the quantified pSS-02-001 data alone. For these new fits, we fixed the promoter and translation parameters to those used in our final model (Supplementary Table S8) and fixed $\delta_s = 0.0003610$ as in all models, fitting only the sRNA–mRNA binding parameter and a scaling factor. These newly fitted simple mass-action models fit the quantified pSS-02-001 data well (Supplementary Fig. S46A–B). However, when they were tested against the 'unseen' pSS-02-001 data, our final model fit that data better (Supplementary Fig. S46E) than either the annihilation or predation model (Supplementary Fig. S46C–D). This further supports the importance of resource competition in understanding our synthetic ChiX circuits, even in the absence of the competing synthetic ChbBC sponge RNA, while also suggesting that the poor fit to the quantified pSS-02-001 dataset may reflect issues specific to that unrepeated experiment.

The following is the simple mass-action model with annihilation for the sRNA circuit (pSS-02-001):

$$\begin{aligned}
\frac{dM}{dt} &= f(V(t)) - (\mu(t) + \delta_m)\, M - \gamma_{sm} SM \\
\frac{dP}{dt} &= \theta M - \mu(t)P - \tau P \\
\frac{dF}{dt} &= \tau P - \mu(t)F \\
\frac{dS}{dt} &= f(O(t)) - (\mu(t) + \delta_s)\, S - \gamma_{sm} SM
\end{aligned} \tag{S16}$$

The following is the simple mass-action model with predation for the sRNA circuit (pSS-02-001):

$$\begin{aligned}
\frac{dM}{dt} &= f(V(t)) - (\mu(t) + \delta_m)\, M - \gamma_{sm} SM \\
\frac{dP}{dt} &= \theta M - \mu(t)P - \tau P \\
\frac{dF}{dt} &= \tau P - \mu(t)F \\
\frac{dS}{dt} &= f(O(t)) - (\mu(t) + \delta_s)\, S
\end{aligned} \tag{S17}$$

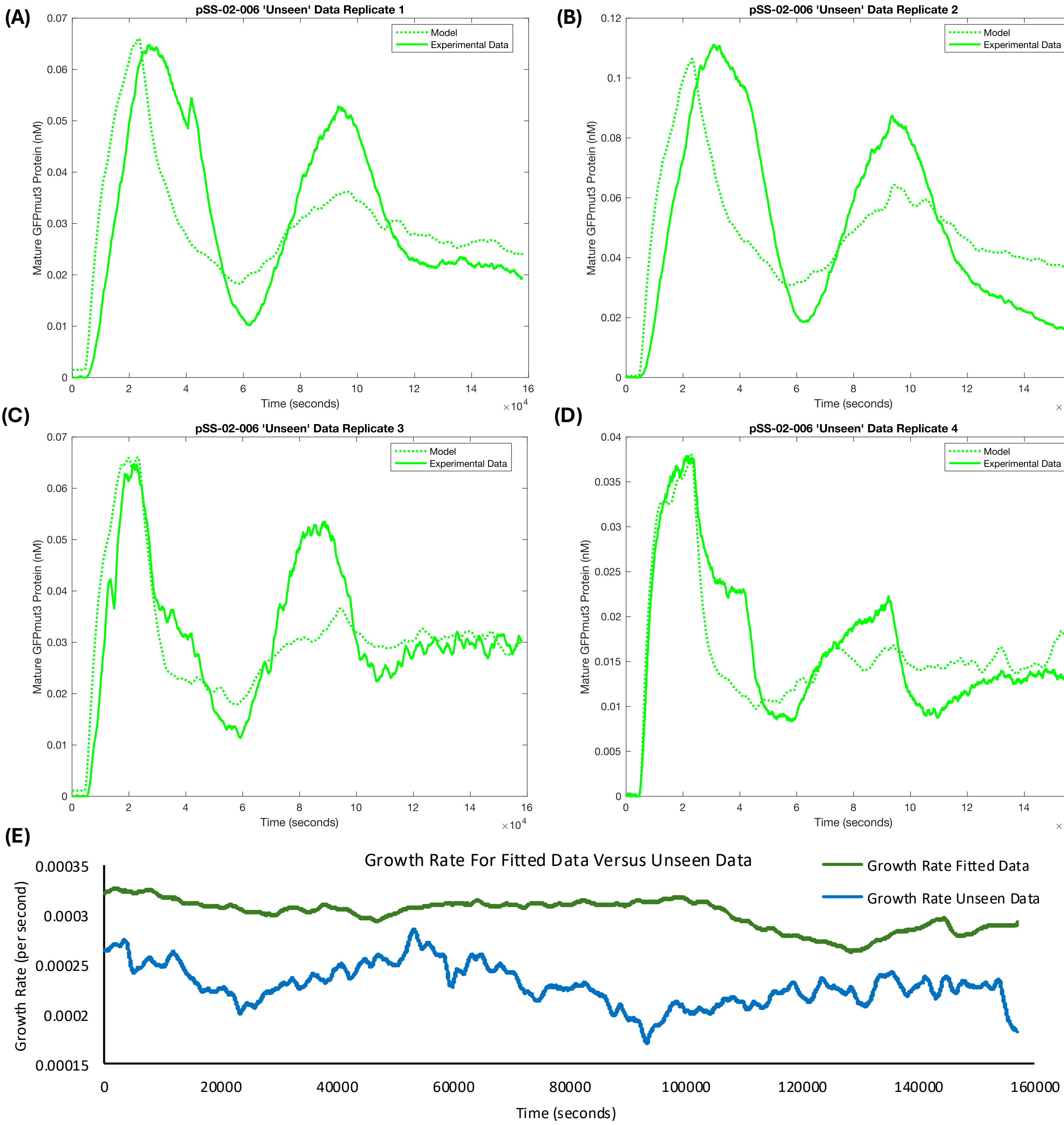


**Figure S45: Validation of our final model against 'unseen' pSS-02-006 data: (A–D)** Four replicate experiments not used in the model fitting process for pSS-02-006 (spRNA circuit with mScarlet-I). The final model was applied to these data by fitting only calibration parameters that convert model output from nM to arbitrary fluorescence units. **(E)** Growth rate was higher at all timepoints in the Chi.Bio experiments used for model fitting. The model largely qualitatively captures the behaviour of pSS-02-006 in these 'unseen' experiments despite the physiological differences indicated by the divergent growth rates.

**Table S11: Newly fitted $\gamma_{sm}$ values for the simple mass-action annihilation and predation models in Supplementary Fig. S46C–D.**

| Parameter | Annihilation | Predation | Units | Source |
|---|---|---|---|---|
| $\gamma_{sm}$ | $2.241 \times 10^{-6}$ | $1.438 \times 10^{-6}$ | $\text{nM}^{-1}\,\text{s}^{-1}$ | Fitted in this work |

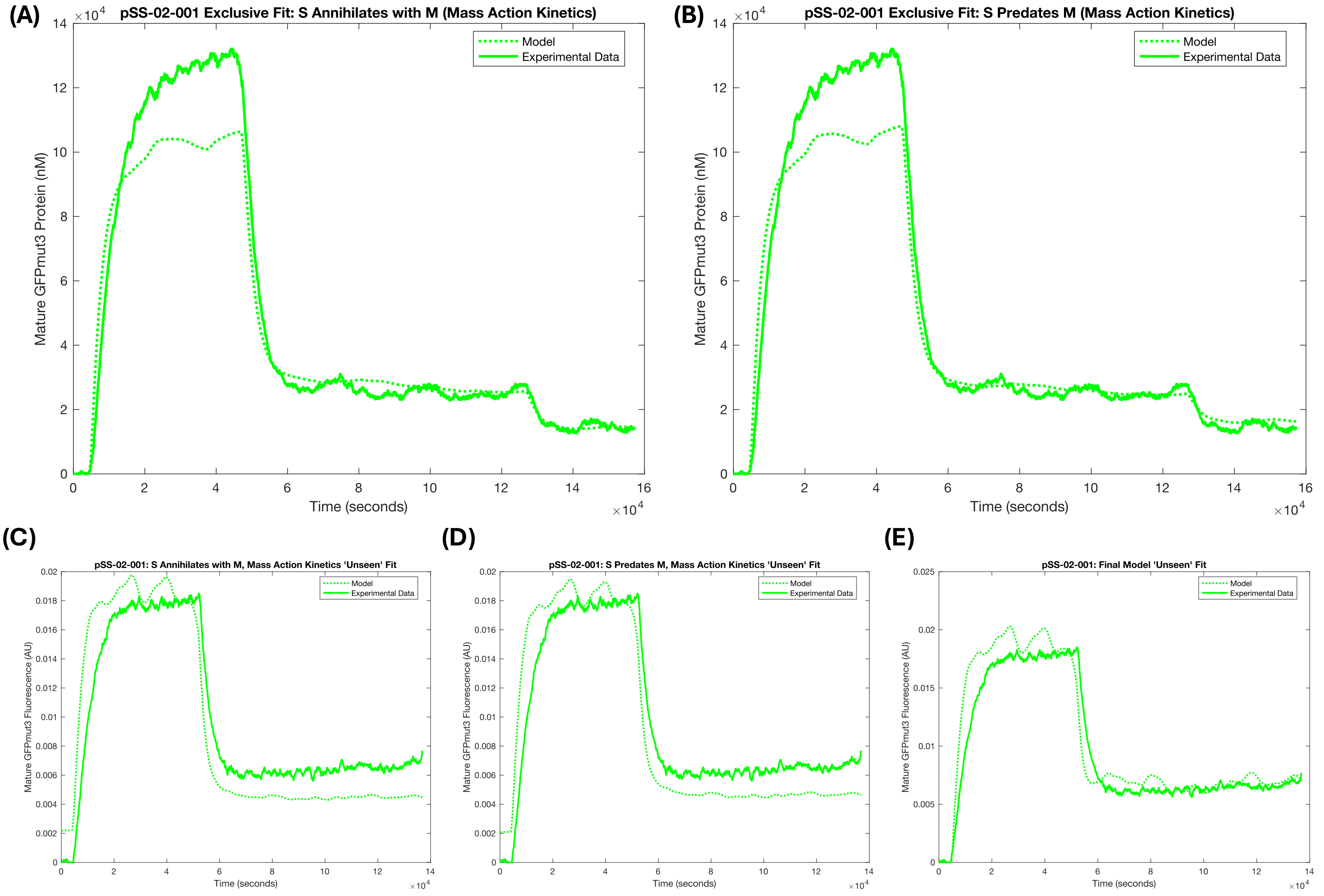


**Figure S46: Validating the final model against sRNA-circuit data: (A)** Simple mass-action annihilation model fit to the quantified pSS-02-001 data; **(B)** simple mass-action predation model fit to the quantified pSS-02-001 data; **(C–E)** the two simple mass-action models and the final model applied to 'unseen' pSS-02-001 data, fitting only conversion parameters from nM to arbitrary fluorescence units. The final model provides a better quantitative fit to these data than the simpler mass-action models.

### S8 Growth-feedback investigation

As described in the Results section of the main text, we investigated why the sponge RNA appeared to perform better in the circuit with mScarlet-I (pSS-02-006) than in the otherwise comparable circuit without mScarlet-I (pSS-02-005). We first introduced a series of stop codons at the beginning of the mScarlet-I coding sequence to generate pSS-02-009 (Supplementary Fig. S47). This produced a sponge RNA with a very similar sequence, and likely similar structure, to the sponge RNA in pSS-02-006, while removing the burden associated with mScarlet-I expression. This allowed us to distinguish between burden-mediated growth feedback and changes to spRNA structure or downstream sequence as explanations for the apparent improvement in performance. As discussed in the main text, removing mScarlet-I expression reduced the apparent spRNA regulatory effect (Fig. 4J), suggesting that at least part of the apparent improvement is due to burden-mediated growth feedback.

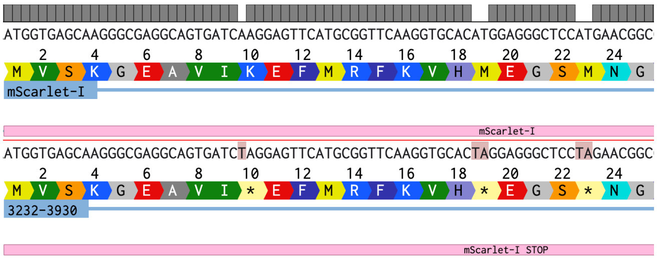


**Figure S47: Sequencing results and alignment showing stop codons introduced into mScarlet-I in pSS-02-006 by site-directed mutagenesis, generating pSS-02-009.**

As discussed in the main text, we next carried out growth-feedback modelling to further investigate this effect and assess its plausibility. We provide additional details on that modelling here.

We derived a growth-feedback variant of the final RNA-circuit model to test this hypothesis. In the main fitting framework, experimentally measured time-varying growth rates were used directly as exogenous inputs to the ODE system. Here, instead, growth rate was predicted from modelled fluorescent protein abundance and then fed back into the model as the dilution rate.

Specifically, growth was modelled phenomenologically as

$$\mu = \mu_{\max} \left( 1 - \frac{P + F}{Q_1} - \frac{P_2 + F_2}{Q_2} \right) , \tag{S18}$$

where $P$ and $F$ are the immature and mature GFPmut3 concentrations, $P_2$ and $F_2$ are the immature and mature mScarlet-I concentrations, $Q_1$ and $Q_2$ are fitted proportionality constants describing the linear growth defect associated with GFPmut3 and mScarlet-I expression, respectively, and $\mu_{\max}$ is the maximal growth rate. Because experimental conditions could vary between replicates, $\mu_{\max}$ was fitted separately for each replicate.

This predicted growth rate replaced the experimentally measured growth-rate input in the dilution terms of the final RNA-circuit model. Parameters were then estimated using a combined cost function that included GFPmut3 trajectories, mScarlet-I trajectories where present, and measured growth-rate data, as described in the Methods. Fitted parameter values are reported in Supplementary Table S12.

To assess whether growth feedback improved explanatory power, we also fitted a fixed-growth version of the model in which $\mu = \mu_{\max}$, while retaining the same fitting framework. In this comparison, different spRNA degradation rates were still allowed for pSS-02-005 and pSS-02-006, so that any improvement in fit could not be attributed solely to differences in inferred spRNA stability.

The growth-feedback model quantitatively captured the observed growth-rate changes for pSS-02-005 and pSS-02-006 (Fig. 4K; Supplementary Fig. S48A) and also fit the fluorescent protein trajectories well (Supplementary Fig. S48B–D). By contrast, the fixed-growth model did not fully capture the relative GFPmut3 changes following spRNA induction (Fig. 4L), particularly for pSS-02-006, where both GFPmut3 and mScarlet-I are expressed and growth feedback would therefore be expected to be stronger. These results support a contribution of burden-mediated growth feedback to the observed circuit behaviour.

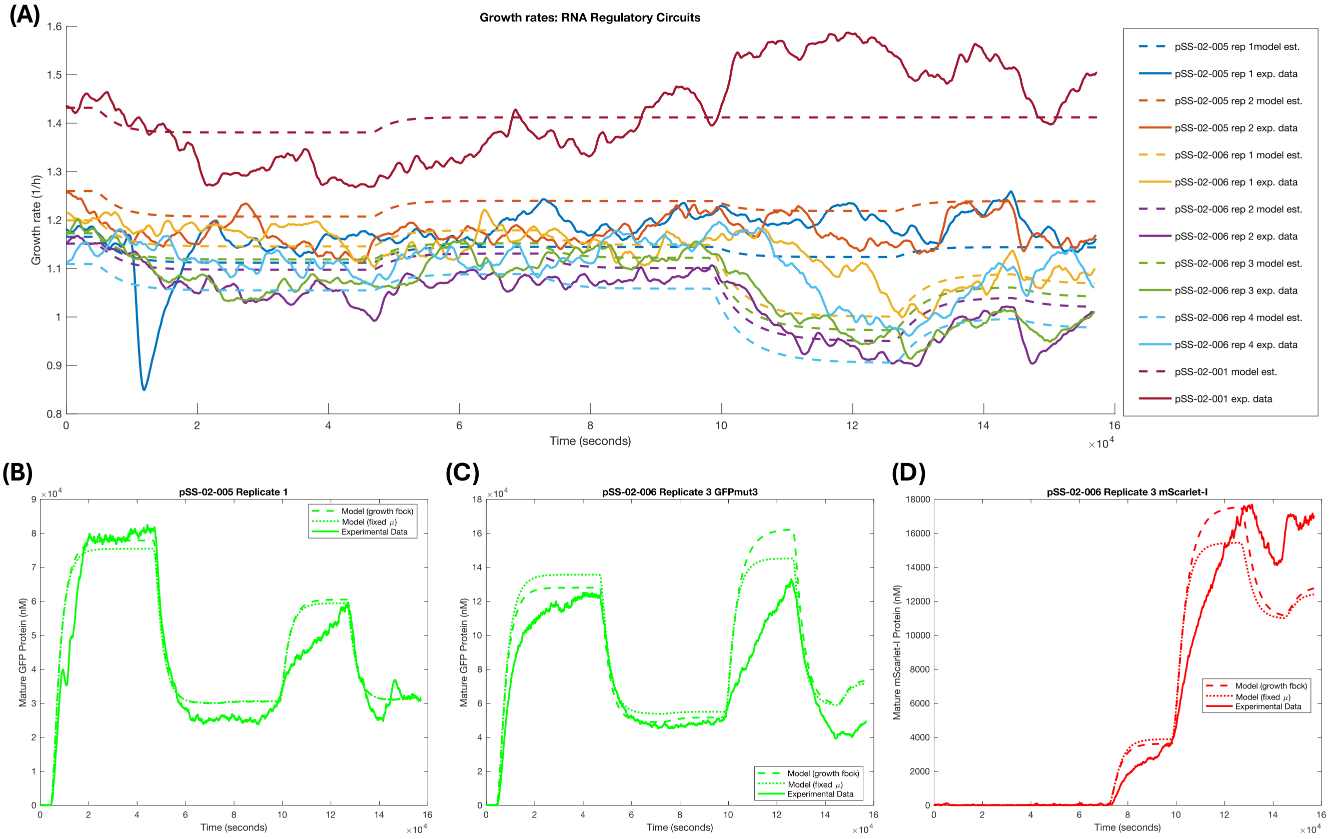


**Figure S48: Growth-feedback model fits: (A)** Fitted growth rates compared with experimental growth rates for the RNA circuits; **(B–D)** representative fits comparing the growth-feedback model, fixed-growth model, and experimental data for the two spRNA circuits (pSS-02-005 and pSS-02-006).

**Table S12: Fitted parameters for the growth-feedback model**

| Parameter | Description | Value | Units | Source |
|---|---|---|---|---|
| $J_{\mathrm{GFP}}$ | Burden threshold for GFPmut3 | 1.372 | mM | Fitted in this work |
| $J_{\mathrm{mScarlet}}$ | Burden threshold for mScarlet-I | 0.1102 | mM | Fitted in this work |
| $\mu_{\max,6,1}$ | Maximal growth rate for pSS-02-006 replicate 1 | 1.200 | $\mathrm{h}^{-1}$ | Fitted in this work |
| $\mu_{\max,6,2}$ | Maximal growth rate for pSS-02-006 replicate 2 | 1.152 | $\mathrm{h}^{-1}$ | Fitted in this work |
| $\mu_{\max,6,3}$ | Maximal growth rate for pSS-02-006 replicate 3 | 1.173 | $\mathrm{h}^{-1}$ | Fitted in this work |
| $\mu_{\max,6,4}$ | Maximal growth rate for pSS-02-006 replicate 4 | 1.109 | $\mathrm{h}^{-1}$ | Fitted in this work |
| $\mu_{\max,5,1}$ | Maximal growth rate for pSS-02-005 replicate 1 | 1.165 | $\mathrm{h}^{-1}$ | Fitted in this work |
| $\mu_{\max,5,2}$ | Maximal growth rate for pSS-02-005 replicate 2 | 1.260 | $\mathrm{h}^{-1}$ | Fitted in this work |
| $\mu_{\max,s}$ | Maximal growth rate for pSS-02-001 | 1.432 | $\mathrm{h}^{-1}$ | Fitted in this work |
| $\mu_{\max,\mathrm{Van},2}$ | Maximal growth rate for pSS-01-001 replicate 2 | 1.063 | $\mathrm{h}^{-1}$ | Fitted in this work |
| $\mu_{\max,\mathrm{Van},3}$ | Maximal growth rate for pSS-01-001 replicate 3 | 1.232 | $\mathrm{h}^{-1}$ | Fitted in this work |
| $\mu_{\max,\mathrm{Van},4}$ | Maximal growth rate for pSS-01-001 replicate 4 | 1.176 | $\mathrm{h}^{-1}$ | Fitted in this work |
| $\mu_{\max,\mathrm{Lux},1}$ | Maximal growth rate for pSS-01-002 replicate 1 | 1.115 | $\mathrm{h}^{-1}$ | Fitted in this work |
| $\mu_{\max,\mathrm{Lux},2}$ | Maximal growth rate for pSS-01-002 replicate 2 | 1.202 | $\mathrm{h}^{-1}$ | Fitted in this work |
| $\mu_{\max,\mathrm{Lux},3}$ | Maximal growth rate for pSS-01-002 replicate 3 | 1.218 | $\mathrm{h}^{-1}$ | Fitted in this work |
| $\mu_{\max,\mathrm{Lux},4}$ | Maximal growth rate for pSS-01-002 replicate 4 | 1.193 | $\mathrm{h}^{-1}$ | Fitted in this work |
| $\mu_{\max,\mathrm{Lux},5}$ | Maximal growth rate for pSS-01-002 replicate 5 | 1.211 | $\mathrm{h}^{-1}$ | Fitted in this work |
| $\mu_{\max,\mathrm{Tac},2}$ | Maximal growth rate for pSS-01-003 replicate 2 | 1.114 | $\mathrm{h}^{-1}$ | Fitted in this work |
| $\mu_{\max,\mathrm{Tac},3}$ | Maximal growth rate for pSS-01-003 replicate 3 | 1.157 | $\mathrm{h}^{-1}$ | Fitted in this work |

### S9 Additional Details for Quantitative Investigation of Synthetic spRNA Systems Biology

Unless otherwise stated, all simulations in this section and the main text were performed with our final model (Supplementary Equations (S13)) using the parameters described in Supplementary Tables S9 and S8.

#### S9.1 Promoter leakiness in sRNA gene expression

The sponge RNA-mediated leakiness tuning described in the main text (Fig. 7I) may be particularly important in sRNA biology, where promoter leakiness can have an outsized impact. To demonstrate this, we simulated steady-state gene-expression dose-response curves (Supplementary Fig. S49) with different levels of promoter leakiness applied either to a promoter driving GFPmut3 expression (Supplementary Equations (S2)) or to the same promoter driving sRNA expression (Supplementary Equations (S14)). Promoter leakiness had little effect when driving the protein itself, but applying the same leakiness to sRNA expression strongly altered protein output and the resulting inducer response functions (Supplementary Fig. S49). This may help explain why spRNAs are common in important tightly regulated cellular processes [36].

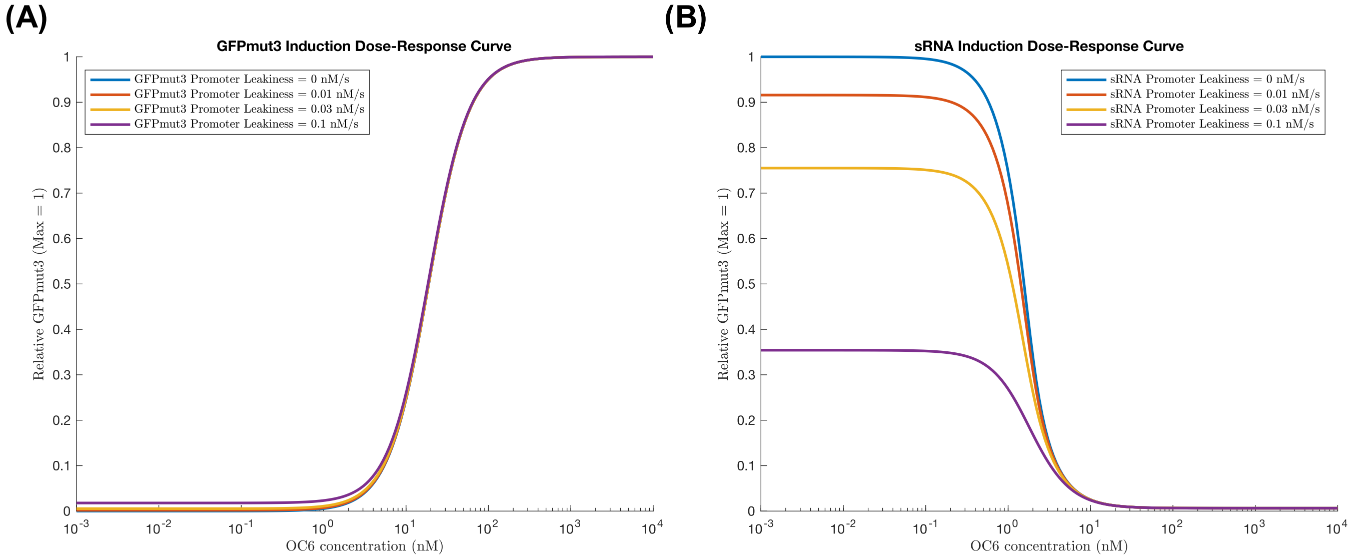


**Figure S49: Promoter leakiness can have different effects when applied directly to protein expression or indirectly through sRNAs: (A)** Fitted $P_{LuxB}$ promoter parameters were used to model the effect of leakiness on GFPmut3 expression, where increasing leakiness had little impact on the resulting GFPmut3 steady-state dose-response curve. **(B)** When $P_{LuxB}$ instead drove sRNA expression, the same changes in leakiness strongly altered the resulting GFPmut3 dose-response curve.

#### S9.2 Response-time investigation

As described in the Results section of the main text, we next investigated the impact of sRNA and spRNA on response times.

##### S9.2.1 Response times for trajectories rising from zero expression

We first investigated rise times for trajectories reaching a chosen steady-state GFPmut3 expression from zero. This could be achieved by different input combinations: switching on GFPmut3; switching on GFPmut3 and sRNA; or switching on GFPmut3, sRNA, and spRNA. For the chosen transcription rates, the GFPmut3 plus sRNA scenario gave the fastest rise times, while inclusion of spRNA increased rise time (Supplementary Fig. S50). We then tested whether this result was specific to the chosen parameters by simulating 10 alternative input combinations that produced the same steady state (Supplementary Fig. S51A), and by analysing rise times for random input combinations with steady-state normalised to one (Supplementary Fig. S51B). For a given sRNA transcription rate, inclusion of spRNA transcription always increased rise time; however, when sRNA transcription rates differed, some spRNA-containing combinations were faster than others without spRNA (Supplementary Fig. S51A). Similarly, across random input combinations, inclusion of spRNA sometimes produced faster rise times (Supplementary Fig. S51B). Nevertheless, in both analyses, mean 50% and 90% rise times were slower with non-zero spRNA transcription (Supplementary Tables S13 and S14).

(A)

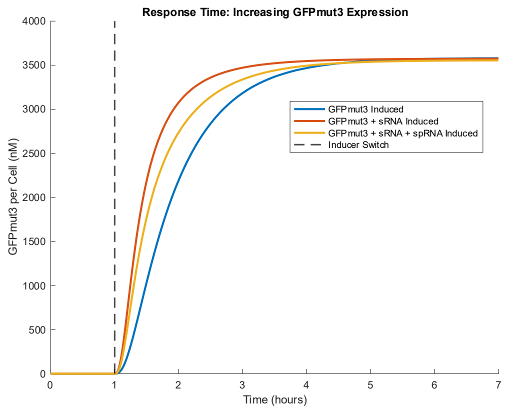


(B)

| Condition | Rise time after induction (hours) | | | |
|---|---|---|---|---|
| | T_10% | T_25% | T_50% | T_90% |
| GFPmut3 induced | 0.26532 | 0.45144 | 0.79371 | 2.0802 |
| GFPmut3 + sRNA induced | 0.1372 | 0.22694 | 0.39353 | 1.1861 |
| GFPmut3 + sRNA + spRNA induced | 0.16254 | 0.27546 | 0.50164 | 1.6148 |

**Figure S50: GFPmut3 rise-time simulations from zero expression:** **(A)** Example trajectories for three conditions: GFPmut3 transcription alone; GFPmut3 plus Syn-ChiX-20D transcription; and GFPmut3, Syn-ChiX-20D, and Syn-ChbBC-20D transcription. Transcription rates were chosen to achieve the same steady-state GFPmut3 expression. **(B)** Associated 10%, 25%, 50%, and 90% rise times. In this example, the fastest response was achieved when GFPmut3 was expressed alongside the sRNA, while the slowest response occurred when only GFPmut3 was transcribed.

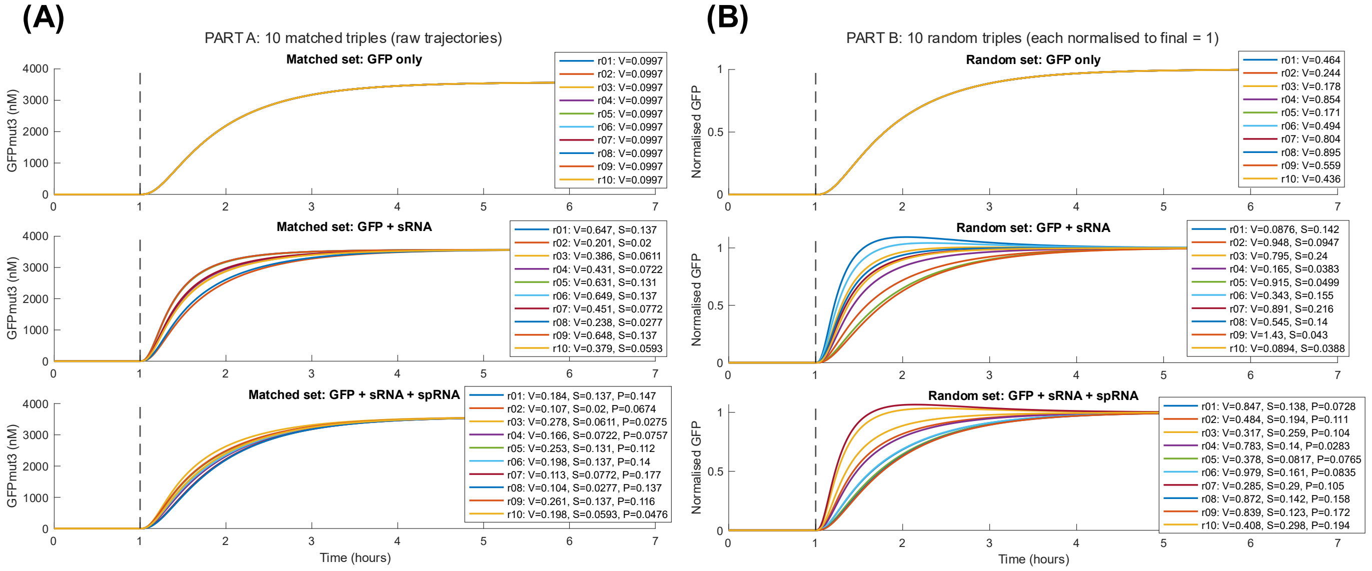


**Figure S51: Effect of transcription-input choices on response times across the three expression conditions:** **(A)** Simulations in which transcription rates were chosen to give the same steady-state GFPmut3 expression, allowing direct comparison of rise times. **(B)** Simulations in which random transcription-rate inputs were chosen and the resulting steady-state GFPmut3 expression was normalised to 1. Across both analyses, the fastest mean rise times were achieved by GFPmut3 plus sRNA expression, although some spRNA-containing parameter sets were faster than some parameter sets without spRNA.

**Table S13: Response-time summary statistics for Supplementary Fig. S51A simulations**

| Condition | Description | Mean $t_{50}$ (h) | SD $t_{50}$ (h) | Mean $t_{90}$ (h) | SD $t_{90}$ (h) |
|---|---|---|---|---|---|
| 1 | GFP only | 0.794 | 0.000 | 2.080 | 0.000 |
| 2 | GFP + sRNA | 0.441 | 0.0905 | 1.330 | 0.297 |
| 3 | GFP + sRNA + spRNA | 0.693 | 0.0779 | 1.963 | 0.105 |

##### S9.2.2 Response times in turbidostat experiments

As described in the Results section of the main text, faster rise times could be achieved when spRNA expression was used to de-repress a gene already under sRNA regulation (Fig. 7J). We next investigated whether this effect would be enhanced in a turbidostat-like setting. In a turbidostat, immediately reducing chemical inducer concentration is difficult because it requires full media replacement. Without full media replacement, inducer concentration decreases

**Table S14: Response-time summary statistics for Supplementary Fig. S51B simulations**

| Condition | Description | Mean $t_{50}$ (h) | SD $t_{50}$ (h) | Mean $t_{90}$ (h) | SD $t_{90}$ (h) |
|---|---|---|---|---|---|
| 1 | GFP only | 0.794 | 0.000 | 2.080 | 0.000 |
| 2 | GFP + sRNA | 0.431 | 0.195 | 1.169 | 0.596 |
| 3 | GFP + sRNA + spRNA | 0.526 | 0.206 | 1.494 | 0.606 |

by growth-rate-dependent dilution rather than instantaneous removal.

To model this, we repeated the step-response simulations while adding dynamic inducer-state variables to the ODE model. Inducer increases were applied instantaneously at the switching time, whereas decreases were modelled as washout by dilution, with inducer concentration decaying at the fixed growth/dilution rate $\mu = 0.0003468419\ \mathrm{s}^{-1}$ ($1.25\ \mathrm{h}^{-1}$). Simulations were run for 1 h before the switch and 6 h after the switch, and response times were calculated relative to the change in mature GFPmut3 concentration between the value at the switching time and the final simulated value. In these simulations, the inducer variables were treated as direct transcription-rate inputs rather than being mapped through promoter Hill functions. Under this washout assumption, switching on the spRNA produced a larger reduction in GFPmut3 rise time than simply switching off sRNA production, consistent with spRNAs providing a rapid post-transcriptional route for relieving existing sRNA-mediated repression (Supplementary Fig. S52). These improved rise times may be particularly important in biotechnological contexts where bioreactors are often used.

(A)

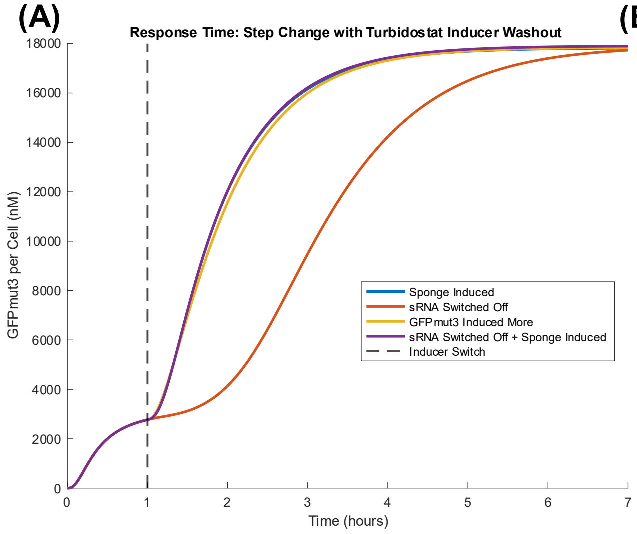


(B)

| Condition | Rise time after condition change (h) | |
|---|---|---|
| | $T_{50\%}$ | $T_{90\%}$ |
| Sponge induced | 0.78884 | 2.0757 |
| sRNA switched off | 2.1297 | 3.8332 |
| GFPmut3 induced more | 0.84163 | 2.1595 |
| Sponge induced + sRNA switched off | 0.79198 | 2.0856 |

**Figure S52: Rise times in turbidostat-like simulations: (A)** Simulated gene-expression trajectories for an sRNA-repressed gene de-repressed to the same steady state by sponge RNA transcription, delayed switch-off of sRNA transcription through inducer dilution, increased transcription of the repressed gene, or combined sRNA switch-off and sponge RNA transcription. **(B)** 50% and 90% rise times for the simulations, showing that sponge RNA transcription improves rise times in this turbidostat-like setting.

#### S9.3 Input filtering

As described in the main text, we also investigated rejection of noisy input disturbances. Here we provide more details on these simulations.

To investigate noise filtering behaviour, we first simulated one of two noisy transcriptional inputs: (i) noisy spRNA transcription with sRNA transcription held constant, or (ii) noisy sRNA transcription with spRNA transcription set to zero (Fig. 7K, bottom). For this parameter set, output noise was lower when the noisy input was applied to the spRNA (Fig. 7K, top). We next tested whether this disturbance rejection generalised across different realisations of input noise. For each of 50 iterations, we generated a single smoothed random noise trace after the perturbation time. This trace was mean-centred, scaled to unit standard deviation, and then applied identically to the two simulated conditions, either as variation in spRNA transcription or as variation in sRNA transcription. In both cases, the noisy input had a mean transcription rate of 0.1 $\mathrm{nM\,s^{-1}}$ and was scaled by an SD of 0.1 $\mathrm{nM\,s^{-1}}$. The resulting noisy inputs were not perfectly Gaussian as negative transcription rates were clipped to zero. Simulations were run for 18 h, with the noisy input applied after 6 h. To avoid the immediate post-perturbation transient, noise attenuation was quantified over the post-perturbation window beginning 1 h after the input change.

For each iteration, we calculated a dimensionless Gain in Coefficient of Variation:

$$\text{Gain in CoV} = \frac{\text{CoV}_{\text{Output}}}{\text{CoV}_{\text{Input}}},$$

where the input CoV was calculated from the noisy transcriptional input and the output CoV was calculated from mature GFPmut3 over the same post-perturbation window. Lower values therefore indicate greater attenuation of input noise. Under these conditions, the spRNA-input case showed a lower mean Gain in CoV than the sRNA-input case (Supplementary Fig. S53). Specifically, when the noisy transcriptional input was applied to the spRNA, the mean Gain in CoV across the 50 iterations was 0.5873, whereas when the noisy transcriptional input was applied to the sRNA, the mean Gain in CoV was 0.7117. These simulations therefore suggest that, for this parameter set and for our circuit, noisy variation in spRNA transcription is more strongly attenuated at the GFPmut3 output than equivalent noisy variation in sRNA transcription.

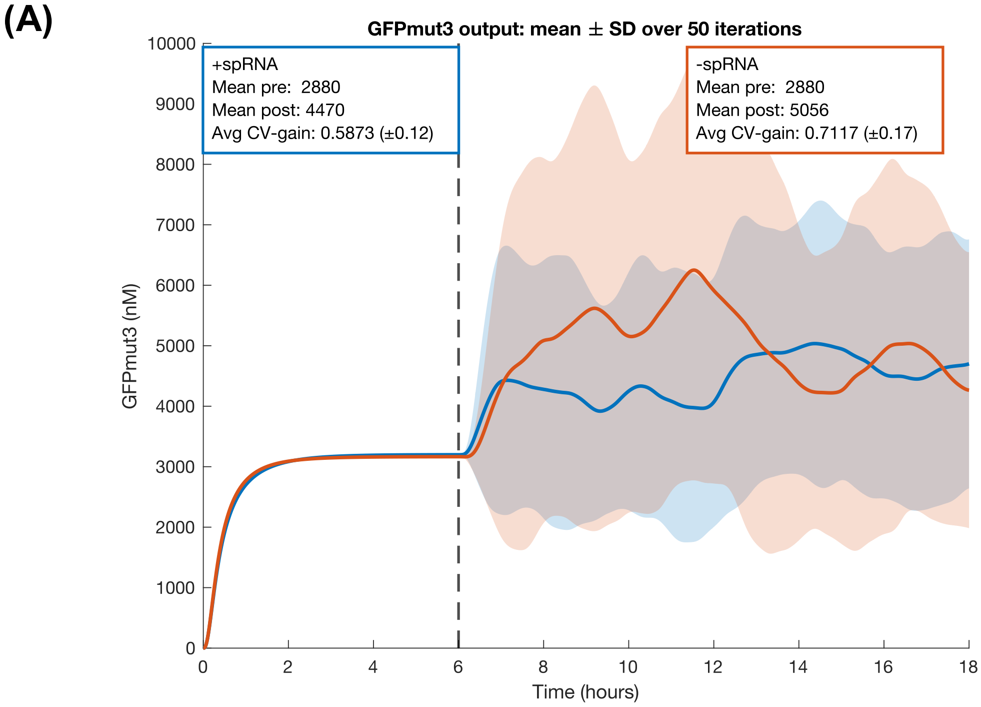


**Figure S53: Iterated noisy-input simulations for estimating disturbance rejection by spRNA-mediated regulation.** For each of 50 iterations, a shared smoothed random noise trace was generated after the perturbation time and applied either to spRNA transcription, with sRNA transcription held constant, or to sRNA transcription, with spRNA transcription set to zero. Mature GFPmut3 trajectories were simulated for each condition and used to calculate a dimensionless Gain in Coefficient of Variation, defined as the output CoV divided by the input CoV over the post-perturbation analysis window. Lower Gain in CoV values indicate stronger attenuation of input noise.

In addition to noisy transcriptional inputs, we also investigated other disturbance types, including spike inputs and square-wave inputs. These simulations again suggested that, in some but not all regimes, spRNAs may improve disturbance rejection (Supplementary Fig. S54). For input spikes, brief transcriptional-input pulses were applied after the system had reached an initial steady state. The same pulse waveform was applied either to spRNA transcription, with sRNA transcription held constant, or to sRNA transcription, with spRNA transcription set to zero. GFPmut3 timecourses were normalised to one at the pre-perturbation time. In our parameter regime, a short transcriptional input pulse ($t = 12.6$ mins) produced no clear change in GFPmut3 expression when applied to sRNA transcription, while the same pulse applied to spRNA transcription produced small changes in GFPmut3 expression (Supplementary Fig. S54A). In contrast, a longer input pulse ($t = 30$ mins) produced a larger transient change in GFPmut3 expression when applied to sRNA transcription, while the same pulse applied to spRNA transcription produced a smaller transient change in GFPmut3 expression (Supplementary Fig. S54B). For our circuit, this suggests that both the duration and frequency of the input pulses are important in determining whether spRNA-mediated regulation can buffer brief transcriptional disturbances.

We then applied square-wave transcriptional inputs to test the response to repeated periodic perturbations. When the same square-wave input was applied to spRNA transcription or to sRNA transcription, the spRNA-input case produced markedly smaller oscillations in mature GFPmut3 (Supplementary Fig. S54C). We next tested the response to square-wave transcriptional perturbations to sRNA transcription in the presence and absence of fixed spRNA transcription. Again, propagation of input disturbances to the GFPmut3 output was reduced in the presence of spRNA (Supplementary Fig. S54D).

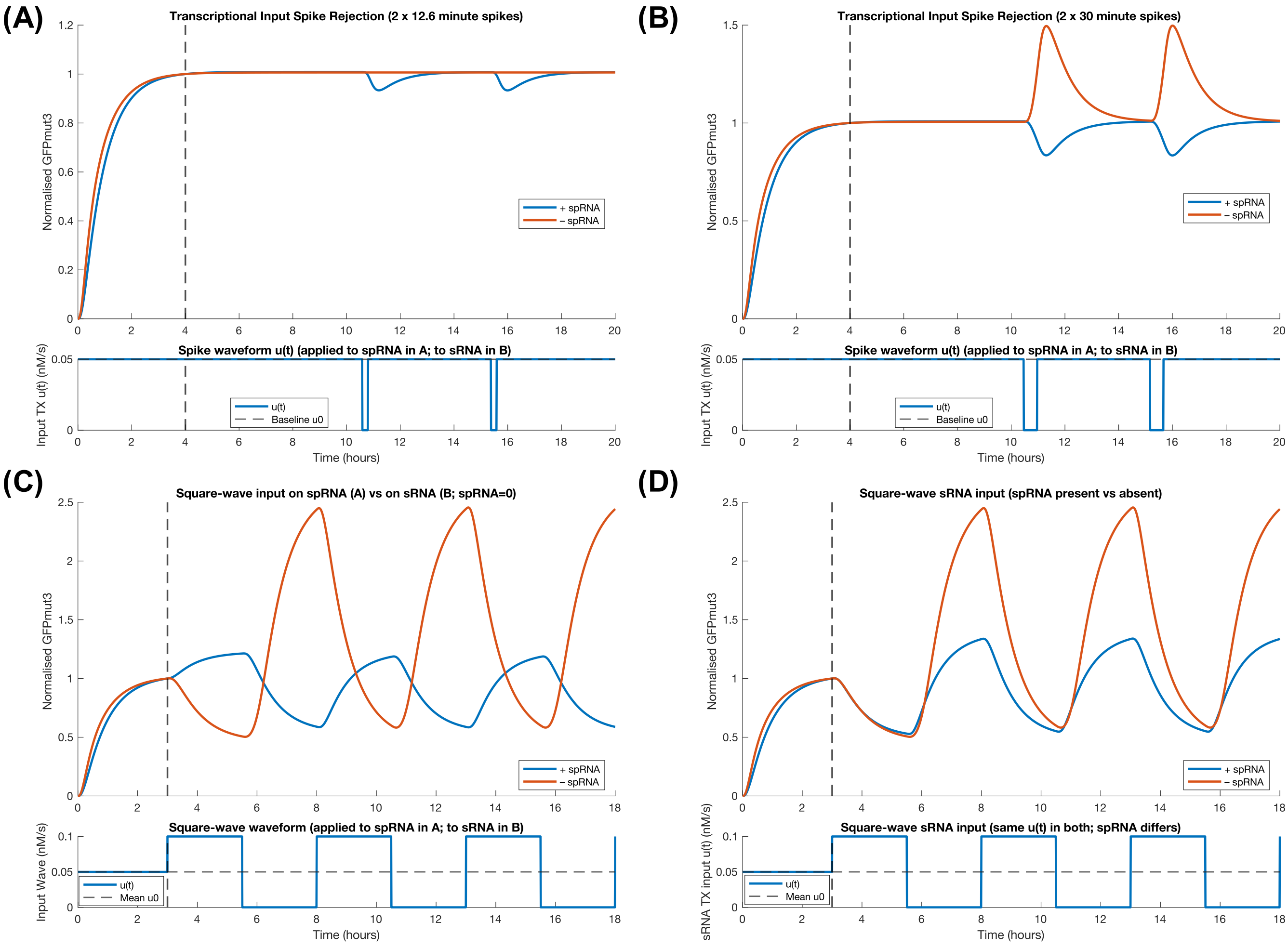


**Figure S54: Additional input-disturbance simulations:** **(A)** Brief transcriptional input spike applied either to sRNA transcription (spRNA transcription = 0) or to spRNA transcription with constant sRNA transcription. For this short pulse, the spRNA-input case produced a small GFPmut3 response, whereas the sRNA-input case produced little change. **(B)** Longer transcriptional input spikes showed the opposite pattern, with lower propagation of the input disturbance to the GFPmut3 output when the pulse was applied to the sponge RNA. **(C)** Square-wave input applied to either spRNA or sRNA transcription. The spRNA-input case produced smaller GFPmut3 output oscillations. **(D)** Square-wave input applied to sRNA transcription in the presence or absence of fixed spRNA transcription, showing reduced input-disturbance propagation when spRNA is present.

#### S9.4 Specificity

As described in the main text, we used simulations to investigate whether sponge RNAs could improve effective specificity when an sRNA represses both an intended on-target and a weaker unintended off-target. These simulations were based on the fitted RNA-interaction model described above (Supplementary Equations (S13)) and the parameter values in Supplementary Tables S9 and S8. Because mScarlet-I protein dynamics were not relevant to this question, these states were omitted and the model was instead extended to include an additional off-target transcript, immature protein, mature protein, and sRNA–off-target complex.

The simulations used the following ODE system:

$$\begin{aligned}
\frac{dM}{dt} &= \alpha_m - (\mu + \delta_m) M - \frac{\gamma_{sm}}{K_{sm}} RMS \\
\frac{dP}{dt} &= \theta M - (\mu + \tau) P \\
\frac{dF}{dt} &= \tau P - \mu F \\
\frac{dS}{dt} &= \alpha_s(t) - (\mu + \delta_s) S - \frac{\gamma_{sm}}{K_{sm}} RSM - \frac{\gamma_{sz}}{K_{sz}} RSZ - \frac{\gamma_{so}}{K_{so}} RSO + n\delta_c C_1 + n\delta_c C_3 \\
\frac{dZ}{dt} &= \alpha_z - (\mu + \delta_z) Z - \frac{\gamma_{sz}}{K_{sz}} RSZ \\
\frac{dC_1}{dt} &= \frac{\gamma_{sm}}{K_{sm}} RSM - (\mu + \delta_c) C_1 \\
\frac{dC_2}{dt} &= \frac{\gamma_{sz}}{K_{sz}} RSZ - (\mu + \delta_c) C_2 \\
\frac{dO}{dt} &= \alpha_o - (\mu + \delta_m) O - \frac{\gamma_{so}}{K_{so}} RSO \\
\frac{dP_o}{dt} &= \theta O - (\mu + \tau) P_o \\
\frac{dF_o}{dt} &= \tau P_o - \mu F_o \\
\frac{dC_3}{dt} &= \frac{\gamma_{so}}{K_{so}} RSO - (\mu + \delta_c) C_3 \,.
\end{aligned} \tag{S19}$$

Here, all species and parameters are as defined in our final model (Supplementary Equations (S13)), with the following additions. $\alpha_m$ and $\alpha_o$ are fixed GFPmut3 and off-target transcription rates, respectively, with $\alpha_m = \alpha_o = 0.1\ \text{nM}\,\text{s}^{-1}$. $\alpha_s(t)$ is the time-dependent transcription rate of the sRNA, which is set to zero before sRNA induction and to $1\ \text{nM}\,\text{s}^{-1}$ after sRNA induction. $\alpha_z$ is the fixed spRNA transcription rate, which is set to $1\ \text{nM}\,\text{s}^{-1}$ in the spRNA-expressing case and to zero in the without-spRNA case. $O$ is the off-target mRNA, $P_o$ is the immature off-target protein, $F_o$ is the mature off-target protein, and $C_3$ is the sRNA–off-target mRNA complex. $\gamma_{so}$ is the newly defined interaction rate between the sRNA and off-target mRNA, which was set to $\gamma_{so} = \gamma_{sm}/10$, while $K_{so}$ is the effective dissociation constant for sRNA–off-target binding, which was set to $K_{so} = 10K_{sm}$.

This analysis makes a number of assumptions: (i) the off-target gene is transcribed, translated, and matured at the same rate as the on-target gene; (ii) the sRNA has the same capacity to act catalytically on both the on-target and off-target mRNAs, and therefore mirrored recycling terms are included for the on-target and off-target complexes ($n\delta_c C_1$ and $n\delta_c C_3$); (iii) the off-target complex is as stable as the on-target complex, such that both are degraded with rate $\delta_c$; and (iv) to model the off-target interaction as a weaker unintended interaction, the sRNA–off-target interaction rate is reduced by a factor of 10 and the effective dissociation constant is increased by a factor of 10 relative to the on-target interaction. This reflects the assumption that the more weakly bound off-target interaction will also compete less effectively for the shared RNA-binding resource. Nevertheless, this off-target interaction still contributes to resource coupling, and so the shared resource pool, $R$, is redefined here as:

$$R = \frac{R_{\text{total}}}{1 + \dfrac{SM}{K_{sm}} + \dfrac{SZ}{K_{sz}} + \dfrac{SO}{K_{so}}} \,.$$

The simulations were run for 12 h, with the sRNA input introduced after 6 h. The total on-target output was plotted as $P + F$, and the total off-target output was plotted as $P_o + F_o$. As outlined in the main text, in the absence of spRNA, induction of the sRNA led to substantial repression of both the intended on-target gene and the weaker unintended off-target gene (Fig. 7L, red curves). In contrast, when the spRNA was expressed, off-target repression was strongly reduced while repression of the intended on-target was largely maintained (Fig. 7L, blue curves).

To further test whether this behaviour could be generalised to other sRNA and spRNA transcription-rate regimes, we tested whether spRNA expression could enable sRNA regulation of both an on-target and an off-target to fall within arbitrary target ranges, for example $\geq 40\%$ repression of the on-target and $\leq 2\%$ repression of the off-target. Using the model and parameters described above, we simulated across a range of post-induction sRNA transcription rates and constitutive spRNA transcription rates. The sRNA transcription rate was varied from 0.01 to 2 $\mathrm{nM\,s^{-1}}$, while the spRNA transcription rate was varied from 0 to 2 $\mathrm{nM\,s^{-1}}$, with the zero-spRNA condition included explicitly as the no-spRNA reference. The on-target and off-target transcription rates were fixed and equal to each other ($\alpha_m = \alpha_o = 0.1$ $\mathrm{nM\,s^{-1}}$). For each pair of sRNA and spRNA transcription rates, the model was first simulated for 8 h without sRNA induction, and then for a further 8 h after sRNA induction. On-target and off-target repression were calculated as the fractional decrease in total protein output after sRNA induction:

$$\text{Repression}_{\text{on}} = \frac{(P+F)_{\text{pre}} - (P+F)_{\text{post}}}{(P+F)_{\text{pre}}}, \qquad \text{Repression}_{\text{off}} = \frac{(P_o+F_o)_{\text{pre}} - (P_o+F_o)_{\text{post}}}{(P_o+F_o)_{\text{pre}}}.$$

Here, the "pre" and "post" values correspond to the final simulated values before and after sRNA induction, respectively. We then produced heat maps showing the sRNA and spRNA transcription-rate regimes in which particular regulatory targets could be achieved. In each case, increasing spRNA transcription expanded the range of sRNA transcription rates able to achieve the desired regulatory target (Supplementary Fig. S55).

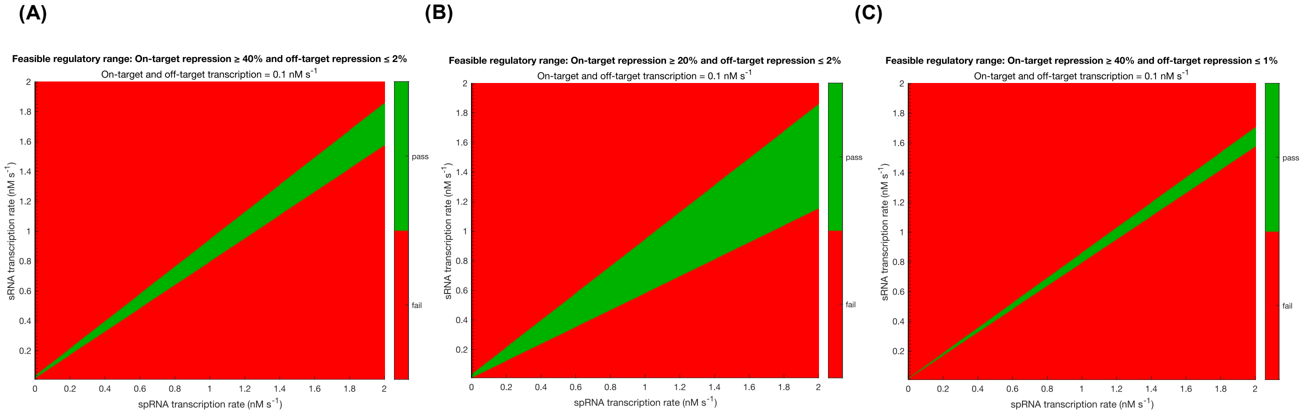


**Figure S55: Heat maps showing sRNA and spRNA transcription-rate regimes in which arbitrary regulatory targets could be achieved.** Feasible regions are shown for **(A)** $\geq 40\%$ repression of the on-target and $\leq 2\%$ repression of the off-target, **(B)** $\geq 20\%$ repression of the on-target and $\leq 2\%$ repression of the off-target, and **(C)** $\geq 40\%$ repression of the on-target and $\leq 1\%$ repression of the off-target. In all cases, increasing spRNA transcription expands the range of sRNA transcription rates that can achieve the regulatory target.

#### S9.5 Evolutionary Stability

As described in the Results section of the main text, we next investigated how circuits would respond to mutations in the sRNA binding region in the presence and absence of spRNA. Using IntaRNA [8], we predicted $\Delta G$ values for sRNA–mRNA and sRNA–spRNA binding for all 164 possible single-nucleotide mutations in the 20-nt Syn-ChiX-20D binding region (60 substitutions, 20 deletions, and 84 insertions). Assuming that effective binding rates are proportional to predicted $\Delta G$ values, we calculated mutant-specific effective binding rates as

$$\gamma_{sm}^{\text{mut}} = \alpha_{sm}\,\Delta G_{sm}^{\text{mut}}, \qquad \gamma_{sz}^{\text{mut}} = \alpha_{sz}\,\Delta G_{sz}^{\text{mut}},$$

where $\alpha_{sm}$ and $\alpha_{sz}$ are constants of proportionality for sRNA–mRNA and sRNA–spRNA interactions, inferred from our fitted values of $\gamma_{sm}$ and $\gamma_{sz}$ for the unmutated Syn-ChiX-20D sequence (Supplementary Table S9). Because the predicted $\Delta G$ values are negative for favourable binding interactions, the inferred proportionality constants are also negative, such that stronger predicted binding gives a larger positive effective interaction rate. The resulting mutant-specific interaction rates are provided in Supplementary Data File 3.

Using these mutant-specific effective binding rates, we ran 164 simulations of our model (Supplementary Equations (S13) and Supplementary Tables S9 and S8) in the presence and absence of spRNA. Parameters were chosen so that the two circuits had similar GFPmut3 expression immediately before mutation. Each system was simulated for 7 h using the fitted wild-type values of $\gamma_{sm}$ and $\gamma_{sz}$, followed by a further 7 h in which $\gamma_{sm}$ and $\gamma_{sz}$ were changed to the mutant-specific values inferred from the IntaRNA predictions and proportionality relationships above. The model output was taken as total GFPmut3 protein, $P + F$, and each trajectory was normalised to its value immediately before the mutation step.

Following mutation, GFPmut3 expression was effectively unchanged in the presence of spRNA, compared with an average $\sim 10\%$ increase in the absence of spRNA (Fig. 7M). Variability across mutants was also substantially larger in the absence of spRNA (Fig. 7M), with $\pm 1\sigma$ around the mean corresponding to approximately 0–20% changes in expression. Representative trajectories for individual mutants (Supplementary Fig. S56) corroborated this, showing substantial heterogeneity in mutational effects on expression in the absence of spRNA, including cases where mutations had minimal impact. In most cases, the presence of spRNA increased the stability of gene expression in response to sRNA mutation, although some mutants showed little change after mutation in both the presence and absence of spRNA (Supplementary Fig. S56D), while others showed the opposite behaviour but typically involved only small changes in expression (Supplementary Fig. S56E).

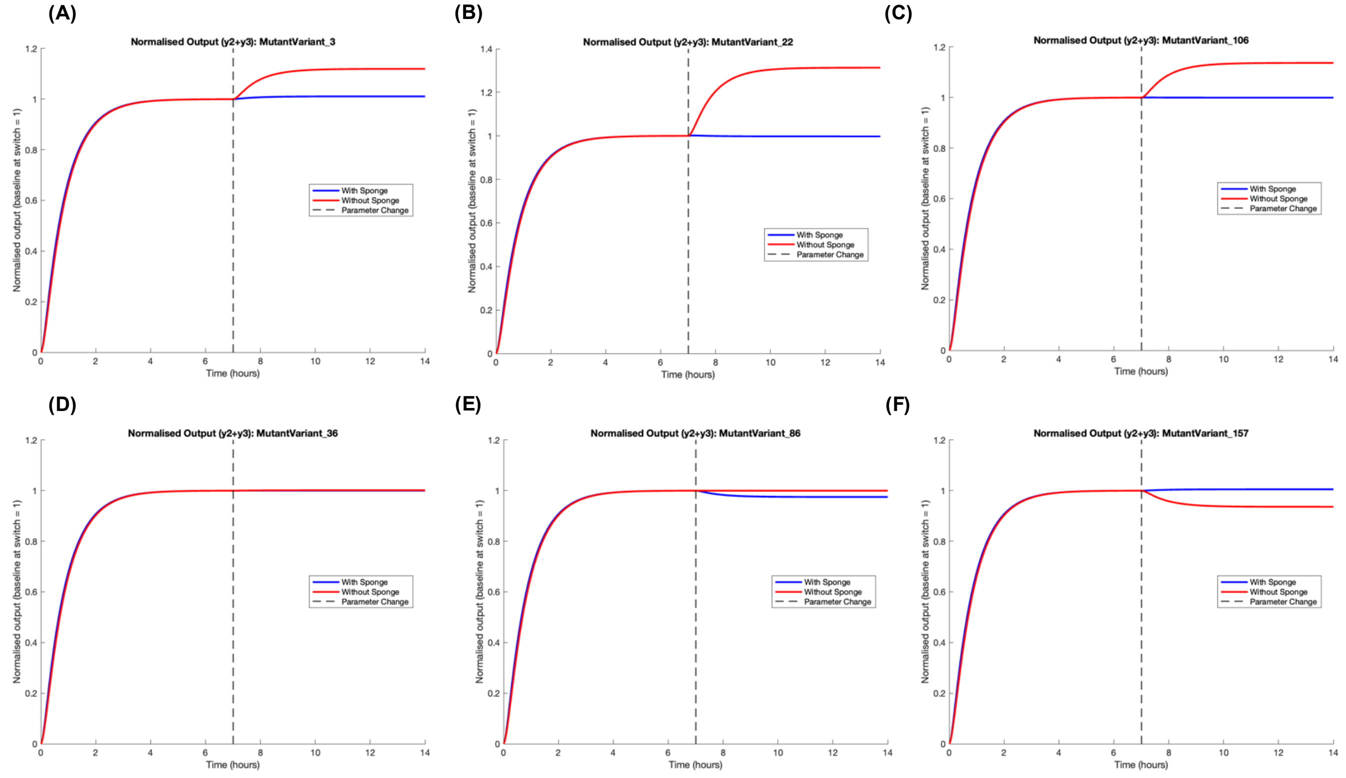


**Figure S56: Example trajectories from the 164 simulated mutant sRNAs in the presence and absence of sponge RNA.** Simulations were run for 7 h using the fitted wild-type sRNA–mRNA and sRNA–spRNA interaction rates, followed by a further 7 h after switching to mutant-specific interaction rates inferred from IntaRNA-predicted binding free energies. GFPmut3 output is shown as total GFPmut3 protein, $P + F$, normalised to the value immediately before the mutation step.

As described in the main text, we also performed a 2D parameter sweep over sRNA and spRNA transcription rates to test whether this mutational buffering was restricted to the specific input conditions used in the initial simulations. In these simulations, GFPmut3 transcription was held fixed at 0.5 $\text{nM}\,\text{s}^{-1}$, while sRNA and spRNA transcription rates were each varied from 0 to 1 $\text{nM}\,\text{s}^{-1}$. For each sRNA–spRNA transcription-rate pair, the model was first simulated for 7 h using the fitted wild-type interaction rates. The sRNA–mRNA and sRNA–spRNA interaction rates were then switched to the mutant-specific values inferred from the IntaRNA predictions, and the model was simulated for a further 7 h for each of the 164 mutant sRNAs.

For each mutant, the total GFPmut3 output was again taken as $P + F$, and the fractional change in expression after mutation was calculated relative to the value immediately before the mutation step:

$$\Delta_{\text{mut}} = \frac{(P+F)_{\text{post}}}{(P+F)_{\text{pre}}} - 1.$$

For each sRNA–spRNA transcription-rate pair, we then calculated the mean value of $\Delta_{\text{mut}}$ across all 164 simulated mutants (Fig. 7N). As spRNA transcription increased, buffering against mutation strengthened, particularly at higher sRNA transcription rates, where the mean normalised change decreased from $\sim 0.25$ in the absence of spRNA to near zero at high spRNA expression. These results suggest that, under certain conditions, the presence of a spRNA in a regulatory network may increase robustness to sRNA mutations and thereby facilitate evolutionary exploration of new sRNA interactions.

##### S9.5.1 Limitations to the evolutionary analysis

There are several important limitations to this analysis. First, the quantitative strength of the conclusion depends on the assumed relationship between predicted binding energy and effective binding rate. Here, we assumed that $\gamma_{sm} = \alpha_{sm}\Delta G_{sm}$ and $\gamma_{sz} = \alpha_{sz}\Delta G_{sz}$, but a more realistic relationship may include an offset, saturation, thresholding, or other non-linearities. Because we have only a single fitted value for each of $\gamma_{sm}$ and $\gamma_{sz}$, we cannot reliably infer a more complex mapping from predicted binding energy to effective interaction rate, and the resulting $\alpha$ values should therefore be interpreted cautiously.

Second, this analysis only considers mutations in the sRNA binding region. In the circuit architecture simulated here, the spRNA can buffer mutations that weaken or alter sRNA binding because the same mutation affects both the sRNA–mRNA and sRNA–spRNA interactions. However, mutations occurring in the spRNA sequence itself would not necessarily have the same effect and could instead reduce spRNA-mediated buffering. More generally, the evolutionary consequences of spRNAs are likely to depend strongly on circuit architecture. For example, in some architectures the sRNA, mRNA target, and spRNA target-recognition regions may be encoded by independent DNA sequences, whereas in others the relevant binding regions may be shared, overlapping, or evolutionarily coupled. A mutation in one genetic element may therefore affect only sRNA binding sites, only mRNA binding sites, only spRNA binding sites, or multiple binding sites simultaneously. These simulations should therefore be interpreted as showing that spRNAs can plausibly buffer one class of mutations in one circuit architecture, rather than as a general statement that spRNAs always increase evolutionary robustness.

#### S10 DNA Sequences and Plasmids

DNA sequences used in this study are provided in Supplementary File 1. All pre-existing biopart sequences (promoters, RBSs, RiboJs, CDSs, terminators, plasmids) that were used in this study were taken from [4] or [3]. Kanamycin resistance cassette sequence was taken from [37]. The CDS sequence for mScarlet-I was taken from [38].

Plasmids used in this study are provided in Supplementary File 2.

#### References


[1] Harrison Steel, Robert Habgood, Ciarán L. Kelly, and Antonis Papachristodoulou. In situ characterisation and manipulation of biological systems with Chi.Bio. *PLOS Biology*, 18(7):e3000794, 2020.

[2] Scott B. Stacey, Ting An Lee, Olivia Gallupova, Eszter Csibra, Antonis Papachristodoulou, Harrison Steel, and Kirill Sechkar. Characterisation of synthetic gene circuits with absolute quantification in continuous culture. In Xiaojun Tian, editor, *In Press*. Springer US, 2025.

[3] Adam J. Meyer, Thomas H. Segall-Shapiro, Emerson Glassey, Jing Zhang, and Christopher A. Voigt. Escherichia coli “Marionette” strains with 12 highly optimized small-molecule sensors. *Nature Chemical Biology*, 15(2):196–204, 2019.

[4] Simon J. Moore, Hung-En Lai, Richard J. R. Kelwick, Soo Mei Chee, David J. Bell, Karen Marie Polizzi, and Paul S. Freemont. EcoFlex: A Multifunctional MoClo Kit for *E. coli* Synthetic Biology. *ACS Synthetic Biology*, 5(10):1059–1069, 2016.

[5] Chunbo Lou, Brynne Stanton, Ying-Ja Chen, Brian Munsky, and Christopher A Voigt. Ribozyme-based insulator parts buffer synthetic circuits from genetic context. *Nature Biotechnology*, 30(11):1137–1142, November 2012.

[6] Pierre Mandin and Susan Gottesman. A genetic approach for finding small RNAs regulators of genes of interest identifies RybC as regulating the DpiA/DpiB two-component system. *Molecular Microbiology*, 72(3):551–565, May 2009.

[7] Anders Aamann Rasmussen, Jesper Johansen, Jesper S. Nielsen, Martin Overgaard, Birgitte Kallipolitis, and Poul Valentin-Hansen. A conserved small RNA promotes silencing of the outer membrane protein YbfM. *Molecular Microbiology*, 72(3):566–577, May 2009.

[8] Martin Mann, Patrick R. Wright, and Rolf Backofen. IntaRNA 2.0: enhanced and customizable prediction of RNA–RNA interactions. *Nucleic Acids Research*, 45(W1):W435–W439, 2017.

[9] Nara Figueroa-Bossi, Martina Valentini, Laurette Malleret, and Lionello Bossi. Caught at its own game: regulatory small RNA inactivated by an inducible transcript mimicking its target. *Genes & Development*, 23:2004–2015, 2009.

[10] Jacqueline Plumbridge, Lionello Bossi, Jacques Oberto, Joseph T. Wade, and Nara Figueroa-Bossi. Interplay of transcriptional and small RNA-dependent control mechanisms regulates chitosugar uptake in *E scherichia coli* and *S almonella*: Coregulation of a sugar porin and PTS transporter. *Molecular Microbiology*, 92(4):648–658, 2014.

[11] Ronny Lorenz, Stephan H. Bernhart, Christian Höner zu Siederdissen, Hakim Tafer, Christoph Flamm, Peter F. Stadler, and Ivo L. Hofacker. ViennaRNA Package 2.0. *Algorithms for Molecular Biology*, 6(1):26, 2011.

[12] Philip Z Johnson and Anne E Simon. RNAcanvas: interactive drawing and exploration of nucleic acid structures. *Nucleic Acids Research*, 51(W1):W501–W508, 2023.

[13] Martin Raden, Syed M Ali, Omer S Alkhnbashi, Anke Busch, Fabrizio Costa, Jason A Davis, Florian Eggenhofer, Rick Gelhausen, Jens Georg, Steffen Heyne, Michael Hiller, Kousik Kundu, Robert Kleinkauf, Steffen C Lott, Mostafa M Mohamed, Alexander Mattheis, Milad Miladi, Andreas S Richter, Sebastian Will, Joachim Wolff, Patrick R Wright, and Rolf Backofen. Freiburg RNA tools: a central online resource for RNA-focused research and teaching. *Nucleic Acids Research*, 46(W1):W25–W29, 2018.

[14] Philip P Adams, Gabriele Baniulyte, Caroline Esnault, Kavya Chegireddy, Navjot Singh, Molly Monge, Ryan K Dale, Gisela Storz, and Joseph T Wade. Regulatory roles of Escherichia coli 5' UTR and ORF-internal RNAs detected by 3' end mapping. *eLife*, 10:e62438, 2021.

[15] Martin Overgaard, Jesper Johansen, Jakob Møller-Jensen, and Poul Valentin-Hansen. Switching off small RNA regulation with trap-mRNA. *Molecular Microbiology*, 73(5):790–800, 2009.

[16] Jiandong Chen, Leann To, Francois de Mets, Xing Luo, Nadim Majdalani, Chin-Hsien Tai, and Susan Gottesman. A fluorescence-based genetic screen reveals diverse mechanisms silencing small RNA signaling in *E. coli*. *PNAS*, 118(27):e2106964118, 2021.

[17] Dokyun Na, Seung Min Yoo, Hannah Chung, Hyegwon Park, Jin Hwan Park, and Sang Yup Lee. Metabolic engineering of Escherichia coli using synthetic small regulatory RNAs. *Nature Biotechnology*, 31(2):170–174, 2013.

[18] Seung Min Yoo, Dokyun Na, and Sang Yup Lee. Design and use of synthetic regulatory small RNAs to control gene expression in Escherichia coli. *Nature Protocols*, 8(9):1694–1707, 2013.

[19] Jinho Yeom, Jong Seong Park, Yong Min Jeon, Beom Seop Song, and Seung Min Yoo. Synthetic fused sRNA for the simultaneous repression of multiple genes. *Applied Microbiology and Biotechnology*, 106(7):2517–2527, 2022.

[20] Eszter Csibra and Guy-Bart Stan. Absolute protein quantification using fluorescence measurements with FPCountR. *Nature Communications*, 13(1):6600, 2022.

[21] Eszter Csibra and Guy-Bart Stan. FPCount protocol - in-lysate (purification free) protocol. 2023.

[22] Talley J. Lambert. FPbase: a community-editable fluorescent protein database. *Nature Methods*, 16(4):277–278, April 2019.

[23] Eszter Csibra. Fpcountr: Fluorescent protein calibration for plate readers. r package, 2021.

[24] Marco Corrao. GOFFREDOpy: probabilistic tracking of growth rate and OD from turbidostat data. Available at https://github.com/marco-corrao/GOFFREDOpy, 2026.

[25] Bin Shao, Jayan Rammohan, Daniel A. Anderson, Nina Alperovich, David Ross, and Christopher A. Voigt. Single-cell measurement of plasmid copy number and promoter activity. *Nature Communications*, 12(1):1475, December 2021.

[26] Jonathan A. Bernstein, Arkady B. Khodursky, Pei-Hsun Lin, Sue Lin-Chao, and Stanley N. Cohen. Global analysis of mRNA decay and abundance in *Escherichia coli* at single-gene resolution using two-color fluorescent DNA microarrays. *Proceedings of the National Academy of Sciences*, 99(15):9697–9702, July 2002.

[27] Kalen P. Clifton, Ethan M. Jones, Sudip Paudel, John P. Marken, Callan E. Monette, Andrew D. Halleran, Lidia Epp, and Margaret S. Saha. The genetic insulator RiboJ increases expression of insulated genes. *Journal of Biological Engineering*, 12(1):23, October 2018.

[28] Lisa Tietze and Rahmi Lale. Importance of the 5 regulatory region to bacterial synthetic biology applications. *Microbial Biotechnology*, 14(6):2291–2315, November 2021.

[29] Scott A Scholz, Chase D Lindeboom, and Lydia Freddolino. Genetic context effects can override canonical *cis* regulatory elements in *Escherichia coli*. *Nucleic Acids Research*, 50(18):10360–10375, October 2022.

[30] Erel Levine, Zhongge Zhang, Thomas Kuhlman, and Terence Hwa. Quantitative Characteristics of Gene Regulation by Small RNA. *PLoS Biology*, 5(9):e229, 2007.

[31] Ciarán L Kelly, Andreas W K Harris, Harrison Steel, Edward J Hancock, John T Heap, and Antonis Papachristodoulou. Synthetic negative feedback circuits using engineered small RNAs. *Nucleic Acids Research*, 46(18):9875–9889, 2018.

[32] Jörg Vogel, Verena Bartels, Thean Hock Tang, Gennady Churakov, Jacoba G. Slagter-Jäger, Alexander Hüttenhofer, and E. Gerhart H. Wagner. RNomics in Escherichia coli detects new sRNA species and indicates parallel transcriptional output in bacteria. *Nucleic Acids Research*, 31(22):6435–6443, November 2003.

[33] Armin M. Zand, Stanislav Anastassov, Timothy Frei, and Mustafa Khammash. Multi-Layer Autocatalytic Feedback Enables Integral Control Amidst Resource Competition and Across Scales. *ACS Synth. Biol.*, 14(4):1041–1061, 2025.

[34] Enrique Balleza, J. Mark Kim, and Philippe Cluzel. Systematic characterization of maturation time of fluorescent proteins in living cells. *Nature Methods*, 15(1):47–51, January 2018.

[35] Kyung Moon and Susan Gottesman. Competition among Hfq-binding small RNAs in Escherichia coli. *Molecular Microbiology*, 82(6):1545–1562, 2011.

[36] Emma L. Denham. The Sponge RNAs of bacteria – How to find them and their role in regulating the post-transcriptional network. *Biochimica et Biophysica Acta (BBA) - Gene Regulatory Mechanisms*, 1863(8):194565, August 2020.

[37] Esteban Martínez-García, Sofía Fraile, Elena Algar, Tomás Aparicio, Elena Velázquez, Belén Calles, Huseyin Tas, Blas Blázquez, Bruno Martín, Clara Prieto, Lucas Sánchez-Sampedro, Morten H H Nørholm, Daniel C Volke, Nicolas T Wirth, Pavel Dvořák, Lorea Alejaldre, Lewis Grozinger, Matthew Crowther, Angel Goñi-Moreno, Pablo I Nikel, Juan Nogales, and Víctor de Lorenzo. SEVA 4.0: an update of the Standard European Vector Architecture database for advanced analysis and programming of bacterial phenotypes. *Nucleic Acids Research*, 51(D1):D1558–D1567, January 2023.

[38] Daphne S. Bindels, Lindsay Haarbosch, Laura van Weeren, Marten Postma, Katrin E. Wiese, Marieke Mastop, Sylvain Aumonier, Guillaume Gotthard, Antoine Royant, Mark A. Hink, and Theodorus W. J. Gadella. mScarlet: a bright monomeric red fluorescent protein for cellular imaging. *Nature Methods*, 14(1):53–56, January 2017.
